# Towards a comprehensive anatomical matrix for crown birds: phylogenetic insights from the pectoral girdle and forelimb skeleton

**DOI:** 10.1101/2025.01.17.633553

**Authors:** Albert Chen, Elizabeth M. Steell, Roger B. J. Benson, Daniel J. Field

## Abstract

Phylogenetic analyses of phenotypic characters in crown-group birds often recover results that are strongly incongruous with the findings of recent phylogenomic analyses. Furthermore, existing morphological datasets for crown birds are frequently limited by restricted taxon or character sampling, inconsistent character construction, incorrect scoring, or a combination of several of these factors. As part of an effort to address these limitations, in this study we focus on identifying phylogenetically informative characters of the avian pectoral girdle and forelimb skeleton, elements of which are commonly preserved as avian fossils. We assembled and vetted a dataset of 204 characters, which were then scored for a phylogenetically diverse range of 75 extant avian taxa and incorporated into phylogenetic analyses. Analyses run without topological constraints exhibited notable conflicts with the results of recent phylogenomic studies, possibly due to functional convergence and rapid cladogenesis in the early evolutionary history of crown birds. Qualitative anatomical comparisons and quantitative metrics of homoplasy further highlighted the fact that similar morphologies in pectoral girdle and forelimb elements have evolved repeatedly in distantly related groups of birds, representing a major confounding factor in avian morphological phylogenetics. However, the implementation of molecular scaffolds allowed identification of diagnostic character combinations for numerous avian clades previously only recognized through molecular data, such as Phaethontimorphae, Aequornithes, and Telluraves. Although large morphological datasets may not guarantee increased congruence with molecular phylogenetic studies, they can nonetheless be valuable tools for identifying anatomical synapomorphies of key clades, placing fossils into phylogenetic context, and studying macroevolutionary patterns within major groups of organisms.

## Introduction

Recent large-scale phylogenomic studies have greatly clarified the phylogenetic interrelationships of crown birds (Jarvis et al., 2014; Prum et al., 2015; Reddy et al., 2017; Kuhl et al., 2021; Stiller et al., 2024; S. Wu et al., 2024). However, areas of contention remain, particularly concerning the branching topology at the base of Neoaves, a clade encompassing over 95% of extant bird diversity (Reddy et al., 2017; Braun et al., 2019; Braun and Kimball, 2021; Bravo et al., 2021; Gatesy and Springer, 2022; N. Wang et al., 2022; Stiller et al., 2024; S. Wu et al., 2024). Furthermore, analyses of existing avian anatomical datasets (e.g., Mayr and Clarke, 2003; Livezey and Zusi, 2007; Worthy et al., 2017; Musser and Cracraft, 2019) frequently fail to recover well-supported clades found by phylogenomic studies. As morphological data are essential for assessing the affinities of most extinct taxa and identifying suitable fossil calibrations for estimating divergence times (Giribet, 2015; Lee and Palci, 2015), this incongruence between molecular and morphological phylogenetic analyses severely hampers our understanding of morphological evolution in the early history of crown birds.

Several outstanding shortcomings of existing avian morphological phylogenetic datasets have been identified. The largest such dataset to date, presented in Livezey and Zusi (2006), has been criticized for its limited sampling of fossil taxa (Mayr, 2007a) and incorrect or doubtful character scorings (Mayr, 2008a). Furthermore, most morphological datasets for crown birds are relatively limited in anatomical and phylogenetic scope, often incorporating fewer than 200 characters or targeted towards resolving the affinities of narrow avian subgroups instead of broader clades. Although Mayr (2008a) also interpreted the results of Livezey and Zusi (2007) as evidence that large morphological datasets potentially contain little phylogenetic signal, this contention has not been tested for crown bird phylogenetics using independent datasets constructed with the explicit aim of correcting the shortcomings of previous studies. In any case, the production of morphological datasets that encompass a truly representative sample of crown bird character and taxic diversity remains underexplored.

Despite the frequently conflicting phylogenetic signals arising from molecular and morphological sources of data, some recent research has suggested that analyses of avian osteological data can potentially produce more congruent phylogenetic topologies with molecular analyses than those of soft tissue characters (Sansom and Wills, 2017; Callender-Crowe and Sansom, 2022; though see Ericson and Qu, 2024). Furthermore, past reassessments of avian morphological data considering strongly supported clades recovered by molecular analyses have often been successful in identifying previously overlooked character support for said groupings (e.g., Mayr, 2004a; Manegold, 2006; Mayr, 2011a; Chen et al., 2019; Mayr, 2019a; McInerney et al., 2019; Steell et al., 2023a). The production of a comprehensive, improved morphological phylogenetic dataset for crown birds could therefore provide a robust independent avenue with which to assess avian phylogenetic relationships, as well as facilitate congruence between morphological and molecular sources of phylogenetic information.

As part of an initiative to assemble such a dataset, we present a new morphological dataset for crown birds focusing on skeletal characters from the pectoral girdle and forelimb. Long recognized as critical to understanding the evolutionary origins of the modern avian bauplan (e.g., Ostrom, 1974; Jenkins, 1993; Ostrom, 1995; Middleton and Gatesy, 2000; Gishlick, 2001; Senter, 2006; Dececchi and Larsson, 2009; Nesbitt et al., 2009; Sullivan et al., 2010; Heers and Dial, 2012; Zheng et al., 2012; Benson and Choiniere, 2013; Botelho et al., 2014; Xu et al., 2014; Zheng et al., 2014; O’Connor et al., 2015; Heers, 2016; Heers et al., 2016; Mayr, 2017a; Nebreda et al., 2020; Novas et al., 2020; Cau et al., 2021; Heers et al., 2021; Karoullas and Nudds, 2021; Novas et al., 2021; Lo Coco et al., 2022; Pittman et al., 2022; S. Wang et al., 2022; Eliason et al., 2023; Lowi-Merri et al., 2023; Wang and Zhou, 2023; Widrig et al., 2023; Q. Wu et al., 2024a; Q. Wu et al., 2024b), this anatomical region is of particular interest in avian morphological phylogenetics for several reasons. For one, though the fossil record of birds is often regarded as relatively sparse, some elements of the avian pectoral girdle and forelimb, such as the coracoid, are commonly recovered in fossil assemblages that preserve remains of birds and their close relatives (e.g., Longrich, 2009; Longrich et al., 2011). In addition, several elements from this anatomical region have been assessed for phylogenetic signal in previous ornithological studies (Fürbringer, 1888; Höfling and Alvarenga, 2001; Mayr, 2014a; Wang and Clarke, 2014; Shatkovska and Ghazali, 2017; Steell et al., 2023a; Shimizu and Anezaki, 2024), and characters from the pectoral girdle and forelimb account for over 30% of the characters used in some phylogenetic datasets focused on crown birds and near-crown stem-birds (Brocklehurst et al., 2012).

## Methods and Materials

### Phylogenetic dataset

We assembled a phylogenetic dataset of 204 morphological characters from the pectoral girdle and forelimb skeleton. Characters were sourced from direct anatomical observations as well as numerous previous studies (primarily Mayr and Clarke, 2003; Livezey and Zusi, 2006; Smith, 2010; Worthy et al., 2017; Ksepka et al., 2019; Musser and Cracraft, 2019; and Musser and Clarke, 2020), and modified based on guidelines for character construction put forth by Hawkins et al. (1997), Sereno (2007), and Simões et al. (2017). Characters from nearly all bony elements of the avian pectoral girdle and forelimb were included, with the exception of the smaller manual phalanges, which are frequently lost in fossil specimens and prepared skeletons of extant species. Complete character descriptions are available as Supplementary Material Appendix S1.

Our dataset includes 77 taxa, including the Cretaceous stem-birds *Gansus* and *Ichthyornis* as well as 75 extant species broadly sampling crown bird diversity (Fig. 1). Only taxa that use the forelimbs in propulsive locomotion (powered aerial flight or wing-propelled diving) were considered for this study, as the convergent reduction of pectoral and forelimb elements in flightless birds has well-known confounding effects on phylogenetic reconstruction (Worthy and Scofield, 2012; Worthy et al., 2017). Taxa were scored primarily based on personal examination of computed-tomography (CT) scans published by Bjarnason and Benson (2021) and Benito et al. (2022a), with the exception of *Gansus*, which was scored based on published fossil descriptions (You et al., 2006; Li et al., 2011; Wang et al., 2016). A complete list of specimens examined is provided in Supplementary Material Appendix S3. Homology of structures in the highly modified humerus and antebrachial bones of *Spheniscus* with those of other birds was inferred based on muscular arrangements described by Watanabe et al. (2021). For some extant taxa, the unfused carpals and/or manual phalanges were absent from the specimens we examined and the characters pertaining to these elements were scored as unknown. Our complete phylogenetic matrix is available as Supplementary Material Appendix S4.

**Figure 1.**
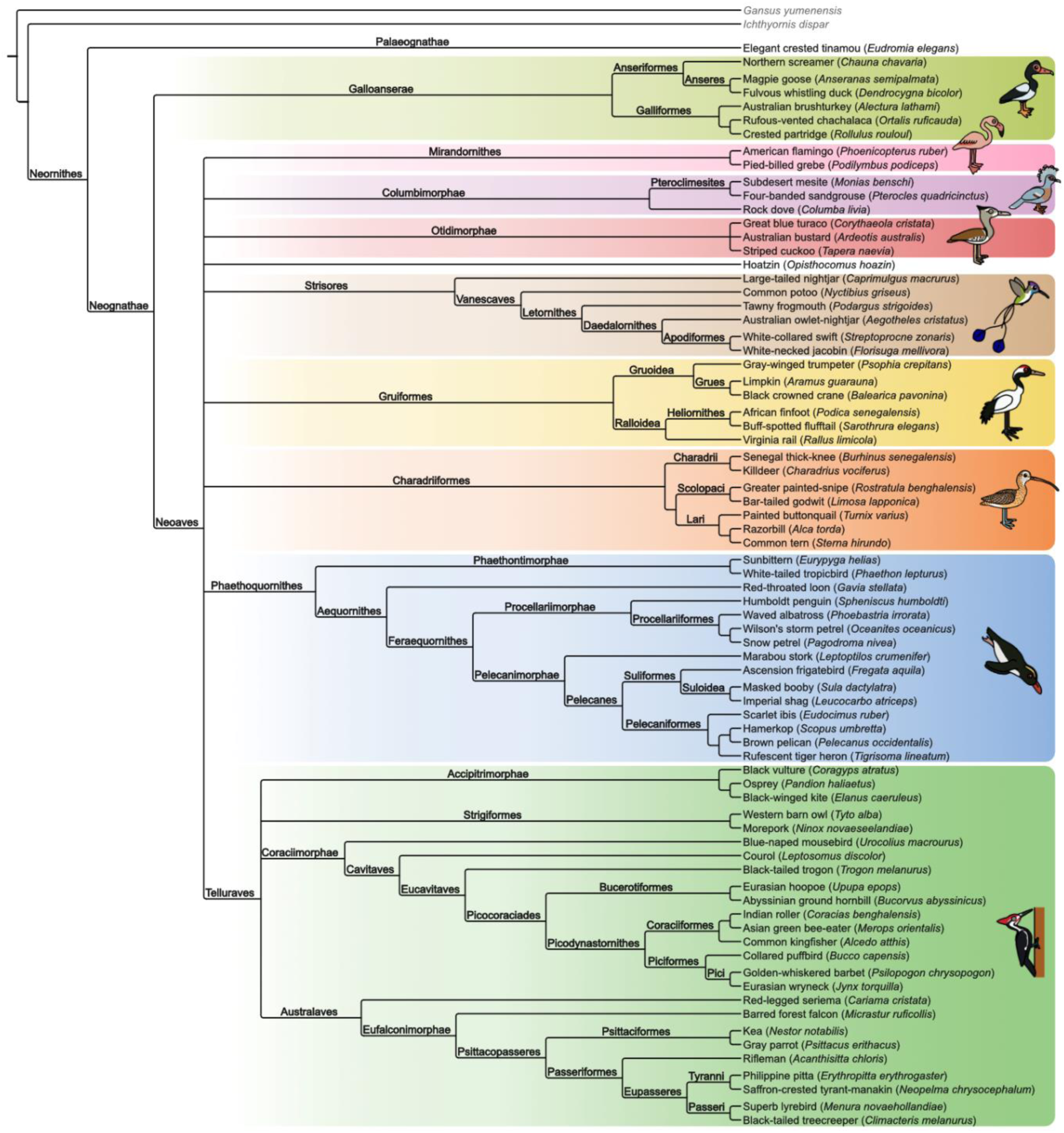
Phylogenetic consensus tree of the species sampled in this study, summarized based on recent phylogenomic studies (Jarvis et al., 2014; Prum et al., 2015; Kuhl et al., 2021; Stiller et al., 2024; S. Wu et al., 2024) and labeled with names of higher-order clades. English names largely follow Gill et al. (2024), though the name “courol” is used for *Leptosomus discolor* instead of “cuckoo-roller” to emphasize its phylogenetic distinctiveness from true rollers (Coraciidae). Illustrations depict representatives of major clades, but not necessarily the precise species examined in the present study.

Anatomical terminology follows the English equivalents of terms used by Baumel and Witmer (1993), with some exceptions as follows. “Lateral trabeculae of the sternum” is used here to indicate the apomorphic, laterally projecting sternal processes found in galliforms instead of the more widely distributed caudolaterally-projecting processes, which are instead called “caudolateral trabeculae” here, following Livezey and Zusi (2006). The radial carpal (“radiale”) and ulnar carpal (“ulnare”) are referred to here as the “scapholunare” and “pisiform” respectively, following the recommendations of Botelho et al. (2014). Anatomical features referenced within this study are illustrated in Figs. 2–6.

**Figure 2.**
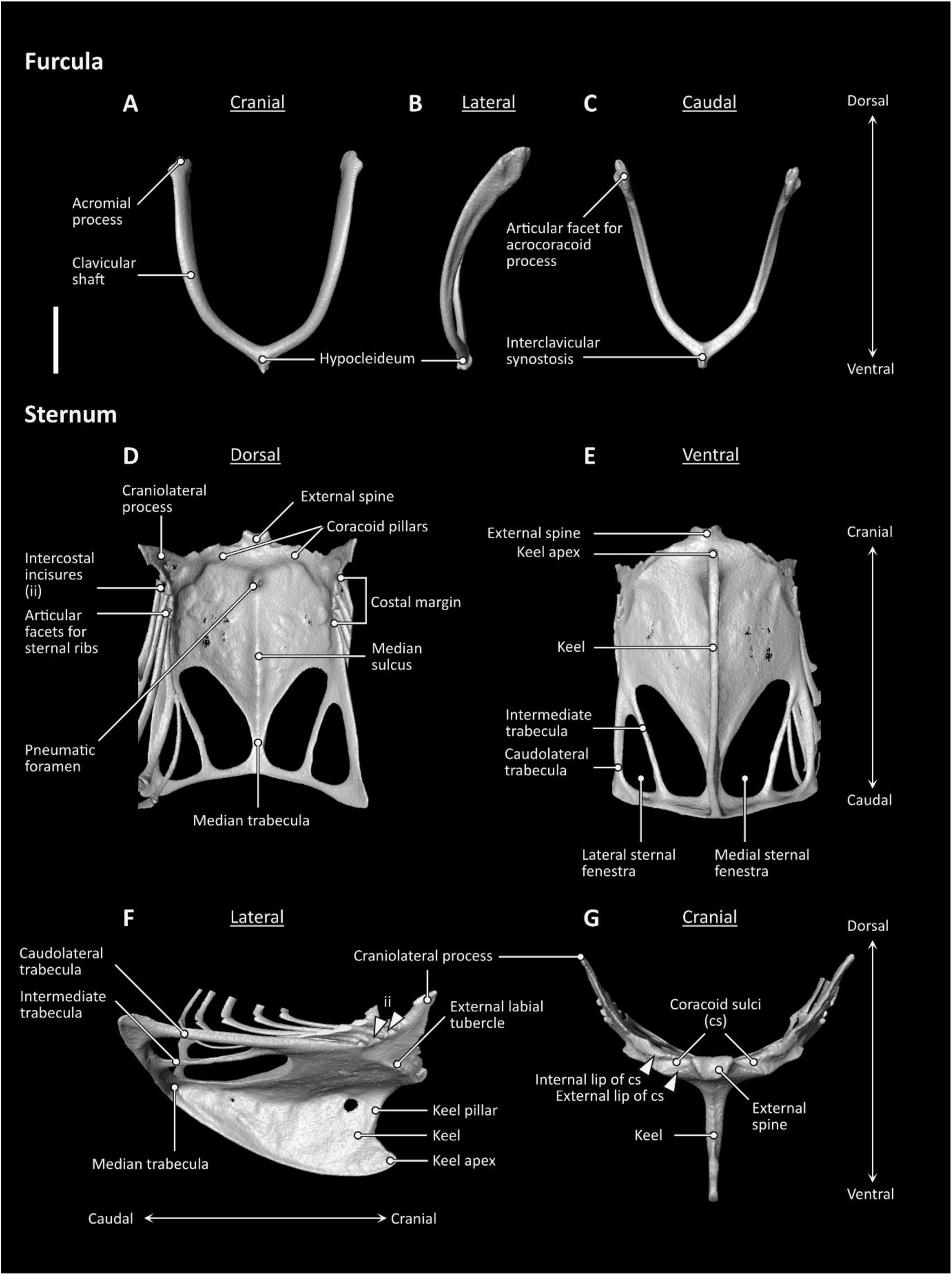
Furcula (**A–C**) and sternum (**D–G**) anatomy of *Aegotheles cristatus* (Australian owlet-nightjar, YPM 124258). Scale bar = 5 mm.

**Figure 3.**
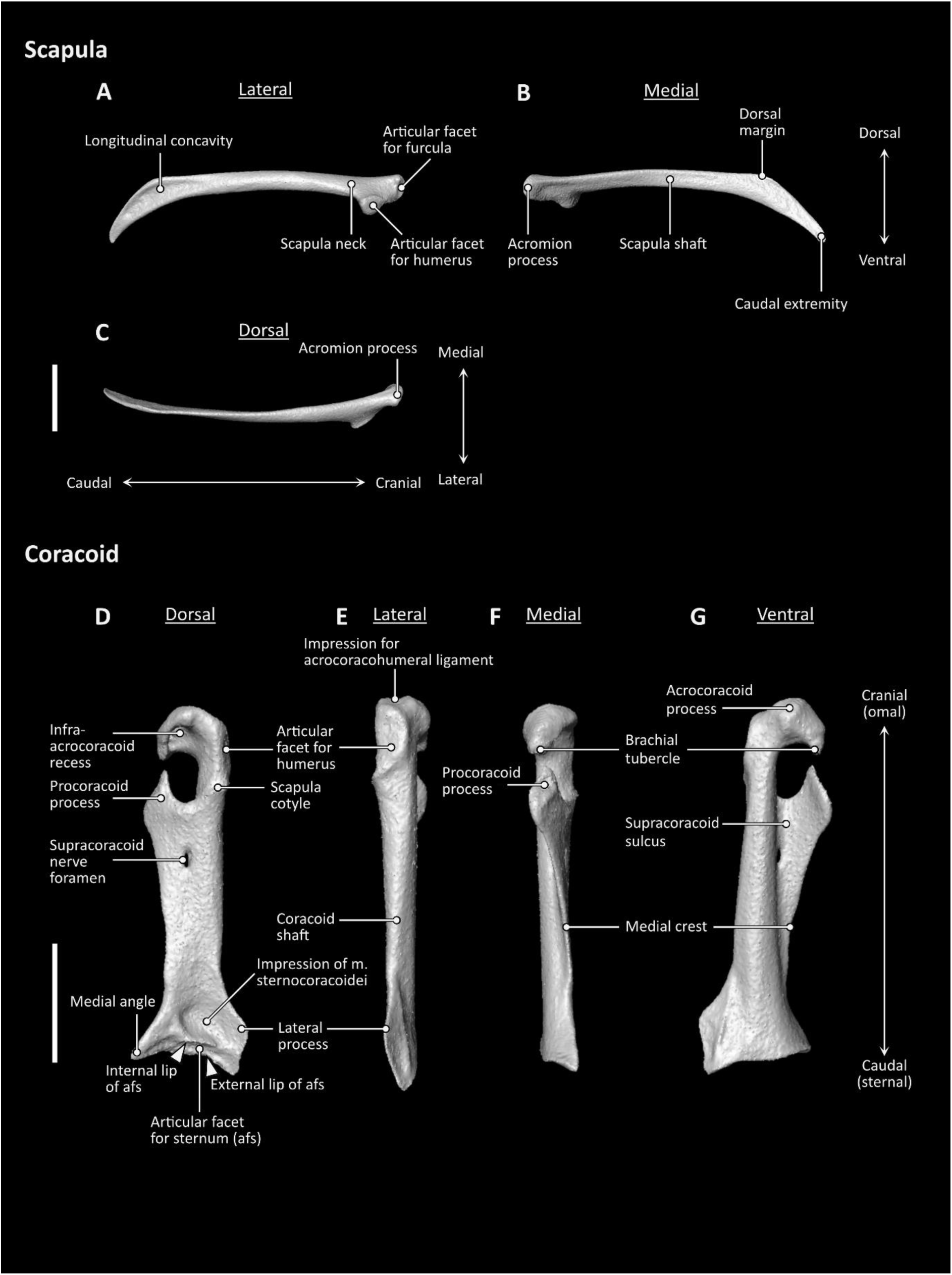
Scapula (**A–C**) and coracoid (**D–G**) anatomy of *Aegotheles cristatus* (Australian owlet-nightjar, YPM 124258). Scale bars = 5 mm.

**Figure 4.**
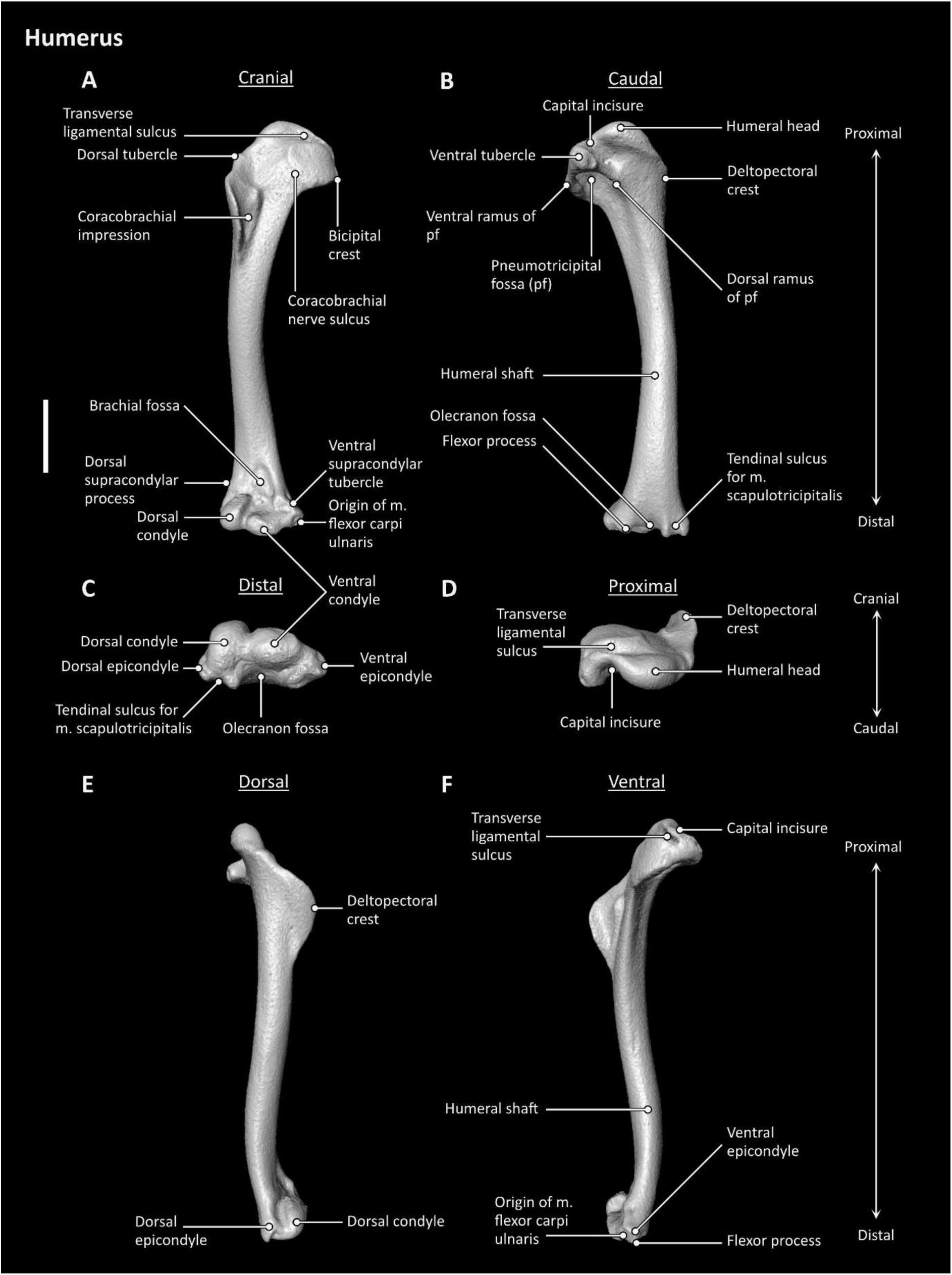
Humerus anatomy of *Aegotheles cristatus* (Australian owlet-nightjar, YPM 124258). Scale bar = 5 mm.

**Figure 5.**
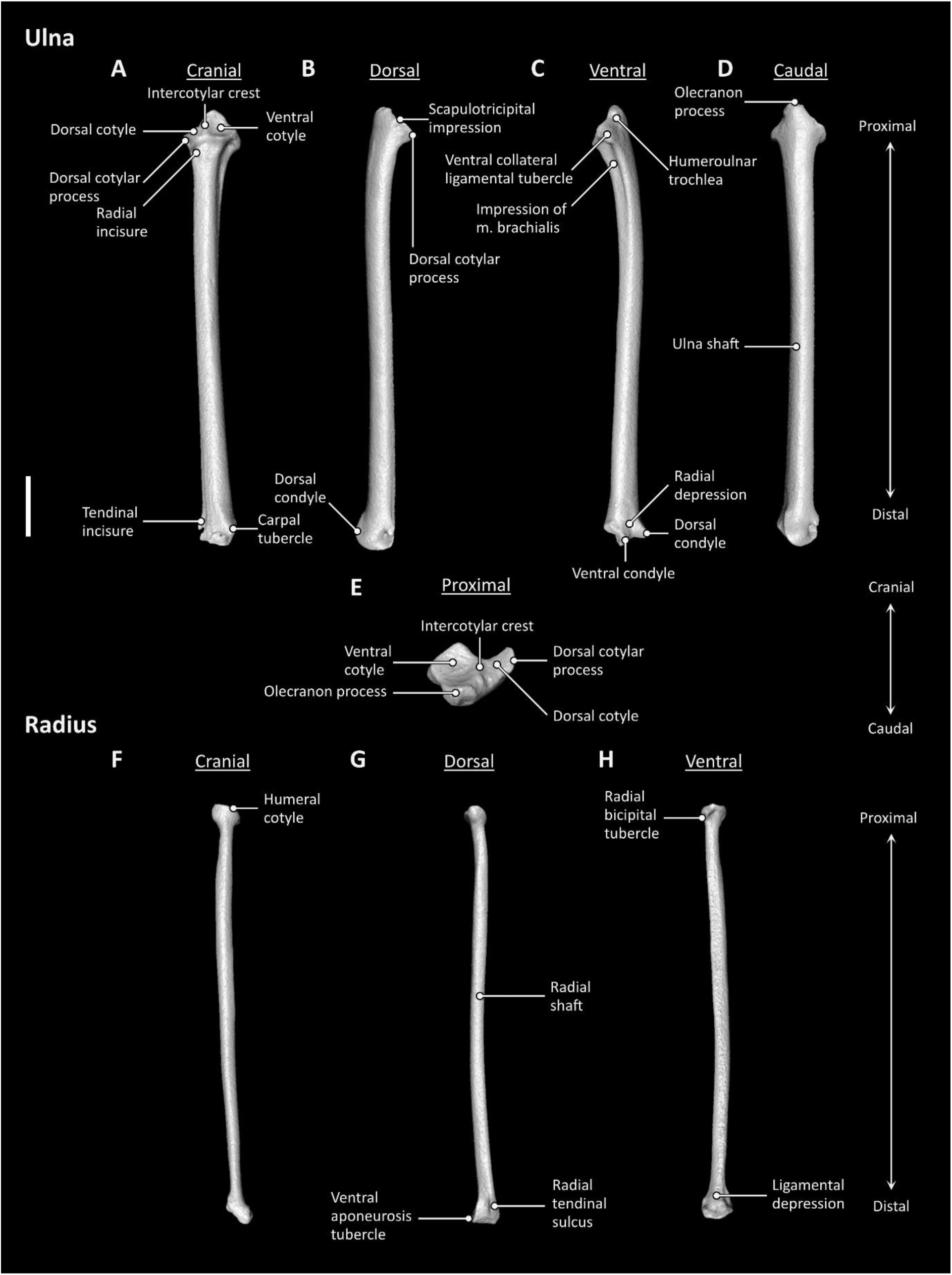
Ulna (**A–D**) and radius (**F–H**) anatomy of *Aegotheles cristatus* (Australian owlet-nightjar, YPM 124258). Scale bar = 5 mm.

**Figure 6.**
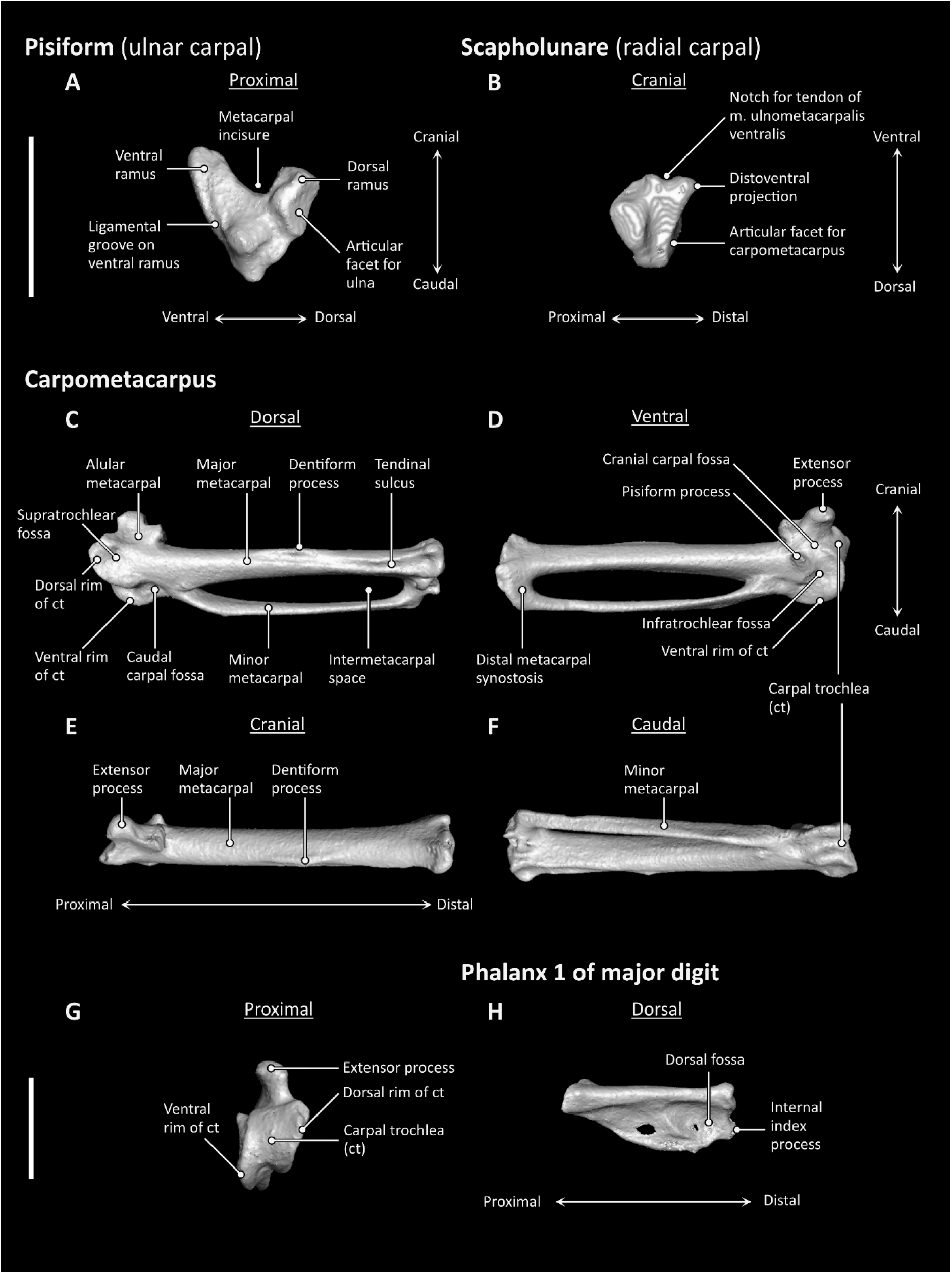
Pisiform (**A**), scapholunare (**B**), carpometacarpus (**C–G**), and manual phalanx II-1 (**H**) anatomy of *Aegotheles cristatus* (Australian owlet-nightjar, YPM 124258). Scale bars = 5 mm.

Where applicable, taxon names for major bird clades follow the phylogenetic definitions of Chen and Field (2020), Mindell (2020), Sangster (2020a), Sangster (2020b), Sangster and Mayr (2021), and Sangster et al. (2022). A phylogenetic consensus tree and nomenclatural framework for the taxa included in this study are shown in Fig. 1.

### Phylogenetic analyses

Twelve different analyses were run under alternative topological constraints and search parameters, as described in Table 1. Details of topological constraints are visualized in Fig. 7.

**Table 1.**
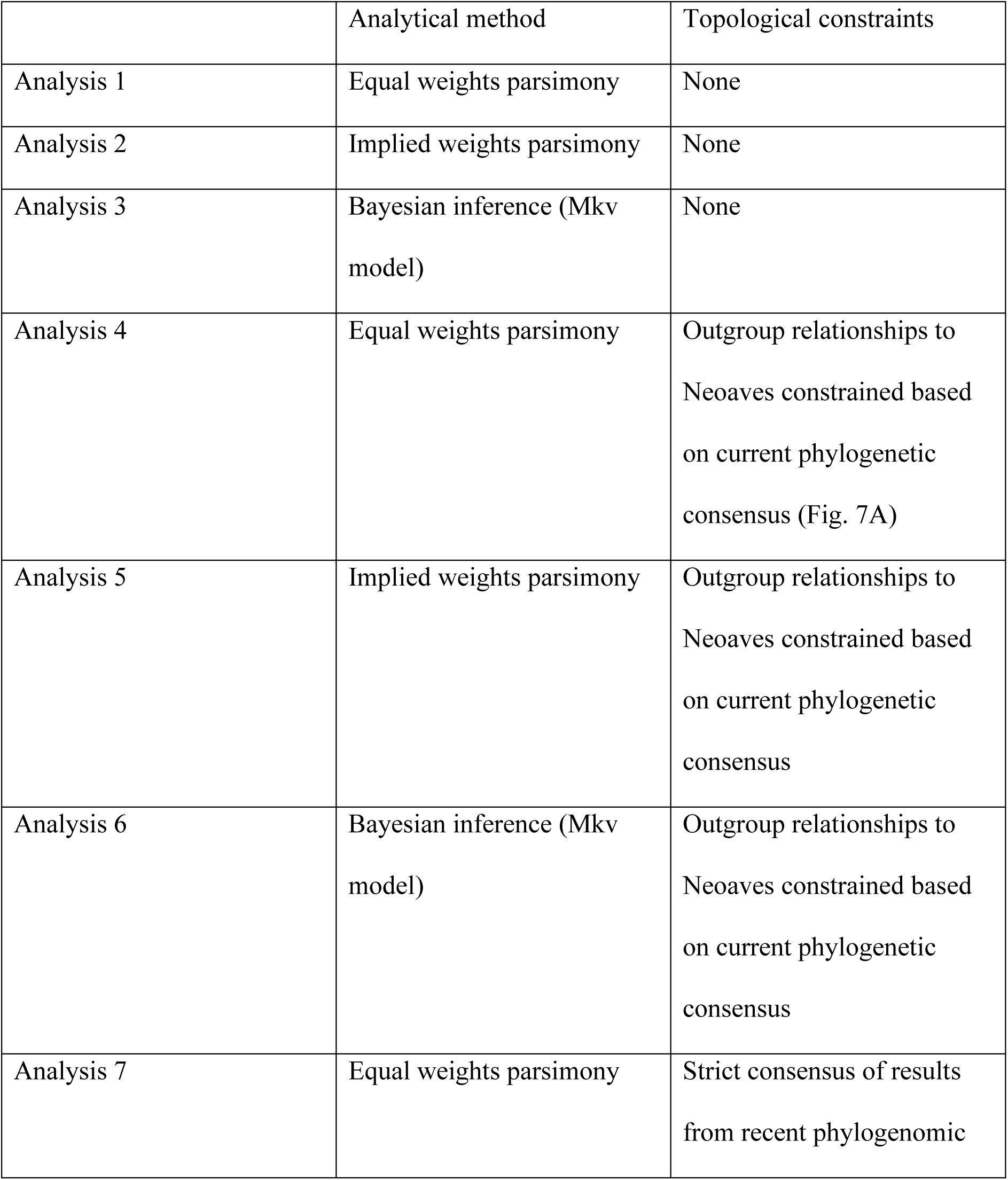
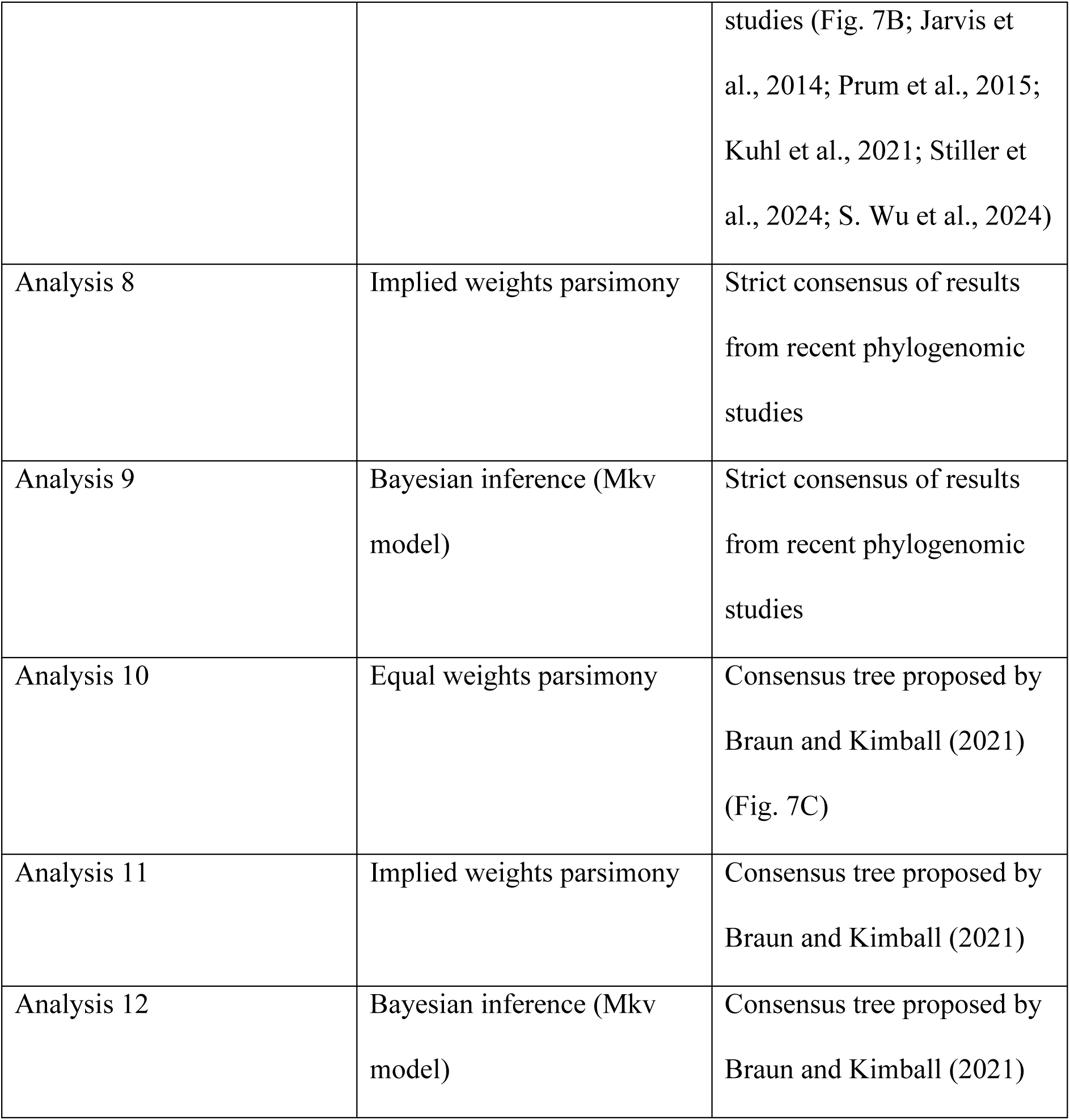
Parameters for phylogenetic analyses run in the present study.

|  | Analytical method | Topological constraints |
| --- | --- | --- |
| Analysis 1 | Equal weights parsimony | None |
| Analysis 2 | Implied weights parsimony | None |
| Analysis 3 | Bayesian inference (Mkv model) | None |
| Analysis 4 | Equal weights parsimony | Outgroup relationships to Neoaves constrained based on current phylogenetic consensus (Fig. 7A) |
| Analysis 5 | Implied weights parsimony | Outgroup relationships to Neoaves constrained based on current phylogenetic consensus |
| Analysis 6 | Bayesian inference (Mkv model) | Outgroup relationships to Neoaves constrained based on current phylogenetic consensus |
| Analysis 7 | Equal weights parsimony | Strict consensus of results from recent phylogenomic |
|  |  | studies (Fig. 7B; Jarvis et al., 2014; Prum et al., 2015; Kuhl et al., 2021; Stiller et al., 2024; S. Wu et al., 2024) |
| Analysis 8 | Implied weights parsimony | Strict consensus of results from recent phylogenomic studies |
| Analysis 9 | Bayesian inference (Mkv model) | Strict consensus of results from recent phylogenomic studies |
| Analysis 10 | Equal weights parsimony | Consensus tree proposed by Braun and Kimball (2021) (Fig. 7C) |
| Analysis 11 | Implied weights parsimony | Consensus tree proposed by Braun and Kimball (2021) |
| Analysis 12 | Bayesian inference (Mkv model) | Consensus tree proposed by Braun and Kimball (2021) |

**Figure 7.**
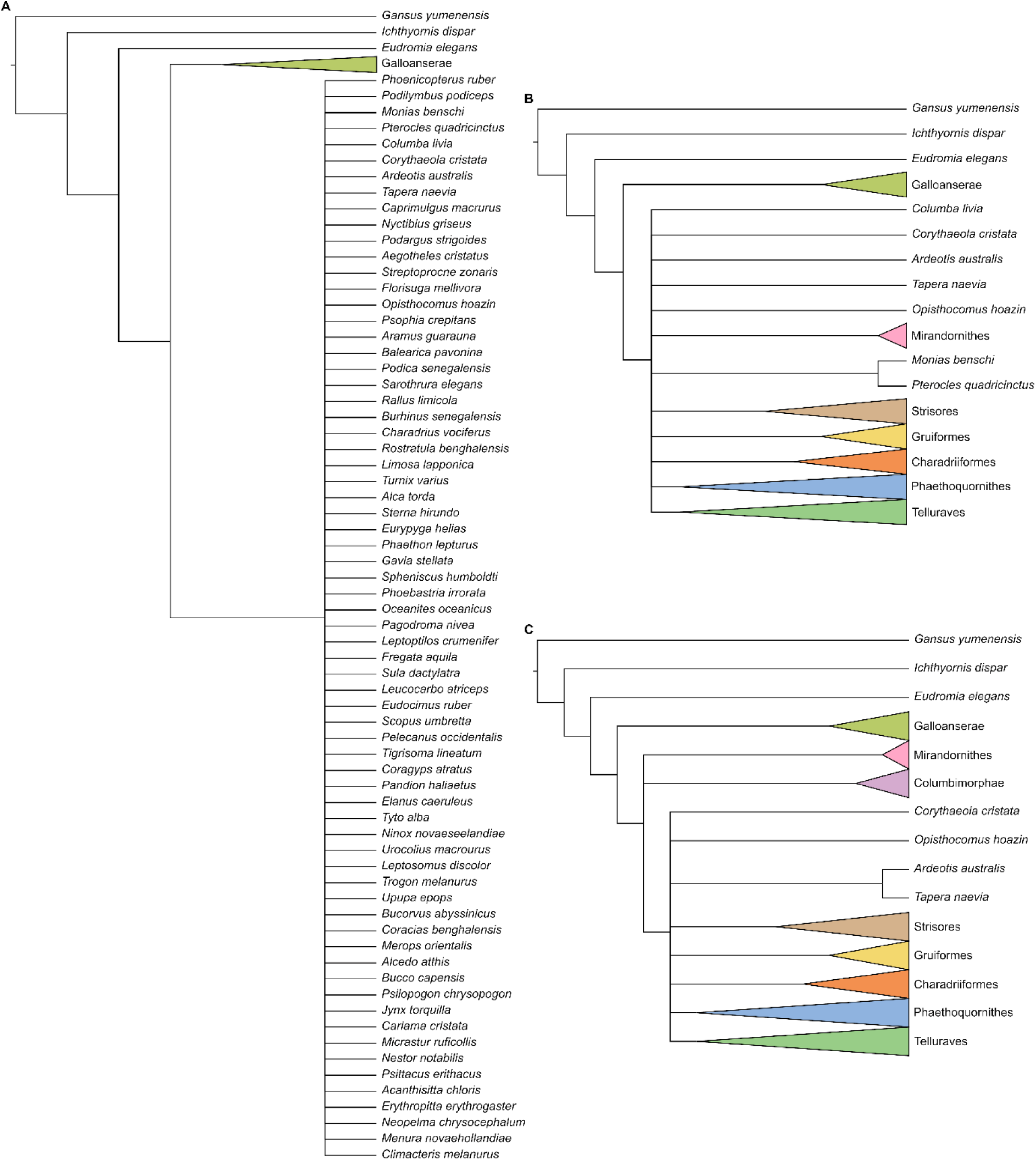
Topological constraints used for Analyses 4–6 (**A**), Analyses 7–9 (**B**), and Analyses 10–12 (**C**). Where constrained to be monophyletic, select major avian clades are collapsed and highlighted following the color coding shown in Fig. 1. Internal relationships of collapsed groups can be assumed to follow those shown in Fig. 1. Further details of analytical parameters are discussed in the main text.

Maximum parsimony analyses were conducted in TNT v.1.5 (Goloboff and Catalano, 2016). After increasing the maximum number of trees to 99,999, a new technology search was run in which a minimum length tree was found in 1000 replicates and default parameters were set for sectorial search, ratchet, tree drift and tree fusion. Implied weights analyses were run with a *k* (concavity constant) value of 12, following Goloboff et al. (2018). Absolute bootstrap frequencies were obtained from 1,000 replicates under a traditional search with default parameters.

Bayesian phylogenetic analyses were conducted in MrBayes (Ronquist et al., 2012) using the CIPRES Science Gateway (Miller et al., 2010). Data were analysed under the Mkv model (Lewis, 2001), and gamma-distributed rate variation was assumed to allow for evolutionary rate heterogeneity across characters. Analyses were conducted using four chains and two independent runs for 30,000,000 generations, with a tree sampled every 4,000 generations and a burn-in of 25%. Analytical convergence was assessed using standard diagnostics provided in MrBayes (average standard deviation of split frequencies < 0.01, potential scale reduction factors = 1, effective sample sizes > 200). Results of independent runs of the same analyses were summarized using the sump and sumt commands. To facilitate comparisons with fully resolved topologies, the “contype = allcompat” command was used to display clades that were present in less than 50% of post-burnin trees but were compatible with the 50% majority-rule consensus tree.

### Comparisons with molecular topologies

As a measure of tree distance, the Robinson–Foulds (RF) index comparing the most parsimonious trees and the “allcompat” consensus trees recovered by analyses of the morphological dataset with the molecular phylogenetic results of Jarvis et al. (2014), Prum et al. (2015), Kuhl et al. (2021), Stiller et al. (2024), and S. Wu et al. (2024) were calculated in R 4.3.1 (R Core Team, 2023) using the package phangorn (Schliep, 2011). The data and R code used for calculating RF indices is available as Supplementary Material Appendix S5. Synapomorphies of major clades recovered by molecular phylogenetic analyses were inferred under maximum parsimony by mapping the morphological dataset onto the aforementioned molecular topologies in TNT and ASADO v.1.61 (Nixon, 2002). Tree lengths based on the inferred number of morphological changes under molecular topologies were also calculated in TNT.

To visualize the structure of the morphological dataset relative to molecular topologies, heatmaps of similarity matrices were produced from our phylogenetic matrix following the procedure outlined by Evers and Benson (2019). Similarity matrices were calculated for the complete morphological dataset as well as partitions of that dataset pertaining to individual elements for which over ten characters were coded, and colors were assigned to the similarity values of each matrix using the package gclus (Hurley, 2019). *Monias benschi* was excluded from the similarity matrix calculated for furcular characters due to its lack of an ossified furcula. The data and R code for calculating similarity and producing the heatmaps is available as Supplementary Material Appendix S6.

### Measuring homoplasy

We applied parsimony-based indices (consistency index [Kluge & Farris, 1969], retention index [Farris, 1989] and relative homoplasy index [Steell et al., 2023b]) to quantify homoplasy in the present dataset with the same molecular constraint topologies as above. For comparison, we also estimated homoplasy in additional published extant bird matrices covering several taxonomic scales: Mayr and Clarke (2003), Livezey and Zusi (2007), Field et al. (2020), and Musser and Clarke (2020) representing general Neornithes matrices; Ksepka et al. (2023a) representing Galliformes; Chen et al. (2019) representing Strisores; Ksepka et al. (2023b) representing Sphenisciformes; Ksepka et al. (2019) representing Telluraves; and Steell et al. (2023a) representing Passeriformes. Recent molecular topological constraints were applied to each matrix, and when applicable, multiple constraints were applied based on the differing topologies of Jarvis et al. (2014), Prum et al. (2015), and Stiller et al. (2024). For the galliform-specific dataset of Ksepka et al. (2023a), molecular constraints followed Kimball et al. (2021), and for the sphenisciform-specific dataset of Ksepka et al. (2023b), molecular constraints followed Pan et al. (2019) and Vianna et al. (2020). Analyses of the datasets from Chen et al. (2019) and Steell et al. (2023a) followed scaffolds employed in those respective studies. Where appropriate, we removed all fossil taxa from comparison matrices prior to analysis.

We applied the relative homoplasy index (RHI) (Steell et al., 2023b) to all applicable molecular topologies per matrix. The RHI measures the amount of homoplasy in a matrix with a given topology against a null distribution of topologies with an estimated maximum amount of homoplasy, therefore enabling accurate comparisons across datasets with different taxon and character samples (Steell et al., 2023b). RHI calculations and matrix preparation were carried out in R 4.3.1 (R Core Team, 2023). All code and custom functions used are available from Supplementary Material Appendix S7. The main dependencies are the packages phangorn (Schliep, 2011), APE (Paradis and Schliep, 2019) and phytools (Revell, 2024). In many cases, the removal of fossil taxa produced invariant characters which were removed prior to analysis. See Supplementary Material Appendix S7 for specific characters removed in each case, except for the Livezey and Zusi (2007) dataset, in which 659 characters were removed due to the large number of non-avian reptile taxa in that matrix. For simplicity, all characters were treated as unordered, polymorphisms and uncertainties were treated the same with the most parsimonious state in the polymorphism being used in the step count, and ‘gaps’ (‘-’) were treated as ambiguities (‘?’). In addition to RHI, consistency (CI) and retention index (RI) were also calculated using phangorn.

## Results

### Phylogenetic analyses

Analysis 1 recovered seven most parsimonious trees (MPT) of 2503 steps (Fig. 8), with a consistency index (CI) of 0.098 and a retention index (RI) of 0.422. The strict consensus of all MPTs bears little topological resemblance to recent phylogenetic consensus. Instead of the universally accepted split between paleognaths and neognaths, the deepest divergence within crown birds was recovered between a clade uniting *Chauna*, *Dendrocygna*, and *Phoenicopterus* and a clade containing all other crown birds. Eleven nodes consistent with current phylogenetic consensus were resolved: the exclusion of *Ichthyornis* from Neornithes, Galliformes (*Alectura* + *Ortalis* + *Rollulus*), Apodiformes (*Streptoprocne* + *Florisuga*), Grues (*Aramus* + *Balearica*), Ralloidea (*Podica* + *Sarothrura* + *Rallus*), Procellariiformes (*Phoebastria* + *Oceanites* + *Pagodroma*), *Oceanites* + *Pagodroma*, Strigiformes (*Tyto* + *Ninox*), Eupasseres (*Erythropitta* + *Neopelma* + *Menura* + *Climacteris*), Tyranni (*Erythropitta* + *Neopelma*), and Passeri (*Menura* + *Climacteris*).

**Figure 8.**
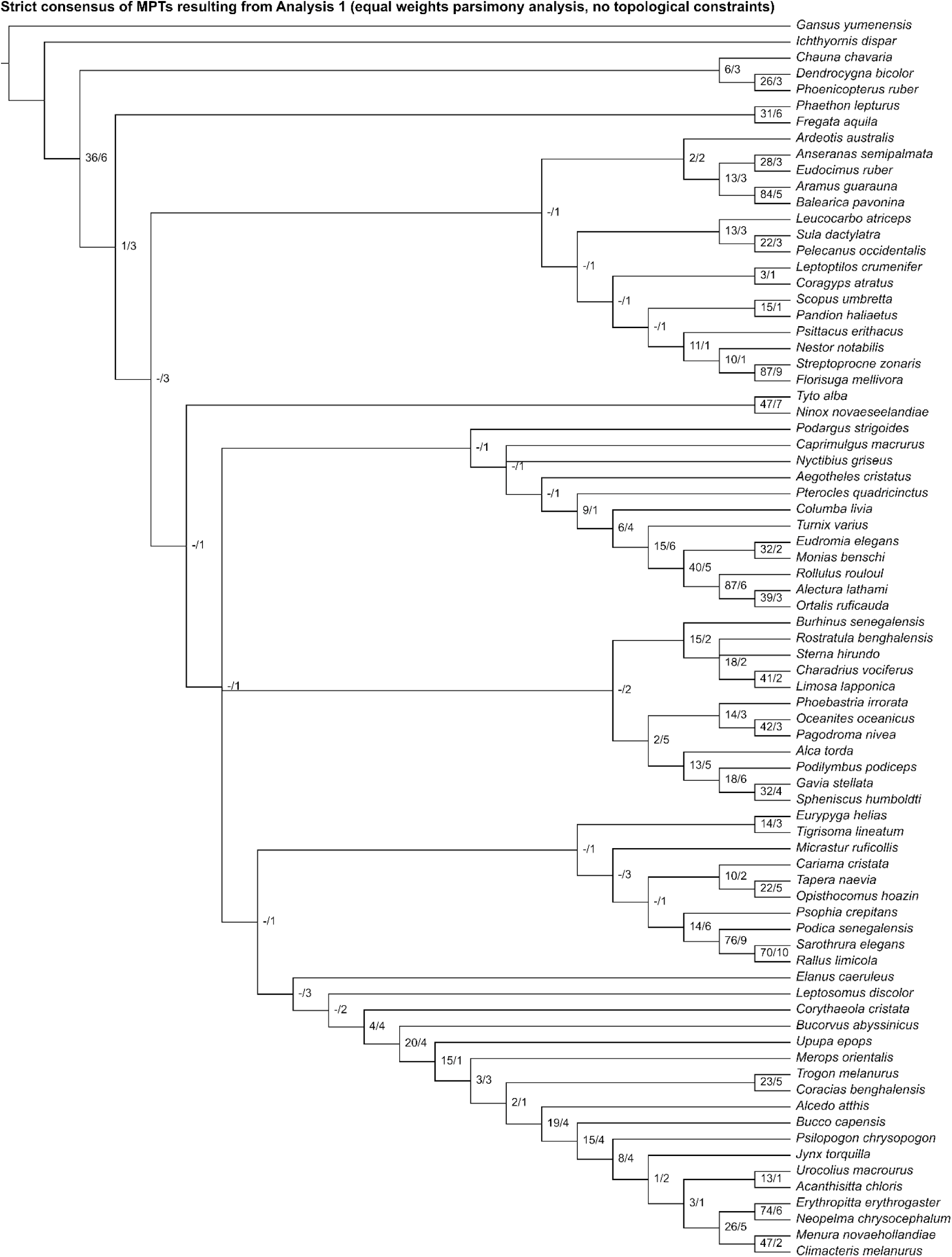
Strict consensus of most parsimonious trees resulting from Analysis 1 (equal weights parsimony analysis without topological constraints). Numbers on nodes represent absolute bootstrap frequencies (left of bar) and Bremer support values (right of bar). A hyphen (−) denotes values of 0.

Analysis 2 recovered one MPT of 2520 steps (Fig. 9), with a CI of 0.097 and a RI of 0.418. This tree is also highly incongruent with current knowledge of avian phylogenetic relationships. *Phoebastria* was recovered as the sister group to all other crown birds. Twelve nodes consistent with current phylogenetic consensus were resolved: the exclusion of *Ichthyornis* from Neornithes, Galliformes (*Alectura* + *Ortalis* + *Rollulus*), Apodiformes (*Streptoprocne* + *Florisuga*), Grues (*Aramus* + *Balearica*), Ralloidea (*Podica* + *Sarothrura* + *Rallus*), *Oceanites* + *Pagodroma*, Strigiformes (*Tyto* + *Ninox*), Bucerotiformes (*Upupa* + *Bucorvus*), Piciformes (*Bucco* + *Psilopogon* + *Jynx*), Psittaciformes (*Nestor* + *Psittacus*), Tyranni (*Erythropitta* + *Neopelma*), and Passeri (*Menura* + *Climacteris*).

**Figure 9.**
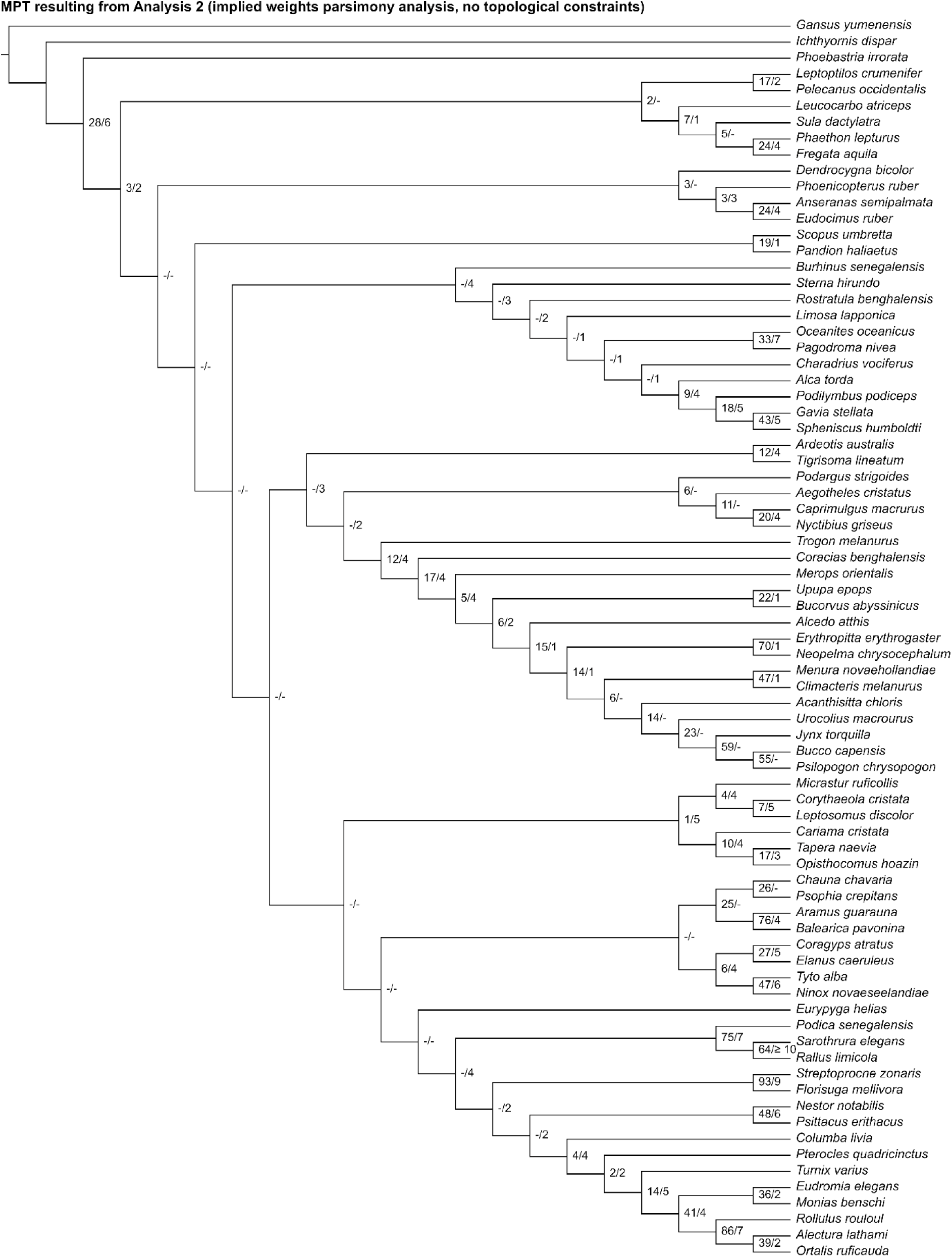
The most parsimonious tree resulting from Analysis 2 (implied weights parsimony analysis without topological constraints). Numbers on nodes represent absolute bootstrap frequencies (left of bar) and Bremer support values (right of bar). A hyphen (−) denotes values of 0.

In the “allcompat” consensus tree recovered by Analysis 3 (Fig. 10), *Phoebastria* was also placed as the sister group to all other crown birds. Thirteen nodes consistent with current phylogenetic consensus were resolved: the exclusion of *Ichthyornis* from Neornithes, Galliformes (*Alectura* + *Ortalis* + *Rollulus*), Apodiformes (*Streptoprocne* + *Florisuga*), Grues (*Aramus* + *Balearica*), Ralloidea (*Podica* + *Sarothrura* + *Rallus*), Heliornithes (*Podica* + *Sarothrura*), *Oceanites* + *Pagodroma*, Strigiformes (*Tyto* + *Ninox*), Bucerotiformes (*Upupa* + *Bucorvus*), Piciformes (*Bucco* + *Psilopogon* + *Jynx*), Eupasseres (*Erythropitta* + *Neopelma* + *Menura* + *Climacteris*), Tyranni (*Erythropitta* + *Neopelma*), and Passeri (*Menura* + *Climacteris*).

**Figure 10.**
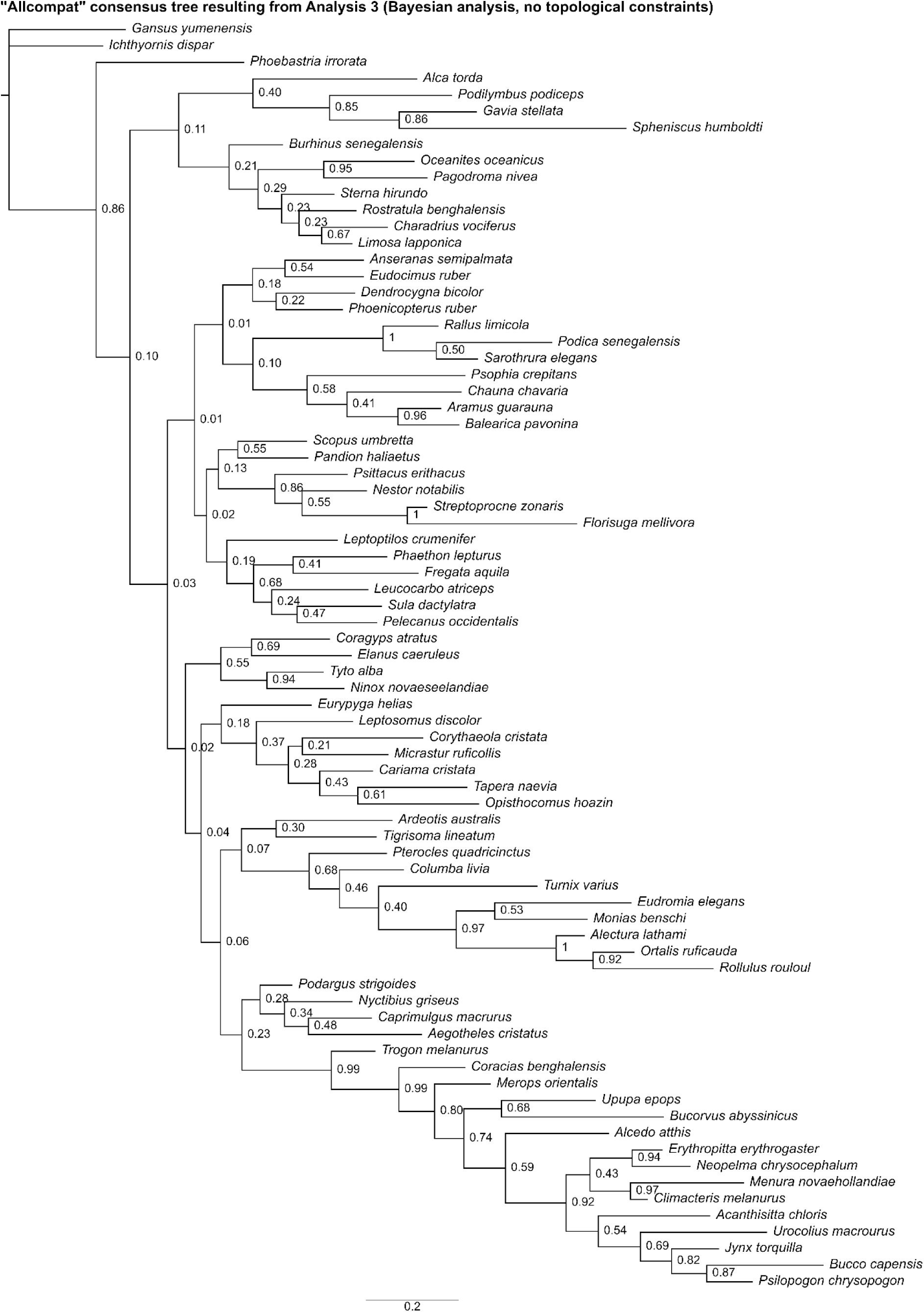
The “allcompat” consensus tree resulting from Analysis 3 (Bayesian analysis without topological constraints). Numbers on nodes represent posterior probabilities.

Analysis 4 recovered 56 MPTs of 2546 steps (Fig. 11), with a CI of 0.096 and a RI of 0.411. The strict consensus of all trees was poorly resolved, but eleven unconstrained nodes consistent with current phylogenetic consensus were recovered: Apodiformes (*Streptoprocne* + *Florisuga*), Grues (*Aramus* + *Balearica*), Ralloidea (*Podica* + *Sarothrura* + *Rallus*), Procellariiformes (*Phoebastria* + *Oceanites* + *Pagodroma*), *Oceanites* + *Pagodroma*, Strigiformes (*Tyto* + *Ninox*), Bucerotiformes (*Upupa* + *Bucorvus*), Psittaciformes (*Nestor* + *Psittacus*), Eupasseres (*Erythropitta* + *Neopelma* + *Menura* + *Climacteris*), Tyranni (*Erythropitta* + *Neopelma*), and Passeri (*Menura* + *Climacteris*). Other results that broadly align with those of recent phylogenetic studies include a clade uniting *Monias*, *Pterocles*, and *Columba* (though *Turnix* was also recovered as a member of this clade) and a clade uniting most members of Charadriiformes other than *Turnix* and *Alca*.

**Figure 11.**
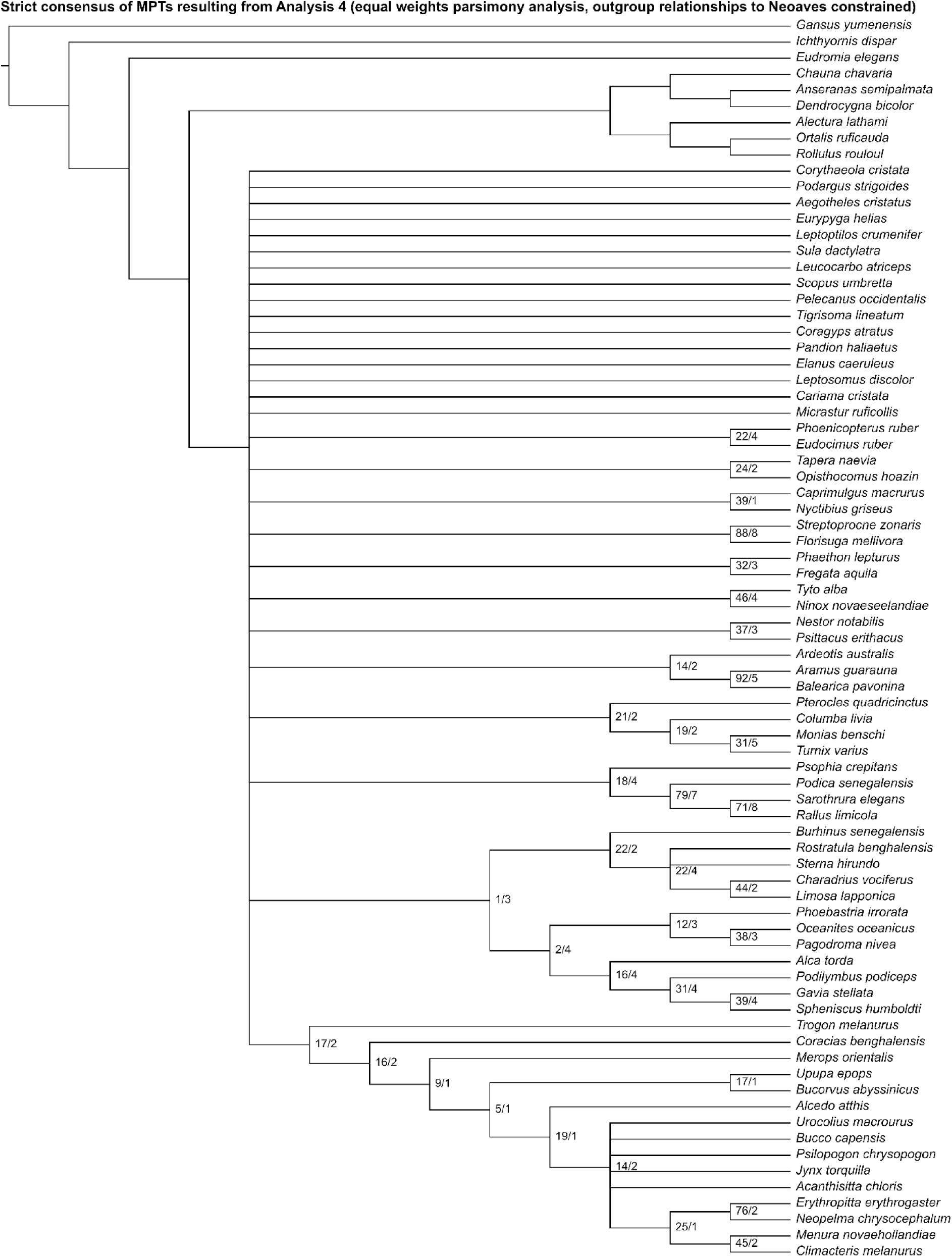
Strict consensus of most parsimonious trees resulting from Analysis 4 (equal weights parsimony analysis run with outgroup relationships to Neoaves constrained). Numbers on nodes represent absolute bootstrap frequencies (left of bar) and Bremer support values (right of bar). A hyphen (−) denotes values of 0. Nodes lacking support values were constrained to be monophyletic.

Analysis 5 recovered one MPT of 2563 steps (Fig. 12), with a CI of 0.096 and a RI of 0.407. A clade uniting *Monias*, *Pterocles*, *Columba*, and *Turnix* was recovered as the sister group to all other neoavians. Twelve unconstrained nodes consistent with current phylogenetic consensus were resolved: Apodiformes (*Streptoprocne* + *Florisuga*), Gruiformes (*Psophia* + *Aramus* + *Balearica* + *Podica* + *Sarothrura* + *Rallus*), Gruoidea (*Psophia* + *Aramus* + *Balearica*), Grues (*Aramus* + *Balearica*), Ralloidea (*Podica* + *Sarothrura* + *Rallus*), *Oceanites* + *Pagodroma*, Strigiformes (*Tyto* + *Ninox*), Bucerotiformes (*Upupa* + *Bucorvus*), Piciformes (*Bucco* + *Psilopogon* + *Jynx*), Psittaciformes (*Nestor* + *Psittacus*), Tyranni (*Erythropitta* + *Neopelma*), and Passeri (*Menura* + *Climacteris*). Other results that broadly align with those of recent phylogenetic studies include a close relationship among *Monias*, *Pterocles*, and *Columba* (though *Turnix* was also recovered as a member of this clade).

**Figure 12.**
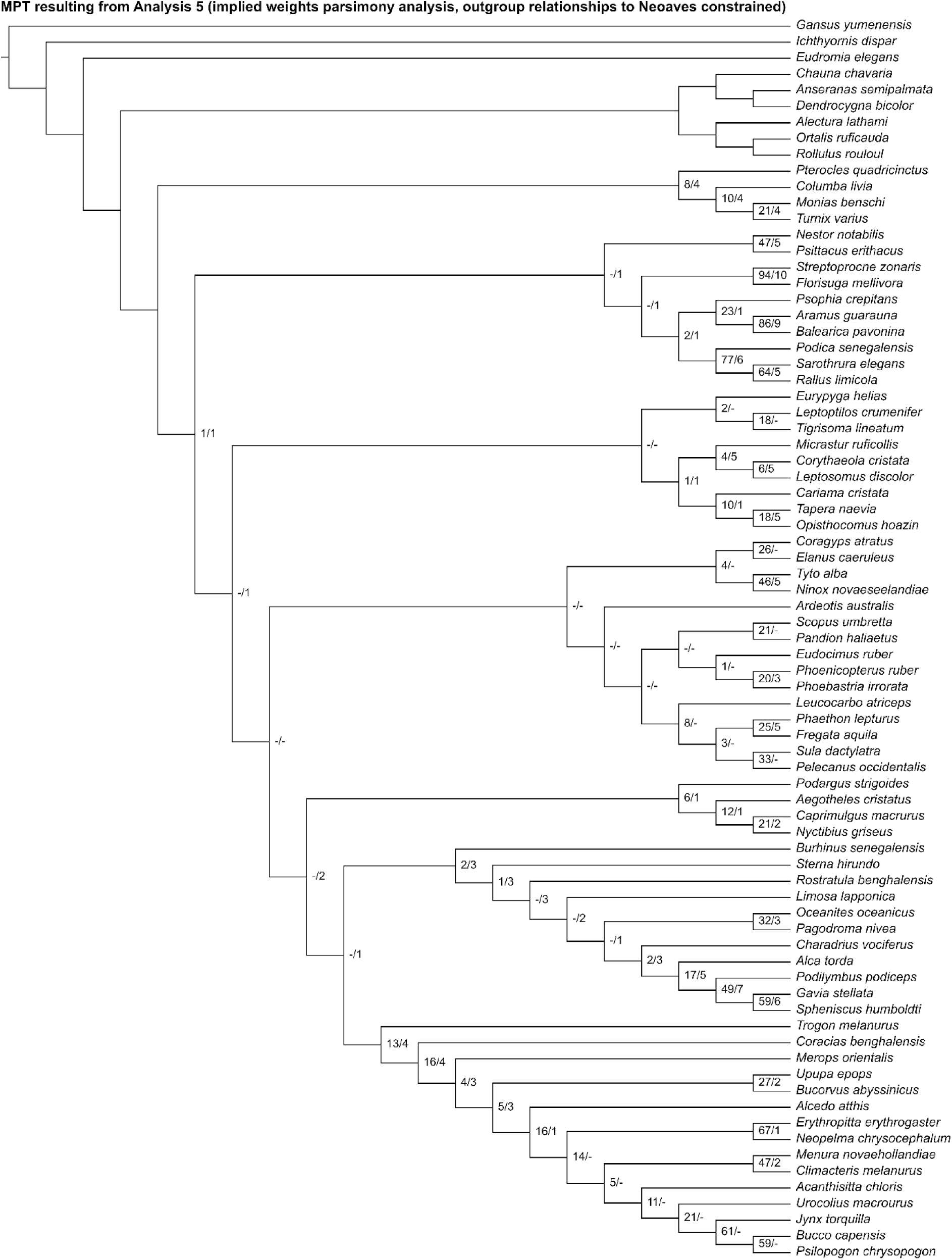
The most parsimonious tree resulting from Analysis 5 (implied weights parsimony analysis run with outgroup relationships to Neoaves constrained). Numbers on nodes represent absolute bootstrap frequencies (left of bar) and Bremer support values (right of bar). A hyphen (−) denotes values of 0. Nodes lacking support values were constrained to be monophyletic.

In the “allcompat” consensus tree recovered by Analysis 6 (Fig. 13), a clade uniting *Phoenicopterus*, *Phoebastria*, *Podilymbus*, *Gavia*, and *Spheniscus* was placed as the sister group to all other neoavians. Twelve unconstrained nodes consistent with current phylogenetic consensus were resolved: Apodiformes (*Streptoprocne* + *Florisuga*), Gruoidea (*Psophia* + *Aramus* + *Balearica*), Grues (*Aramus* + *Balearica*), Ralloidea (*Podica* + *Sarothrura* + *Rallus*), Heliornithes (*Podica* + *Sarothrura*), *Oceanites* + *Pagodroma*, Strigiformes (*Tyto* + *Ninox*), Bucerotiformes (*Upupa* + *Bucorvus*), Piciformes (*Bucco* + *Psilopogon* + *Jynx*), Eupasseres (*Erythropitta* + *Neopelma* + *Menura* + *Climacteris*), Tyranni (*Erythropitta* + *Neopelma*), and Passeri (*Menura* + *Climacteris*). Other results that broadly align with those of recent phylogenetic studies include a clade uniting *Monias*, *Pterocles*, and *Columba* (though *Turnix* was also recovered as a member of this clade) and a clade uniting most members of Charadriiformes other than *Turnix* (though *Oceanites* and *Pagodroma* were also recovered in this clade).

**Figure 13.**
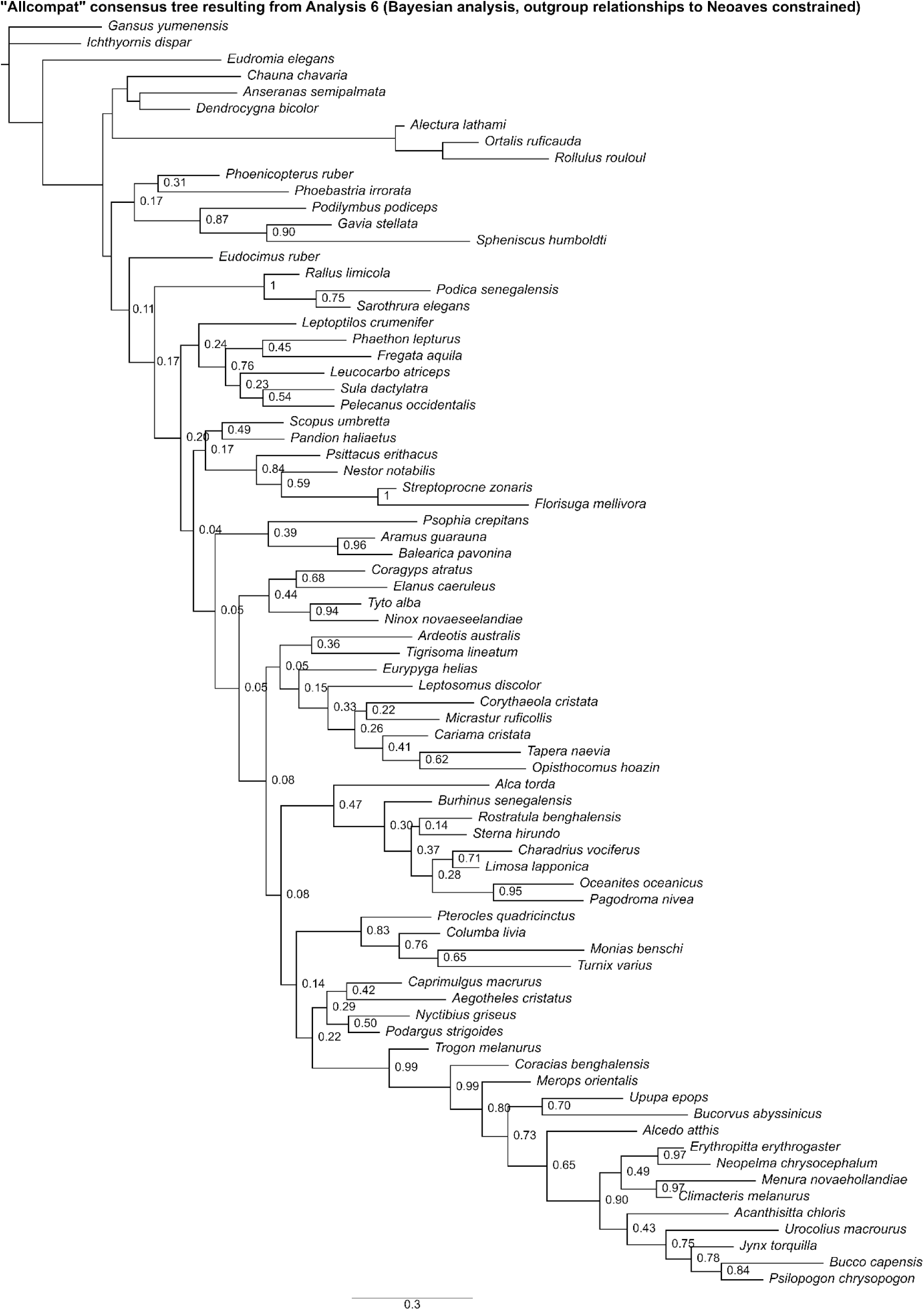
The “allcompat” consensus tree resulting from Analysis 6 (Bayesian analysis run with outgroup relationships to Neoaves constrained). Numbers on nodes represent posterior probabilities. Nodes lacking support values were constrained to be monophyletic.

Analysis 7 recovered eight MPTs of 2764 steps (Fig. 14), with a CI of 0.089 and a RI of 0.355. In the strict consensus of all MPTs, the interrelationships within Neoaves were poorly resolved, but *Ardeotis* was recovered as the sister group to Strisores, *Tapera* was recovered as the sister taxon to *Opisthocomus*, and Mirandornithes was recovered as the sister taxon to Phaethoquornithes. Within Telluraves, Accipitrimorphae was recovered as the sister group to Strigiformes, forming Hieraves; this clade was in turn recovered as the sister group to remaining telluravians.

**Figure 14.**
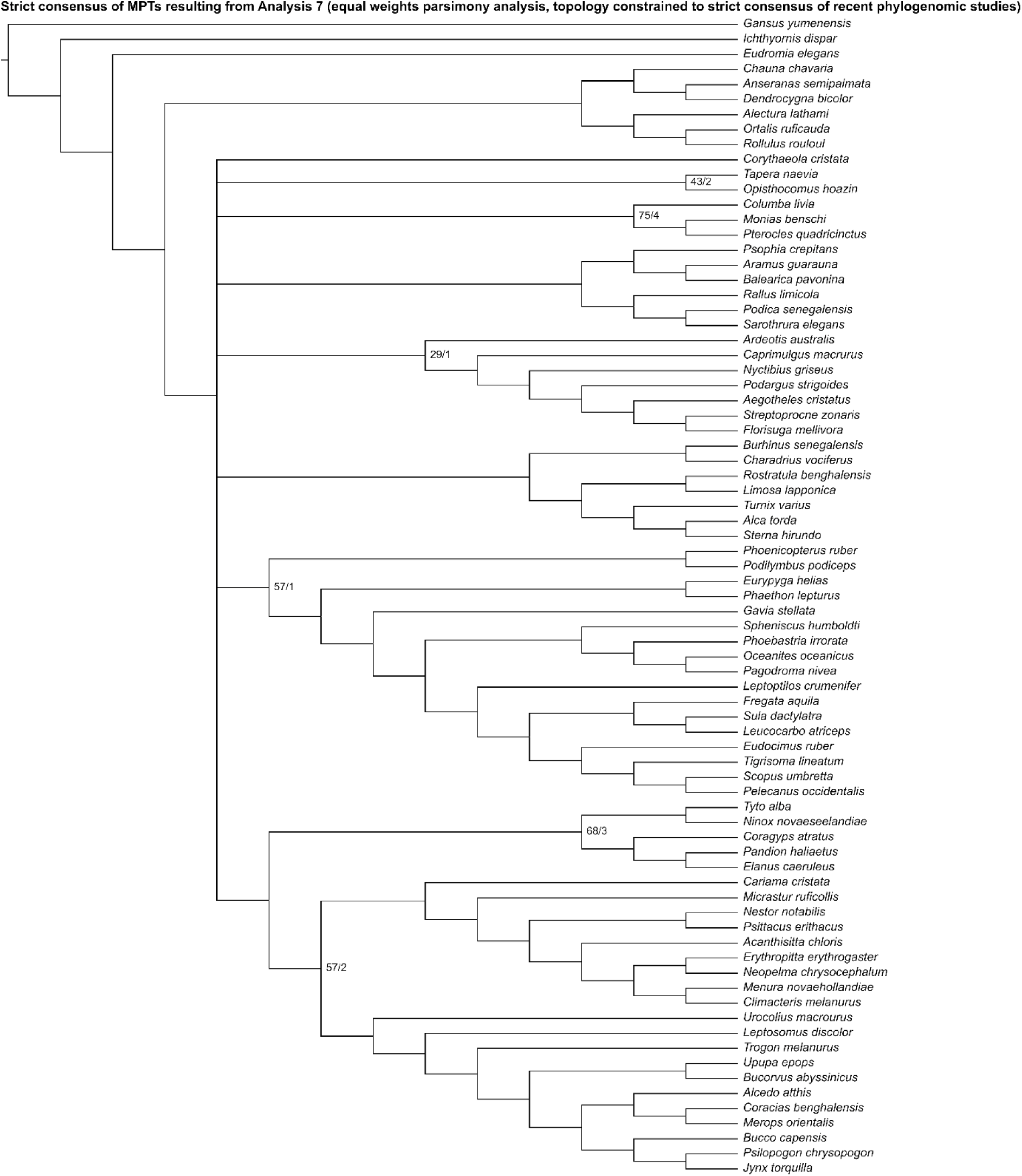
Strict consensus of most parsimonious trees resulting from Analysis 7 (equal weights parsimony analysis run with the topology constrained to a strict consensus of results from recent phylogenomic studies). Numbers on nodes represent absolute bootstrap frequencies (left of bar) and Bremer support values (right of bar). A hyphen (−) denotes values of 0. Nodes lacking support values were constrained to be monophyletic.

Analysis 8 recovered one MPT of 2766 steps (Fig. 15), with a CI of 0.089 and a RI of 0.355. Gruiformes was recovered as the sister group to all other neoavians. The remaining neoavians were divided into one clade consisting of Columbimorphae, *Ardeotis*, Strisores, Charadriiformes, Mirandornithes, and Phaethoquornithes, and another consisting of *Corythaeola*, *Tapera*, *Opisthocomus*, and Telluraves. Within Telluraves, Accipitrimorphae was recovered as the sister group to Strigiformes, forming Hieraves; this clade was in turn recovered as the sister group to remaining telluravians.

**Figure 15.**
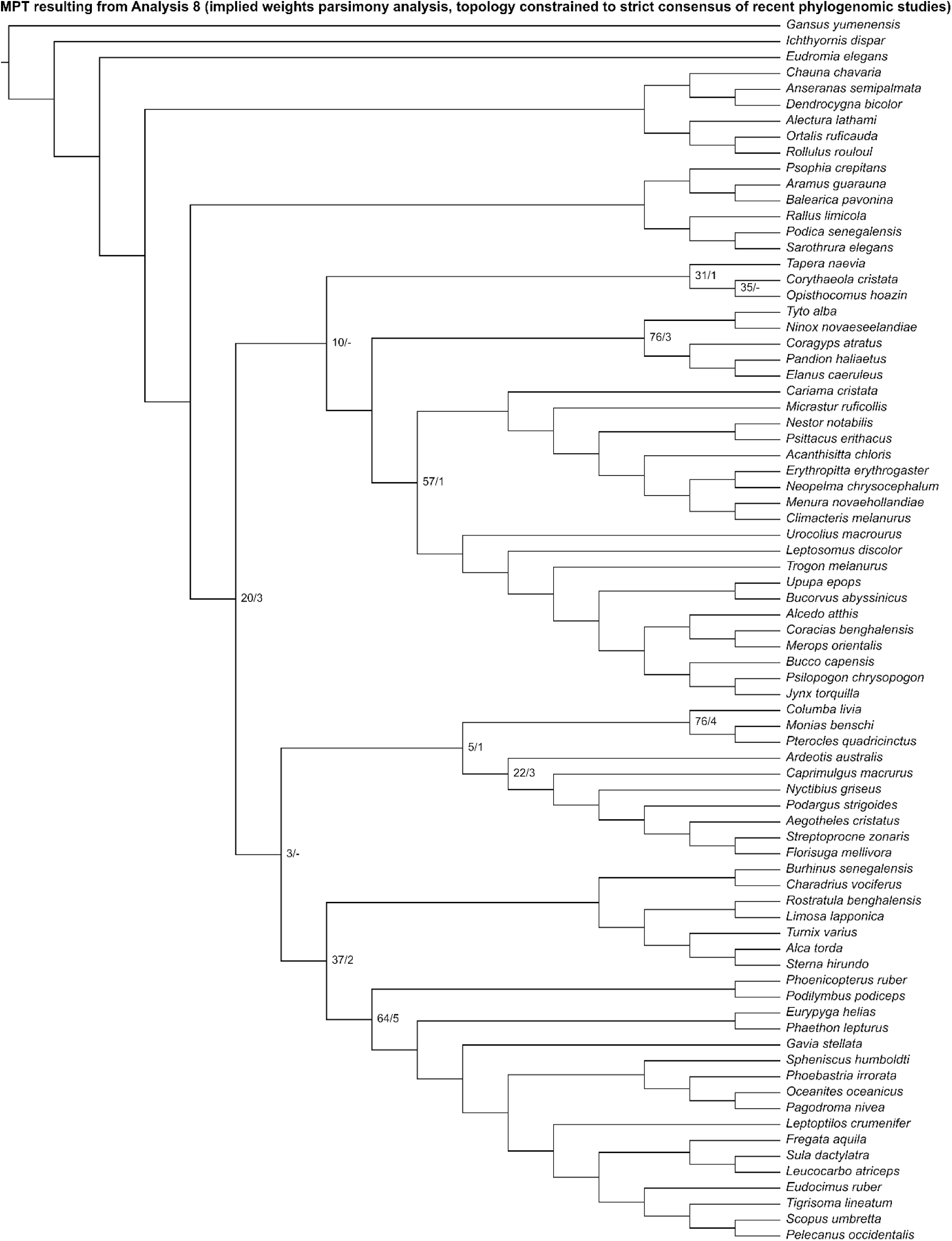
The most parsimonious tree resulting from Analysis 8 (implied weights parsimony analysis run with the topology constrained to a strict consensus of results from recent phylogenomic studies). Numbers on nodes represent absolute bootstrap frequencies (left of bar) and Bremer support values (right of bar). A hyphen (−) denotes values of 0. Nodes lacking support values were constrained to be monophyletic.

In the “allcompat” consensus tree recovered by Analysis 9 (Fig. 16), Mirandornithes was placed as the sister group to all other neoavians. Phaethoquornithes, Gruiformes, and *Ardeotis* were recovered as successively closer relatives to a group containing Telluraves and a clade consisting of Strisores, *Corythaeola*, *Tapera*, *Opisthocomus*, Columbimorphae, and Charadriiformes. Within Telluraves, Accipitrimorphae was recovered as the sister group to Strigiformes, forming Hieraves; this clade was in turn recovered as the sister group to remaining telluravians.

**Figure 16.**
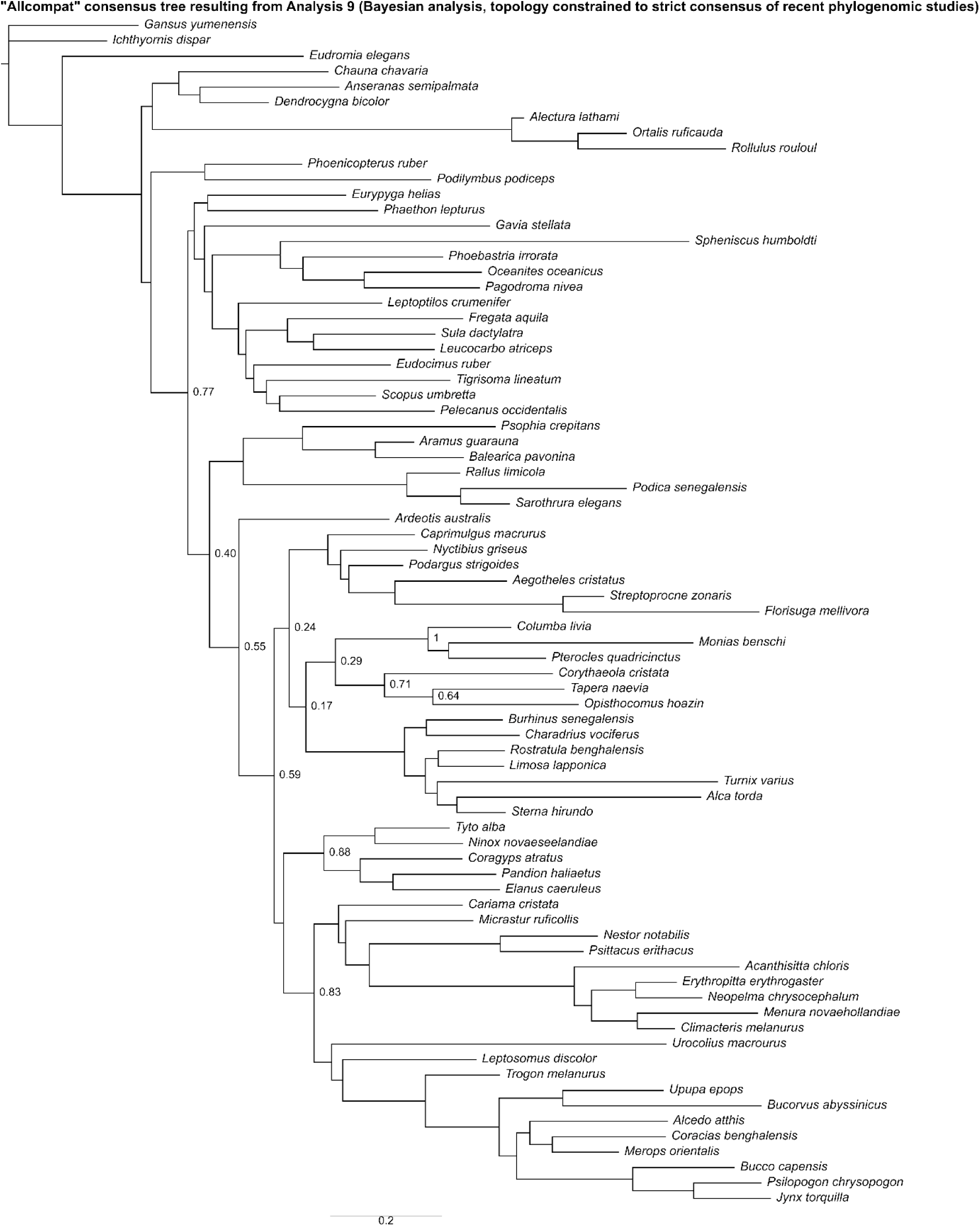
The “allcompat” consensus tree resulting from Analysis 9 (Bayesian analysis run with the topology constrained to a strict consensus of results from recent phylogenomic studies). Numbers on nodes represent posterior probabilities. Nodes lacking support values were constrained to be monophyletic.

Analysis 10 recovered one MPT of 2780 steps (Fig. 17), with a CI of 0.088 and a RI of 0.351. Mirandornithes was recovered as the sister group to the rest of Neoaves. Charadriiformes, Strisores, *Ardeotis* + *Tapera*, *Corythaeola*, *Opisthocomus*, and Gruiformes + Phaethoquornithes were resolved as successively closer relatives to Telluraves. Within Telluraves, Strigiformes was recovered as the sister group to Accipitrimorphae, forming Hieraves, with both being excluded from a clade containing all other telluravians.

**Figure 17.**
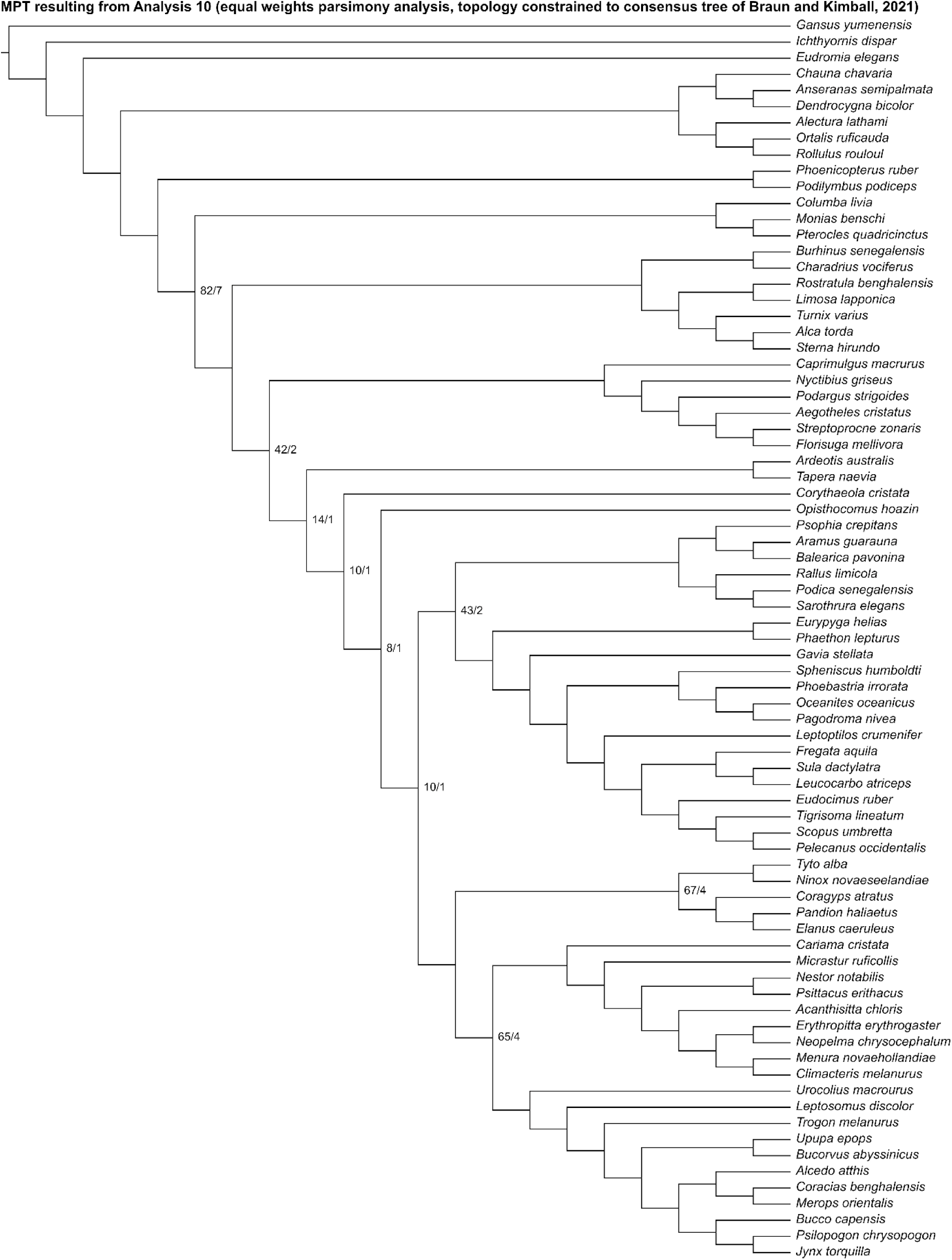
The most parsimonious tree resulting from Analysis 10 (equal weights parsimony analysis run with the topology constrained to the consensus tree proposed by Braun and Kimball, 2021). Numbers on nodes represent absolute bootstrap frequencies (left of bar) and Bremer support values (right of bar). A hyphen (−) denotes values of 0. Nodes lacking support values were constrained to be monophyletic.

Analysis 11 recovered one MPT of 2781 steps (Fig. 18), with a CI of 0.088 and a RI of 0.351. Mirandornithes was recovered as the sister group to all other neoavians. Strisores + Charadriiformes and *Ardeotis* + *Tapera* + *Corythaeola* + *Opisthocomus* + Gruiformes + Phaethoquornithes were resolved as successively closer relatives to Telluraves. Within Telluraves, Strigiformes was recovered as the sister group to Accipitrimorphae, forming Hieraves, with both being excluded from a clade containing all other telluravians.

**Figure 18.**
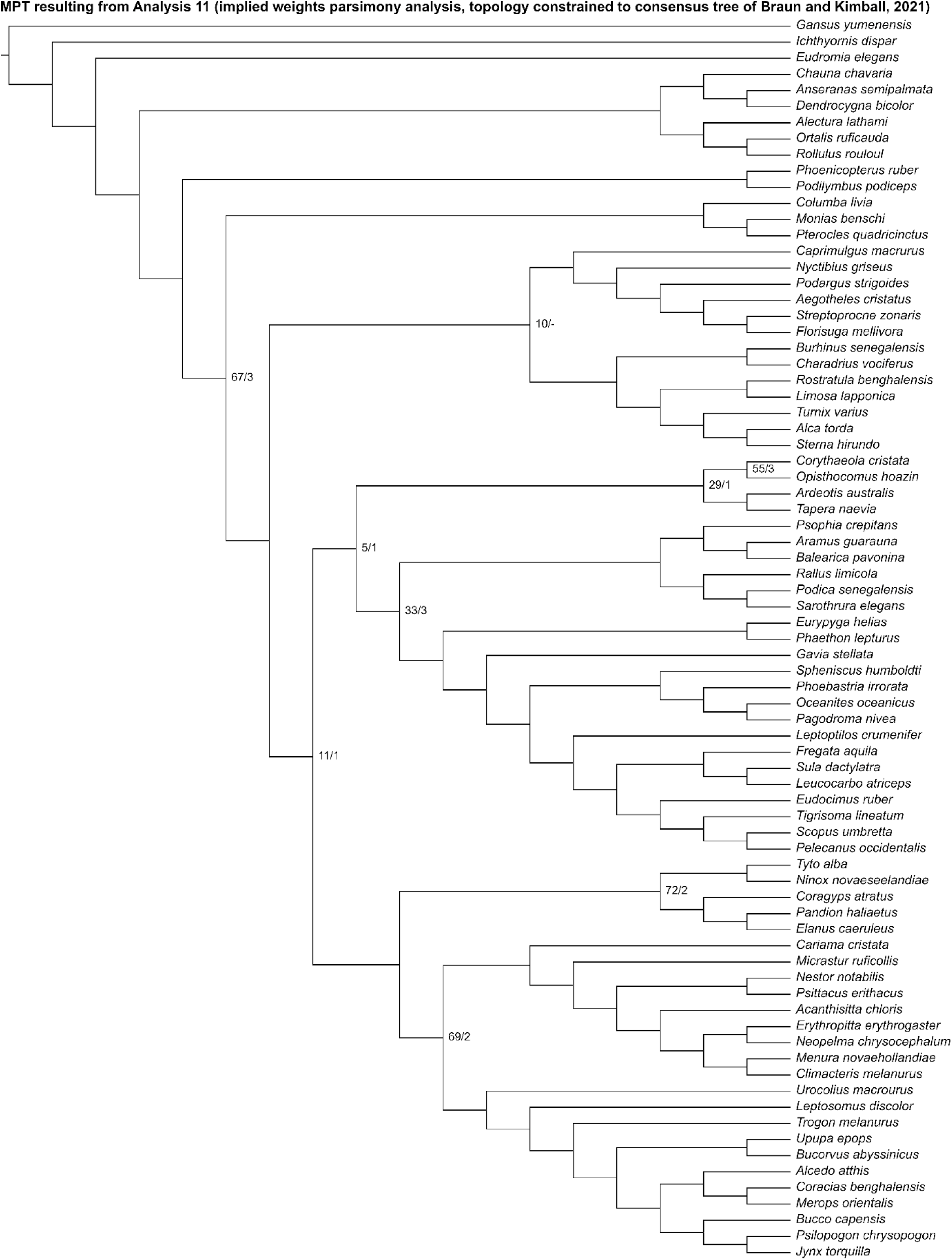
The most parsimonious tree resulting from Analysis 11 (implied weights parsimony analysis run with the topology constrained to the consensus tree proposed by Braun and Kimball, 2021). Numbers on nodes represent absolute bootstrap frequencies (left of bar) and Bremer support values (right of bar). A hyphen (−) denotes values of 0. Nodes lacking support values were constrained to be monophyletic.

In the “allcompat” consensus tree recovered by Analysis 12 (Fig. 19), Mirandornithes was placed as the sister group to all other neoavians. Phaethoquornithes, Gruiformes, *Ardeotis* + *Tapera* + *Corythaeola* + *Opisthocomus*, and Strisores + Charadriiformes were resolved as successively closer relatives to Telluraves. Within Telluraves, Strigiformes was recovered as the sister group to Accipitrimorphae, forming Hieraves, with both being excluded from a clade containing all other telluravians.

**Figure 19.**
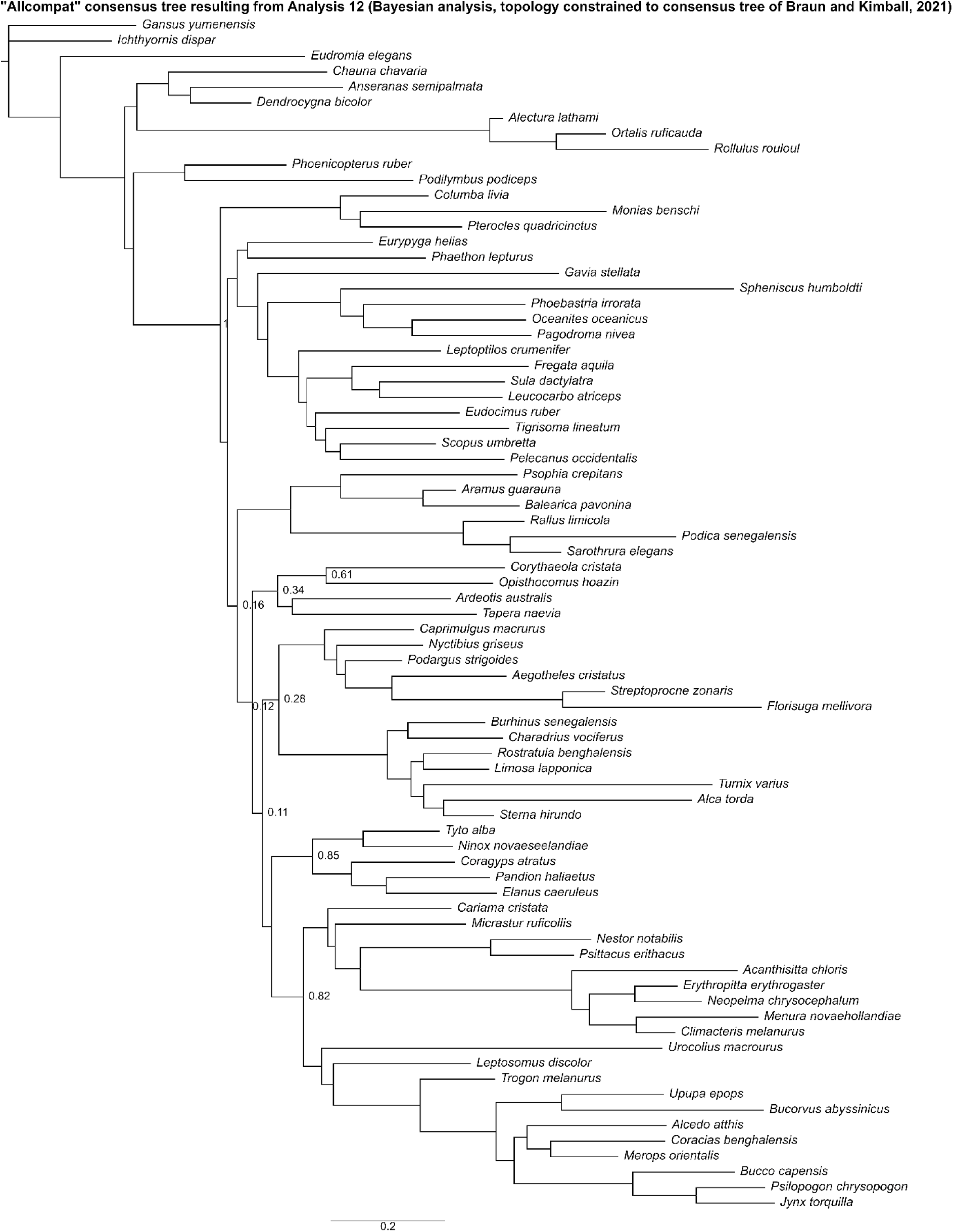
The “allcompat” consensus tree resulting from Analysis 12 (Bayesian analysis run with the topology constrained to the consensus tree proposed by Braun and Kimball, 2021). Numbers on nodes represent posterior probabilities. Nodes lacking support values were constrained to be monophyletic.

### Comparisons with molecular topologies

Values for tree distance metrics comparing the most parsimonious trees and “allcompat” consensus trees recovered by the morphological analyses in the present study to topologies recovered by recent avian phylogenomic studies are provided in Table 2. In general, analyses constrained to the consensus tree of Braun and Kimball (2021) (Analyses 10–12) produced trees with the shortest distances from molecular topologies. Under analyses where no topological constraints were applied (Analyses 1–3), results of implied weights parsimony analysis and especially Bayesian inference exhibited shorter distances from molecular topologies than those of equal weights parsimony analysis. However, in analyses implementing topological constraints, implied weights parsimony and Bayesian analyses often yielded values for RF distances comparable to or exceeding those based on equivalent equal weights parsimony analyses.

**Table 2.** Values for RF distances comparing the most parsimonious trees and “allcompat” consensus trees recovered by the morphological analyses in the present study to topologies recovered by recent avian phylogenomic studies.

|  | Jarvis et al.<br>(2014) | Prum et al.<br>(2015) | Kuhl et al.<br>(2021) | Stiller et al.<br>(2024) | S. Wu et al.<br>(2024) |
| --- | --- | --- | --- | --- | --- |
| T1:<br>Unconstrained<br>topology<br>analyzed under<br>equal weights<br>parsimony | 126 (mean<br>= 126) | 126 (mean<br>= 126) | 126 (mean<br>= 126) | 126 (mean<br>= 126) | 126 (mean<br>= 126) |
| T2:<br>Unconstrained<br>topology<br>analyzed under<br>implied weights<br>parsimony | 124 | 124 | 124 | 124 | 124 |
| T3:<br>Unconstrained<br>topology<br>analyzed under<br>Bayesian<br>inference | 120 | 120 | 120 | 120 | 120 |
| T4: Outgroups<br>constrained and<br>analyzed under<br>equal weights<br>parsimony | 106–110<br>(mean =<br>108.8) | 106–110<br>(mean =<br>108.8) | 106–110<br>(mean =<br>108.8) | 106–110<br>(mean =<br>108.8) | 106–110<br>(mean =<br>108.8) |
| T5: Outgroups<br>constrained and | 108 | 108 | 108 | 108 | 108 |
| analyzed under<br>implied weights<br>parsimony |  |  |  |  |  |
| T6: Outgroups<br>constrained and<br>analyzed under<br>Bayesian<br>inference | 108 | 108 | 108 | 108 | 108 |
| T7: Constrained<br>to strict<br>consensus of all<br>reference trees<br>and analyzed<br>under equal<br>weights<br>parsimony | 22 (mean =<br>22) | 20–22<br>(mean =<br>21.5) | 24 (mean =<br>24) | 20 (mean =<br>20) | 20 (mean =<br>20) |
| T8: Constrained<br>to strict<br>consensus of all<br>reference trees<br>and analyzed<br>under implied<br>weights<br>parsimony | 22 | 20 | 24 | 20 | 20 |
| T9: Constrained<br>to strict<br>consensus of all<br>reference trees<br>and analyzed<br>under Bayesian<br>inference | 22 | 22 | 22 | 18 | 20 |
| T10: Constrained<br>to consensus tree<br>of Braun and<br>Kimball and<br>analyzed under<br>equal weights<br>parsimony | 20 | 20 | 22 | 16 | 20 |
| T11: Constrained<br>to consensus tree<br>of Braun and<br>Kimball and<br>analyzed under<br>implied weights<br>parsimony | 20 | 20 | 22 | 16 | 20 |
| T12: Constrained<br>to consensus tree<br>of Braun and<br>Kimball and | 20 | 20 | 22 | 16 | 20 |
| analyzed under<br>Bayesian<br>inference |  |  |  |  |  |

The molecular topologies studied did not differ from each other substantially in distance from each morphological tree recovered, but on average the Stiller et al. (2024) topology exhibited slightly shorter RF distances from morphological trees than the others when results were constrained to consensus trees of molecular studies, whereas the Kuhl et al. (2021) topology exhibited slightly higher RF distances.

When topologies recovered by recent phylogenomic analyses were enforced under the current dataset, the Jarvis et al. (2014) and S. Wu et al. (2024) topologies resulted in the longest trees and the lowest CI and RI, and the latter also exhibits the highest RHI. The Prum et al. (2015) and Stiller et al. (2024) topologies resulted in the shortest trees, highest RI and CI, and lowest RHI. Specific values for these metrics are reported in Table 3.

**Table 3.** Characterization of trees resulting from enforcing the results of recent avian phylogenomic studies on the current morphological dataset with ordered characters included.

|  | Jarvis et al.<br>(2014) | Prum et al.<br>(2015) | Kuhl et al.<br>(2021) | Stiller et al.<br>(2024) | S. Wu et al.<br>(2024) |
| --- | --- | --- | --- | --- | --- |
| Tree length | 2807 steps | 2796 steps | 2801 steps | 2796 steps | 2807 steps |
| CI | 0.087 | 0.088 | 0.087 | 0.088 | 0.087 |
| RI | 0.344 | 0.347 | 0.346 | 0.347 | 0.344 |
| RHI | 0.791 | 0.788 | 0.791 | 0.788 | 0.792 |

The heatmap generated from the similarity matrix incorporating all characters in the present study suggests that our morphological dataset contains relatively low phylogenetic signal (Fig. 20). Although relatively high intra-group similarity is seen in some clades (e.g., Anseriformes, Galliformes, Columbimorphae, some Strisores, Grues, Ralloidea, most Charadriiformes, Accipitrimorphae, Strigiformes, Piciformes, Psittaciformes, and Passeriformes), many species exhibit comparable or greater similarity to distantly related taxa in comparison with their closest relatives included in the dataset.

**Figure 20.**
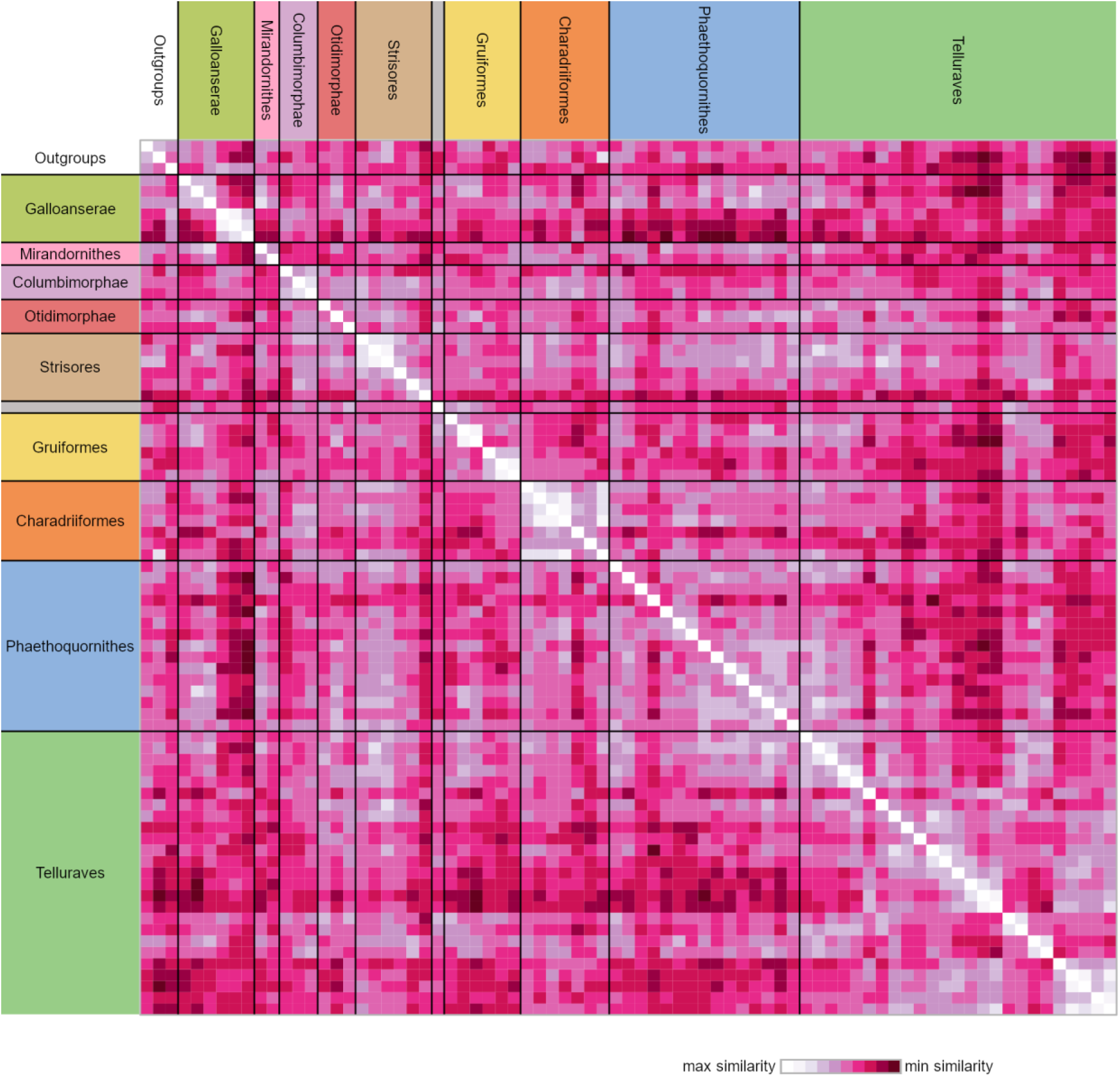
Heatmap of symmetric pairwise-taxon similarity matrix using our complete character dataset (n = 204). The unlabeled monotypic group positioned between Strisores and Gruiformes represents *Opisthocomus*.

This general pattern is also seen in heatmaps generated from subsets of our character dataset partitioned by individual skeletal elements (Fig. 21). Of note, however, is that characters from the humerus and carpometacarpus appear to more closely reflect phylogenetic relationships compared to those of the other elements analyzed.

**Figure 21.**
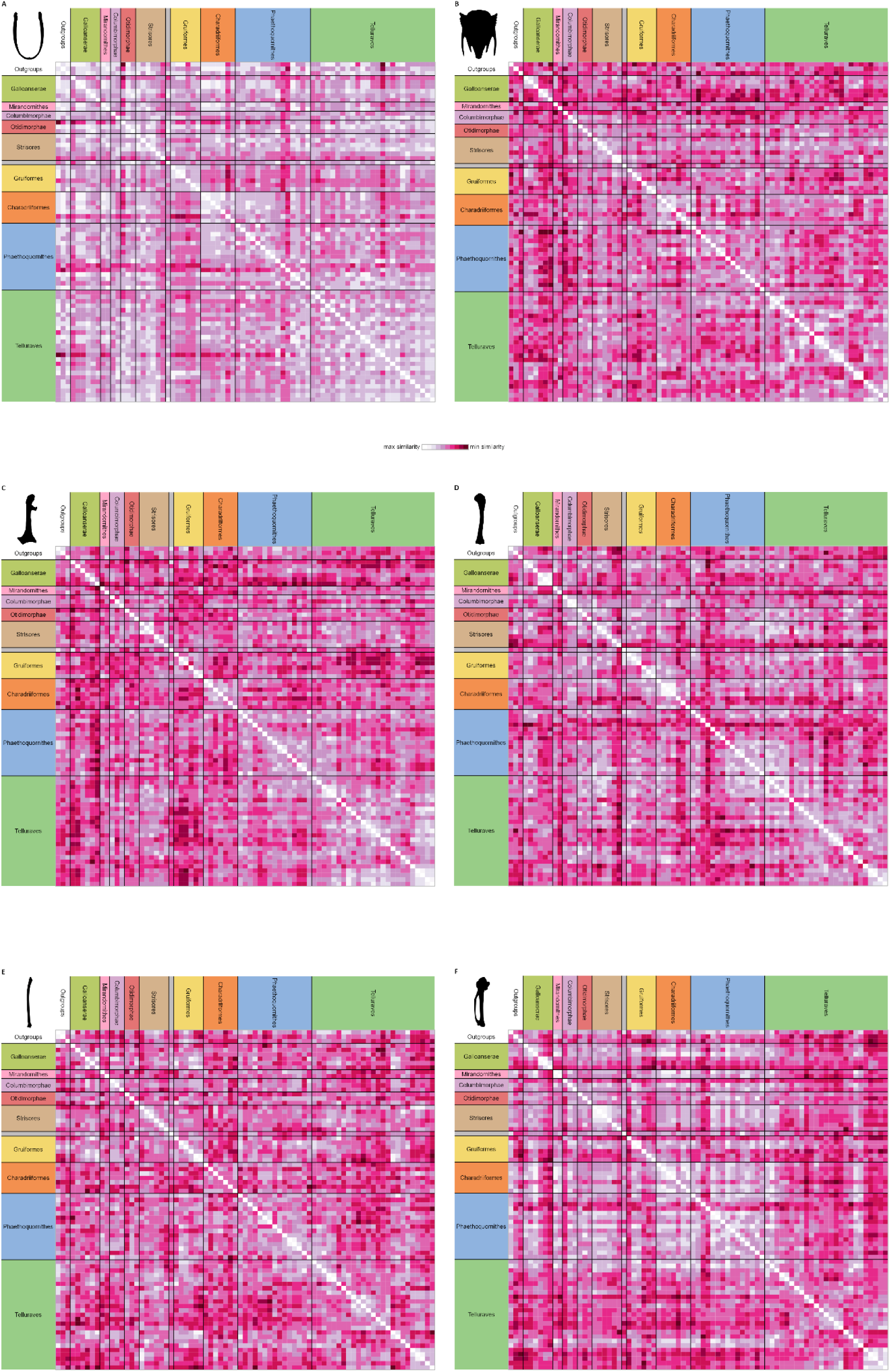
Heatmaps of symmetric pairwise-taxon similarity matrices of subsets of our character dataset partitioned by individual element, including the furcula (**A**, n = 13), sternum (**B**, n = 47), coracoid (**C**, n = 26), humerus (**D**, n = 45), ulna (**E**, n = 15), and carpometacarpus (**F**, n = 31). The unlabeled monotypic group positioned between Strisores and Gruiformes represents *Opisthocomus*.

### Homoplasy across avian morphological matrices

Our comparative homoplasy analysis shows that the present matrix is highly homoplastic with an RHI value of 0.79–0.80 when all characters are treated as unordered (Fig. 22, Table 4). A value of zero or close to zero would indicate no or little homoplasy within the dataset, and a value of one or close to one would indicate high levels of homoplasy comparable to a matrix with no phylogenetic signal present with regards to the empirical tree. Indeed, when states within the empirical pectoral matrix are permuted without replacement within characters (therefore representing a matrix with no phylogenetic signal with respect to any external tree), RHI values range between 0.99–1.00 for the published topological constraints. We find no significant difference in values across different topological constraints per matrix with near-identical RHI values and overlapping confidence intervals for all datasets sampled (Table 4). The Mayr and Clarke (2003) crown bird dataset exhibits comparable levels of homoplasy to the present matrix (Fig. 22, Table 4), and together both matrices are the most homoplastic out of the datasets sampled. The Ksepka et al. (2023b) sphenisciform dataset shows the lowest relative homoplasy with an RHI of 0.19.

**Figure 22.**
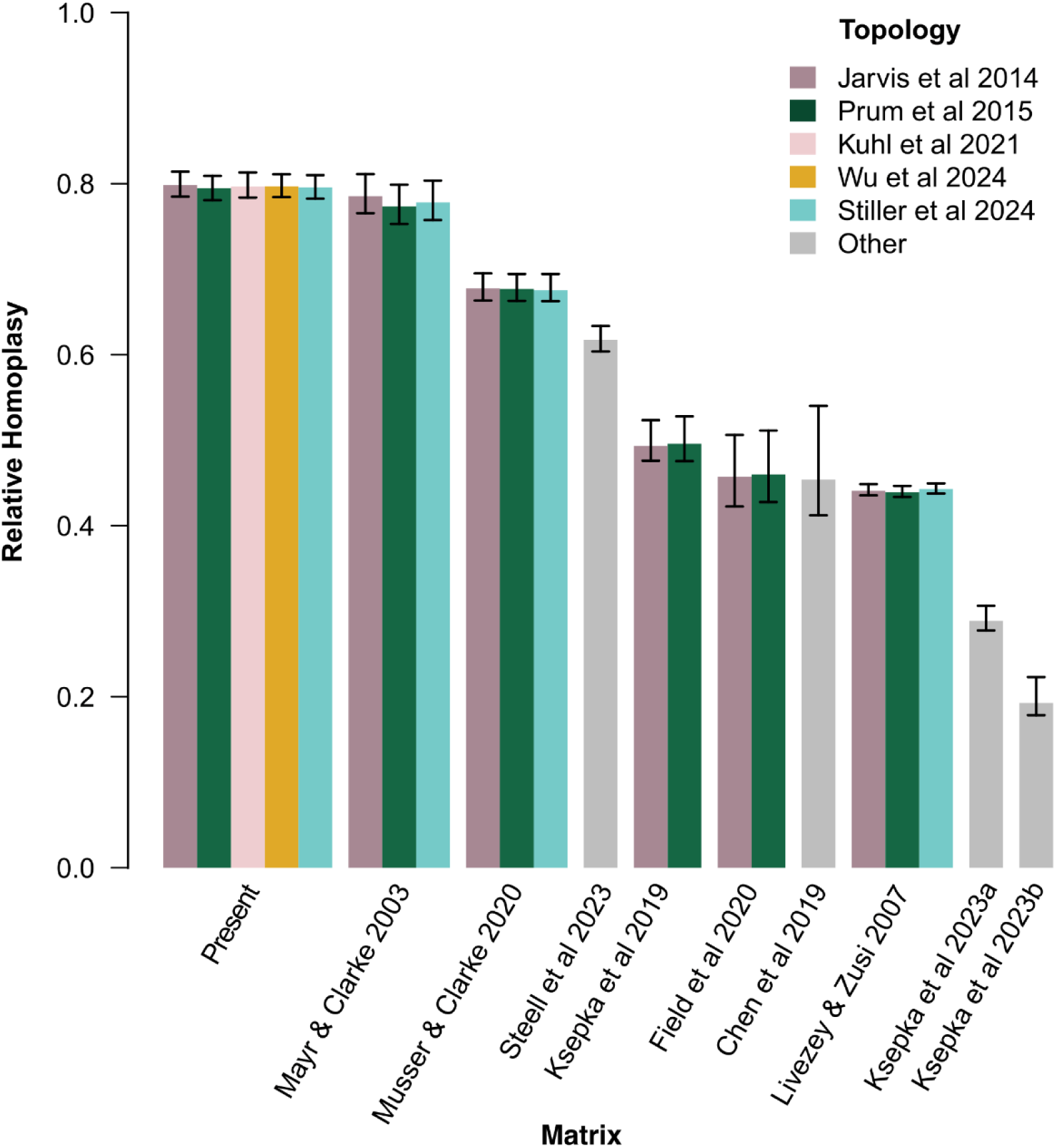
Bar plot showing relative homoplasy index values for the present matrix and published matrices of extant birds for a range of molecular topologies. Error bars indicate the 5% and 95% quantile ranges. A value of 1 indicates maximum possible homoplasy. Where topology is denoted as “other”, details of molecular reference trees are indicated in the main text and Table 4.

**Table 4.** Summary of the relative homoplasy results for published extant avian datasets in addition to consistency and retention indices. Note that RHI measures homoplasy so that zero indicates no homoplasy and one indicates maximum possible homoplasy, whereas CI and RI indicate the opposite.

| Study | Taxa | No.<br>Taxa | No.<br>Characters | Constraint | Tree<br>length | RHI | RHI<br>5% | RHI<br>95% | CI | RI |
| --- | --- | --- | --- | --- | --- | --- | --- | --- | --- | --- |
| Present | Neornithes | 75 | 203 | Jarvis et al.<br>2014 | 2766 | 0.80 | 0.78 | 0.81 | 0.09 | 0.34 |
|  |  |  |  | Prum et al.<br>2015 | 2755 | 0.79 | 0.78 | 0.81 | 0.09 | 0.34 |
|  |  |  |  | Kuhl et al.<br>2021 | 2760 | 0.80 | 0.78 | 0.81 | 0.09 | 0.34 |
|  |  |  |  | S. Wu et al.<br>2024 | 2764 | 0.80 | 0.78 | 0.81 | 0.09 | 0.34 |
|  |  |  |  | Stiller et<br>al. 2024 | 2757 | 0.80 | 0.78 | 0.81 | 0.09 | 0.34 |
| Mayr &<br>Clarke<br>2003 | Neornithes | 43 | 145 | Jarvis et al.<br>2014 | 824 | 0.79 | 0.76 | 0.81 | 0.20 | 0.33 |
|  |  |  |  | Prum et al.<br>2015 | 813 | 0.77 | 0.75 | 0.80 | 0.20 | 0.35 |
|  |  |  |  | Stiller et<br>al. 2024 | 817 | 0.78 | 0.76 | 0.80 | 0.20 | 0.34 |
| Livezey &<br>Zusi<br>2007 | Neornithes | 150 | 2295 | Jarvis et al.<br>2014 | 17967 | 0.44 | 0.44 | 0.45 | 0.19 | 0.61 |
|  |  |  |  | Prum et al.<br>2015 | 17929 | 0.44 | 0.43 | 0.45 | 0.19 | 0.61 |
|  |  |  |  | Stiller et<br>al. 2024 | 18046 | 0.44 | 0.44 | 0.45 | 0.19 | 0.61 |
| Field et<br>al. 2020 | Neornithes | 26 | 293 | Jarvis et al.<br>2014 | 1094 | 0.45 | 0.42 | 0.50 | 0.36 | 0.63 |
|  |  |  |  | Prum et al.<br>2015 | 1100 | 0.46 | 0.43 | 0.51 | 0.36 | 0.63 |
| Musser &<br>Clarke<br>2020 | Neornithes | 52 | 692 | Jarvis et al.<br>2014 | 5247 | 0.68 | 0.66 | 0.70 | 0.16 | 0.45 |
|  |  |  |  | Prum et al.<br>2015 | 5246 | 0.68 | 0.66 | 0.70 | 0.16 | 0.45 |
|  |  |  |  | Stiller et<br>al. 2024 | 5237 | 0.68 | 0.66 | 0.69 | 0.16 | 0.45 |
| Chen et<br>al. 2019 | Strisores | 28 | 114 | Chen et al.<br>2019 | 233 | 0.46 | 0.41 | 0.55 | 0.54 | 0.63 |
| Ksepka<br>et al.<br>2019 | Telluraves | 34 | 144 | Jarvis et al.<br>2014 | 548 | 0.49 | 0.47 | 0.53 | 0.33 | 0.57 |
|  |  |  |  | Prum et al.<br>2015 | 549 | 0.50 | 0.48 | 0.53 | 0.33 | 0.57 |
| Ksepka et al. 2023a | Galliformes | 61 | 135 | Kimball et al. 2021 | 442 | 0.29 | 0.28 | 0.31 | 0.34 | 0.75 |
| Ksepka et al. 2023b | Sphenisciformes | 25 | 245 | Pan et al. 2019 & Vianna et al. 2020 | 422 | 0.19 | 0.18 | 0.22 | 0.76 | 0.83 |
| Steell et al. 2023a | Passeriformes | 142 | 49 | Steell et al. 2023a | 990 | 0.62 | 0.63 | 0.60 | 0.06 | 0.51 |

### Optimization of synapomorphies onto molecular topologies

Potential synapomorphies from the pectoral and forelimb skeleton were recovered for most avian clades strongly supported by molecular phylogenetic analyses, though many of these characters appear to be homoplastic across crown bird phylogeny and some were not inferred to be synapomorphic under all alternative molecular topologies. Specific characters optimized as synapomorphies for major clades and their phylogenetic distributions are detailed below.

## Discussion

### Distribution of Optimized Osteological Synapomorphies for Well-Supported Clades

Although potential morphological synapomorphies could be optimized for most of the avian clades recognized by molecular phylogenetic analyses, not all of these character states are equally convincing as synapomorphies when their distribution among crown birds and relevant fossil taxa is considered. Here, remarks on character state distributions in fossil taxa are limited to characters that can be assessed based on previously published fossil descriptions. Synapomorphies optimizing onto specific molecular phylogenetic topologies are denoted below as J14 (for Jarvis et al., 2014), P15 (for Prum et al., 2015), K21 (for Kuhl et al., 2021), S24 (for Stiller et al., 2024), and W24 (for S. Wu et al., 2024).

#### Neornithes

Thirteen character states were optimized as potential synapomorphies of Neornithes (the avian crown group), of which nine were consistently inferred as such across all alternative molecular topologies studied.

- Four articular facets on costal margin of sternum or fewer (char. 36: 2 > 1, reversed in Anseriformes, Mirandornithes, *Monias*, *Corythaeola*, *Podargus*, Apodiformes, *Opisthocomus*, Gruiformes, Charadriiformes, Phaethoquornithes, Accipitrimorphae, and Australaves); P15.
- Single pair of caudal fenestrae in sternum (char. 49: 2 > 1, reversed in *Pterocles*, *Columba*, *Corythaeola*, *Ardeotis*, *Tapera*, Vanescaves, *Opisthocomus*, Charadriiformes, Phaethontimorphae, *Pagodroma*, *Eudocimus*, *Coragyps*, *Ninox*, Coraciimorphae, and *Acanthisitta*); K21, S24.
- Median trabecula of sternum extends caudal of caudolateral trabeculae (char. 60: 1 > 0, reversed in *Dendrocygna*, Mirandornithes, Strisores, Heliornithes, *Phaethon*, *Spheniscus*, *Phoebastria*, Suliformes, *Scopus*, *Pelecanus*, *Pandion* + *Elanus*, *Tyto*, Pici, *Micrastur*, and *Nestor*); J14, P15, K21, S24, W24.
- Scapular cotyle on coracoid a shallow concavity (char. 66: 0 > 1, reversed in *Chauna*, *Dendrocygna*, *Phoenicopterus*, *Pterocles*, *Ardeotis*, Apodiformes, *Opisthocomus*, Charadriiformes, *Gavia*, *Leptoptilos*, *Pelecanus*, and *Tigrisoma*); J14, P15, K21, S24, W24.
- Procoracoid process of coracoid modest or reduced (char. 68: 2 > 1, reversed in *Chauna*, *Pterocles*, *Columba*, *Corythaeola*, *Tapera*, Letornithes, *Opisthocomus*, Gruoidea, *Podica*, Charadriiformes, *Phaethon*, *Spheniscus*, *Oceanites* + *Pagodroma*, Pelecanimorphae, Strigiformes, *Leptosomus*, Picocoraciades, *Cariama*, *Micrastur*, *Psittacus*, and Tyranni); P15, K21, S24.
- Impression of m. sternocoracoidei on coracoid shallow or hard to discern (char. 76: 1 > 0, reversed in *Chauna*, *Phoenicopterus*, *Aegotheles*, *Opisthocomus*, Grues, *Sarothrura*, *Rallus*, *Charadrius*, *Turnix*, *Sterna*, *Spheniscus*, *Phoebastria*, *Coragyps*, *Coracias*, and *Cariama*); J14, P15, K21, S24, W24.
- Lateral process of coracoid unhooked or weakly hooked (char. 80: 1 > 0, reversed in *Anseranas*, *Rollulus*, *Ardeotis*, *Tapera*, Strisores, Grues, *Burhinus*, *Sterna*, *Phaethon*, Procellariiformes, *Leptoptilos*, *Fregata*, *Leucocarbo*, *Scopus* + *Pelecanus*, *Pandion*, *Trogon*, *Bucrovus*, *Coracias*, *Bucco*, and *Psittacus*); J14, P15, K21, S24, W24.
- Ulna less than or equal to 40% of humerus + ulna + carpometacarpus length (char. 98: 1 > 0, reversed in *Chauna*, *Phoenicopterus*, *Columba*, *Ardeotis*, Strisores, Grues, *Burhinus*, *Rostratula*, *Sterna*, Phaethontimorphae, *Phoebastria*, Pelecanimorphae, Accipitrimorphae, Strigiformes, Cavitaves, and Eufalconimorphae); J14, P15, K21, S24, W24.
- Pneumatic foramen in humeral pneumotricipital fossa (char. 109: 0 > 1, reversed in *Podilymbus*, Ralloidea, Charadriiformes, *Gavia*, *Spheniscus*, *Oceanites*, *Pagodroma*, *Leucocarbo*, *Urocolius*, and *Acanthisitta*); J14, P15, K21, S24, W24.
- Deltopectoral crest a small eminence, extending less than one third the total length of the humerus (char. 122: 0 > 1, reversed in *Corythaeola*, *Ardeotis*, *Nyctibius*, *Podargus*, *Spheniscus*, *Leptoptilos*, *Leucocarbo*, *Eudocimus*, *Scopus*, Accipitrimorphae, Strigiformes, Cavitaves, *Micrastur*, and Passeri); J14, P15, K21, S24, W24.
- Radial depression on ulna weak (char. 162: 1 > 0, reversed in *Chauna*, *Dendrocygna*, *Podilymbus*, *Corythaeola*, *Streptoprocne*, *Charadrius*, *Limosa*, *Alca* + *Sterna*, *Phaethon*, *Leptoptilos*, *Fregata*, *Pelecanus*, *Pandion* + *Elanus*, *Leptosomus*, *Bucorvus*, Picodynastornithes, *Psittacus*, and *Climacteris*); J14, P15, K21, S24, W24.
- Minor metacarpal strongly bowed (char. 174: 0 > 1, reversed in *Dendrocygna*, Mirandornithes, Apodiformes, Charadriiformes, *Phaethon*, *Gavia*, Procellariimorphae, *Fregata*, *Eudocimus*, *Pandion*, Picodynastornithes, and Psittacopasseres); J14, P15, S24, W24.
- Extensor process on carpometacarpus surpasses distal articular facet for alular digit by half the width of the facet or more (char. 182: 1 > 2, reversed in Mirandornithes, *Gavia*, and *Spheniscus*); J14, P15, K21, S24, W24.

However, many of these features are absent in the extinct Lithornithidae, which have been recovered as stem-paleognaths (Nesbitt and Clarke, 2016; Worthy et al., 2017) or near-crown stem-birds (Livezey and Zusi, 2007; Hartman et al., 2019) in recent phylogenetic analyses. Based on previous descriptions (Houde, 1988; Bourdon and Lindow, 2015; Nesbitt and Clarke, 2016; Mayr and Kitchener, 2025), lithornithids typically lack sternal incisures (char. 49: 0) and have a deep, cup-like scapular cotyle of the coracoid (char. 66: 0), a deep impression of the m. sternocoracoidei on the coracoid (char. 76: 1), a sharply hooked lateral process of the coracoid (char. 80: 1), a well-developed deltopectoral crest (char. 122: 0), and a weakly bowed minor metacarpal (char. 174: 0). This distribution of character states might reflect a position of lithornithids outside of crown-group birds, but it is also plausible that character optimization at this node may be confounded by convergent acquisition of character states in tinamous (Tinamidae) and galliforms, perhaps correlated with specialization in these groups for burst flight (Houde, 1988; Widrig et al., 2023). Based on comparisons with fossil crown birds and near-crown stem-birds, Mayr (2021a) previously inferred that a deep, cup-like scapular cotyle is likely plesiomorphic for crown birds, despite the presence of a shallow or flattened facet in extant tinamids and galliforms. Of the above characters, lithornithids do exhibit a reduced procoracoid process of the coracoid, ulnae 40% or less the combined length of the humerus, ulna, and carpometacarpus, a pneumatic foramen in the pneumotricipital fossa of the humerus, and a well-developed extensor process on the carpometacarpus, the last of which has previously been identified as a synapomorphy of ornithuran avialans crownward of *Ichthyornis* (Clarke, 2004).

Other pectoral and forelimb features that have been considered potential synapomorphies of crown birds include pneumatic pores in the intercostal incisures of the sternum and a pneumatized coracoid (Clarke, 2004; Turner et al., 2012). However, minute and possibly pneumatic pores have subsequently been identified in the intercostal incisures of *Ichthyornis* (Benito et al., 2022a), and as a result, this character state was not optimized as a synapomorphy of Neornithes in our analyses. Coracoid pneumatization was not coded as a single character in the present study, as the positioning of pneumatic foramina in the coracoid differs among crown birds with pneumatic coracoids. In lithornithids (Houde, 1988) and tinamids (Bertelli et al., 2014; Suzuki et al., 2014; Widrig et al., 2023) the coracoid is dorsally penetrated by one or more pneumatic foramina near the scapular cotyle, whereas in many galloanserans and some neoavians the pneumatic foramen is positioned near the sternal facet.

#### Neognathae

Six character states were optimized as potential synapomorphies of Neognathae, of which four were consistently inferred as such across all alternative molecular topologies studied.

- Median trabecula of sternum squared off at caudal termination (char. 59: 0 > 1, reversed in *Ortalis* + *Rollulus*, *Monias*, *Columba*, *Florisuga*, Gruiformes, *Turnix*, *Alca*, *Eurypyga*, Aequornithes, Accipitrimorphae, *Tyto*, *Urocolius*, *Bucorvus*, *Bucco*, Psittaciformes, and *Menura)*; J14, P15, K21, S24, W24.
- Length of coracoid between three and four times the width of articular facet for sternum (char. 72: 1 > 2, reversed in *Anseranas*, *Phoenicopterus*, *Corythaeola*, *Nyctibius*, Grues, Charadrii, *Limosa*, *Phaethon*, *Gavia*, Procellariiformes, *Fregata*, *Eudocimus*, and Accipitriformes); J14, P15, K21, S24, W24.
- Olecranon fossa on humerus deep (char. 127: 0 > 1, reversed in *Anseranas*, Mirandornithes, *Monias*, *Ardeotis*, *Nyctibius*, *Opisthocomus*, Grues, *Alca*, *Gavia*, *Leptoptilos*, and *Eudocimus*); J14, P15, W24.
- Tendinal sulcus of m. scapulotricipitalis on humerus present (char. 128: 0 > 1, reversed in *Nyctibius*, *Podargus*, *Phoebastria*, *Pagodroma*, and *Menura*); J14, P15, K21, S24, W24.
- Proximal third of minor metacarpal more than half as thick of major metacarpal dorsoventrally (char. 179: 0 > 1, reversed in *Anseranas*, *Ardeotis*, *Nyctibius*, *Podargus*, Apodiformes, *Sula*, and *Pelecanus*); J14, P15, K21, S24, W24.
- Extensor process on carpometacarpus surpasses distal articular facet for alular digit by the width of the facet or more (char. 182: 2 > 3, reversed in Mirandornithes, *Monias*, *Corythaeola*, *Opisthocomus*, *Charadrius*, *Phaethon*, *Gavia*, *Spheniscus*, Bucerotiformes, *Nestor*, and *Climacteris*); J14, P15, W24.

In lithornithids, the extensor process on the carpometacarpus surpasses the distal articular facet for the alular digit by approximately the width of the facet (Nesbitt and Clarke, 2016), so this character may be synapomorphic for a more inclusive clade than Neognathae.

Although a moderately elongate coracoid is found in stem-galliforms (Mourer-Chauviré, 1992a; Mourer-Chauviré, 2000; Mayr and Weidig, 2004; Mayr and Kitchener, 2023a) and the putative stem-anseriform (Houde et al., 2023) or stem-galliform (Mayr et al., 2023a) *Danielsavis*, stockier coracoids (in which the total length of the bone is only two to three times the width of the sternal facet) are present in *Anachronornis* (Houde et al., 2023), *Nettapterornis* (Olson, 1999), and Presbyornithidae (Howard, 1955; Ericson, 1999; Worthy et al., 2023), which are potential stem-anseriforms (Tambussi et al., 2019; Field et al., 2020; Houde et al., 2023; Crane et al., 2024). A similarly stocky coracoid is found in *Juncitarsus* (Ericson, 1999), a putative stem member of Mirandornithes (Mayr, 2014b), which is a clade that has been recovered as the sister group to all other neoavians in some molecular analyses (Braun and Kimball, 2021; Kuhl et al., 2021; Stiller et al., 2024). Therefore, a more elongate coracoid may have been convergently acquired by several neognath clades.

The presence of a tendinal sulcus on the humerus for the m. scapulotricipitalis as a synapomorphy of Neognathae is consistent with previously described fossil evidence, as this feature is absent in most near-crown stem-birds (Clarke, 2004; Clarke et al., 2006; Wang et al., 2016) and lithornithids (Nesbitt and Clarke, 2016), whereas it is present (though inconspicuous) in *Anachronornis* (Houde et al., 2023), presbyornithids (Ericson, 2000; Worthy et al., 2023), and the stem-galliform *Paraortygoides messelensis* (Mayr, 2000a). However, this sulcus is lacking in *Danielsavis* (Houde et al., 2023). A deep olecranon fossa is similarly absent in near-crown stem-birds (Clarke, 2004; Wang et al., 2016) but present in *Anachronornis* (Houde et al., 2023), presbyornithids (Worthy et al., 2023), and the possible stem-anseriform *Conflicto* (Tambussi et al., 2019). This character is absent in Mirandornithes, however, and was therefore not optimized as a neognath synapomorphy under the topologies of Kuhl et al. (2021) and Stiller et al. (2024).

#### Galloanserae

Two character states were optimized as potential synapomorphies of Galloanserae, of which one was consistently inferred as such across all alternative molecular topologies studied.

- Large pneumatic foramen in impression of m. sternocoracoidei in coracoid (char. 77: 0 > 1, reversed in *Dendrocygna*; also found in *Pterocles*, *Opisthocomus*, *Psophia*, *Balearica*, and *Phoebastria*); J14, P15, K21, S24, W24.
- Extensor process on carpometacarpus surpasses distal articular facet for alular digit by more than the width of the facet (char. 182: 3 > 4, reversed in *Dendrocygna*; also found in *Pterocles*, Strisores, Gruoidea, *Sarothrura*, *Burhinus*, Scolopaci, *Sterna*, *Leptoptilos*, *Fregata*, *Scopus*, *Tigrisoma*, Accipitrimorphae, Cavitaves, *Cariama*, *Micrastur*, *Psittacus*, and Tyranni); J14, S24, K21.

Previously reported morphological synapomorphies of Galloanserae have been largely limited to features of the skull (Cracraft and Clarke, 2001; Mayr, 2011a; Field et al., 2020; Crane et al., 2024), so the identification of potential pectoral and forelimb synapomorphies is noteworthy. A large pneumatic foramen in the impression of the m. sternocoracoidei is present in non-anatid crown anseriforms, many crown galliforms, *Nettapterornis* (Olson 1999), and the stem-galliform *Ameripodius* (Mourer-Chauviré, 2000), but is absent in *Anachronornis*, *Danielsavis* (Houde et al., 2023), presbyornithids (Howard, 1955; Worthy et al., 2023), and the stem-galliforms *Gallinuloides* (Mayr and Weidig, 2004), *Paraortygoides* (Mayr, 2006a; Mayr and Kitchener, 2023a), *Waltonortyx* (Mayr and Kitchener, 2023a), *Paraortyx*, and *Quercymegapodius* (Mourer-Chauviré, 1992a). Whether this character was present in the last common ancestor of Galloanserae is therefore uncertain. However, an evolutionary tendency towards the formation of a pneumatic foramen near the sternal end of the coracoid may plausibly be an underlying synapomorphy of Galloanserae, as suggested by Mayr (2006a).

That galloanserans ancestrally possessed an extensor process on the carpometacarpus that surpasses the distal articular facet for the alular digit by more than the width of the facet is supported by the morphology of *Anachronornis* (Houde et al., 2023), *Nettapterornis* (Olson, 1999), *Conflicto* (Tambussi et al., 2019), presbyornithids (Howard, 1955), and stem-galliforms (Mourer-Chauviré, 1992a; Mayr, 2000a; Mourer-Chauviré, 2000; Mayr and Weidig, 2004), though *Danielsavis* appears to have a relatively low extensor process (Houde et al., 2023). Whether or not this feature is optimized as a synapomorphy of Galloanserae is sensitive to the internal topology of Neoaves.

The presence of several prominent muscle scars that diagonally traverse the dorsal surface of the coracoid may be a synapomorphy of Galloanserae, as this feature is found in most crown anseriforms, presbyornithids (Ericson, 1997), *Danielsavis* (Houde et al., 2023), *Gallinuloides* (Mayr and Weidig, 2004), *Paraortygoides* (Mayr, 2006a), *Waltonortyx* (Mayr and Kitchener, 2023a), and *Ameripodius* (Mourer-Chauviré, 2000). Although this character was included in the present study (char. 74: 1), the extant-only taxon sampling for crown birds prevented it from being optimized as a galloanseran synapomorphy, as it is absent in most crown galliforms.

#### Neoaves

Seven character states were optimized as potential synapomorphies of Neoaves, but none of these were consistently inferred as such across all alternative molecular topologies studied. No unambiguous synapomorphies for this clade were inferred under the Stiller et al. (2024) topology.

- Ventrolateral sulcus on sternum distinct (char. 34: 0 > 1, reversed in Mirandornithes, *Caprimulgus*, *Aegotheles*, *Sterna*, Gruoidea, *Eurypyga*, *Phoebastria*, *Leptoptilos*, Pelecaniformes, *Bucorvus*, and *Nestor*; also found in *Rollulus*); W24.
- Coracoid keeled at ventromedial margin of supracoracoid sulcus (char. 69: 0 > 1, reversed in Mirandornithes, *Pterocles*, *Ardeotis*, *Tapera*, *Aegotheles*, *Streptoprocne*, Gruiformes, Aequornithes, *Bucorvus*, *Nestor*, and *Acanthisitta*; also found in *Dendrocygn*a and *Ortalis* + *Rollulus*); P15, W24.
- Little expansion of external lip of coracoid (char. 84: 1 > 0, reversed in Columbimorphae, *Ardeotis*, *Tapera*, Grues, *Rostratula*, *Alca*, *Sterna*, *Oceanites*, Suliformes, *Scopus*, *Pelecanus*, *Coragyps*, *Pandion*, *Leptosomus*, *Upupa*, Pici, and *Nestor*); K21.
- Prominent expansion of internal lip of coracoid (char. 86: 0 > 1, reversed in *Podilymbus*, *Pterocles*, *Caprimulgus*, *Psophia*, Ralloidea, *Charadrius*, *Rostratula*, *Turnix*, and *Urocolius*; also found in *Anseranas* and *Alectura*); J14, W24.
- Coracobrachial impression on humerus prominent (char. 114: 0 > 1, reversed in Mirandornithes, *Monias*, *Tapera*, Apodiformes, *Sarothrura*, *Rallus*, *Charadrius*, *Limosa*, *Alca*, *Gavia*, *Oceanites* + *Pagodroma*, Pelecanimorphae, *Pandion* + *Elanus*, *Urocolius*, Pici, *Nestor*, and *Acanthisitta*; also found in *Anseranas* and *Ortalis*); P15, W24.
- Distal radius strongly curved dorsoventrally (char. 147: 0 > 1, reversed in *Phoenicopterus*, *Ardeotis*, Apodiformes, *Psophia*, *Balearica*, *Rallus*, *Phaethon*, Aequornithes, *Coragyps*, *Ninox*, *Trogon*, *Bucorvus*, *Cariama*, *Nestor*, and *Acanthisitta*; also found in *Anseranas* and *Ortalis* + *Rollulus*); J14, P15, W24.
- Internal index on phalanx 1 of major digit well developed (char. 201: 0 > 1, reversed in *Podilymbus*, *Monias*, *Corythaeola*, *Opisthocomus*, Gruiformes, *Turnix*, *Spheniscus*, *Tigrisoma*, *Ninox*, *Urocolius*, *Bucorvus*, *Merops*, Piciformes, and Australaves; also found in *Ichthyornis* and Anseres); J14.

Neoavians exhibit a great degree of anatomical disparity, and morphological synapomorphies characterizing the entire clade have therefore been challenging to identify, with previously proposed synapomorphies being largely limited to cranial and soft tissue traits (Livzey and Zusi, 2007; Mayr, 2008a; Mayr, 2011a). The characters listed above are highly variable within Neoaves and can also be found in non-neoavian birds, so their identification as potential synapomorphies should be considered tentative.

#### Mirandornithes

22 character states were optimized as potential synapomorphies of Mirandornithes, of which four were consistently inferred as such across all alternative molecular topologies studied.

- Strong craniocaudal curvature of furcula (char. 2: 0 > 1, also found in Anseres, *Nyctibius*, Charadriiformes, Aequornithes, *Pandion*, *Trogon*, and *Coracias* + *Merops*); J14, K21, S24, W24.
- Ventrolateral sulcus on sternum indistinct (char. 34: 1 > 0, also found in non-neoavian birds, *Caprimulgus*, *Aegotheles*, Gruoidea, *Sterna*, *Eurypyga*, *Phoebastria*, *Leptoptilos*, Pelecaniformes, *Bucorvus*, and *Nestor*); P15, W24.
- Pneumatic pores in intercostal incisures of sternum absent (char. 37: 1 > 0, also found in *Rollulus*, *Caprimulgus*, *Aegotheles*, Heliornithes, Charadriiformes, *Gavia*, *Spheniscus*, *Oceanites*, *Pagodroma*, *Leucocarbo*, *Urocolius*, Piciformes, *Neopelma*, and *Menura*); J14, K21, S24, W24.
- Caudolateral trabeculae of sternum extend caudal of median trabecula (char. 60: 0 > 1, also found in *Gansus*, *Ichthyornis*, *Dendrocygna*, Strisores, Heliornithes, *Phaethon*, *Spheniscus*, *Phoebastria*, Suliformes, *Scopus*, *Pelecanus*, *Pandion* + *Elanus*, *Tyto*, Pici, *Micrastur*, and *Nestor*); J14, P15, K21, S24, W24.
- Impression for acrocoracohumeral ligament on coracoid deep (char. 64: 0 > 1, also found in *Eudromia*, *Chauna*, *Dendrocygna*, *Streptoprocne*, *Rallus*, *Rostratula*, *Turnix*, *Pagodroma*, *Leptoptilos*, Suloidea, *Eudocimus*, *Pelecanus*, Accipitrimorphae, Strigiformes, *Urocolius*, Eucavitaves, and Psittacopasseres); J14, P15, W24.
- Procoracoid process of coracoid modest or reduced (char. 68: 2 > 1, also found in Anseres, Galliformes, *Monias*, *Ardeotis*, *Caprimulgus*, *Nyctibius*, Apodiformes, *Sarothrura*, *Rallus*, *Eurypyga*, *Gavia*, *Phoebastria*, *Leucocarbo*, Pelecaniformes, Accipitrimorphae, *Urocolius*, *Trogon*, Pici, *Nestor*, *Acanthisitta*, and Passeri); P15, W24.
- Coracoid rounded and relatively thick at ventromedial margin of supracoracoid sulcus (char. 69: 1 > 0, also found in non-neoavian birds, *Pterocles*, *Ardeotis*, *Tapera*, *Aegotheles*, *Streptoprocne*, Gruiformes, Aequornithes, *Bucorvus*, *Nestor*, and *Acanthisitta*); P15, W24.
- Strong lateral curvature of scapular shaft (char. 96: 0 > 1, also found in *Ichthyornis*, Anseriformes, *Charadrius*, *Limosa*, *Eurypyga*, Aequornithes, and *Leptosomus*); J14.
- Dorsal tubercle of humerus pointed (char. 105: 0 > 1, also found in Anseriformes, *Nyctibius*, *Florisuga*, *Aramus*, Ralloidea, Phaethoquornithes, *Pandion*, Strigiformes, *Trogon*, *Upupa*, *Coracias*, and Passeri); J14, K21, S24.
- Coracobrachial impression on humerus absent or weak (char. 114: 1 > 0, also found in non-neoavian birds, *Monias*, *Tapera*, Apodiformes, *Sarothrura*, *Rallus*, *Charadrius*, *Limosa*, *Alca*, *Gavia*, *Oceanites* + *Pagodroma*, Pelecanimorphae, *Pandion* + *Elanus*, *Urocolius*, Pici, *Nestor*, and *Acanthisitta*); P15, W24.
- Deltopectoral crest rounded dorsally (char. 121: 1 > 0, also found in *Gansus*, *Chauna*, *Anseranas*, *Ardeotis*, *Aegotheles*, *Opisthocomus*, Gruiformes, *Charadrius*, *Rostratula*, *Alca*, *Eudocimus*, Eucavitaves, and Australaves); J14, P15, K21, S24.
- Humeral shaft straight (char. 125: 0 > 1, also found in *Pterocles*, *Columba*, Apodiformes, *Opisthocomus*, Charadriiformes, *Phaethon*, Procellariimorphae, *Fregata*, *Leucocarbo*, *Urocolius*, *Psilopogon*, *Psittacus*, *Neopelma*, and Passeri); K21, S24.
- Olecranon fossa on humerus shallow (char. 127: 1 > 0, also found in non-neognath birds, *Anseranas*, *Monias*, *Ardeotis*, *Nyctibius*, *Opisthocomus*, Grues, *Alca*, *Gavia*, *Leptoptilos*, and *Eudocimus*); J14, P15.
- Proximal margin of dorsal condyle on humerus pointed (char. 139: 0 > 1, also found in *Dendrocygna*, *Podargus*, *Aegotheles*, *Balearica*, *Charadrius*, *Alca*, Procellariiformes, *Leptoptilos*, Suloidea, *Scopus* + *Pelecanus*, Accipitrimorphae, *Leptosomus*, *Trogon*, *Coracias*, *Micrastur*, *Psittacus*, and Eupasseres); J14, P15, K21, S24, W24.
- Proximal margin of dorsal condyle on humerus separated from brachial fossa by smooth area of bone (char. 140: 1 > 0, also found in *Gansus*, *Eudromia*, Anseres, *Ardeotis*, *Nyctibius*, *Podargus*, *Aramus*, *Burhinus*, *Alca*, *Phaethon*, Aequornithes, Strigiformes, *Leptosomus*, *Trogon*, *Bucorvus*, and *Coracias*); J14.
- Radial incisure on ulna prominent (char. 156: 0 > 1, also found in *Ichthyornis*, *Eudromia*, *Chauna*, *Anseranas*, Mirandornithes, *Ardeotis*, *Nyctibius*, Grues, Charadriiformes, *Phaethon*, *Phoebastria*, *Pagodroma*, *Leptoptilos*, *Fregata*, *Sula*, *Scopus* + *Pelecanus*, Accipitrimorphae, and Psittaciformes); W24.
- Scapulotricipital impression on ulna deep (char. 157: 0 > 1, also found in Columbimorphae, Strisores, *Rallus*, Charadriiformes, *Phaethon*, *Gavia*, Procellariiformes, Suliformes, *Eudocimus*, *Scopus*, *Pandion*, Strigiformes, *Trogon*, *Bucco*, *Psilopogon*, and Psittacopasseres); S24.
- Minor metacarpal weakly bowed (char. 174: 1 > 0, also found in *Gansus*, *Ichthyornis*, *Dendrocygna*, Apodiformes, Charadriiformes, *Phaethon*, *Gavia*, Procellariimorphae, *Fregata*, *Eudocimus*, *Pandion*, Picodynastornithes, and Psittacopasseres); J14, S24.
- Minor metacarpal prominently narrows distally in caudal view (char. 180: 0 > 1, also found in *Ichthyornis*, Anseres, Columbimorphae, *Aegotheles*, Gruiformes, Charadriiformes, Phaethontimorphae, Pelecanimorphae, *Leptosomus*, *Coracias*, Accipitrimorphae, Strigiformes, and Australaves); W24.
- Extensor process on carpometacarpus slightly surpasses distal articular facet for alular digit (Fig. 23, char. 182: 2 > 1, also found in *Ichthyornis*); J14, P15, K21, S24, W24.
- Distal synostosis of major and minor metacarpals longer than craniocaudal width (char. 198: 0 > 1, also found in *Ichthyornis*, *Eudromia*, Anseriformes, *Monias*, *Psophia*, Ralloidea, *Charadrius*, Scolopaci, *Sterna*, *Gavia*, *Eudocimus*, and *Cariama*); W24.
- Phalanx 1 of major digit elongate (char. 202: 0 > 1, also found in *Fregata*); J14, P15, K21, S24, W24.

**Figure 23.**
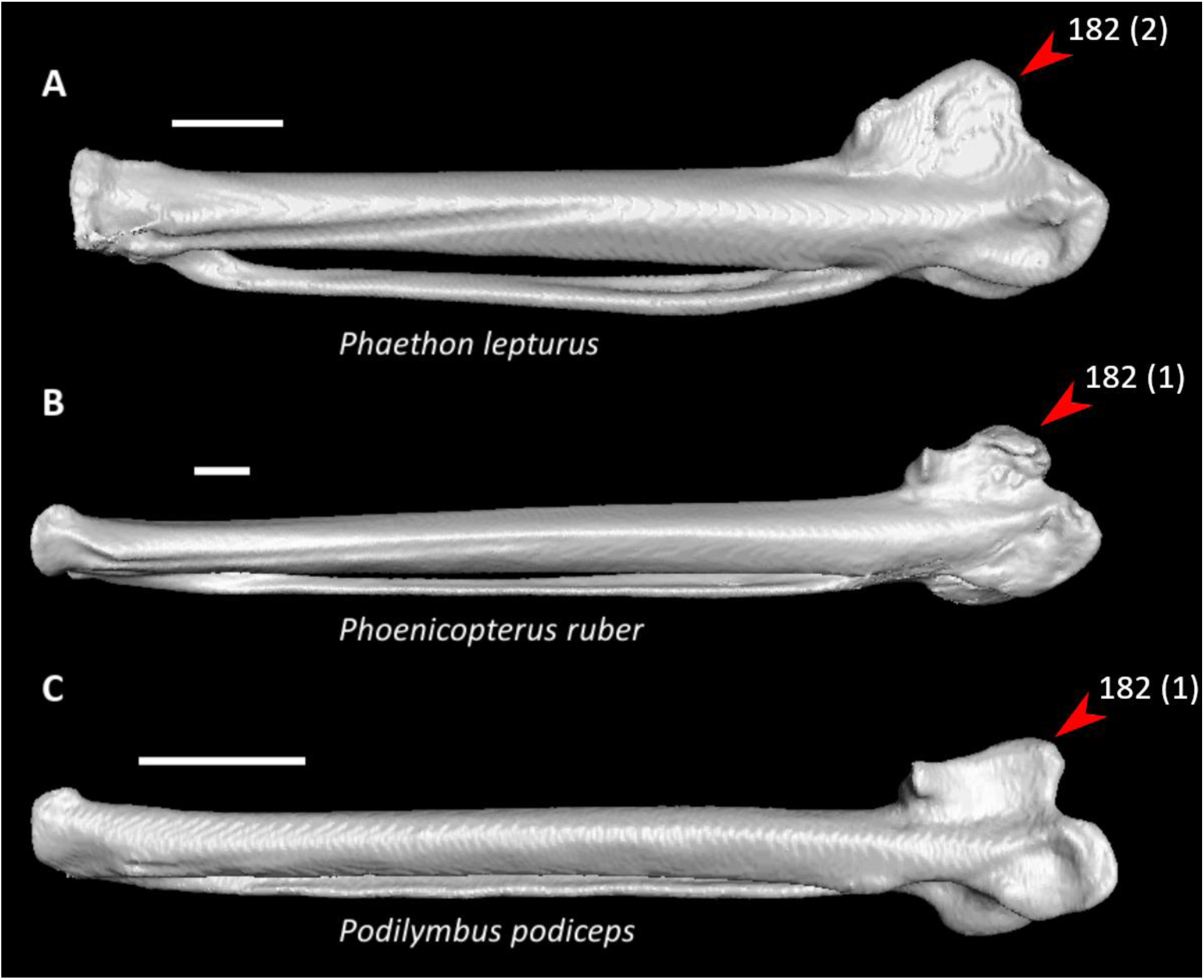
Carpometacarpi of *Phaethon lepturus* (**A**, NHMUK 1876.3.16.3, left element), *Phoenicopterus ruber* (**B**, UMZC 346.B, left element), and *Podilymbus podiceps* (**C**, UMMZ 205087, right element, mirrored) in dorsal view. Scale bars = 5 mm. Arrows indicate the extensor process. *Phaethon* exhibits state 2 for character 182 (extensor process surpassing the distal articular facet for alular digit by roughly half the width of the facet), whereas *Phoenicopterus* and *Podilymbus* exhibit state 1 (low extensor process on the carpometacarpus only slightly surpassing the distal articular facet, optimized as a synapomorphy of Mirandornithes).

That only a few of these characters were recovered as synapomorphies across all alternative topologies indicates that their optimization is highly sensitive to the placement of Mirandornithes within Neoaves. An elongate phalanx 1 of the major digit was previously considered a synapomorphy of Mirandornithes by Mayr (2004a). Another mirandornithean synapomorphy identified by Mayr (2004a) was the presence of an oval depression on the humerus at the insertion site of the m. scapulohumeralis cranialis. This character state was included in the present study (char. 108: 1), but it was not optimized as a synapomorphy of this clade because it happens to be absent or indistinct in *Phoenicopterus ruber* (Mayr, 2004a; Mayr, 2014b), the species of flamingo (Phoenicopteridae) used in our dataset. In general, this feature is less prominent in extant phoenicopterids than in grebes (Podicipedidae), stem-phoenicopterids, and *Juncitarsus* (Mayr, 2014b), though a distinct depression has been reported in *Phoenicopterus chilensis* (Mayr, 2004a). The recognition of sternal caudolateral trabeculae that caudally surpass the median trabecula, a rounded deltopectoral crest, and an extensor process of the carpometacarpus only slightly surpassing the distal articular facet for the alular digit as potential synapomorphies of Mirandornithes is, to our knowledge, novel.

Most of the above characters cannot be assessed in published descriptions of *Juncitarsus* (Olson and Feduccia, 1980; Peters, 1987; Ericson, 1999; Mayr, 2014b), though Mayr (2014b) noted that *Juncitarsus* lacks an elongate phalanx 1 of the major digit and interpreted this as evidence supporting its placement outside of crown-group Mirandornithes.

#### Columbimorphae

Eleven character states were optimized as potential synapomorphies of Columbimorphae, but none of these were consistently inferred as such across all alternative molecular topologies studied. Two character states were optimized as potential synapomorphies under all molecular reference topologies except Kuhl et al. (2021), in which cuckoos (Cuculidae) are included in this clade.

- Pair of cranially extending flanges absent from dorsal lip of coracoid sulci on sternum (char. 22: 1 > 0, also found in *Ichthyornis*, *Eudromia*, *Alectura*, *Ortalis*, Apodiformes, *Phoebastria*, *Fregata*, *Urocolius*, and Psittaciformes); P15, S24, W24.
- Cranial extent of lateral fenestrae greater than one-third of total sternum length (char. 50: 0 > 1, also found in *Eudromia*, *Rollulus*, *Caprimulgus*, Letornithes, *Sarothrura*, *Rallus*, *Limosa*, *Turnix*, *Gavia*, *Spheniscus*, *Urocolius*, Piciformes, and *Acanthisitta*); J14, P15, S24, W24.
- Transverse expansion on caudolateral trabecula of sternum (char. 52: 0 > 1, also found in *Rollulus*, *Podilymbus*, *Tapera*, *Aegotheles*, *Opisthocomus*, *Turnix*, *Coragyps*, *Ninox*, *Urocolius*, Eucavitaves, *Acanthisitta*, *Erythropitta*, and *Climacteris*); P15, K21, S24.
- Lateral extent of lateral process on coracoid half the width of sternal facet or greater (char. 79: 0 > 1, also found in *Corythaeola*, *Tapera*, *Caprimulgus*, *Podargus*, Charadriiformes, *Phoebastria*, *Pagodroma*, *Fregata*, *Sula*, *Scopus*, *Pelecanus*, *Elanus*, *Tyto*, *Leptosomus*, *Upupa*, *Coracias* + *Merops*, and *Bucco*); J14.
- Ventral tubercle on humerus largely concealing pneumotricipital fossa (Fig. 24, char. 103: 0 > 1, also found in *Gansus*, *Florisuga*, *Turnix*, *Alca*, *Spheniscus*, *Pagodroma*, *Urocolius*, *Alcedo*, Piciformes, *Micrastur*, *Neopelma*, and *Climacteris*); J14, P15, S24, W24.
- Capital shaft ridge on humerus absent (char. 106: 1 > 0, also found in *Eudromia*, Galliformes, *Corythaeola*, *Caprimulgus*, Daedalornithes, *Opisthocomus*, Gruoidea, *Alca*, *Eurypyga*, *Spheniscus*, *Oceanites*, *Leptoptilos*, *Eudocimus*, *Tigrisoma*, *Urocolius*, *Cariama*, *Micrastur*, and *Acanthisitta*); J14, W24.
- Deltopectoral crest caudally convex (char. 124: 0 > 1, also found in *Eudromia*, Galliformes, *Podilymbus*, *Ardeotis*, Daedalornithes, Ralloidea, *Rostratula*, *Turnix*, *Alca*, *Spheniscus*, *Trogon*, *Alcedo*, Psittaciformes, and *Acanthisitta*); P15, S24, W24.
- Brachial fossa on humerus shallow (char. 134: 1 > 0, also found in *Gansus*, *Eudromia*, Galliformes, *Podilymbus*, *Tapera*, *Sarothrura*, *Rallus*, *Rostratula*, *Turnix*, *Alca*, *Eurypyga*, *Phoebastria*, *Oceanites*, *Tigrisoma*, *Elanus*, *Ninox*, *Upupa*, *Alcedo*, *Cariama*, *Psittacus*, and *Erythropitta*); P15, K21.
- Scapulotricipital impression on ulna deep (char. 157: 0 > 1, also found in Mirandornithes, Strisores, *Rallus*, Charadriiformes, *Phaethon*, *Gavia*, Procellariiformes, Suliformes, *Eudocimus*, *Scopus*, *Pandion*, Strigiformes, *Trogon*, *Bucco*, *Psilopogon*, and Psittacopasseres); P15, S24.
- Minor metacarpal prominently narrows distally in caudal view (char. 180: 0 > 1, also found in *Ichthyornis*, Anseres, Mirandornithes, *Aegotheles*, Gruiformes, Charadriiformes, Phaethontimorphae, Pelecanimorphae, *Leptosomus*, *Coracias*, Accipitrimorphae, Strigiformes, and Australaves); W24.
- Cranial projection on distal end of major metacarpal weak (char. 197: 1 > 0, also found in *Gansus*, *Ortalis* + *Rollulus*, *Corythaeola*, *Tapera*, *Opisthocomus*, *Psophia*, *Aramus*, Heliornithes, *Rostratula*, *Turnix*, *Spheniscus*, *Tigrisoma*, Strigiformes, *Upupa*, *Merops*, *Bucco*, *Cariama*, and Passeriformes); J14.

**Figure 24.**
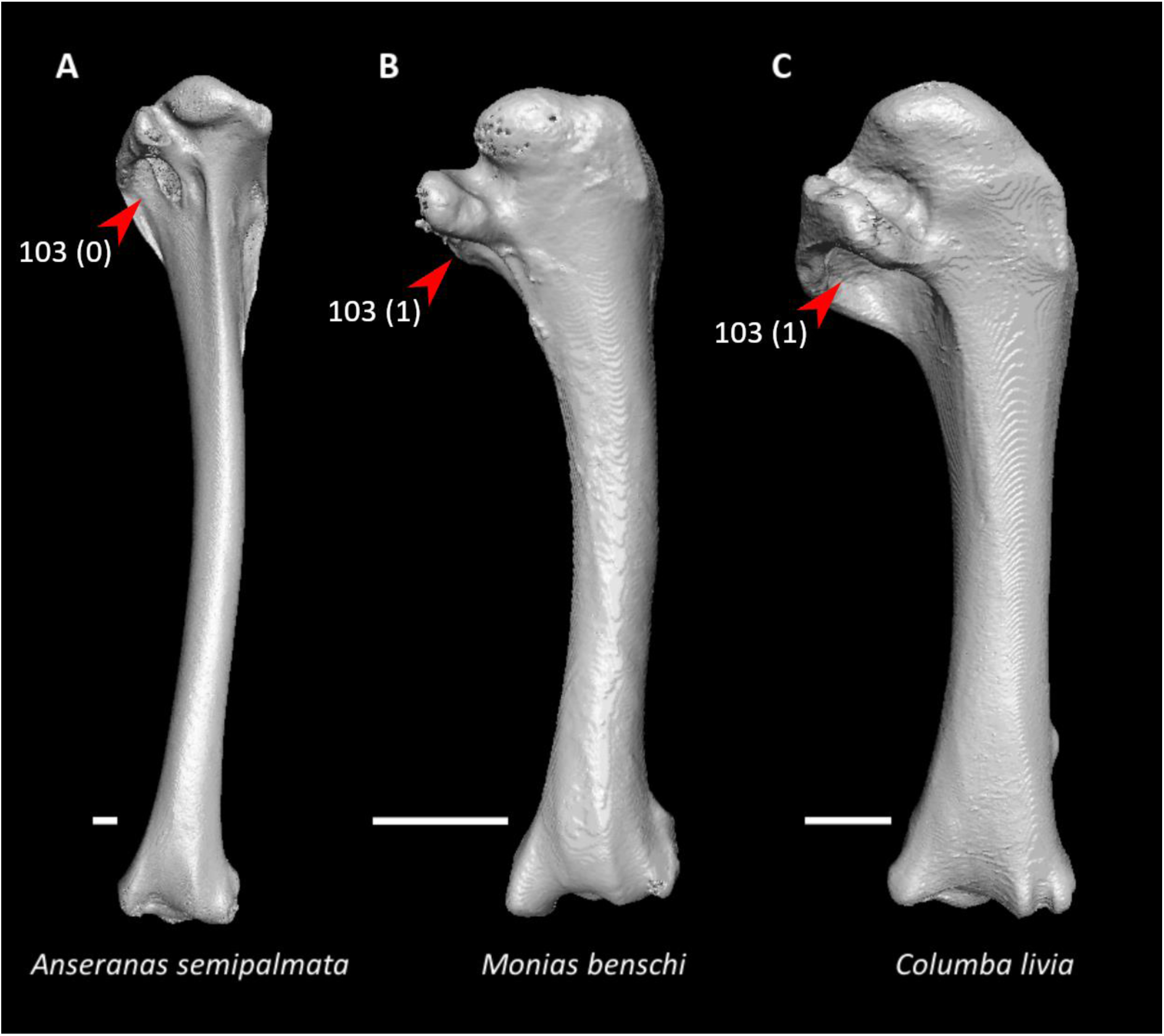
Humeri of *Anseranas semipalmata* (**A**, NHMUK 1852.7.22.1, left element, mirrored), *Monias benschi* (**B**, NHMUK 1924.11, right element), and *Columba livia* (**C**, FMNH 347273, left element, mirrored) in caudal view. Scale bars = 5 mm. Arrows indicate the pneumotricipital fossa. *Anseranas* exhibits state 0 for character 103 (pneumotricipital fossa largely exposed in caudal view), whereas *Monias* and *Columba* exhibit state 1 (pneumotricipital fossa largely concealed by the ventral tubercle in caudal view, optimized as a synapomorphy of Columbimorphae).

Using the phylogenetic matrix of Livezey and Zusi (2006), Sangster et al. (2022) also inferred a proximodistally elongated dorsal tubercle on the humerus as a potential synapomorphy of this clade. The fossil record of all potential columbimorph groups is extremely poorly known, and few stem members of the major constituent clades have been convincingly identified (Mayr, 2017b; Mayr, 2022a). The stem-sandgrouse (stem-Pteroclidae) *Archaeoganga larvatus* and *Leptoganga* do, however, exhibit a ventral tubercle that largely conceals the pneumotricipital fossa, the absence of a capital ridge shaft, and a shallow brachial fossa on the humerus (Pl. 2 in Mourer-Chauviré, 1993).

#### Pteroclimesites

Eight character states were optimized as potential synapomorphies of Pteroclimesites, of which two were consistently inferred as such across all alternative molecular topologies studied.

- External spine on sternum distinctly tapered (char. 16: 0 > 1, also found in *Ardeotis*, *Caprimulgus*, *Nyctibius*, *Balearica*, *Turnix*, *Phaethon*, *Phoebastria*, *Leptoptilos*, *Tigrisoma*, *Ninox*, *Urocolius*, *Alcedo*, Piciformes, and Passeriformes); J14, P15, S24, W24.
- Keel sulcus present on sternum (char. 39: 0 > 1, also found in *Eudromia*, *Dendrocygna*, Galliformes, *Phoenicopterus*, *Tapera*, *Psophia*, *Rallus*, *Charadrius*, *Limosa*, *Eurypyga*, *Pagodroma*, *Leptoptilos*, *Coragyps*, and *Micrastur*); J14, P15, S24, W24.
- Longitudinal concavity of scapula prominent (char. 93: 0 > 1, also found in *Gansus*, *Anseranas*, Galliformes, *Tapera*, *Opisthocomus*, *Aramus*, Heliornithes, *Eudocimus*, *Urocolius*, *Leptosomus*, *Trogon*, Pici, *Cariama*, *Micrastur*, and Tyranni); J14, P15, S24, W24.
- Ventral tubercle on humerus distal to dorsal tubercle (Fig. 25, char. 104: 1 > 0, also found in *Eudromia*, *Dendrocygna*, Galliformes, *Phoenicopterus*, *Florisuga*, *Turnix*, *Alca*, *Gavia*, *Spheniscus*, *Urocolius*, Piciformes, Tyranni, and *Climacteris*); J14, P15, K21, S24, W24.
- Ventral epicondyle on humerus equal or distal to ventral condyle (char. 130: 0 > 1, also found in Galliformes, *Phoenicopterus*, *Tapera*, *Streptoprocne*, *Opisthocomus*, *Psophia*, *Aramus*, *Sarothrura*, *Charadrius*, Scolopaci, *Alca*, *Gavia*, Procellariimorphae, *Fregata*, *Pelecanus* + *Tigrisoma*, *Elanus*, *Ninox*, Coraciimorphae, and Australaves); J14, P15, S24.
- Ventral aponeurosis tubercle on radius rounded (char. 149: 1 > 0, also found in *Nyctibius*, *Streptoprocne*, *Opisthocomus*, Grues, Heliornithes, *Turnix*, *Eurypyga*, *Phoebastria*, *Fregata*, *Elanus*, *Tyto*, *Urocolius*, *Leptosomus*, *Merops*, Piciformes, and Australaves); J14, P15, K21, S24, W24.
- Distinct ridge connecting caudal end of minor metacarpal and pisiform process (char. 173: 0 > 1, also found in *Tapera*, *Psophia*, *Rallus*, *Tyto*, *Leptosomus*, *Upupa*, *Cariama*, *Micrastur*, and *Acanthisitta*); J14, P15, S24, W24.
- Pneumatic foramen in infratrochlear fossa on carpometacarpus absent (char. 191: 1 > 0, also found in Mirandornithes, Strisores, *Opisthocomus*, *Rallus*, Scolopaci + Lari, Procellariimorphae, *Leucocarbo*, *Eudocimus*, *Tigrisoma*, *Ninox*, *Urocolius*, *Merops*, *Jynx*, *Pandion*, *Nestor*, and Passeriformes); K21.

**Figure 25.**
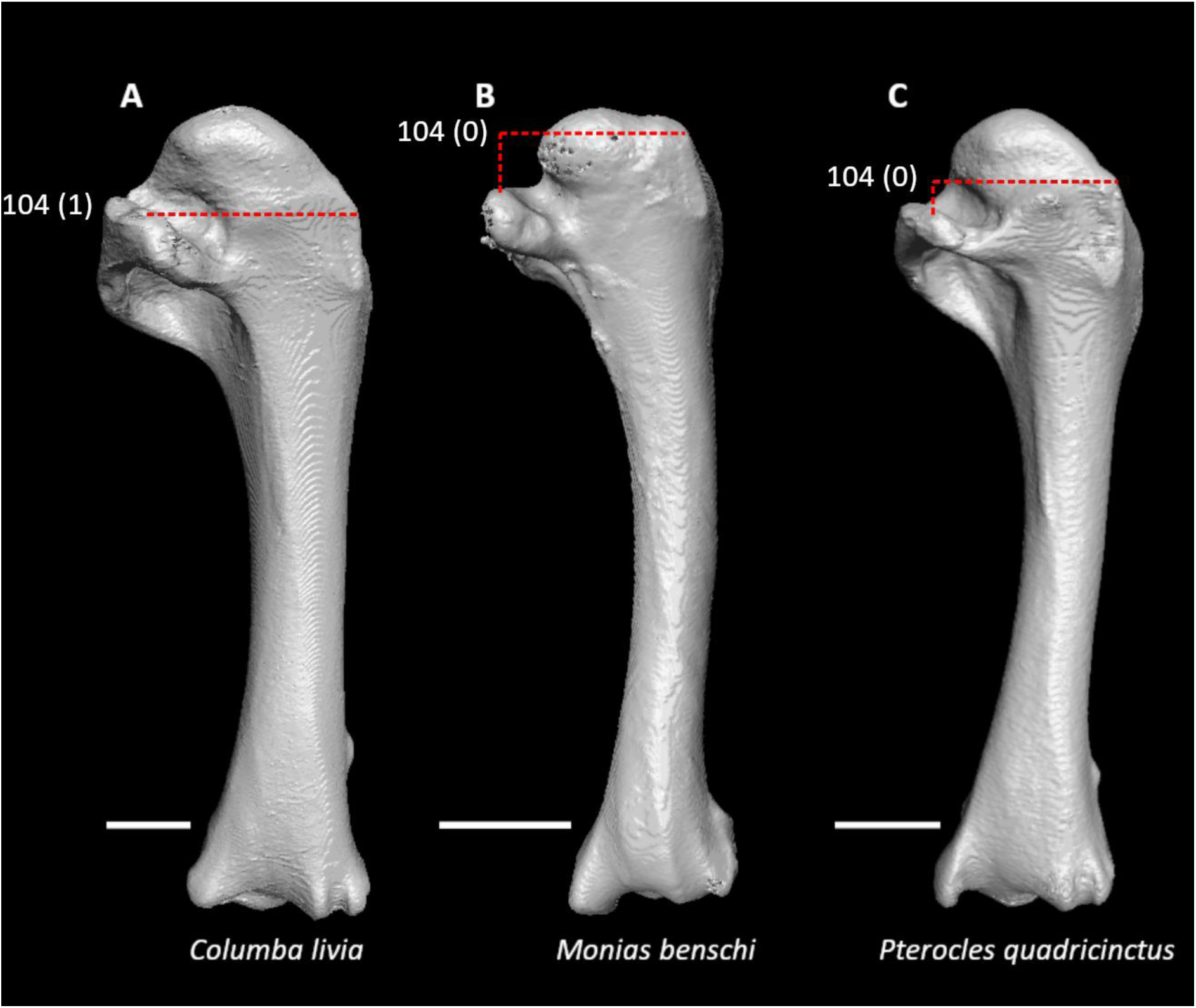
Humeri of *Columba livia* (**A**, FMNH 347273, left element, mirrored), *Monias benschi* (**B**, NHMUK 1924.11, right element), and *Pterocles quadricinctus* (**C**, FMNH 319937, left element, mirrored) in caudal view. Scale bars = 5 mm. Dotted lines indicate the proximal extent of the dorsal tubercle relative to that of the ventral tubercle. *Columba* exhibits state 1 for character 104 (ventral tubercle subequal in elevation to dorsal tubercle), whereas *Monias* and *Pterocles* exhibit state 0 (ventral tubercle distal to dorsal tubercle, optimized as a synapomorphy of Pteroclimesites).

Most of these characters are difficult to assess in previously identified stem-pteroclid fossils based on published descriptions, though *Archaeoganga* and *Leptoganga* do have a ventral tubercle distal to the dorsal tubercle and a ventral epicondyle equal or distal to the ventral condyle on the humerus (Pl. 2 in Mourer-Chauviré, 1993).

Using the phylogenetic matrix of Livezey and Zusi (2006), Sangster et al. (2022) optimized a sternum with a median trabecula equal in length or longer than the caudolateral trabeculae, which in turn are equal in length or longer than the intermediate trabeculae, as a potential synapomorphy of Pteroclimesites. However, this character is not strictly applicable to mesites (Mesitornithidae), which only have a single pair of sternal incisures (and thus lack intermediate trabeculae).

#### Strisores

15 character states were optimized as potential synapomorphies of Strisores, of which three were consistently inferred as such across all alternative molecular topologies studied.

- Hypocleideum less than 5% the total length of the furcula (char. 9: 1 > 0, reversed in *Streptoprocne*; also found in *Podilymbus*, Scolopaci, *Alca*, Phaethoquornithes, *Coragyps*, *Bucorvus*, *Alcedo*, *Cariama*, and *Menura*); J14, P15, K21.
- External spine of sternum less than 5% total sternum length (char. 15: 1 > 0, also found in *Chauna*, *Pterocles*, *Ardeotis*, Gruiformes, *Alca*, *Phaethon*, *Gavia*, *Phoebastria*, *Leptoptilos*, *Leucocarbo*, *Eudocimus*, *Coragyps*, and *Leptosomus*); J14, P15.
- Costal margin less than 25% of total sternum length (char. 35: 1 > 0, reversed in *Streptoprocne*; also found in *Gansus*, *Eudromia*, Galliformes, Mirandornithes, Columbimorphae, *Ardeotis*, *Tapera*, *Sarothrura*, *Burhinus*, *Rostratula*, *Turnix*, *Tigrisoma*, *Ninox*, *Urocolius*, Eucavitaves, and Eupasseres); S24.
- Dorsoventral orientation of acrocoracoid process essentially coplanar with main craniocaudal axis of coracoid (char. 62: 0 > 1, also found in Galliformes, *Pterocles*, *Columba*, *Psophia*, Heliornithes, Charadriiformes, *Urocolius*, Eucavitaves, Tyranni, and *Climacteris*); J14, P15, K21, S24.
- Lateral process of coracoid strongly hooked (char. 80: 0 > 1, reversed in Daedalornithes; also found in *Gansus*, *Ichthyornis*, *Anseranas*, *Rollulus*, *Ardeotis*, *Tapera*, Grues, *Burhinus*, *Sterna*, *Phaethon*, Procellariiformes, *Leptoptilos*, *Fregata*, *Leucocarbo*, *Scopus* + *Pelecanus*, *Pandion*, *Trogon*, *Bucorvus*, *Coracias*, *Bucco*, and *Psittacus*); P15, K21, W24.
- Ulna greater than 40% of humerus + ulna + carpometacarpus length (char. 98: 0 > 1, reversed in Apodiformes; also found in *Gansus*, *Ichthyornis*, *Chauna*, *Phoenicopterus*, *Columba*, *Ardeotis*, Grues, *Burhinus*, *Rostratula*, *Sterna*, Phaethontimorphae, *Phoebastria*, Pelecanimorphae, Accipitrimorphae, Strigiformes, Cavitaves, and Eufalconimorphae); J14, P15, K21, W24.
- Transverse sulcus on humerus intermediate in length (char. 115: 0 > 1, reversed in *Podargus*; also found in *Anseranas*, *Balearica*, *Charadrius*, *Gavia*, *Oceanites* + *Pagodroma*, Pelecanimorphae, *Coragyps*, *Elanus*, *Tyto*, *Coracias*, and Psittaciformes); J14, P15, K21, W24.
- Ventral epicondyle on humerus proximal to ventral condyle (char. 130: 1 > 0, reversed in *Streptoprocne*; also found in *Ichthyornis*, *Eudromia*, Anseriformes, *Podilymbus*, *Columba*, *Corythaeola*, *Ardeotis*, *Balearica*, *Podica*, *Rallus*, *Burhinus*, *Turnix*, *Sterna*, *Phaethon*, *Oceanites*, *Leptoptilos*, Suloidea, *Eudocimus*, *Coragyps*, *Pandion*, *Tyto*, and *Nestor*); S24.
- Radius craniocaudally curved (char. 144: 0 > 1, reversed in Apodiformes; also found in Anseriformes, *Phoenicopterus*, *Ardeotis*, Grues, Charadrii, Feraequornithes, *Coragyps*, *Pandion*, Strigiformes, Cavitaves, *Micrastur*, Tyranni, and *Climacteris*); J14, P15, K21, S24, W24.
- Apex of dorsal cotylar process of ulna approximately coplanar with dorsal surface of ulnar main body (char. 150: 0 > 1, reversed in *Aegotheles* and *Streptoprocne*; also found in *Ichthyornis*, Anseriformes, *Podilymbus*, *Corythaeola*, *Opisthocomus*, Charadriiformes, *Urocolius*, Cavitaves, *Nestor*, and Passeriformes); J14.
- Scapulotricipital impression on ulna deep (char. 157: 0 > 1, reversed in Daedalornithes; also found in Mirandornithes, Columbimorphae, *Rallus*, Charadriiformes, *Phaethon*, *Gavia*, Procellariiformes, Suliformes, *Eudocimus*, *Scopus*, *Pandion*, Strigiformes, *Trogon*, *Bucco*, *Psilopogon*, and Psittacopasseres); J14, P15.
- Ventral collateral ligamental tubercle on ulna strongly developed (char. 158: 0 > 1, also found in *Ichthyornis*, *Ortalis*, *Corythaeola*, *Tapera*, *Psophia*, Charadriiformes, Procellariiformes, *Fregata*, *Pelecanus*, *Elanus*, Strigiformes, Coraciimorphae, *Nestor*, and Passeriformes); P15, K21, S24.
- Proximal termination of carpal trochlea strongly angular in dorsal view (char. 176: 0 > 2, reversed in Apodiformes; also found in *Monias*, *Columba*, *Balearica*, Ralloidea, *Alca*, *Eudocimus*, *Pelecanus*, *Elanus*, *Trogon*, *Coracias*, *Psilopogon*, and *Cariama*); J14, P15, K21, S24, W24.
- Extensor process on carpometacarpus surpasses distal articular facet for alular digit by more than the width of the facet (char. 182: 3 > 4, also found in Galloanserae, *Pterocles*, Gruoidea, *Sarothrura*, *Burhinus*, Scolopaci, *Sterna*, *Leptoptilos*, *Fregata*, *Scopus*, *Tigrisoma*, Accipitrimorphae, Cavitaves, *Cariama*, *Micrastur*, *Psittacus*, and Tyranni); J14, K21, S24.
- Phalanx 1 of major digit fenestrated (char. 204: 1, reversed in *Podargus*; also found in *Pterocles*, *Sterna*, *Phaethon*, *Fregata*, *Elanus*, and *Tyto*); J14, P15, K21, S24, W24.

Using the phylogenetic matrix of Livezey and Zusi (2006), Ericson and Qu (2024) also inferred a spatulate articulation with the furcula on the acrocoracoid process of the coracoid as a potential synapomorphy of this clade. A prominent transverse sulcus on the humerus was considered a possible synapomorphy of Strisores by Cracraft (1988) and Chen et al. (2019). This feature has been documented in a wide range of fossil strisoreans, such as the putative stem-nightjar (stem-Caprimulgidae) clade Archaeotrogonidae (Mourer-Chauviré, 1980; Mayr, 2021b), the stem-potoo (stem-Nyctibiidae) *Paraprefica kelleri* (Mayr, 1999a), the putative stem-apodiforms *Aegialornis* and *Eocypselus vincenti*, and the stem-swift (stem-Apodidae) *Scaniacypselus wardi* (Harrison, 1984). Archaeotrogonids additionally exhibit a short external spine on the sternum (Mayr, 1998a), and a deep scapulotricipital impression and prominently projecting (though small) ventral collateral tubercle on the ulna (Mayr, 2021b). However, the proximal termination of their carpal trochlea is only weakly angular (Figs. 3 and 4 in Mourer-Chauviré, 1980). An ulna greater than 40% of humerus + ulna + carpometacarpus length has been recorded in many fossil strisoreans (Olson, 1987; Mayr, 1998a; Mayr, 1999a; Mayr, 2010a), including some that are thought to be members of crown-group Apodiformes (Karhu, 1999; Mayr, 2003a; Mayr, 2015a). An elongated extensor process on the carpometacarpus is known to be present in archaeotrogonids (Mourer-Chauviré, 1980; Mayr and Kitchener, 2024a), *Aegialornis* (Mourer-Chauviré, 1988a), and *Fluvioviridavis* (Mayr and Kitchener, 2024a), which has been proposed to be a stem-frogmouth (stem-Podargidae; Nesbitt et al., 2011) or stem-oilbird (stem-Steatornithidae; Chen et al., 2019). The hypocleideum is short in the archaeotrogonid *Archaeodromus* (Fig. 2i in Mayr and Kitchener, 2024a), *Fluvioviridavis* (Fig. 8 in Mayr and Kitchener, 2024a), *Eocypselus rowei* (Ksepka et al., 2013) and the stem-apodiform *Primapus* (Mayr and Kitchener, 2024b), but altogether absent in the archaeotrogonid *Hassiavis* (Mayr, 1998a). *Fluvioviridavis* also possesses a sharply hooked lateral process on the coracoid (Fig. 8b in Mayr and Kitchener, 2024a), but has a more elongate external spine on the sternum than do extant strisoreans (Mayr and Kitchener, 2024a).

A pair of depressions on the dorsal surface of phalanx 1 of the major digit was suggested by Mayr (2002a) to be a synapomorphy of a putative clade uniting Caprimulgidae, Nyctibiidae, owlet-nightjars (Aegothelidae), and Apodiformes. Although the monophyly of this assemblage has not been upheld by recent molecular phylogenetic analyses (Prum et al., 2015; Chen et al., 2019; White and Braun, 2019; Kuhl et al., 2021), at least some of the features proposed to characterize this group may be synapomorphies of Strisores as a whole that have undergone secondary reversals in the oilbird (*Steatornis caripensis*) and podargids (Chen et al., 2019). Possessing a pair of dorsal depressions on phalanx 1 of the major digit (char. 203: 1) is itself a widespread trait among crown birds, but the present study optimizes fenestrae in these depressions as a potential strisorean synapomorphy. The presence of these fenestrae is variable in fossil strisoreans, being present in *Archaeodromus* (Mayr, 2021b), the stem-steatornithid *Prefica* (Mayr, 1999a), *Aegialornis* (Mourer-Chauviré, 1988a), the stem-hummingbird (stem-Trochilidae) *Argornis* (Karhu, 1999), and some species of *Fluvioviridavis* (Mayr and Kitchener, 2024a), but absent in *Hassiavis* (Mayr, 2021b), *Paraprefica* (Mayr, 2005a), *Fluvioviridavis platyrhamphus* (Mayr and Kitchener, 2024a), *Eocypselus* (Mayr, 2010a), and the stem-trochilids *Parargornis* (Mayr, 2003a) and *Eurotrochilus noniewiczi* (Bochenski and Bochenski, 2008). Nonetheless, the widespread distribution of this feature within Strisores may suggest that it is an underlying synapomorphy of the clade.

Other pectoral and forelimb characters that have been proposed as putative synapomorphies of Strisores include an elongated ventral ramus of the pisiform (Mayr, 2010b; char. 166: 0), an unhooked acrocoracoid process of the coracoid (Chen et al., 2019; char. 63: 0), and a caudally prominent ventral tubercle of the humerus (Ericson and Qu, 2024; char. 102: 1) but these were not optimized as unambiguous synapomorphies for this clade under the topologies studied here.

#### Gruiformes

Twelve character states were optimized as potential synapomorphies of Gruiformes, of which five were consistently inferred as such across all alternative molecular topologies studied.

- Hypocleideum ventrocranially deflected from furcular shafts (char. 10: 1 > 2, also found in *Pterocles*, Apodiformes, *Oceanites*, *Leptoptilos*, *Coragyps*, *Psittacus*, and *Menura*); J14, P15, K21, S24, W24.
- External spine of sternum less than 5% total sternum length (char. 15: 1 > 0, reversed in *Podica*; also found in *Chauna*, *Pterocles*, *Ardeotis*, Strisores, *Alca*, *Phaethon*, *Gavia*, *Phoebastria*, *Leptoptilos*, *Leucocarbo*, *Eudocimus*, *Coragyps*, and *Leptosomus*); J14, P15, K21.
- Two laminae on dorsal surface of sternum immediately caudal to cranial margin (Fig. 26, char. 30: 0 > 2, reversed in *Podica*; also found in *Chauna*, *Monias*, *Columba*, Apodiformes, *Eurypyga*, *Upupa*, and *Coracias*); J14, P15, K21, S24, W24.
- Six or more articular facets on costal margin of sternum (char. 36: 2 > 3, reversed in *Sarothrura*; also found in Anseriformes, *Phoenicopterus*, *Alca*, *Sterna*, *Phaethon*, *Gavia*, Procellariimorphae, *Sula*, *Eudocimus*, Scopus, *Pandion* + *Elanus*, and Psittaciformes); J14, P15, K21, S24, W24.
- Median trabecula of sternum rounded at caudal termination (char. 59: 1 > 0, reversed in *Aramus*; also found in non-neognath birds, *Ortalis* + *Rollulus*, *Monias*, *Columba*, *Florisuga*, *Turnix*, *Alca*, *Eurypyga*, Aequornithes, Accipitrimorphae, *Tyto*, *Urocolius*, *Bucorvus*, *Bucco*, Psittaciformes, and *Menura*); J14, P15, K21, S24, W24.
- Acrocoracoid process on coracoid straight (char. 63: 1 > 0, reversed in *Podica*; also found in *Gansus*, Anseriformes, *Ortalis*, *Phoenicopterus*, *Monias*, *Ardeotis*, Strisores, *Opisthocomus*, *Gavia*, *Pagodroma*, Pelecanimorphae, Accipitrimorphae, Strigiformes, *Urocolius*, *Micrastur*, and *Acanthisitta*); K21.
- Coracoid rounded and relatively thick at ventromedial margin of supracoracoid sulcus (Fig. 26, char. 69: 1 > 0, also found in non-neoavian birds, Mirandornithes, *Pterocles*, *Ardeotis*, *Tapera*, *Aegotheles*, *Streptoprocne*, Aequornithes, *Bucorvus*, *Nestor*, and *Acanthisitta*); J14, P15, K21, S24, W24.
- Supracoracoid nerve foramen in coracoid present (char. 70: 0 > 1, also found in *Ichthyornis*, *Anseranas*, *Phoenicopterus*, *Corythaeola*, Daedalornithes, Charadrii, *Alca*, *Sterna*, *Phaethon*, Procellariiformes, *Leptoptilos*, *Eudocimus*, *Pelecanus*, Accipitrimorphae, Strigiformes, *Leptosomus*, and *Micrastur*); P15.
- Pronounced ventral curvature of scapula (char. 92: 0 > 1, reversed in *Balearica*; also found in *Caprimulgus*, *Nyctibius*, *Florisuga*, *Opisthocomus*, *Eurypyga*, *Spheniscus*, *Oceanites* + *Pagodroma*, *Leptoptilos*, *Tigrisoma*, *Coragyps*, *Pandion*, *Tyto*, *Merops*, *Alcedo*, *Psilopogon*, *Nestor*, *Acanthisitta*, and Tyranni); K21.
- Deltopectoral crest rounded dorsally (char. 121: 1 > 0, reversed in *Podica*; also found in *Gansus*, *Chauna*, *Anseranas*, Mirandornithes, *Ardeotis*, *Aegotheles*, *Opisthocomus*, *Charadrius*, *Rostratula*, *Alca*, *Eudocimus*, Eucavitaves, and Australaves); P15, K21, W24.
- Humeral shaft width approximately constant throughout length (char. 126: 2 > 0, reversed in *Podica*; also found in *Gansus*, *Eudromia*, Anseriformes, *Ortalis*, Mirandornithes, *Monias*, *Ardeotis*, *Limosa*, *Alca*, *Phaethon*, *Gavia*, *Phoebastria*, *Pagodroma*, Suliformes, *Eudocimus*, *Coragyps*, *Elanus*, Strigiformes, *Urocolius*, *Merops*, *Alcedo*, *Bucco*, *Micrastur*, and *Acanthisitta*); S24.
- Internal index on phalanx 1 of major digit poorly developed (char. 201: 1 > 0, reversed in *Balearica*; also found in *Gansus*, *Eudromia*, *Chauna*, *Alectura*, *Rollulus*, *Podilymbus*, *Monias*, *Corythaeola*, *Opisthocomus*, *Rostratula*, *Turnix*, *Spheniscus*, *Tigrisoma*, *Ninox*, *Urocolius*, *Bucorvus*, *Merops*, Piciformes, and Australaves); K21, W24.

**Figure 26.**
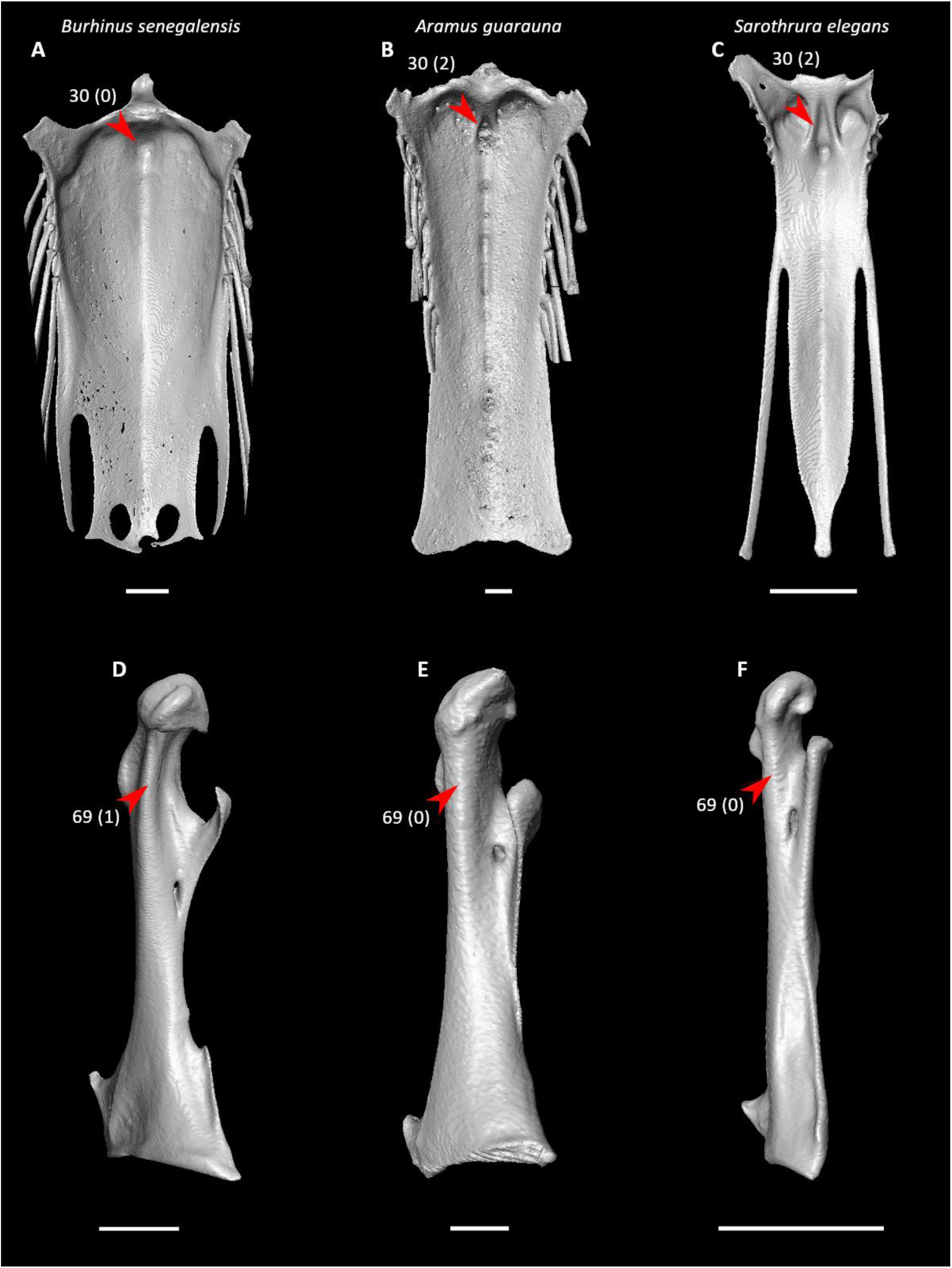
Sterna of *Burhinus senegalensis* (**A**, FMNH 313704), *Aramus guarauna* (**B**, FMNH 376078), and *Sarothrura elegans* (**C**, NHMUK S1997.34.2) in dorsal view and coracoids of *B. senegalensis* (**D**, FMNH 313704, left element, mirrored), *A. guarauna* (**E**, FMNH 376078, right element), and *S. elegans* (**F**, NHMUK S1997.34.2, right element) in ventromedial view. Scale bars = 5 mm. Arrows indicate the region immediately caudal to the cranial margin of the sternum (**A–C**) and the ventromedial margin of supracoracoid sulcus (**D–F**). The right craniolateral process of the sternum in *Sarothrura* is broken. *Burhinus* exhibits state 0 for character 30 (laminae absent from region immediately caudal to the cranial margin of the sternum), whereas *Aramus* and *Sarothrura* exhibit state 2 (two laminae in region immediately caudal to the cranial margin of the sternum, optimized as a synapomorphy of Gruiformes). *Burhinus* also exhibits state 1 for character 69 (coracoid compressed and keeled at ventromedial margin of supracoracoid sulcus), whereas *Aramus* and *Sarothrura* exhibit state 0 (coracoid rounded and relatively thick at ventromedial margin of supracoracoid sulcus, optimized as a synapomorphy of Gruiformes).

The optimization of six or more costal facets as a synapomorphy of this clade is consistent with the presence of six costal facets on the sternum of the messelornithids *Messelornis nearctica* (Hesse, 1992) and *Songzia acutunguis* (Wang et al., 2012), which recent studies suggest are stem members of Ralloidea (Mayr, 2004b; Musser et al., 2019). Messelornithids possess a supracoracoid nerve foramen (Hesse, 1988; Mourer-Chauviré, 1995; Mayr and Kitchener, 2024c), but have a dorsocaudally deflected hypocleideum (Hesse, 1988; Hesse, 1990), a hooked acrocoracoid process of the coracoid (Hesse, 1988; Pl. VII in Hesse, 1990; Figs. 4–7 in Mourer-Chauviré, 1995; Fig. 3 in Mayr and Kitchener, 2024c), and a well-developed internal index process on phalanx 1 of the major digit (Hesse, 1990).

#### Charadriiformes

Twelve character states were optimized as potential synapomorphies of Charadriiformes, of which one was consistently inferred as such across all alternative molecular topologies studied. Five character states were inferred as synapomorphies under all molecular reference topologies except for Prum et al. (2015), in which Charadriiformes was recovered as the extant sister group of Mirandornithes instead of Gruiformes.

- Strong craniocaudal curvature of furcula (char. 2: 0 > 1, reversed in *Turnix*; also found in Anseres, Mirandornithes, *Nyctibius*, Aequornithes, *Pandion*, *Trogon*, and *Coracias* + *Merops*); J14, K21, S24, W24.
- Pointed acromial processes on furcula (char. 5: 0 > 1, also found in *Ichthyornis*, Anseriformes, *Phoenicopterus*, *Pterocles*, *Nyctibius*, Apodiformes, *Sarothrura*, *Eurypyga*, *Gavia*, Procellariimorphae, *Sula*, *Coragyps*, *Pandion*, *Urocolius*, and *Acanthisitta*); J14, K21, S24, W24.
- Hypocleideum dorsocaudally deflected from furcular shafts (char. 10: 1 > 0, reversed in *Limosa* and *Sterna*; also found in *Ichthyornis*, *Rollulus*, *Podilymbus*, *Podargus*, Phaethontimorphae, *Phoebastria*, *Sula*, *Scopus*, *Tigrisoma*, *Urocolius*, *Bucorvus*, Coraciiformes, *Acanthisitta*, and *Erythropitta*); J14, K21, S24, W24.
- Pneumatic pores in intercostal incisures of sternum absent (char. 37: 1 > 0, also found in *Rollulus*, Mirandornithes, *Caprimulgus*, *Aegotheles*, Heliornithes, *Gavia*, *Spheniscus*, *Oceanites*, *Pagodroma*, *Leucocarbo*, *Urocolius*, Piciformes, *Acanthisitta*, *Erythropitta*, and *Climacteris*); J14, K21, S24, W24.
- Dorsoventral orientation of acrocoracoid process essentially coplanar with main craniocaudal axis of coracoid (char. 62: 0 > 1, reversed in *Limosa* and *Alca*; also found in Galliformes, *Pterocles*, *Columba*, Strisores, *Psophia*, Heliornithes, *Urocolius*, Eucavitaves, Tyranni, and *Climacteris*); P15.
- Acrocoracoid process on coracoid forms pronounced hook (char. 63: 0 > 1, also found in *Ichthyornis*, *Eudromia*, *Alectura*, *Rollulus*, *Podilymbus*, *Pterocles*, *Columba*, *Corythaeola*, *Tapera*, *Aegotheles*, *Streptoprocne*, *Podica*, Phaethontimorphae, Procellariimorphae, Cavitaves, *Cariama*, Psittaciformes, and Eupasseres); S24, W24.
- Scapular cotyle on coracoid deep and cup-like (char. 66: 1 > 0, reversed in *Alca*; also found in *Gansus*, *Ichthyornis*, *Chauna*, *Dendrocygna*, *Phoenicopterus*, *Pterocles*, Apodiformes, *Opisthocomus*, *Eurypyga*, *Gavia*, *Leptoptilos*, *Pelecanus*, and *Tigrisoma*); K21, W24.
- Strong ventral curvature of sternal articular facet of coracoid (char. 83: 0 > 1, also found in *Chauna*, *Rollulus*, Gruoidea, *Sarothrura*, *Spheniscus*, *Oceanites*, *Coragyps*, Psittaciformes, and *Acanthisitta*); P15.
- Moderate ventral curvature of scapula (char. 92: 1 > 0, also found in most birds outside of Gruiformes, *Opisthocomus*, and some Strisores, Phaethoquornithes, and Telluraves); S24.
- Apex of dorsal cotylar process of ulna approximately coplanar with dorsal surface of ulnar main body (char. 150: 0 > 1, reversed in *Limosa*; also found in *Ichthyornis*, Anseriformes, *Podilymbus*, *Corythaeola*, Strisores, *Opisthocomus*, *Urocolius*, Cavitaves, *Nestor*, and Passeriformes); K21.
- Radial incisure on ulna prominent (char. 156: 0 > 1, reversed in *Turnix*; also found in *Ichthyornis*, *Eudromia*, *Chauna*, *Anseranas*, Mirandornithes, *Ardeotis*, *Nyctibius*, Grues, *Phaethon*, *Phoebastria*, *Pagodroma*, *Leptoptilos*, *Fregata*, *Sula*, *Scopus* + *Pelecanus*, Accipitrimorphae, and Psittaciformes); J14, K21, S24, W24.
- Tubercle at insertion point of humerocarpal ligament on pisiform (char. 167: 0 > 1, also found in *Pterocles*, *Columba*, *Aegotheles*, *Florisuga*, Pelecanimorphae, *Coragyps*, *Tyto*, *Micrastur*, *Erythropitta*, and Passeri); J14, P15, K21, S24, W24.

Using the phylogenetic matrix of Livezey and Zusi (2006), Ericson and Qu (2024) also inferred a caudoventrally placed pneumotricipital fossa on the humerus in distal view as a potential synapomorphy of this clade. A hooked acrocoracoid process of the coracoid and the presence of a tubercle at the insertion point of the humerocarpal ligament on the pisiform have been considered characteristic of Charadriiformes by previous studies (Mayr and Knopf, 2007; Mayr, 2008a; Mayr, 2011b; Mayr and Kitchener, 2023b). Additionally, strong cranial curvature of the sternal articular facet (char. 82: 1) of the coracoid as well as a long transverse sulcus (char. 115: 2) and a prominent supracondylar process (char. 136: 2) on an apneumatic (char. 109: 0) humerus are distinctive features widely found in charadriiforms (Mayr, 2000b; Mayr and Knopf, 2007; Mayr, 2008a; Mayr, 2011b; Mayr, 2016a; Hood et al., 2019; Mayr et al., 2022; Burton et al., 2023; Mayr and Kitchener, 2023b), but their optimization as synapomorphies of this clade is likely sensitive to ingroup taxon sampling and outgroup interrelationships.

Pectoral girdle and forelimb elements are mostly unknown in the most completely preserved putative stem-charadriiforms, *Scandiavis* (Bertelli et al., 2013; Heingård et al., 2021) and *Nahmavis* (Musser and Clarke, 2020), though Mayr and Kitchener (2023b) assigned a distal humerus from the London Clay Formation to *Scandiavis* and tentatively referred additional pectoral girdle and forelimb material from the same locality to a closely related taxon. Similarities of these specimens to crown-group charadriiforms include a hooked acrocoracoid process and the presence of a dorsal supracondylar tubercle on the humerus, though this is weakly developed (Mayr and Kitchener, 2023b).

#### Phaethoquornithes

Three character states were optimized as potential synapomorphies of Phaethoquornithes, but none of these were consistently inferred as such across all alternative molecular topologies studied.

- Hypocleideum less than 5% the total length of the furcula (char. 9: 1 > 0, reversed in *Oceanites*; also found in *Podilymbus*, Strisores, Scolopaci, *Alca*, *Coragyps*, *Bucorvus*, *Alcedo*, *Cariama*, and *Menura*); J14, K21.
- Dorsal tubercle of humerus pointed (char. 105: 0 > 1, reversed in Procellariimorphae and *Tigrisoma*; also found in Anseriformes, Mirandornithes, *Nyctibius*, *Florisuga*, *Aramus*, Ralloidea, *Pandion*, Strigiformes, *Trogon*, *Upupa*, *Coracias*, and Passeri); J14, K21, S24.
- Prominent ligamental groove on ventral ramus of pisiform (char. 168: 0 > 1, reversed in *Phoebastria*, *Oceanites*, *Eudocimus*, and *Tigrisoma*; also found in *Ichthyornis*, *Dendrocygna*, *Columba*, *Tapera*, *Nyctibius*, *Streptoprocne*, Grues, Ralloidea, *Charadrius*, Scolopaci, *Sterna*, *Pandion*, Strigiformes, *Upupa*, *Coracias*, *Alcedo*, *Psittacus*, *Erythropitta*, and Passeri); J14, P15, K21, W24.

A short hypocleideum is visible in the stem-loon (stem-Gaviidae) *Nasidytes* (Fig. 3F in Mayr and Kitchener, 2022), whereas a small projection for articulation with the sternum can be seen in this region of the furcula in the stem-tropicbird (stem-Phaethontidae) *Prophaethon* (Mayr, 2015b). *Prophaethon* additionally exhibits a prominent ligamental groove on the ventral ramus of the pisiform (Fig. 5n in Mayr, 2015b).

#### Phaethontimorphae

Seven character states were optimized as potential synapomorphies of Phaethontimorphae, of which one was consistently inferred as such across all alternative molecular topologies studied.

- Hypocleideum dorsocaudally deflected from clavicular shafts (char. 10: 1 > 0, also found in *Ichthyornis*, *Rollulus*, *Podilymbus*, *Podargus*, Charadriiformes, *Phoebastria*, *Sula*, *Scopus*, *Tigrisoma*, *Urocolius*, *Bucorvus*, Coraciiformes, *Acanthisitta*, and *Erythropitta*); J14, K21, S24, W24.
- Coracoid sulci on sternum crossed (char. 25: 0 > 1, also found in *Ichthyornis*, *Phoenicopterus*, *Corythaeola*, Grues, Procellariimorphae, *Scopus*, *Tigrisoma*, *Elanus*, *Leptosomus*, and *Micrastur*); J14, P15, K21, S24, W24.
- Acrocoracoid process on coracoid forms pronounced hook (char. 63: 0 > 1, also found in *Ichthyornis*, *Eudromia*, *Alectura*, *Rollulus*, *Podilymbus*, *Pterocles*, *Columba*, *Corythaeola*, *Tapera*, *Aegotheles*, *Streptoprocne*, *Podica*, Charadriiformes, Procellariimorphae, Cavitaves, *Cariama*, Psittaciformes, and Eupasseres); S24, W24.
- Ulna greater than 40% of humerus + ulna + carpometacarpus length (char. 98: 0 > 1, also found in *Gansus*, *Ichthyornis*, *Chauna*, *Phoenicopterus*, *Columba*, *Ardeotis*, Strisores, Grues, *Burhinus*, *Rostratula*, *Sterna*, *Phoebastria*, Pelecanimorphae, Accipitrimorphae, Strigiformes, Cavitaves, and Eufalconimorphae); P15, W24.
- Olecranon fossa on humerus deep (char. 127: 0 > 1, also found in Procellariimorphae, Suliformes, *Scopus* + *Pelecanus*, and most non-phaethoquornithean neognaths); W24.
- Brachial fossa on humerus positioned medially on shaft (Fig. 27, char. 133: 0 > 1, also found in Anseres, *Ardeotis*, *Nyctibius*, *Opisthocomus*, Gruoidea, Procellariiformes, *Tyto*, and *Leptosomus*); J14, P15, K21, S24.
- Minor metacarpal prominently narrows distally in caudal view (char. 180: 0 > 1, also found in *Ichthyornis*, Anseres, Mirandornithes, Columbimorphae, *Aegotheles*, Gruiformes, Charadriiformes, Pelecanimorphae, *Leptosomus*, *Coracias*, Accipitrimorphae, Strigiformes, and Australaves); W24.

**Figure 27.**
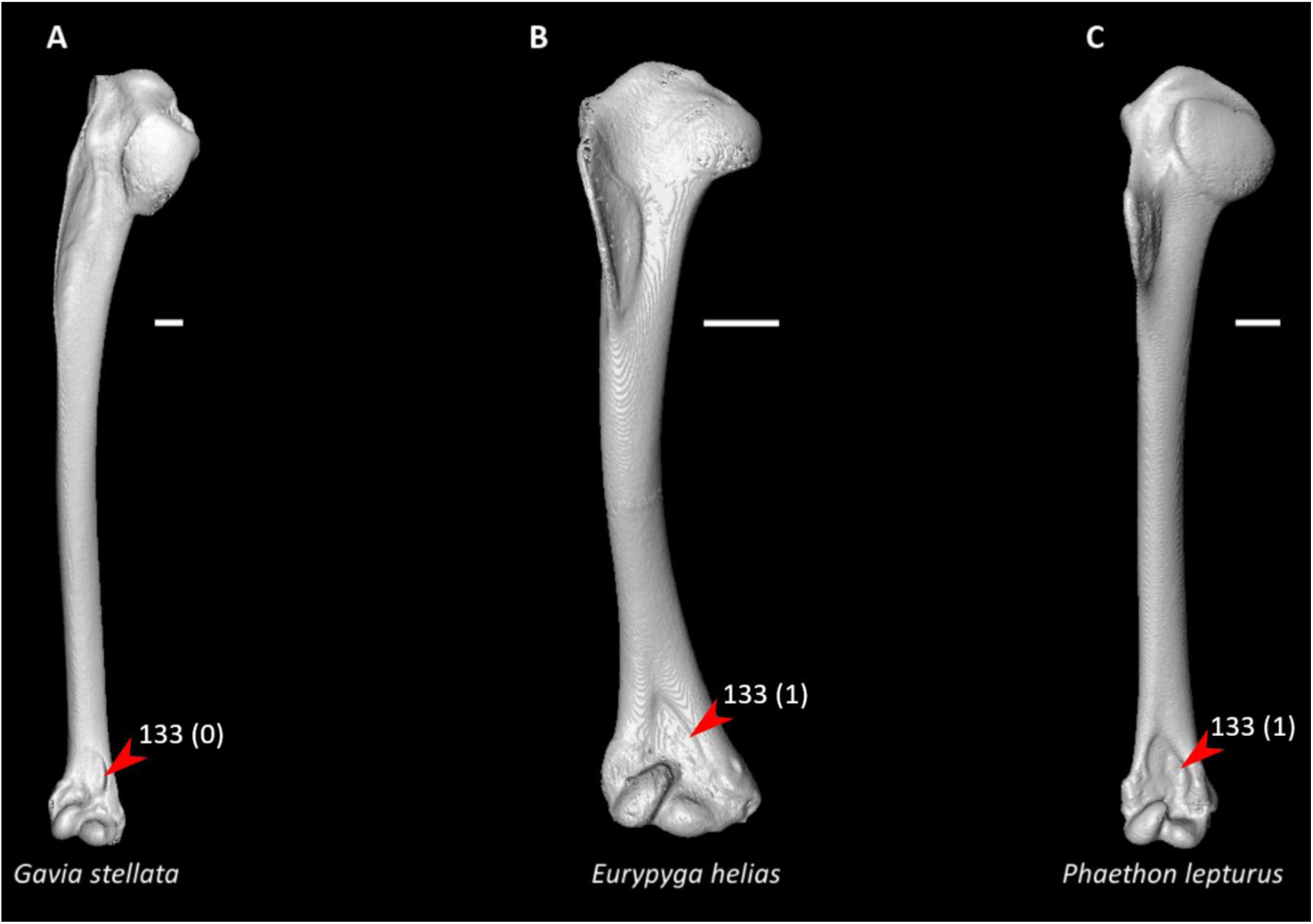
Humeri of *Gavia stellata* (**A**, NHMUK 1891.7.20.132, right element), *Eurypyga helias* (**B**, FMNH 317341, right element), and *Phaethon lepturus* (**C**, NHMUK 1876.3.16.3, right element) in cranial view. Scale bars = 5 mm. Arrows indicate the brachial fossa. *Gavia* exhibits state 0 for character 133 (brachial fossa positioned ventrally on humeral shaft), whereas *Eurypyga* and *Phaethon* exhibit state 1 (brachial fossa positioned medially on humeral shaft, optimized as a synapomorphy of Phaethontimorphae).

The identification of potential morphological synapomorphies for Phaethontimorphae has been challenging (Mayr, 2014c; Mayr, 2019b; Mayr, 2022a; Sangster et al., 2022; Mayr and Kitchener, 2024d) and no well-corroborated fossils of total-group Eurypygiformes have been formally described (Mayr, 2022a). In the Paleocene stem-phaethontid *Clymenoptilon*, the ulna is proportionately shorter than in extant phaethontids and does not exceed 40% of humerus + ulna + carpometacarpus length (Mayr et al., 2023b). Nonetheless, some of the above characters have been documented in stem-phaethontid fossils—*Prophaethon* exhibits dorsocaudally-directed projections on the furcular symphysis (Mayr, 2015b), and both *Prophaethon* and *Lithoptila* exhibit a medially positioned brachial fossa on the humerus (Fig. S2b in Mayr, 2015b; Bourdon et al., 2008; Fig. 1 in Mayr and Kitchener, 2024d). Of note is that messelornithids, which were originally interpreted as stem-eurypygids (Hesse, 1990), exhibit a ventrally placed humeral brachial fossa (Mourer-Chauviré, 1995), consistent with the recent consensus that they are stem-ralloids.

The crossed coracoid sulci on the sternum and medially positioned brachial fossa on the humerus distinguish phaethontimorphs from members of Suliformes, which have traditionally been thought to be closely related to phaethontids (e.g., Cracraft, 1985; Livezey and Zusi, 2007). However, some recent morphological analyses proposed a closer relationship between Phaethontidae and Procellariiformes (Mayr, 2003b; Smith, 2010), which is consistent with the fact that all of these potential phaethontimorph synapomorphies also appear to be homoplastically present in at least some procellariiforms.

One of the two extant species of Eurypygiformes, the kagu (*Rhynochetos jubatus*), lacks a caudal projection on the furcular symphysis and crossed coracoid sulci (Parker, 1869). However, this species is essentially flightless (Hunt, 1996) and is likely to possess a highly autapomorphic pectoral girdle and forelimb skeleton as a result.

#### Aequornithes

Four character states were optimized as potential synapomorphies of Aequornithes, of which two were consistently inferred as such across all alternative molecular topologies studied.

- Strong craniocaudal curvature of furcula (char. 2: 0 > 1, reversed in *Phoebastria*, *Fregata*, *Leucocarbo*, *Pelecanus*, and *Tigrisoma*; also found in Anseres, Mirandornithes, *Nyctibius*, Charadriiformes, *Pandion*, *Trogon*, and *Coracias* + *Merops*); J14, K21, S24, W24.
- Single pair of caudal fenestrae in sternum (char. 49: 2 > 1, reversed in *Pagodroma* and *Eudocimus*; also found in *Eudromia*, Anseres, Galliformes, Mirandornithes, *Monias*, *Caprimulgus*, *Balearica*, Ralloidea, *Turnix*, *Pandion* + *Elanus*, *Tyto*, Bucerotiformes, and Australaves); J14, S24.
- Coracoid rounded and relatively thick at ventromedial margin of supracoracoid sulcus (char. 69: 1 > 0, reversed in *Phoebastria*, *Pagodroma*, *Leucocarbo*, and *Tigrisoma*; also found in non-neoavian birds, Mirandornithes, *Pterocles*, *Ardeotis*, *Tapera*, *Aegotheles*, *Streptoprocne*, Gruiformes, *Bucorvus*, *Nestor*, and *Acanthisitta*); J14, P15, K21, S24, W24.
- Scapular acromion subequal or caudal to coracoid tubercle (Fig. 28, char. 89: 1 > 0, reversed in *Phoebastria* and Pelecanes; also found in *Ichthyornis*, *Chauna*, *Anseranas*, *Ardeotis*, *Nyctibius*, *Florisuga*, *Alca*, and *Sterna*); J14, P15, K21, S24, W24.

**Figure 28.**
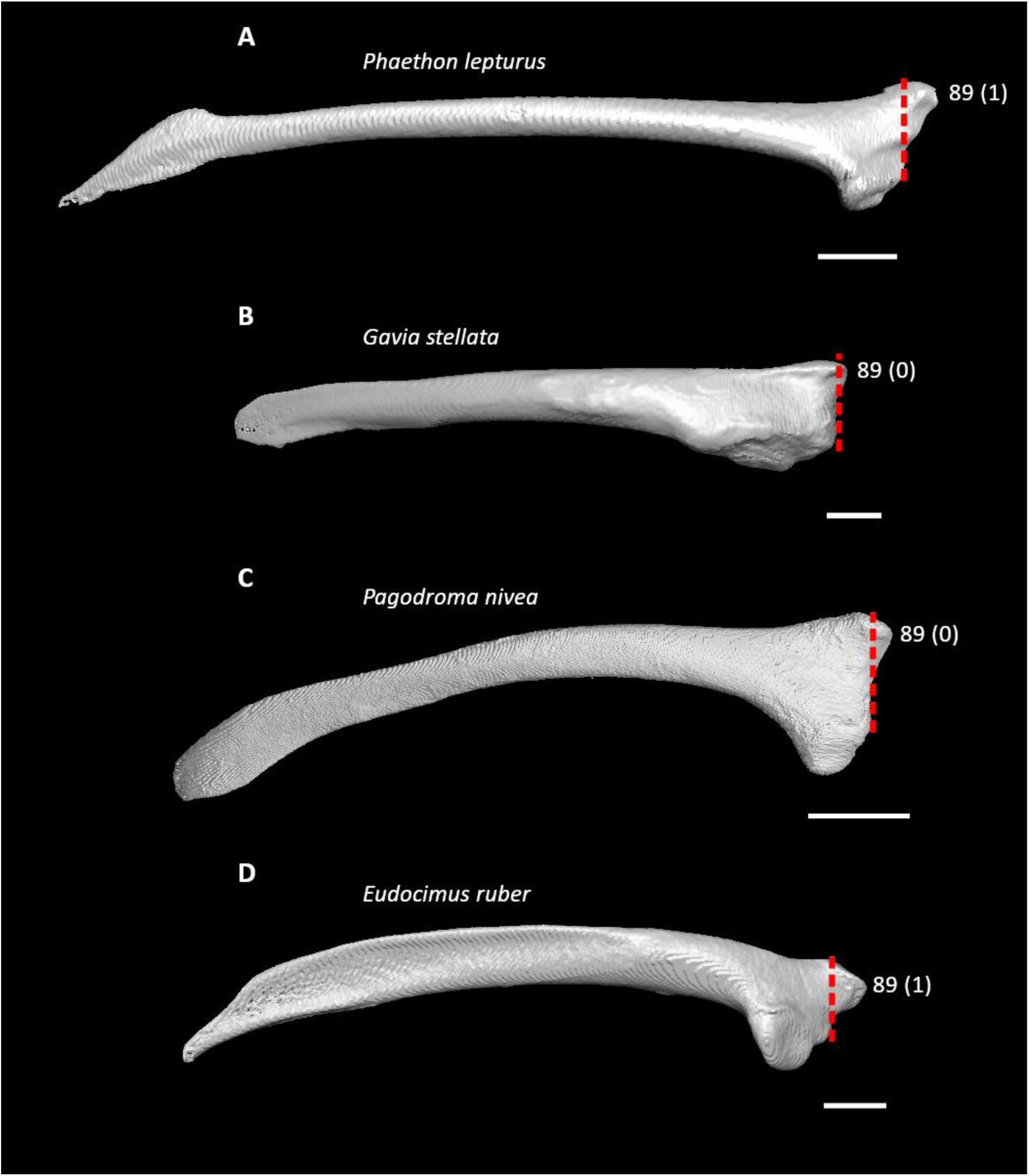
Scapulae of *Phaethon lepturus* (**A**, NHMUK 1876.3.16.3, left element, mirrored), *Gavia stellata* (**B**, NHMUK 1891.7.20.132, right element), *Pagodroma nivea* (**C**, NHMUK S1998.55, right element), and *Eudocimus ruber* (**D**, NHMUK S1999.8.1, right element) in lateral (**A, C–D**) or craniolateral (**B**) view. Scale bars = 5 mm. Dotted lines indicate cranial extent of the coracoid tubercle. *Phaethon* exhibits state 1 for character 89 (acromion extends distinctly cranial to coracoid tubercle), whereas *Gavia* and *Pagodroma* exhibit state 0 (acromion subequal or caudal to coracoid tubercle, optimized as a synapomorphy of Aequornithes). *Eudocimus* also exhibits state 1 for character 89, a reversal to the plesiomorphic condition (optimized as a synapomorphy of Pelecanes).

A scapular acromion equal or caudal to the coracoid tubercle is found in Paleocene stem-penguins (stem-Spheniscidae) (Mayr et al., 2017a; Mayr et al., 2017b; Blokland et al., 2019; Mayr et al., 2021) and the possible stem-procellariiform *Rupelornis* (Mayr et al., 2002; De Pietri et al., 2010; Mayr and Smith, 2012). *Rupelornis* (Mayr et al., 2002) and the stem-gaviid *Nasidytes* (Mayr and Kitchener, 2022) are known to have had a single pair of caudal incisions in the sternum, as does the stem-ibis (stem-Threskiornithidae) *Rhynchaeites* (Mayr, 2002b), which is noteworthy given that crown threskiornithids have a four-notched sternum. Conversely, the stem-frigatebird (stem-Fregatidae) *Limnofregata* has two pairs of caudal sternal incisions (Olson, 1977), perhaps representing another reversal to this condition within Aequornithes. *Nasidytes* exhibits a more elongate scapular acromion and a less craniocaudally curved furcula compared to extant gaviids (Mayr and Kitchener, 2022), which may suggest either their apomorphic presence in *Nasidytes* or multiple losses of these traits among Aequornithes.

#### Feraequornithes

Two character states were optimized as potential synapomorphies of Feraequornithes, both of which were consistently inferred as such across all alternative molecular topologies studied.

- Median trabecula of sternum distinctly tapered (char. 58: 0 > 1, reversed in *Leucocarbo* and *Pelecanus*; also found in *Gansus*, *Alectura*, *Ortalis*, *Monias*, *Ardeotis*, Letornithes, *Sarothrura*, *Rallus*, *Charadrius*, *Turnix*, *Coragyps*, *Ninox*, Coraciimorphae, and *Cariama*); J14, P15, K21, S24, W24.
- Radius craniocaudally curved (Fig. 29, char. 144: 0 > 1, reversed in *Oceanites*; also found in Anseriformes, *Phoenicopterus*, *Ardeotis*, Strisores, Grues, Charadrii, *Coragyps*, *Pandion*, Strigiformes, Cavitaves, *Micrastur*, Tyranni, and *Climacteris*); J14, P15, K21, S24, W24.

**Figure 29.**
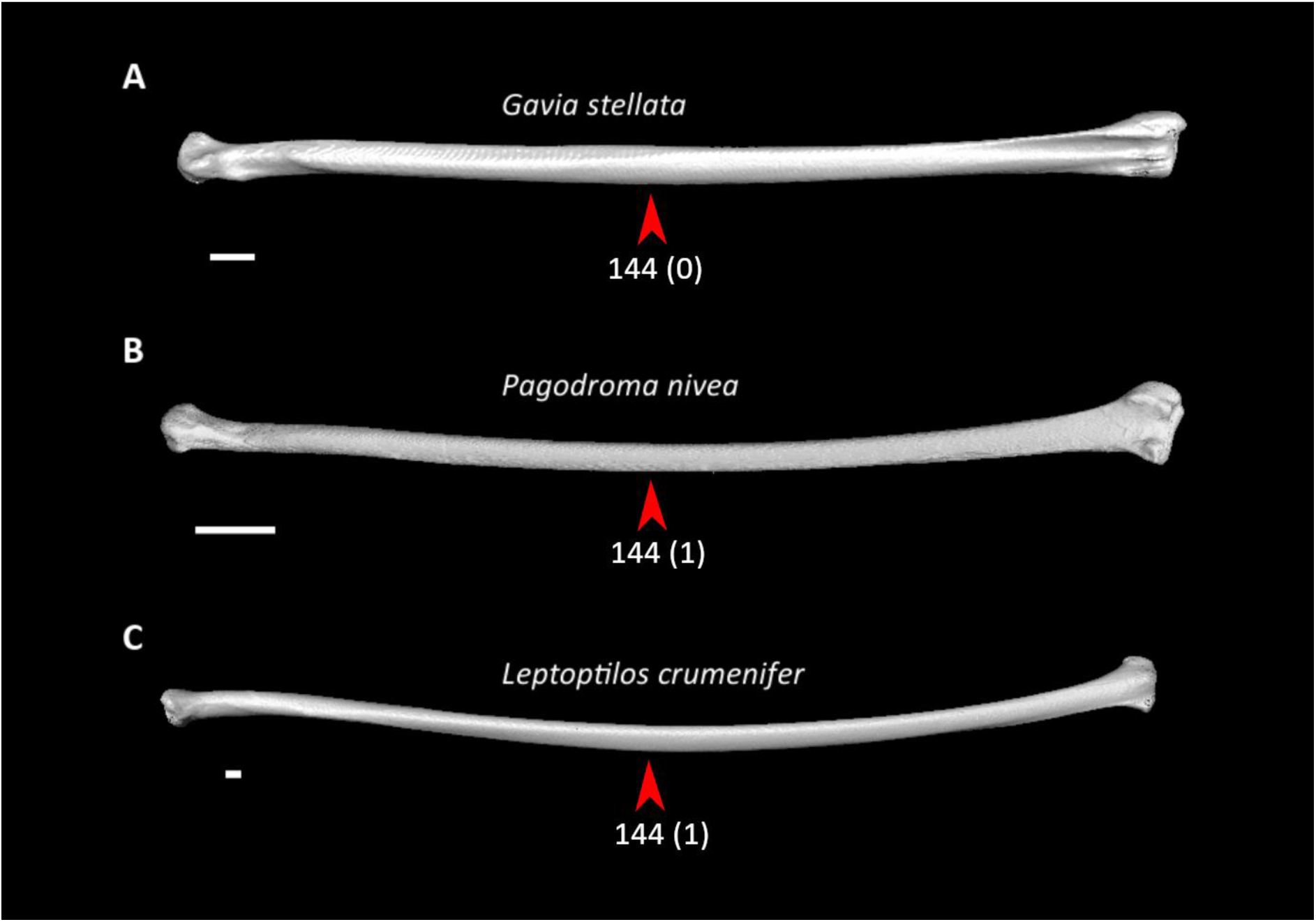
Radii of *Gavia stellata* (**A**, NHMUK 1891.7.20.132, right element), *Pagodroma nivea* (**B**, NHMUK S1998.55, right element), and *Leptoptilos crumenifer* (**C**, NHMUK S1952.3.182, right element) in dorsal view. Scale bars = 5 mm. Arrows indicate the radial shaft. *Gavia* exhibits state 0 for character 144 (radial shaft craniocaudally straight), whereas *Pagodroma* and *Leptoptilos* exhibit state 1 (radial shaft craniocaudally curved, optimized as a synapomorphy of Feraequornithes).

The inference of a craniocaudally curved radius as a synapomorphy of this clade is consistent with the straight radius in the stem-gaviid *Colymboides* (Storer, 1956; Cheneval, 1984) and the strongly curved radius in Paleocene stem-spheniscids (Mayr et al., 2017a; Blokland et al., 2019; Mayr et al., 2020a). Previously recognized morphological synapomorphies of Feraequornithes have been limited to cranial characters (Sangster and Mayr, 2021).

#### Procellariimorphae

Seven character states were optimized as potential synapomorphies of Procellariimorphae, of which three were consistently inferred as such across all alternative molecular topologies studied.

- Acrocoracoid process on coracoid forms pronounced hook (char. 63: 0 > 1, also found in *Ichthyornis*, *Eudromia*, *Alectura*, *Rollulus*, *Podilymbus*, *Pterocles*, *Columba*, *Corythaeola*, *Tapera*, *Aegotheles*, *Streptoprocne*, *Podica*, Charadriiformes, Phaethontimorphae, Cavitaves, *Cariama*, Psittaciformes, and Eupasseres); S24, W24.
- Mound-shaped tuberosity absent from lateral surface of scapula (char. 94: 1 > 0, also found in *Phaethon*, *Scopus*, *Tigrisoma*, and most non-phaethoquornithean birds); P15.
- Distal portion of scapula spatulate (char. 95: 1 > 0, also found in *Rollulus* and *Pterocles*); J14, P15. K21, S24, W24.
- Dorsal tubercle of humerus rounded (char. 105: 1 > 0, reversed in *Pagodroma*; also found in *Tigrisoma* and many non-phaethoquornithean birds); J14, P15, K21, S24, W24.
- Humeral shaft straight (char. 125: 0 > 1, reversed in *Oceanites*; also found in Mirandornithes, *Pterocles*, *Columba*, Apodiformes, *Opisthocomus*, Charadriiformes, *Phaethon*, *Fregata*, *Leucocarbo*, *Urocolius*, *Psilopogon*, *Psittacus*, *Neopelma*, and Passeri); J14, P15, K21, S24.
- Olecranon fossa on humerus deep (char. 127: 0 > 1, also found in Phaethontimorphae, Suliformes, *Scopus* + *Pelecanus*, and most non-phaethoquornithean neognaths); W24.
- Pneumatic foramen in infratrochlear fossa on carpometacarpus absent (char. 191: 1 > 0, reversed in *Pagodroma*; also found in Mirandornithes, Pteroclimesites, Strisores, *Opisthocomus*, *Rallus*, Scolopaci + Lari, *Leucocarbo*, *Eudocimus*, *Tigrisoma*, *Ninox*, *Urocolius*, *Merops*, *Jynx*, *Pandion*, *Nestor*, and Passeriformes); J14, P15, K21, S24, W24.

Sangster et al. (2022) previously inferred a straight humeral shaft as a synapomorphy of this clade, based on the phylogenetic dataset of Livezey and Zusi (2006). However, as with the present study, only extant members of Procellariimorphae were sampled in that dataset. Whereas this character occurs in *Rupelornis* (Mayr et al., 2002), Paleocene stem-spheniscids consistently exhibit a more strongly curved humeral shaft than crown spheniscids (Slack et al., 2006; Jadwiszczak et al., 2013; Mayr et al., 2017a; Blokland et al., 2019; Ksepka et al., 2023b), suggesting that this feature evolved independently in crown-group spheniscids and procellariiforms. Many of the above characters are difficult to assess in fossil procellariimorphs based on preservation and available descriptions, and detailed examination of relevant specimens will likely be necessary to confidently infer potential synapomorphies of this clade.

#### Pelecanimorphae

Seven character states were optimized as potential synapomorphies of Pelecanimorphae, of which three were consistently inferred as such across all alternative molecular topologies studied.

- Rounded acromial processes on furcula (char. 5: 1 > 0, reversed in *Sula*; also found in *Phaethon* and many non-phaethoquornithean birds); P15.
- Pneumatic foramen in median sulcus of sternum immediately caudal to cranial margin (char. 29: 0 > 1, reversed in *Leucocarbo*, *Scopus*, and *Pelecanus*; also found in *Ichthyornis*, *Dendrocygna*, *Alectura*, *Ortalis*, *Phoenicopterus*, *Pterocles*, *Ardeotis*, *Tapera*, Strisores, *Opisthocomus*, Grues, *Podica*, *Phaethon*, *Coragyps*, Strigiformes, Picocoraciades, *Micrastur*, Psittaciformes, *Neopelma*, and *Menura*); J14, P15, K21, S24.
- Infra-acrocoracoid recess in coracoid deep (Fig. 30, char. 65: 0 > 1, reversed in *Leucocarbo* and *Tigrisoma*; also found in *Ichthyornis*, *Pterocles*, *Columba*, *Nyctibius*, *Aegotheles*, *Streptoprocne*, *Psophia*, Charadriiformes, *Phaethon*, *Pandion* + *Elanus*, *Ninox*, Cavitaves, and Australaves); J14, P15, K21, S24, W24.
- Ulna greater than 40% of humerus + ulna + carpometacarpus length (char. 98: 0 > 1, also found in *Gansus*, *Ichthyornis*, *Chauna*, *Phoenicopterus*, *Columba*, *Ardeotis*, Strisores, Grues, *Burhinus*, *Rostratula*, *Sterna*, Phaethontimorphae, *Phoebastria*, Accipitrimorphae, Strigiformes, Cavitaves, and Eufalconimorphae); P15, W24.
- Quill knobs on ulna strongly developed (Fig. 30, char. 161: 0 > 1, reversed in *Sula*; also found in *Anseranas*, *Podilymbus*, *Monias*, *Columba*, *Corythaeola*, *Ardeotis*, *Tapera*, *Caprimulgus*, *Aramus*, *Charadrius*, Scolopaci + Lari, *Oceanites*, Accipitrimorphae, Picocoraciades, and Passeriformes); J14, P15, K21, S24, W24.
- Tubercle at insertion point of humerocarpal ligament on pisiform (char. 167: 0 > 1, reversed in *Leucocarbo* and *Scopus*; also found in *Pterocles*, *Columba*, *Aegotheles*, *Florisuga*, Charadriiformes, *Coragyps*, *Tyto*, *Micrastur*, *Erythropitta*, and Passeri); J14, P15, K21, S24, W24.
- Minor metacarpal prominently narrows distally in caudal view (char. 180: 0 > 1, reversed in Suloidea and *Pelecanus*; also found in *Ichthyornis*, Anseres, Mirandornithes, Columbimorphae, *Aegotheles*, Gruiformes, Charadriiformes, Phaethontimorphae, *Leptosomus*, *Coracias*, Accipitrimorphae, Strigiformes, and Australaves); W24.

**Figure 30.**
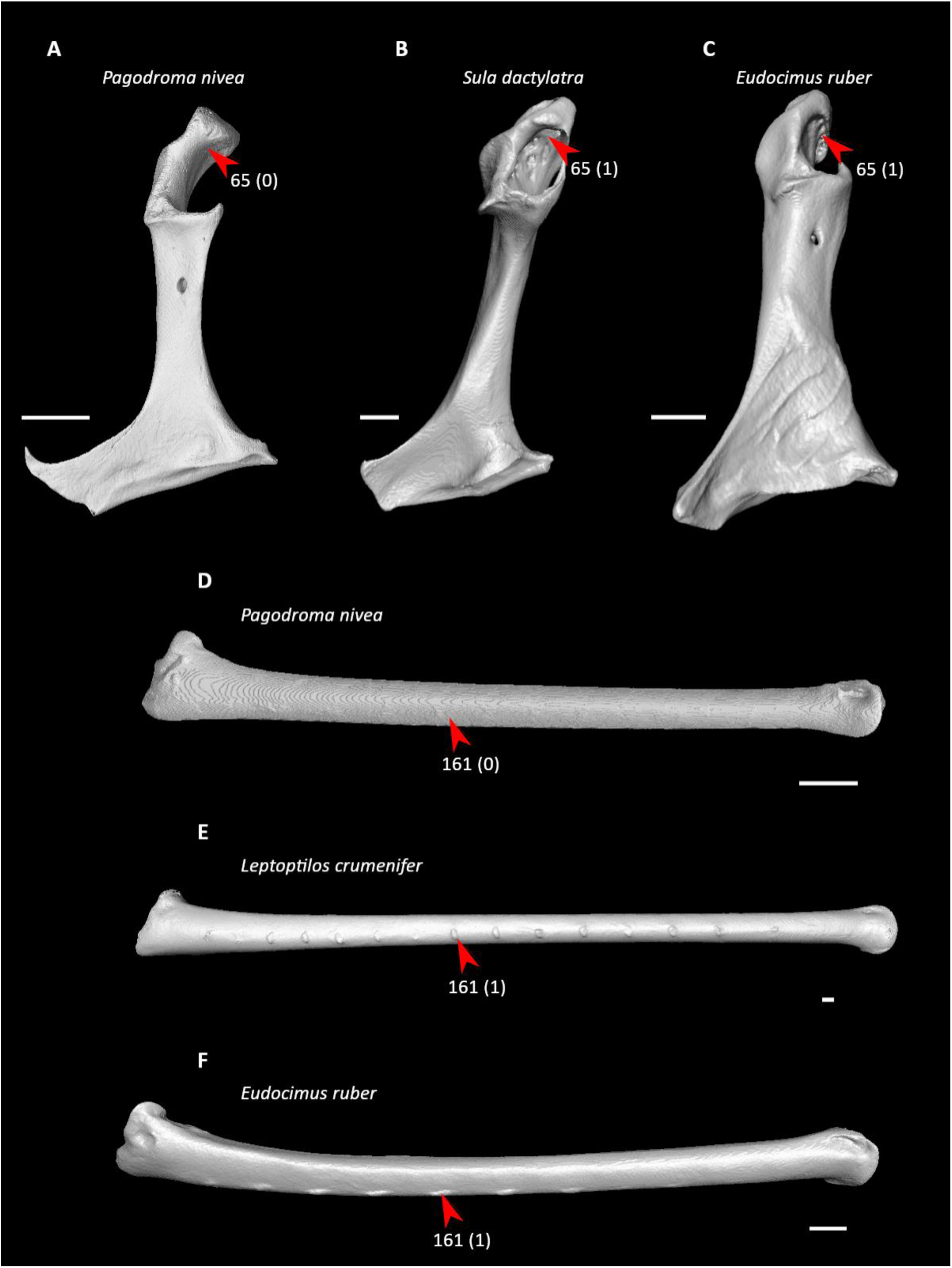
Coracoids of *Pagodroma nivea* (**A**, NHMUK S1998.55, right element, mirrored), *Sula dactylatra* (**B**, NHMUK 1890.11.3.10, right element, mirrored), and *Eudocimus ruber* (**C**, NHMUK S1999.8.1, left element) in dorsal view, and ulnae of *P. nivea* (**D**, NHMUK S1998.55, right element), *Leptoptilos crumenifer* (**E**, NHMUK S1952.3.182, right element), and *E. ruber* (**F**, NHMUK S1999.8.1, left element, mirrored) in caudal view. Scale bars = 5 mm. Arrows indicate the infra-acrocoracoid recess (**A–C**) and the ulnar shaft (**D–F**). *Pagodroma* exhibits state 0 for character 65 (infra-acrocoracoid recess shallow or absent), whereas *Sula* and *Eudocimus* exhibit state 1 (infra-acrocoracoid recess deep, optimized as a synapomorphy of Pelecanimorphae). *Pagodroma* also exhibits state 0 for character 161 (quill knobs absent or weakly developed), whereas *Leptoptilos* and *Eudocimus* exhibit state 1 (quill knobs strongly developed, optimized as a synapomorphy of Pelecanimorphae).

Most of these characters cannot be assessed from published descriptions of the oldest known well-corroborated members of Pelecanimorphae (Smith and Ksepka, 2015), *Limnofregata* (Olson, 1977; Olson and Matsuoka, 2005) and *Rhynchaeites* (Peters, 1983; Mayr, 2002b; Mayr and Bertelli, 2011; Mayr and Kitchener, 2023c), though Olson (1977) noted that the quill knobs of *Limnofregata* are “not nearly as well developed as” in *Fregata*, and the acromial processes on the furcula of *Rhynchaeites* appear to be more pointed than in crown threskiornithids (Fig. 7 in Mayr and Kitchener, 2023c). The ulna of *Limnofregata* exceeds 40% of the combined length of the humerus, ulna, and carpometacarpus (Olson, 1977), whereas the ulna represents approximately 40% or less of this length in *Rhynchaeites*.

Compared to many other members of Pelecanimorphae, the infra-acrocoracoid recess is modestly developed in *Leptoptilos* and apparently other storks (Ciconiidae), including the Miocene *Grallavis* (Fig. 2 in De Pietri and Mayr, 2014). A deep infra-acrocoracoid recess may therefore be a synapomorphy of a less inclusive clade, such as Pelecanes.

#### Pelecanes

Four character states were optimized as potential synapomorphies of Pelecanes, of which two were consistently inferred as such across all alternative molecular topologies studied.

- Internal lip of coracoid expanded along entire width (char. 87: 1 > 0, reversed in *Leucocarbo* and *Scopus* + *Pelecanus*, also found in Galloanserae, Columbimorphae, *Aramus*, Charadriiformes, *Gavia*, *Pagodroma*, and Telluraves); S24, W24.
- Scapular acromion extends distinctly craniad of coracoid tubercle (Fig. 28, char. 89: 0 > 1, reversed in *Tigrisoma*; also found in most non-aequornithean birds); J14, P15, K21, S24, W24.
- Moderate ventral curvature of scapula (char. 92: 1 > 0, reversed in *Tigrisoma*; also found in *Phaethon*, *Gavia*, *Phoebastria*, and most birds outside of Gruiformes, *Opisthocomus*, and some Strisores and Telluraves); S24.
- Distal radius strongly curved dorsoventrally (char. 147: 0 > 1, reversed in *Leucocarbo* and *Pelecanus*; also found in *Anseranas*, *Ortalis* + *Rollulus*, *Eurypyga*, *Pagodroma*, and many non-phaethoquornithean neoavians); J14, P15, K21, S24, W24.

Both *Limnofregata* (Olson, 1977) and *Rhynchaeites* (Mayr and Kitchener, 2023c) exhibit an acromion process that extends cranial to the coracoid tubercle on the scapula, though this is expressed to a lesser extent in the former compared to most extant members of Pelecanes other than herons (Ardeidae) (Olson, 1977). Previously proposed morphological synapomorphies for Pelecanes concern features of the hindlimbs (Sangster et al., 2022).

#### Suliformes

Six character states were optimized as potential synapomorphies of Suliformes, of which five characters were consistently inferred as such across all alternative molecular topologies studied.

- Sternal keel ends caudally before termination of median trabecula (Fig. 31, char. 48: 0 > 1, also found in *Gansus*, Anseriformes, *Phoenicopterus*, *Pterocles*, *Corythaeola*, *Balearica*, *Alca*, *Phaethon*, *Gavia*, *Phoebastria*, *Pelecanus*, *Elanus*, *Psilopogon*, *Acanthisitta*, and *Menura*); J14, P15, K21, S24, W24.
- Median trabecula of sternum squared off at caudal termination (char. 59: 0 > 1, reversed in *Sula*; also found in *Phaethon*, *Tigrisoma*, and many non-phaethoquornithean birds); J14, P15, K21, S24, W24.
- Caudolateral trabeculae of sternum extend caudal of median trabecula (char. 60: 0 > 1, also found in *Gansus*, *Ichthyornis*, *Dendrocygna*, Mirandornithes, Strisores, Heliornithes, *Phaethon*, *Spheniscus*, *Phoebastria*, *Scopus*, *Pelecanus*, *Pandion* + *Elanus*, *Tyto*, Pici, *Micrastur*, and *Nestor*); J14, P15, K21, S24, W24.
- External lip of coracoid greatly expanded (Fig. 31, char. 84: 0 > 1, also found in Columbimorphae, *Ardeotis*, *Tapera*, Grues, *Rostratula*, *Alca*, *Sterna*, *Oceanites*, *Scopus*, *Pelecanus*, *Coragyps*, *Pandion*, *Leptosomus*, *Upupa*, Pici, and *Nestor*); J14, P15, K21, S24, W24.
- Transverse sulcus on humerus long (char. 115: 1 > 2, also found in *Limosa*, *Sterna*, and *Phaethon*); J14, P15, K21, S24, W24.
- Olecranon fossa on humerus deep (char. 127: 0 > 1, also found in Phaethontimorphae, Procellariimorphae, *Scopus* + *Pelecanus*, and most non-phaethoquornithean neognaths); W24.

**Figure 31.**
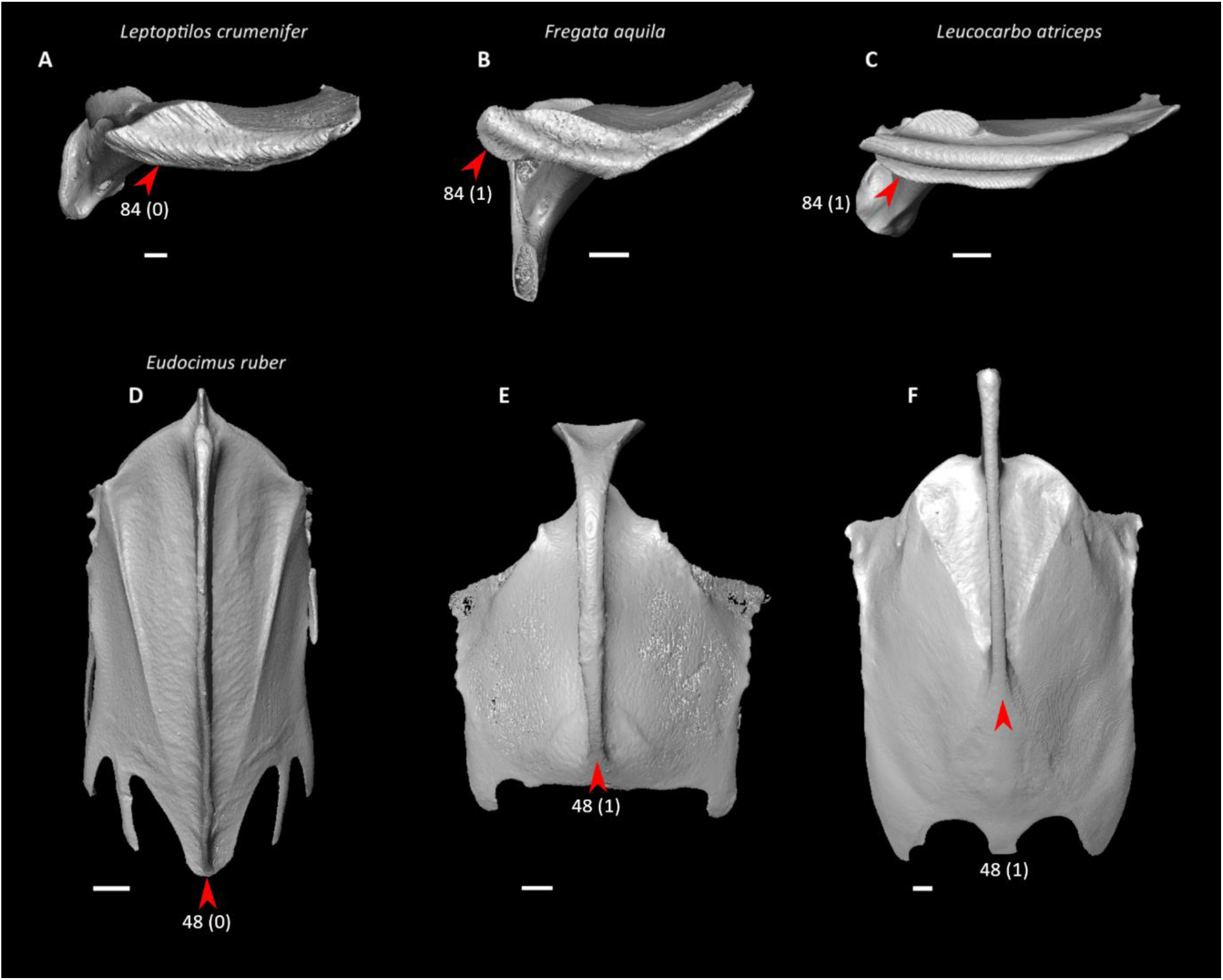
Coracoids of *Leptoptilos crumenifer* (**A**, NHMUK S1952.3.182, left element, mirrored), *Fregata aquila* (**B**, NHMUK 1890.11.3.3, right element), and *Leucocarbo atriceps* (**C**, NHMUK S2012.36, right element) in sternal view and sterna of *Eudocimus ruber* (**D**, NHMUK S1999.8.1), *F. aquila* (**E**, NHMUK 1890.11.3.3), and *L. atriceps* (**F**, NHMUK S2012.36) in ventral view. Scale bars = 5 mm. The furcula of *F. aquila*, which is fused to the sternum and coracoid, has been digitally removed. Arrows indicate the external lip of the coracoid (**A–C**) and the caudal termination of the sternal keel (**D–F**). *Leptoptilos* exhibits state 0 for character 84 (little ventral expansion of external lip of coracoid), whereas *Fregata* and *Leucocarbo* exhibit state 1 (prominent ventral expansion of external lip of coracoid, optimized as a synapomorphy of Suliformes). *Eudocimus* exhibits state 0 for character 48 (sternal keel extending to caudal terminus of median trabecula), whereas *Fregata* and *Leucocarbo* exhibit state 1 (sternal keel ending cranial to caudal terminus of median trabecula, optimized as a synapomorphy of Suliformes).

Traditionally, members of Suliformes were thought to be closely related to pelicans (Pelecanidae) based on anatomical similarities (e.g., Cracraft, 1985; Mayr, 2003b; Livezey and Zusi, 2007; Smith, 2010), and indeed three of the above traits are also found in *Pelecanus*, emphasizing the morphological convergence between these two separate lineages within Pelecanes. *Phaethon*, which as discussed previously was also once hypothesized to form a clade with *Pelecanus* and members of Suliformes, shares four of these traits.

Of these characters, *Limnofregata* has been reported to possess caudolateral trabeculae that extend caudal to the median trabecula of the sternum and an expanded external lip on the coracoid (though to a lesser extent than in *Fregata*) (Olson, 1977). Furthermore, though Olson (1977) considered the sternal keel of *Limnofregata* and *Fregata* to traverse the entire length of the sternum, a flattened region caudal to the keel is nonetheless visible on the ventral surface of the sternum in these taxa.

Of note is that the extinct clade Plotopteridae, a group of flightless marine neoavians that have been alternatively considered either members of Suliformes (Olson, 1980; Smith, 2010, Mayr et al., 2015; Mayr et al., 2021) or closely related to spheniscids (Mayr, 2005b; Mayr et al., 2015), exhibit at least two of the potential suliform synapomorphies listed above that are absent in spheniscids, namely a sternal keel that does not extend to the caudal termination of the median trabecula (Olson, 1980) and a greatly expanded external lip of the coracoid (Mayr et al., 2021). In addition, plotopterids possess an elongated acromion process of the scapula (Olson, 1980; Ando and Fukata, 2018; Mayr et al., 2021), here optimized as a synapomorphy of Pelecanes (though Mayr et al., 2021 reported that some stem-spheniscids display an enlarged acromion compared to that of extant spheniscids, the acromion in these specimens still does not extend as far cranially as it does in most members of Pelecanes besides ardeids). These observations may therefore add to the growing support for a placement of plotopterids within Suliformes and further suggest that their similarities to total-group spheniscids represent a remarkable example of convergent evolution driven by specialization towards a wing-propelled diving lifestyle (Olson, 1980; Smith, 2010; Mayr et al., 2021).

Smith (2010) additionally noted a distinct proximal ridge extending from the ventral supracondylar tubercle on the humerus as well as major and minor metacarpals being subequal in length as potential synapomorphies of Suliformes. However, the latter character state (char. 194: 0) is recovered here as plesiomorphic for neognaths. Mayr (2011c) recovered the absence of a supracoracoid nerve foramen as a suliform synapomorphy, but this character state (char. 70: 0) is inferred to be plesiomorphic for crown birds in this study. Using the phylogenetic matrix of Livezey and Zusi (2006), Ericson and Qu (2024) inferred a caudally inflated flexor process on the humerus and the presence of an interclavicle dorsal process on the furcula (char. 7: 1) as potential synapomorphies of Suliformes, though the latter character state was not optimized as such in the present study as it was only observed in *Leucocarbo* among the suliform taxa sampled here.

#### Pelecaniformes

One character state was optimized as a potential synapomorphy of Pelecaniformes, which was consistently inferred as such across all alternative molecular topologies studied.

- Procoracoid process of coracoid moderately prominent (char. 68: 2 > 1, also found in Anseres, *Phoenicopterus*, *Ardeotis*, *Nyctibius*, *Streptoprocne*, *Sarothrura*, *Rallus*, *Eurypyga*, *Gavia*, *Phoebastria*, Accipitrimorphae, *Trogon*, and *Nestor*); J14, P15, K21, S24, W24.

However, *Rhynchaeites* exhibits a more elongate procoracoid process than extant threskiornithids do (Mayr and Kitchener, 2023c), which is here optimized as a plesiomorphic trait for Phaethoquornithes. Similarly, the putative stem-threskiornithid *Vadaravis* has an elongated procoracoid process (Smith et al., 2013). Also observable in *Vadaravis* are a foramen in the median sulcus of the sternum near its cranial margin and a tubercle at the insertion point of humerocarpal ligament on the pisiform (Smith et al., 2013), suggesting that a phylogenetic placement of this genus within Pelecanimorphae is plausible. Unlike most extant members of Pelecanes other than ardeids, however, *Vadaravis* has a short acromion process of the scapula (Smith et al., 2013).

#### Telluraves

Four character states were optimized as potential synapomorphies of Telluraves, but none of these were consistently inferred as such across all alternative molecular topologies studied.

- Protruding articular facets for acrocoracoids on furcula prominent (char. 3: 0 > 1, reversed in *Tyto*, *Ninox* [though present in most other Strigidae, Mayr, 2005c; Mayr and Kitchener, 2023d], *Trogon*, *Upupa*, *Coracias*, and *Nestor*; also found in *Columba*, *Corythaeola*, *Nyctibius*, Daedalornithes, *Alca*, *Sterna*, *Spheniscus*, *Oceanites*, Suloidea, *Scopus*, and *Pelecanus*); J14, K21, S24, W24.
- Keel low, less than two-thirds total height of sternum (char. 47: 1 > 0, reversed in *Upupa*, Psittaciformes, and *Menura*; also found in *Chauna*, *Anseranas*, Mirandornithes, *Tapera*, Vanescaves, *Opisthocomus*, *Psophia*, Charadrii, Limosa, and Phaethoquornithes), W24.
- Ulna greater than 40% of humerus + ulna + carpometacarpus length (char. 98: 0 > 1, reversed in *Urocolius*, *Cariama*, *Nestor*, and *Menura*; also found in *Gansus*, *Ichthyornis*, *Chauna*, *Phoenicopterus*, *Columba*, *Ardeotis*, Strisores, Grues, *Burhinus*, *Rostratula*, *Sterna*, Phaethontimorphae, *Phoebastria*, and Pelecanimorphae); P15.
- Prominent tubercle on minor metacarpal distal to proximal synostosis of metacarpals (Fig. 32, char. 186: 0 > 1, reversed in *Pandion*, *Urocolius*, Eucavitaves [regained in total-group Coracioidea, Mayr and Mourer-Chauviré, 2000; Mayr et al., 2004; Bourdon et al., 2016; Mayr, 2022b], and Psittacopasseres; also found in *Ichthyornis*, *Eudromia*, *Rollulus*, *Psophia*, *Burhinus*, *Turnix*, *Sterna*, and *Phaethon*); J14, P15, K21.

**Figure 32.**
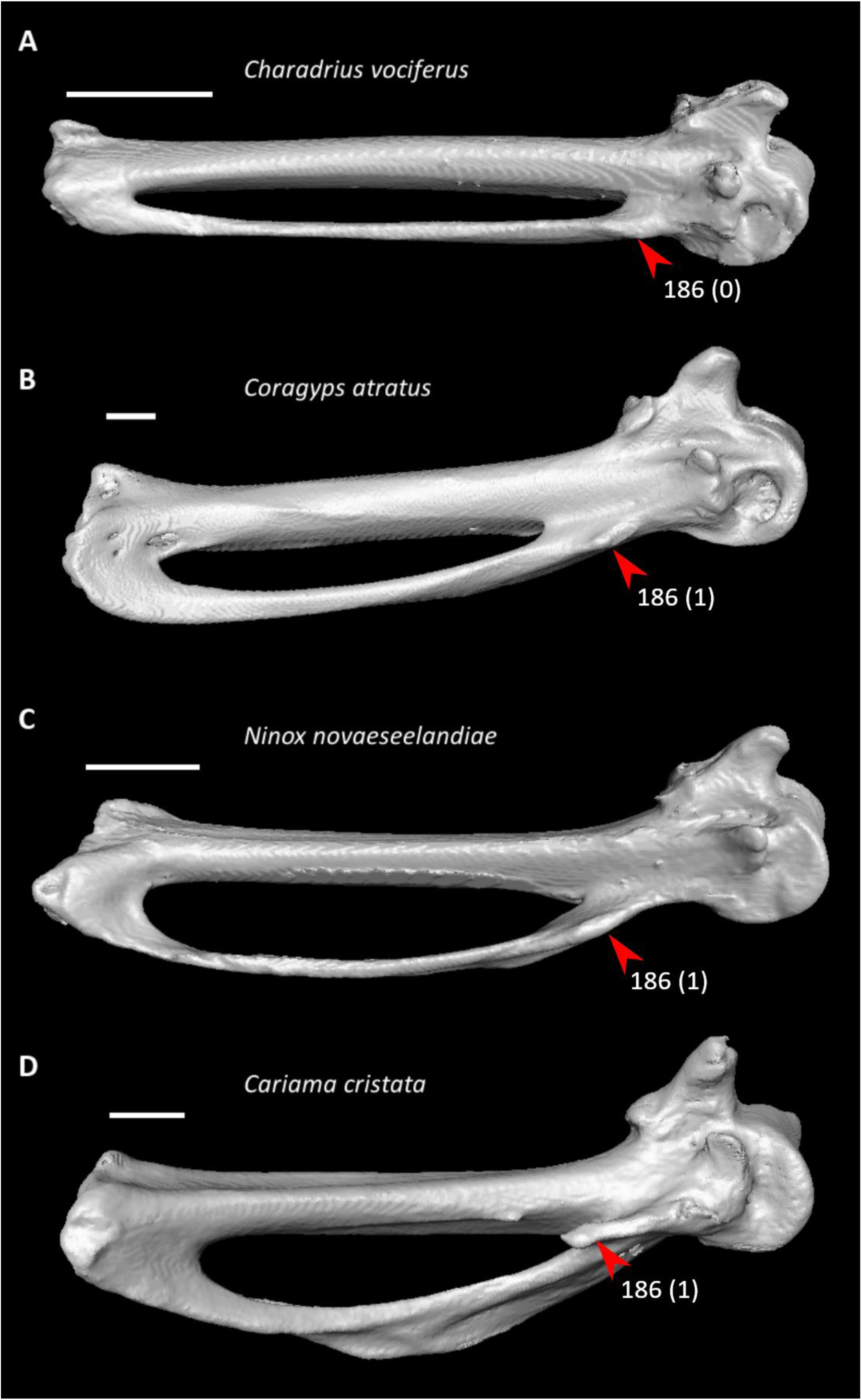
Carpometacarpi of *Charadrius vociferus* (**A**, FMNH 470173, left element, mirrored), *Coragyps atratus* (**B**, UMMZ 71891, right element), *Ninox novaeseelandiae* (**C**, UMZC 497.B, left element, mirrored), and *Cariama cristata* (**D**, FMNH 105653, right element) in ventral view. Scale bars = 5 mm. Arrows indicate the tubercle at the base of the minor metacarpal. *Charadrius* exhibits state 0 for character 186 (tubercle on minor metacarpal small or absent), whereas *Coragyps*, *Ninox*, and *Cariama* exhibit state 1 (tubercle on minor metacarpal prominent, optimized as a synapomorphy of Telluraves).

Probably owing to the extensive morphological disparity of this clade, few anatomical synapomorphies of Telluraves have previously been identified (Mayr, 2011a; Sangster et al., 2022). Even so, potential support for some of the proposed synapomorphies above may be found in the fossil record. The presence of a prominent ventral tubercle on the minor metacarpal has been documented in the stem-mousebird (stem-Coliidae) Sandcoleidae (Houde and Olson, 1992), the putative stem-courol (stem-Leptosomidae) *Waltonavis* (Mayr and Kitchener, 2023e), and many stem-seriemas (stem-Cariamidae) (Mourer-Chauviré, 1983; Mayr, 2016b). A relatively elongate ulna (longer than the humerus) has been previously noted as being common in “higher land birds” (Mayr, 2005d; Mayr et al., 2004).

Prominent articular facets for the acrocoracoid are present on the furcula of the stem-passeriform (Mayr, 2004c; Mayr, 2008b; Mayr, 2015c; Ksepka et al., 2019; Mayr and Kitchener, 2023e) clade Zygodactylidae (Mayr, 1998b; Mayr, 2008b; Mayr and Kitchener, 2023f), but lacking in the stem-owl (stem-Strigiformes) *Ypresiglaux* (Mayr and Kitchener, 2023d), sandcoleids (Houde and Olson, 1992), *Waltonavis* and the stem-leptosomid *Plesiocathartes* (Mayr and Kitchener, 2023e), the stem-falcon (stem-Falconidae) clade Masillaraptoridae (Mayr and Kitchener, 2021), and the putative stem-psittacopasserean (Mayr, 2015c; Ksepka et al., 2019; Mayr and Kitchener, 2023f) clades Messelasturidae (Mayr, 2005c; Mayr and Kitchener, 2023f) and Halcyornithidae (Ksepka et al., 2011; Mayr and Kitchener, 2024e), suggesting that this character state may have evolved multiple times within Telluraves.

#### Accipitrimorphae

Seven character states were optimized as potential synapomorphies of Accipitrimorphae, of which four were consistently inferred as such across all alternative molecular topologies studied.

- Acromial processes on furcula strongly caudally inflected (char. 6: 0 > 1, also found in *Chauna*, *Rollulus*, *Pterocles*, *Florisuga*, *Rostratula*, *Alca*, *Gavia*, *Spheniscus*, *Fregata*, *Leucocarbo*, *Pelecanus*, *Ninox*, *Urocolius*, *Upupa*, *Bucco*, and *Psilopogon*); P15.
- Median trabecula of sternum rounded at caudal termination (char. 59: 1 > 0, also found in non-neognath birds, *Ortalis* + *Rollulus*, *Monias*, *Columba*, *Florisuga*, Gruiformes, *Turnix*, *Alca*, *Eurypyga*, Aequornithes, *Tyto*, *Urocolius*, *Bucorvus*, *Bucco*, Psittaciformes, and *Menura*); P15.
- Procoracoid process of coracoid moderately prominent (char. 68: 2 > 1, also found in Anseres, *Phoenicopterus*, *Ardeotis*, *Nyctibius*, *Streptoprocne*, *Sarothrura*, *Rallus*, *Eurypyga*, *Gavia*, *Phoebastria*, Pelecaniformes, *Trogon*, and *Nestor*); J14, P15, K21, S24, W24.
- Supracoracoid nerve foramen in coracoid present (char. 70: 0 > 1, also found in *Ichthyornis*, *Anseranas*, *Phoenicopterus*, *Corythaeola*, Daedalornithes, Gruiformes, Charadrii, *Alca*, *Sterna*, *Phaethon*, Procellariiformes, *Leptoptilos*, *Eudocimus*, *Pelecanus*, Strigiformes, *Leptosomus*, and *Micrastur*); P15.
- Dorsal tubercle of humerus pointed (char. 139: 0 > 1, also found in *Dendrocygna*, Mirandornithes, *Podargus*, *Aegotheles*, *Balearica*, *Charadrius*, *Alca*, Procellariiformes, *Leptoptilos*, Suloidea, *Scopus* + *Pelecanus*, *Leptosomus*, *Trogon*, *Coracias*, *Micrastur*, *Psittacus*, and Eupasseres); J14, P15, K21, S24, W24.
- Radial incisure on ulna prominent (char. 156: 0 > 1, also found in *Ichthyornis*, *Eudromia*, *Chauna*, *Anseranas*, Mirandornithes, *Ardeotis*, *Nyctibius*, Charadriiformes, Grues, *Phaethon*, *Phoebastria*, *Pagodroma*, *Leptoptilos*, *Fregata*, *Sula*, *Scopus* + *Pelecanus*, and Psittaciformes); J14, P15, K21, S24, W24.
- Quill knobs on ulna strongly developed (char. 161: 0 > 1, also found in *Anseranas*, *Podilymbus*, *Monias*, *Columba*, *Corythaeola*, *Ardeotis*, *Tapera*, *Caprimulgus*, *Aramus*, *Charadrius*, Scolopaci + Lari, *Oceanites*, Pelecanimorphae, Picocoraciades, and Passeriformes); J14, P15, K21, S24, W24.

Using the phylogenetic matrix of Livezey and Zusi (2006), Ericson and Qu (2024) also inferred the presence of an impression for the m. coracobrachialis caudalis placed within an impression for the m. supracoracoideus on the sternum as a potential synapomorphy of this clade. Few fossils that bear on the ancestral morphology of Accipitrimorphae have been described, though *Horusornis*, which may have close affinities to this clade, exhibits a moderately prominent procoracoid process and a supracoracoid nerve foramen in the coracoid (Mourer-Chauviré, 1991). These features are also found in the Oligocene hawk (Accipitridae) *Archaehierax*, which additionally exhibits a prominent radial incisure on the ulna and ulnar quill knobs comparable to those of extant accipitrids (Mather et al., 2022).

#### Coraciimorphae

Six character states were optimized as potential synapomorphies of Coraciimorphae, of which two were consistently inferred as such across all alternative molecular topologies studied.

- Cranial margin of sternal keel not recurved (char. 42: 1 > 0, reversed in *Trogon*, *Bucorvus*, *Alcedo*, and *Jynx*; also found in *Chauna*, *Dendrocygna*, *Monias*, *Corythaeola*, *Sarothrura*, *Scopus* + *Pelecanus*, *Pandion*, *Tyto*, and Psittaciformes); S24, W24.
- Longitudinal concavity of scapula prominent (char. 93: 0 > 1, reversed in Picocoraciades; also found in *Gansus*, *Anseranas*, Galliformes, Pteroclimesites, *Tapera*, *Opisthocomus*, *Aramus*, Heliornithes, *Eudocimus*, *Cariama*, *Micrastur*, and Tyranni); J14, K21.
- Ventral supracondylar tubercle on humerus expanded (Fig. 33, char. 141: 0 > 1, reversed in Picocoraciades; also found in *Ichthyornis*, *Chauna*, Galliformes, *Columba*, *Tapera*, *Limosa*, *Turnix*, *Gavia*, *Oceanites*, *Fregata*, *Leucocarbo*, and *Climacteris*); J14, P15, K21, S24, W24.
- Ventral collateral ligamental tubercle on ulna strongly developed (char. 158: 0 > 1, reversed in *Bucorvus* and *Merops*; also found in *Ichthyornis*, *Ortalis*, *Corythaeola*, *Tapera*, Strisores, *Psophia*, Charadriiformes, Procellariiformes, *Fregata*, *Pelecanus*, *Elanus*, Strigiformes, *Nestor*, and Passeriformes); W24.
- Distal extent of dorsal rim of carpal trochlea falls considerably short of ventral rim (Fig. 33, char. 177: 1 > 0, reversed in *Trogon*; also found in *Eudromia*, Galliformes, *Monias*, *Corythaeola*, *Psophia*, *Sarothrura*, *Rallus*, *Burhinus*, *Gavia*, *Spheniscus*, *Fregata*, *Cariama*, and Passeriformes); J14, P15, K21, S24, W24.
- Minor metacarpal projecting substantially distally beyond major metacarpal (char. 194: 0 > 1, reversed in *Coracias*; also found in *Eudromia*, *Alectura*, *Ortalis*, *Monias*, *Corythaeola*, *Caprimulgus*, *Florisuga*, *Psophia*, *Spheniscus*, *Elanus*, *Ninox*, *Micrastur*, and Passeriformes); S24, W24.

**Figure 33.**
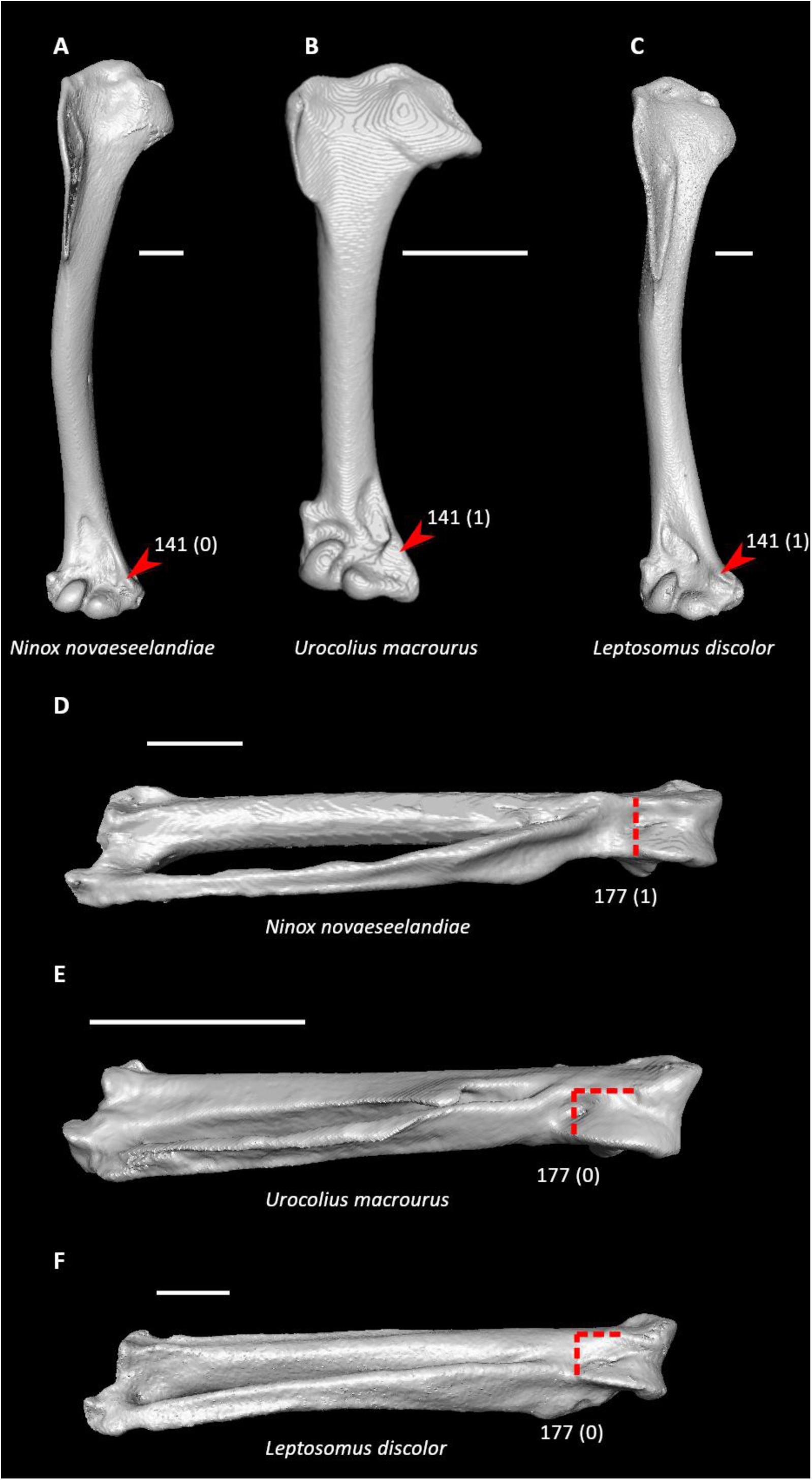
Humeri of *Ninox novaeseelandiae* (**A**, UMZC 497.B, right element), *Urocolius macrourus* (**B**, FMNH 368959, left element, mirrored), and *Leptosomus discolor* (**C**, NHMUK A1968.30.38, left element, mirrored) in cranial view and carpometacarpi of *N. novaeseelandiae* (**D**, UMZC 497.B, left element), *U. macrourus* (**E**, FMNH 368959, right element, mirrored), and *L. discolor* (**F**, NHMUK A1968.30.38, left element) in caudal view. Scale bars = 5 mm. Arrows indicate the ventral supracondylar tubercle (**A–C**) and dotted lines indicate the distal extent of the ventral rim of the carpal trochlea relative to that of the dorsal rim (**D–F**). *Ninox* exhibits state 0 for character 141 (ventral supracondylar tubercle moderately developed), whereas *Urocolius* and *Leptosomus* exhibit state 1 (ventral supracondylar tubercle expanded, forming large triangular platform approximately equal in width to ventral condyle and in dorsal extent to dorsal condyle, optimized as a synapomorphy of Coraciimorphae). *Ninox* also exhibits state 1 for character 177 (dorsal rim of carpal trochlea subequal in distal extent to that of ventral rim), whereas *Urocolius* and *Leptosomus* exhibit state 0 (dorsal rim of carpal trochlea falls considerably short of ventral rim in distal extent, optimized as a synapomorphy of Coraciimorphae).

The status of these character states is largely unclear in fossil specimens that bear on the ancestral morphology of Coraciimorphae, such as *Plesiocathartes* (Mayr, 2002c; Weidig, 2006; Mayr and Kitchener, 2023e). However, an expanded ventral supracondylar tubercle on the humerus is present in sandcoleids (Houde and Olson, 1992; Mayr and Peters, 1998; Ksepka et al., 2017), *Waltonavis* (Fig. 9g in Mayr and Kitchener, 2023e), and stem-trogons (stem-Trogonidae) (Mayr et al., 2023).

#### Cavitaves

Six character states were optimized as potential synapomorphies of Cavitaves, but none of these were consistently inferred as such across all alternative molecular topologies studied.

- Keel apex on sternum cranial to main body of sternum (char. 41: 0 > 1, reversed in *Bucco*; also found in *Cariama*, Psittaciformes, Eupasseres, and many non-telluravian birds); J14, P15, K21.
- Acrocoracoid process on coracoid forms pronounced hook (char. 63: 0 > 1, reversed in *Psilopogon*; also found in *Ichthyornis*, *Eudromia*, *Alectura*, *Rollulus*, *Podilymbus*, *Pterocles*, *Columba*, *Corythaeola*, *Tapera*, *Aegotheles*, *Streptoprocne*, *Podica*, Charadriiformes, Phaethontimorphae, Procellariimorphae, *Cariama*, Psittaciformes, and Eupasseres); J14, P15, K21.
- Deltopectoral crest well developed, extending at least one third the total length of the humerus (char. 122: 1 > 0, reversed in Coraciiformes; also found in *Gansus*, *Ichthyornis*, *Corythaeola*, *Ardeotis*, *Nyctibius*, *Podargus*, *Spheniscus*, *Leptoptilos*, *Leucocarbo*, *Eudocimus*, *Scopus*, Accipitrimorphae, Strigiformes, *Micrastur*, and Passeri); W24.
- Humerus narrowest near midshaft point (char. 126: 0 > 2, reversed in *Merops*, *Alcedo*, and *Bucco*; also found in *Ichthyornis*, *Alectura*, *Rollulus*, *Pterocles*, *Columba*, *Corythaeola*, *Tapera*, Strisores, *Opisthocomus*, *Podica*, Charadrii, *Rostratula*, *Sterna*, *Eurypyga*, *Spheniscus*, *Oceanites*, *Leptoptilos*, *Pelecanus* + *Tigrisoma*, *Pandion*, *Cariama*, Psittaciformes, and Eupasseres); J14, P15, K21.
- Radius craniocaudally curved (char. 144: 0 > 1, reversed in *Coracias*; also found in Anseriformes, *Phoenicopterus*, *Ardeotis*, Strisores, Grues, Charadrii, Feraequornithes, *Coragyps*, *Pandion*, Strigiformes, *Micrastur*, Tyranni, and *Climacteris*); J14, P15, K21, W24.
- Ventral interosseal sulcus on carpometacarpus forms prominent groove (char. 196: 0 > 1, reversed in *Bucorvus*, *Coracias*, and *Alcedo*; also found in *Sarothrura*, *Gavia*, *Fregata*, *Scopus*, Strigiformes, *Psittacus*, *Erythropitta*, and Passeri); S24, W24.

A hooked acrocoracoid process on the coracoid has been reported in *Waltonavis* (Mayr and Kitchener, 2023e), *Plesiocathartes* (Mayr, 2002c), and stem-trogonids (Mayr, 1999b; Mayr, 2009; Mayr et al., 2023). This character state is also found in *Ypresiglaux* (Mayr and Kitchener, 2023d) and sandcoleids (Houde and Olson, 1992; Ksepka et al., 2017), however, which suggests that it could be a synapomorphy of a more inclusive clade. *Waltonavis* additionally has a narrow humeral midshaft (Fig. 9g in Mayr and Kitchener, 2023e), but the remaining characters listed above are otherwise difficult to assess in relevant fossil taxa.

Although not optimized as an unambiguous synapomorphy of Cavitaves, a bifurcated acromion process of the scapula (char. 88: 1) may also be synapomorphic for this group, as it is found in all cavitavians sampled in the present study except for *Leptosomus*, *Bucorvus*, and *Bucco*. A slightly bifurcated acromion process has been reported in *Waltonavis* (Mayr and Kitchener, 2023e); the stem-trogonid *Eotrogon* (Mayr et al., 2023c); a potential stem member of the hoopoe + woodhoopoe clade (Upupides), *Messelirrisor* (Mayr, 2000c); stem members of the roller + ground roller clade (Coracioidea), *Septencoracias* (Mayr and Kitchener, 2024f; *contra* Bourdon et al., 2016) and *Geranopterus* (Mayr and Mourer-Chauviré, 2000); the putative stem-piciform *Gracilitarsus* (Mayr, 2005e); and the stem-Pici taxon *Rupelramphastoides* (Mayr, 2005f). However, this character is also present in *Ypresiglaux* (Mayr and Kitchener, 2023d) and the stem-coliid *Ypresicolius* (Mayr and Kitchener, 2024g). If Strigiformes and Coraciimorphae are extant sister taxa, as recovered by Jarvis et al. (2014), Prum et al. (2015), and Kuhl et al. (2021), this may hint at multiple evolutionary acquisitions and losses of this feature within the clade uniting both.

#### Eucavitaves

Five character states were optimized as potential synapomorphies of Eucavitaves, of which four were consistently inferred as such across all alternative molecular topologies studied.

- Notch in medial margin of sternal end of coracoid (char. 75: 0 > 1, reversed in *Bucorvus* and *Merops*; also found in *Rollulus*, *Tapera*, *Podica*, *Burhinus*, *Alca*, *Spheniscus*, and Passeriformes); J14, P15, K21, S24, W24.
- Deltopectoral crest rounded dorsally (char. 121: 1 > 0, reversed in *Upupa*, *Merops*, and *Bucco*; also found in *Gansus*, *Chauna*, *Anseranas*, Mirandornithes, *Ardeotis*, *Aegotheles*, *Opisthocomus*, Gruiformes, *Charadrius*, *Rostratula*, *Alca*, *Eudocimus*, and Australaves); J14, P15, K21, S24, W24.
- Dorsal supracondylar process on humerus extremely small (char. 136: 1 > 0, reversed in *Upupa*, *Coracias*, and *Jynx*; also found in *Podilymbus*, *Aegotheles*, *Opisthocomus*, *Balearica*, *Spheniscus*, *Sula*, and Psittaciformes); J14, P15, K21, S24, W24.
- Dorsal epicondyle on humerus prominent (char. 138: 0 > 1, reversed in *Coracias*; also found in *Dendrocygna*, Galliformes, Daedalornithes, *Psophia*, *Phaethon*, *Eudocimus*, *Pelecanus*, and Passeriformes); J14, P15, K21, S24, W24.
- Ventral aponeurosis tubercle on radius pointed (char. 149: 0 > 1, reversed in *Merops* and Piciformes; also found in *Coragyps*, *Pandion*, *Ninox*, *Psittacus*, *Climacteris*, and many non-telluravian birds); P15.

A notched medial margin on the sternal end of the coracoid was previously considered a synapomorphy of Piciformes by Mayr et al. (2003), though as noted by Mayr (2005e), this character also occurs in other birds, including many taxa now recognized as members of Eucavitaves. Among fossil eucavitavians, it has been reported in *Eotrogon* (Mayr et al., 2023c), putative stem-upupideans (Mayr, 2000c; Mayr, 2006b; Mayr et al., 2020b), *Gracilitarsus* (Mayr, 2005e), the putative stem-piciform (Duhamel et al., 2020; Mayr, 2022a) *Pristineanis kistneri* (Houde and Olson, 1989), and *Rupelramphastoides* (Mayr, 2005f). However, this feature is not as pronounced in stem-trogonids and stem-upupideans as it is in their respective crown groups (Mayr, 2006b; Mayr et al., 2020b; Mayr et al., 2023), and it appears to be absent in the stem-trogonid *Primotrogon* (Mayr, 1999b).

A rounded deltopectoral crest is visible on the humerus of *Eotrogon*, though the same element appears to exhibit a more prominent dorsal supracondylar process and a less prominent dorsal epicondyle than found in crown-group trogonids (Fig. 5 in Mayr et al., 2023c). *Messelirrisor* has a prominent dorsal epicondyle on the humerus (Mayr, 2000c).

#### Picocoraciades

Ten character states were optimized as potential synapomorphies of Picocoraciades, of which eight were consistently inferred as such across all alternative molecular topologies studied.

- Pneumatic foramen in median sulcus of sternum immediately caudal to cranial margin (char. 29: 0 > 1, reversed in Pici; also found in *Ichthyornis*, *Dendrocygna*, *Alectura*, *Ortalis*, *Phoenicopterus*, *Pterocles*, *Ardeotis*, *Tapera*, Strisores, *Opisthocomus*, Grues, *Podica*, *Phaethon*, Pelecanimorphae, *Coragyps*, Strigiformes, *Micrastur*, Psittaciformes, *Neopelma*, and *Menura*); J14, P15, K21, S24, W24.
- Internal lip of coracoid expanded in lateral portion (char. 87: 0 > 2, reversed in *Coracias* and *Psilopogon*; also found in *Aegotheles* and *Oceanites*); J14, S24, W24.
- Longitudinal concavity of scapula absent or weak (char. 93: 1 > 0, reversed in Pici; also found in Accipitrimorphae, Strigiformes, Psittacopasseres, and many non-telluravian birds); J14, P15, K21, S24, W24.
- Cranial surface of bicipital crest planar or slightly concave (char. 118: 0 > 1, reversed in *Bucco*; also found in *Turnix*, *Phoebastria*, *Urocolius*, *Psittacus*, *Acanthisitta*, and Tyranni); J14, P15, K21, S24, W24.
- Ventral supracondylar tubercle on humerus moderately developed (char. 141: 1 > 0, reversed in Pici; also found in many non-coraciimorph birds); J14, P15, K21, S24, W24.
- Olecranon process on ulna narrow and sharply projected (char. 159: 0 > 1, also found in *Chauna*, *Caprimulgus*, Apodiformes, *Alca*, *Sula*, and Passeriformes); J14, P15, K21, S24, W24.
- Quill knobs on ulna strongly developed (char. 161: 0 > 1, reversed in *Bucco*; also found in *Anseranas*, *Podilymbus*, *Monias*, *Columba*, *Corythaeola*, *Ardeotis*, *Tapera*, *Caprimulgus*, *Aramus*, *Charadrius*, Scolopaci + Lari, *Oceanites*, Pelecanimorphae, Accipitrimorphae, and Passeriformes); J14, P15, K21, S24, W24.
- Sulcus for tendon of m. extensor longus alulae on scapholunare well defined (char. 164: 0 > 1); J14, P15, K21, S24, W24.
- Tendinal sulcus on carpometacarpus wraps around cranial surface of bone (char. 172: 0 > 1, reversed in *Bucco*; also found in *Nyctibius*, *Florisuga*, *Balearica*, *Coragyps*, *Elanus*, *Tyto*, *Micrastur*, and Passeriformes); S24, W24.
- Dorsal fossa on phalanx 1 of major digit indistinct (char. 203: 1 > 2, reversed in *Coracias* and *Merops*; also found in *Spheniscus*, *Pelecanus*, *Urocolius*, *Acanthisitta*, and *Climacteris*); J14, P15, K21, S24, W24.

Mayr (2014a) previously identified a well-defined sulcus for the tendon of m. extensor longus alulae on the scapholunare as a potential synapomorphy of this clade. The stem-upupideans *Waltonirrisor* (Mayr and Kitchener, 2024f) and *Laurillardia* (Mayr et al., 2020b), the stem-coracioid *Eocoracias* (Mayr and Mourer-Chauviré, 2000), the potential stem-piciform *Sylphornis* (Mourer-Chauviré, 1988b), and *Rupelramphastoides* (Mayr, 2005f) have a well-developed olecranon process on the ulna, though this character is absent in the early coraciiform *Quasisyndactylus* (Mayr, 1998b), *Gracilitarsus* (Mayr, 2001), and *Jacamatia*, a putative stem-member of the jacamar + puffbird clade (Galbulae) (Duhamel et al., 2020). Quill knobs similar to those of extant Upupides have been reported in *Messelirrisor* (Mayr, 2000c) and are markedly developed in *Rupelramphastoides* (Mayr, 2005f; Mayr, 2006c), but are poorly developed or absent in *Laurillardia* (Mayr et al., 2020b), the stem-coracioid *Primobucco* (Ksepka and Clarke, 2010), and *Jacamatia* (Duhamel et al., 2020). *Primobucco* also lacks dorsal pneumatic foramina in the sternum (Ksepka and Clarke, 2010).

#### Picodynastornithes

Two character states were optimized as potential synapomorphies of Picodynastornithes, both of which were consistently inferred as such across all alternative molecular topologies studied.

- Minor metacarpal weakly bowed (Fig. 34, char. 174: 1 > 0, reversed in *Jynx*; also found in *Gansus*, *Ichthyornis*, *Dendrocygna*, Mirandornithes, Apodiformes, Charadriiformes, *Phaethon*, *Gavia*, Procellariimorphae, *Fregata*, *Eudocimus*, *Pandion*, and Psittacopasseres); J14, P15, K21, S24, W24.
- Intermetacarpal process on carpometacarpus well developed (char. 183: 0 > 1, reversed in *Merops*, also found in *Rollulus*. *Florisuga*, *Urocolius*, and Passeriformes); J14, P15, K21, S24, W24.

**Figure 34.**
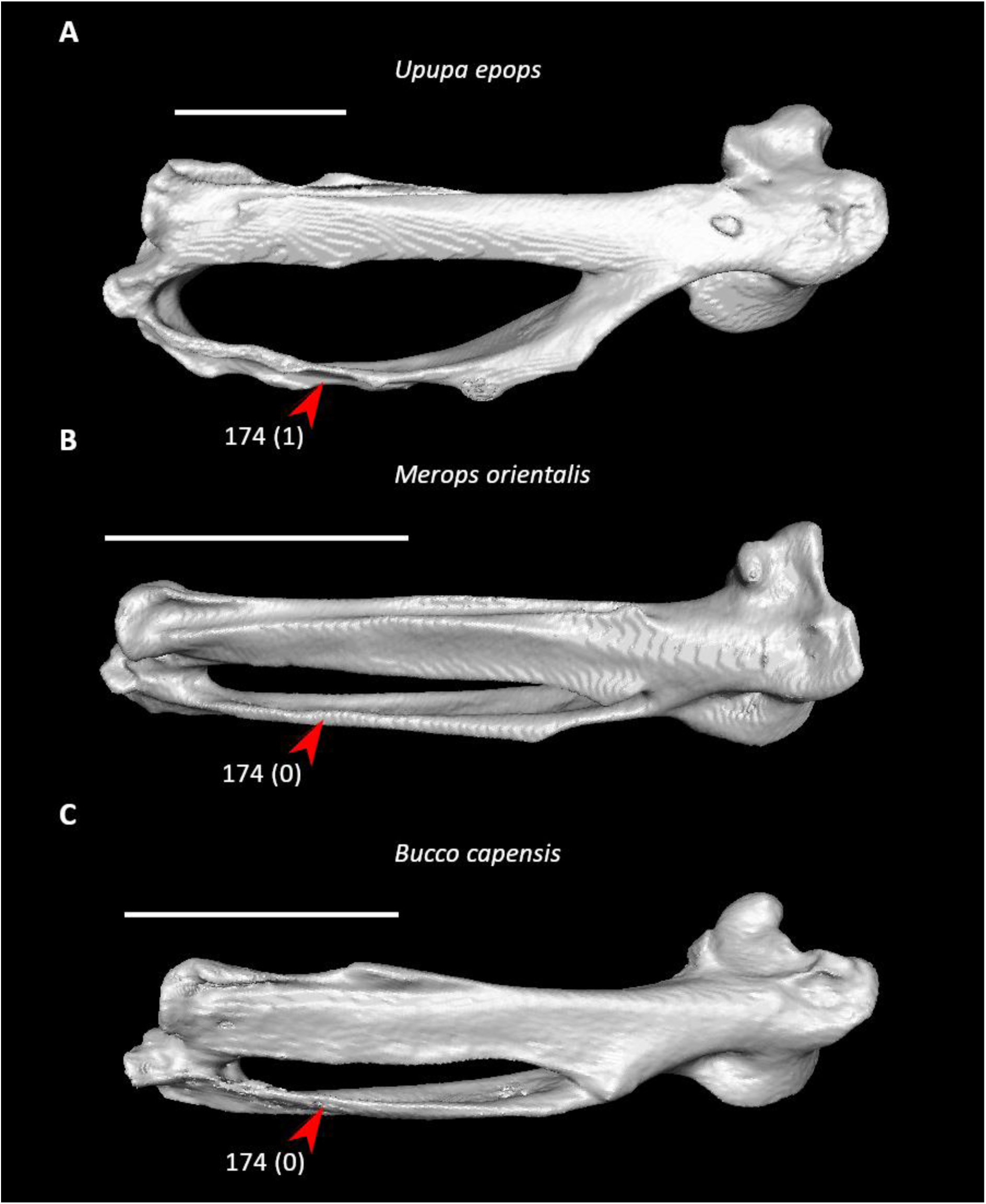
Carpometacarpi of *Upupa epops* (**A**, FMNH 352821, left element), *Merops orientalis* (**B**, UMMZ 216577, right element, mirrored), and *Bucco capensis* (**C**, FMNH 330305, left element) in dorsal view. Scale bars = 5 mm. Arrows indicate the minor metacarpal. *Upupa* exhibits state 1 for character 174 (minor metacarpal strongly bowed), whereas *Merops* and *Bucco* exhibit state 0 (minor metacarpal weakly bowed, optimized as a synapomorphy of Picodynastornithes).

The minor metacarpal is weakly bowed in *Eocoracias* (Mayr and Mourer-Chauviré, 2000), the stem-coracioid *Paracoracias* (Clarke et al., 2009), *Gracilitarsus* (Mayr, 2001; Mayr, 2005e), and *Sylphornis* (Duhamel et al., 2020), though it is more strongly bowed in *Septencoracias* (Mayr, 2022b; Mayr and Kitchener, 2024f) and especially so in *Jacamatia* (Duhamel et al., 2020). A well-developed intermetacarpal process on the carpometacarpus is also found in *Waltonirrisor* (Mayr and Kitchener, 2024f) and *Messelirrisor* (Mayr, 2006b) and may thus represent a synapomorphy of a more inclusive clade. This character appears to have been secondarily reacquired in Coracioidea, as it is absent in stem-coracioids (Mayr and Mourer-Chauviré, 2000; Mayr et al., 2004; Clarke et al., 2009; Bourdon et al., 2016; Mayr, 2022b; Mayr and Kitchener, 2024f).

The above character states, along with some potential synapomorphies of several more inclusive clades like Picocoraciades, Eucavitaves, and Cavitaves, are convergently present in at least some members of Passeriformes, which may have contributed to some members of these groups being hypothesized as close relatives of passeriforms in previous morphological studies (e.g., Höfling and Alvarenga, 2001; Livezey and Zusi, 2007).

#### Australaves

Nine character states were optimized as potential synapomorphies of Australaves, of which three were consistently inferred as such across all alternative molecular topologies studied.

- Pneumatic pores along cranial margin of dorsal sternum (char. 32: 0 > 1, reversed in Passeriformes; also found in Anseriformes, *Ortalis*, *Phoenicopterus*, *Monias*, *Columba*, *Ardeotis*, *Tapera*, *Caprimulgus*, Apodiformes, *Opisthocomus*, Gruoidea, *Podica*, *Phaethon*, *Leptoptilos*, *Sula*, *Eudocimus*, *Scopus*, *Pelecanus*, *Pandion*, Strigiformes, *Trogon*, *Coracias*, and *Merops*); K21, S24.
- Single pair of caudal fenestrae in sternum (char. 49: 2 > 1, reversed in *Acanthisitta*; also found in *Eudromia*, Anseres, Galliformes, Mirandornithes, *Monias*, *Caprimulgus*, *Balearica*, Ralloidea, *Turnix*, Aequornithes, *Pandion* + *Elanus*, *Tyto*, and Bucerotiformes); J14, S24.
- Pneumatic foramen in cranial end of scapula (char. 90: 0 > 1, reversed in *Nestor* and *Acanthisitta*; also found in *Chauna*, *Anseranas*, *Corythaeola*, *Florisuga*, *Opisthocomus*, Gruoidea, *Sula*, *Ninox*, Bucerotiformes, *Coracias*, and *Merops*); J14, P15, K21, S24, W24.
- Deltopectoral crest rounded dorsally (Fig. 35, char. 121: 1 > 0, reversed in *Psittacus* and *Climacteris*; also found in *Gansus*, *Chauna*, *Anseranas*, Mirandornithes, *Ardeotis*, *Aegotheles*, *Opisthocomus*, Gruiformes, *Charadrius*, *Rostratula*, *Alca*, *Eudocimus*, and Eucavitaves); J14, P15, K21, S24, W24.
- Tendinal sulcus of m. scapulotricipitalis on humerus shallow (char. 129: 1 > 0, reversed in Psittaciformes and *Neopelma*; also found in Galloanserae, *Phoenicopterus*, *Monias*, *Corythaeola*, *Opisthocomus*, *Psophia*, *Podica*, *Rallus*, *Eurypyga*, *Gavia*, *Leptoptilos*, *Elanus*, *Leptosomus*, *Bucorvus*, and Picodynastornithes); P15.
- Flexor process on humerus strongly developed (char. 131: 0 > 1, reversed in Psittaciformes; also found in *Monias*, *Corythaeola*, *Tapera*, Heliornithes, *Gavia*, *Spheniscus*, *Urocolius*, Bucerotiformes, *Alcedo*, and *Jynx*); J14, P15, K21, S24, W24.
- Scar of m. flexor carpi ulnaris on flexor process of humerus present as two scars (char. 142: 1 > 2, reversed in *Acanthisitta* and *Erythropitta*; also found in *Pandion*, *Leptosomus*, *Psilopogon*, and many non-telluravian birds); P15.
- Caudal scar for for m. flexor carpi ulnaris on flexor process of humerus shallower than cranial scar (char. 143: 0 > 1, reversed in Psittaciformes; also found in *Chauna*, *Anseranas*, *Podilymbus*, *Ardeotis*, *Podargus*, *Aegotheles*, *Balearica*, *Burhinus*, *Alca*, *Sterna*, *Gavia*, *Pagodroma*, *Sula*, *Eudocimus*, *Tigrisoma*, and *Leptosomus*); J14, K21, S24, W24.
- Internal index on phalanx 1 of major digit poorly developed (char. 201: 1 > 0, reversed in Psittaciformes and Tyranni; also found in *Gansus*, *Eudromia*, *Chauna*, *Alectura*, *Rollulus*, *Podilymbus*, *Monias*, *Corythaeola*, *Opisthocomus*, Gruiformes, *Rostratula*, *Turnix*, *Spheniscus*, *Tigrisoma*, *Ninox*, *Urocolius*, *Bucorvus*, *Merops*, and Piciformes); J14, K21.

**Figure 35.**
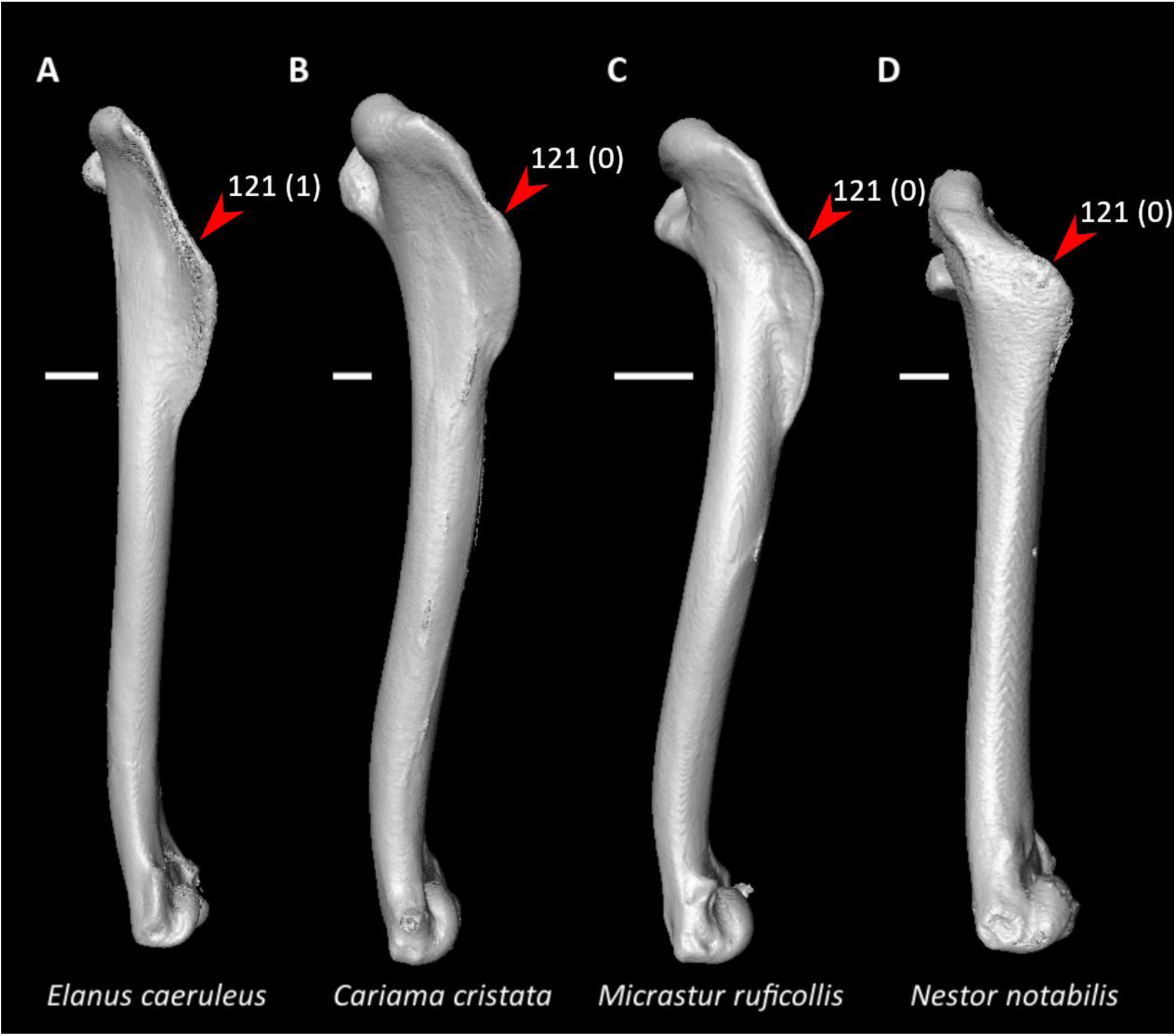
Humeri of *Elanus caeruleus* (**A**, NHMUK 1850.8.15.159, right element), *Cariama cristata* (**B**, FMNH 105653, right element), *Micrastur ruficollis* (**C**, FMNH 330226, right element), and *Nestor notabilis* (**D**, FMNH 23530, right element) in dorsal view. Scale bars = 5 mm. Arrows indicate the proximal dorsal margin of the deltopectoral crest. *Elanus* exhibits state 1 for character 121 (proximal dorsal margin of deltopectoral crest essentially straight), whereas *Cariama*, *Micrastur*, and *Nestor* exhibit state 0 (proximal dorsal margin of deltopectoral crest forming rounded bulge, optimized as a synapomorphy of Australaves).

A proximodorsally rounded deltopectoral crest has been reported in masillaraptorids (Mayr and Kitchener, 2021), messelasturids (Mayr, 2000d; Mayr, 2011d; Mayr and Kitchener, 2023f), halcyornithids (Mayr, 1998c; Fig. 4 in Ksepka et al., 2011; Fig. 3 in Mayr and Kitchener, 2024e), the potentially stem-passeriform (Mayr, 2015c; Ksepka et al., 2019; Mayr and Kitchener, 2023f; Mayr and Kitchener, 2023g) clades Psittacopedidae (Mayr, 2020) and Morsoravidae (Mayr, 1999c), and zygodactylids (Mayr and Kitchener, 2023g). Other character states in the above list have a more variable distribution among fossil taxa with proposed australavian affinities. A shallow scapulotricipital sulcus on the humerus is found in messelasturids (Mayr, 2000d; Mayr, 2005c; Mayr, 2011d), halcyornithids (Mayr, 2002d; Ksepka et al., 2011), and morsoravids (Mayr, 1999c; Mayr and Kitchener, 2023h), but this sulcus is deep in the stem-cariamid *Bathornis* (Mayr, 2016b) and the putative psittacopedid *Eofringillirostrum* (Ksepka et al., 2019). A well-developed flexor process on the humerus is seen in psittacopedids (Mayr and Daniels, 1998; Mayr, 2020), morsoravids (Mayr, 1999c; Mayr and Kitchener, 2023h), and zygodactylids (Mayr, 1998b; Mayr, 2008b; Smith et al., 2018; Mayr and Kitchener, 2023g), and a shorter but still distinct process is present in stem-cariamids (Mayr, 2016b), *Masillaraptor* (Mayr and Kitchener, 2021), and halcyornithids (Mayr, 2002d; Ksepka et al., 2011). In contrast, the humeral flexor process is short in the masillaraptorid *Danielsraptor* (Mayr and Kitchener, 2021), the putative stem-psittacopasserean (Mayr and Kitchener, 2023f) *Vastanavis* (Mayr et al., 2013), and messelasturids (Mayr, 2000d; Mayr, 2005c; Mayr, 2011d; Mayr and Kitchener, 2023f), like in extant Psittaciformes. A poorly developed internal index process on phalanx 1 of the major digit has been found in halcyornithids (Mayr, 2002d; Ksepka et al., 2011), psittacopedids (Mayr and Daniels, 1998; Fig. 5 in Mayr, 2020; Fig 5 in Mayr and Kitchener, 2023g), morsoravids (Mayr, 1999c), and zygodactylids (Mayr and Kitchener, 2023g), whereas this structure is more strongly developed in *Masillaraptor* (Mayr and Kitchener, 2021) and *Vastanavis* (Fig. 1o in Mayr et al., 2013).

The presence of a pneumatic foramen in the cranial end of the scapula is difficult to confirm from fossil material, but this foramen is absent in *Bathornis* (Mayr, 2016b) and apparently in the halcyornithid *Cyrilavis* (Ksepka et al., 2011). Contrary to the condition in most extant members of the clade, many fossil australavians have two pairs of sternal incisions, as seen in masillaraptorids (Mayr and Kitchener, 2021), messelasturids (Mayr, 2021c), halcyornithids (Mayr, 1998c; Mayr, 2000e; Mayr, 2007b; Ksepka et al., 2011; Mayr and Kitchener, 2024e), psittacopedids (Mayr and Daniels, 1998), morsoravids (Mayr, 1999c), and zygodactylids (Mayr, 1998b; Weidig, 2010; Smith et al., 2018; Hieronymus et al., 2019).

As has been noted by previous authors, the putative stem-cariamid *Elaphrocnemus* exhibits several features atypical of total-group cariamids (Mourer-Chauviré, 1983; Mayr, 2016b), such as a deltopectoral crest with an essentially straight proximodorsal margin. Together with the absence of a tubercle on the ventral surface of the minor metacarpal (which was optimized as a synapomorphy of Telluraves in the present study), these features may hint at a placement of this taxon outside total-group Cariamidae, or even Telluraves, with closer affinities to *Opisthocomus* suggested by Mayr (2016b).

#### Eufalconimorphae

No unambiguous synapomorphies were optimized for Eufalconimorphae in this study.

#### Psittacopasseres

Six character states were optimized as potential synapomorphies of Psittacopasseres, of which four were consistently inferred as such across all alternative molecular topologies studied.

- External spine on sternum 10–20% of total sternum length (Fig. 36, char. 15: 1 > 2, also found in *Ortalis* + *Rollulus*, *Phoenicopterus*, *Corythaeola*, *Tapera*, *Turnix*, *Oceanites*, and Eucavitaves); J14, P15, K21, S24, W24.
- External spine of sternum dorsally oriented (Fig. 36, char. 17: 1 > 0, also found in *Monias*, *Columba*, Charadrii, *Turnix*, *Alca*, *Gavia*, *Ninox*, *Urocolius*, *Trogon*, Coraciiformes, and Pici); J14, P15, K21, S24, W24.
- Impression for acrocoracohumeral ligament on coracoid deep (char. 64: 0 > 1, also found in *Eudromia*, *Chauna*, *Dendrocygna*, Mirandornithes, *Streptoprocne*, *Rallus*, *Rostratula*, *Turnix*, *Pagodroma*, *Leptoptilos*, Suloidea, *Eudocimus*, *Pelecanus*, Accipitrimorphae, Strigiformes, *Urocolius*, and Eucavitaves); J14, K21, S24.
- Scapulotricipital impression on ulna deep (char. 157: 0 > 1, also found in Mirandornithes, Columbimorphae, Strisores, *Rallus*, Charadriiformes, *Phaethon*, *Gavia*, Procellariiformes, Suliformes, *Eudocimus*, *Scopus*, *Pandion*, Strigiformes, *Trogon*, *Bucco*, and *Psilopogon*); J14, P15, K21, S24, W24.
- Minor metacarpal weakly bowed (char. 174: 1 > 0, reversed in *Menura*; also found in *Gansus*, *Ichthyornis*, *Dendrocygna*, Mirandornithes, Apodiformes, Charadriiformes, *Phaethon*, *Gavia*, Procellariimorphae, *Fregata*, *Eudocimus*, *Pandion*, and Picodynastornithes); J14, P15, K21, S24, W24.
- Small tubercle on minor metacarpal distal to proximal synostosis of metacarpals absent (char. 186: 1 > 0, also found in *Pandion*, *Urocolius*, Eucavitaves, and most non-telluravian neognaths); J14, P15, K21, W24.

**Figure 36.**
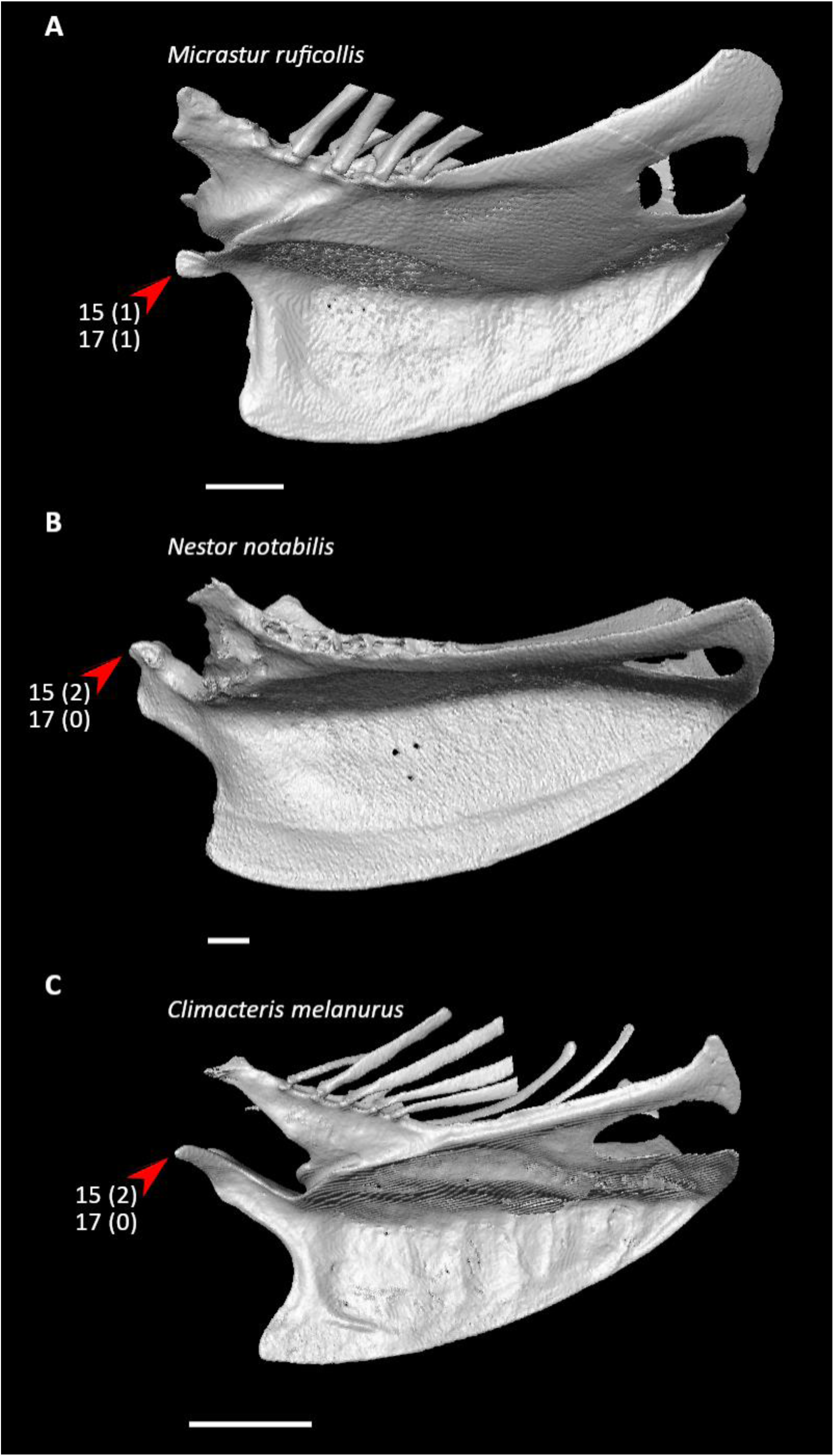
Sterna of *Micrastur ruficollis* (**A**, FMNH 330226), *Nestor notabilis* (**B**, FMNH 23530), and *Climacteris melanurus* (**C**, UMMZ 214303) in left (**A, C**) or right (**B**, mirrored) lateral view. Scale bars = 5 mm. Arrows indicate the external spine. *Micrastur* exhibits state 1 for character 15 (external spine 5–10% of total sternum length), whereas *Nestor* and *Climacteris* exhibit state 2 (external spine 10–20% of total sternum length, optimized as a synapomorphy of Psittacopasseres). *Micrastur* also exhibits state 1 for character 17 (external spine cranially oriented), whereas *Nestor* and *Climacteris* exhibit state 0 (external spine dorsally oriented, optimized as a synapomorphy of Psittacopasseres).

A marked impression for the acrocoracohumeral ligament is visible on the coracoid of *Vastanavis* (Fig. 1 in Mayr et al., 2007; Fig. 1b in Mayr et al., 2013), despite having been described as “shallow” in this taxon (Mayr et al., 2007), as well as that of the stem-psittaciform *Quercypsitta* (Pl. 2 in Mourer-Chauviré, 1992b). An elongated, dorsally oriented external spine of the sternum is found in halcyornithids (albeit not as dorsally inflected as in crown-group Psittaciformes, Mayr, 2007b; Fig. 3 in Mayr and Kitchener, 2024e), psittacopedids (Mayr and Daniels, 1998; Mayr, 2015c; Mayr, 2020), morsoravids (Mayr and Kitchener, 2023h), zygodactylids (Mayr, 1998b; Mayr, 2008b; Mayr and Kitchener, 2023g), and the stem-passeriform *Minutornis* (Mayr and Kitchener, 2023g), though its exact proportions relative to the rest of the sternum are often difficult to determine in fossils due to incomplete preservation. Messelasturids exhibit a shorter external spine (Mayr, 2021c; Mayr and Kitchener, 2023f), which may suggest a stemward phylogenetic placement compared to the aforementioned taxa, as found by Mayr (2021c) and Mayr and Kitchener (2023f). A deep scapuoltricipital impression on the ulna has been reported in *Messelastur* (Mayr, 2011d).

The presence of a weakly bowed minor metacarpal in messelasturids (Mayr, 2000d; Mayr, 2005c), halcyornithids (Mayr, 1998c; Ksepka et al., 2011), *Eofringillirostrum* (Ksepka et al., 2019), *Psittacopes* (Mayr and Daniels, 1998; Mayr and Kitchener, 2023g), morsoravids (Mayr, 1999c), zygodactylids (Fig. 26 in Mayr, 1998b; Mayr and Kitchener, 2023g), and *Minutornis* (Fig. 6o in Mayr and Kitchener, 2023g) is consistent with affinities of these taxa to total-group Psittacopasseres within Telluraves. However, a more strongly bowed minor metacarpal is present in *Vastanavis* (Mayr et al., 2013) and the psittacopedids *Psittacomimus* and *Parapsittacopes* (Mayr and Kitchener, 2023g), which may support a relatively stemward placement of *Vastanavis* in total-group Psittacopasseres, as found by Mayr and Kitchener (2023f).

Although not optimized as an unambiguous synapomorphy of Psittacopasseres in the present study, another pectoral character worthy of consideration as such is a bifurcated external spine on the sternum (char. 18: 1), given that this feature is thought to be plesiomorphic for both crown-group Psittaciformes (Mayr, 2010c) and Passeriformes (Mayr and Kitchener, 2023g). Among fossil pan-psittacopassereans, the external spine is weakly bifurcated in messelasturids (Mayr, 2021c), morsoravids (Mayr and Kitchener, 2023h), and the zygodactylid *Primoscens* (Mayr and Kitchener, 2023g), and strongly bifurcated in *Minutornis* (Mayr and Kitchener, 2023g), but not bifurcated in halcyornithids (Mayr, 2007b), psittacopedids (Mayr and Daniels, 1998; Mayr, 2020), and other zygodactylids (Mayr, 1998b; Mayr, 2008b; Smith et al., 2018). If the assignment of all these taxa to total-group Psittacopasseres is correct, this may indicate that a bifurcated external spine has been secondarily lost or convergently evolved multiple times within this lineage. It is also possible that this trait has deeper phylogenetic origins, as a slightly bifurcated external spine has been reported in the type specimen of *Danielsraptor*, though it is not present in a tentatively referred second specimen (Mayr and Kitchener, 2021).

### Character Support for Alternative Phylogenetic Topologies

#### Deep Divergences Within Neoaves

The phylogenetic relationships among Neoaves inferred from recent phylogenomic studies notably conflict with each other in several respects. For example, the extant sister group to all other neoavians has been variously recovered as Mirandornithes (the non-coding tree of Braun and Kimball, 2021; Kuhl et al., 2021; Stiller et al., 2024), a clade uniting Mirandornithes and Columbimorphae (Jarvis et al., 2014; Reddy et al., 2017; Houde et al., 2019; the coding exon tree of Braun and Kimball, 2021 following RY coding; the combined retroelement and sequenced-based gene tree of Gatesy and Springer, 2022), or Strisores (Prum et al., 2015; the retroelement tree of Gatesy and Springer, 2022).

Using the present dataset, eight osteological character states were optimized as potential synapomorphies of a clade containing all neoavians except for Mirandornithes, of which five were inferred as such under both the Kuhl et al. (2021) and Stiller et al. (2024) topologies.

- Ventrolateral sulcus on sternum distinct (char. 34: 0 > 1, reversed in *Caprimulgus*, *Aegotheles*, *Sterna*, Gruoidea, *Eurypyga*, *Phoebastria*, *Leptoptilos*, Pelecaniformes, *Bucorvus*, and *Nestor*; also found in *Rollulus*); K21, S24.
- Two pairs of caudal fenestrae in sternum (char. 49: 1 > 2, reversed in *Monias*, *Caprimulgus*, Gruiformes, *Turnix*, Aequornithes, *Pandion* + *Elanus*, *Tyto*, Bucerotiformes, and Australaves; also found in *Gansus* and *Ichthyornis*), S24.
- Procoracoid process of coracoid elongate (char. 68: 1 > 2, reversed in *Ardeotis*, *Caprimulgus*, *Nyctibius*, *Streptoprocne*, *Sarothrura*, *Rallus*, *Eurypyga*, *Gavia*, *Phoebastria*, *Leucocarbo*, Pelecaniformes, *Urocolius*, *Trogon*, Pici, Accitrimorphae, *Nestor*, and Passeriformes; also found in *Gansus*, *Ichthyornis*, and *Chauna*); K21, S24.
- Coracoid compressed and keeled at ventromedial margin of supracoracoid sulcus (char. 69: 0 > 1, reversed in *Pterocles*, *Ardeotis*, *Tapera*, *Aegotheles*, *Streptoprocne*, Gruiformes, Aequornithes, *Bucorvus*, *Nestor*, and *Acanthisitta*; also found in *Dendrocygna*, *Ortalis*, and *Rollulus*); K21, S24.
- Coracobrachial impression on humerus prominent (char. 114: 0 > 1, reversed in *Monias*, *Tapera*, Apodiformes, *Charadrius*, *Limosa*, *Alca*, *Sarothrura*, *Rallus*, *Gavia*, *Oceanites* + *Pagodroma*, Pelecanimorphae, *Urocolius*, Pici, *Pandion* + *Elanus*, *Nestor*, and *Acanthisitta*; also found in *Anseranas* and *Ortalis*); K21, S24.
- Humerus narrowest near midshaft point (char. 126: 0 > 2, reversed in *Monias*, *Ardeotis*, Gruiformes, *Limosa*, *Alca*, *Phaethon*, *Gavia*, *Phoebastria*, *Pagodroma*, Suliformes, *Eudocimus*, *Coragyps*, *Elanus*, Strigiformes, *Urocolius*, *Merops*, *Alcedo*, *Bucco*, *Micrastur*, and *Acanthisitta*; also found in *Ichthyornis*, *Alectura*, and *Rollulus*); S24.
- Radial incisure on ulna absent or indistinct (char. 156: 1 > 0, reversed in *Ardeotis*, *Nyctibius*, Grues, Charadriiformes, *Phaethon*, *Phoebastria*, *Pagodroma*, *Leptoptilos*, *Fregata*, *Sula*, *Scopus* + *Pelecanus*, Accipitrimorphae, and Psittaciformes; also found in *Dendrocygna* and Galliformes); K21, S24.
- Distal synostosis of major and minor metacarpals shorter than craniocaudal width (char. 198: 1 > 0, reversed in *Monias*, *Psophia*, Ralloidea, *Charadrius*, Scolopaci, *Sterna*, *Gavia*, *Eudocimus*, and *Cariama*; also found in *Gansus* and Galliformes); S24.

Two character states were optimized as potential synapomorphies of all neoavians excluding both Mirandornithes and Columbimorphae under the Jarvis et al. (2014) topology.

- Coracobrachial nerve sulcus on humerus prominent (char. 113: 0 > 1, reversed in *Tapera*, *Podargus*, *Psophia*, Lari, *Eurypyga*, *Spheniscus*, *Oceanites* + *Pagodroma*, Suloidea, *Eudocimus*, *Scopus*, *Pandion*, *Ninox*, *Leptosomus*, *Bucorvus*, *Merops*, *Alcecdo*, *Cariama*, *Nestor*, *Acanthisitta*, and Tyranni; also found *Anseranas*, *Alectura*, *Ortalis*, *Podilymbus*, and *Pterocles*).
- Brachial fossa on humerus deep (char. 134: 0 > 1, reversed in *Tapera*, *Sarothrura*, *Rallus*, *Rostratula*, *Turnix*, *Alca*, *Phoebastria*, *Oceanites*, *Tigrisoma*, *Elanus*, *Ninox*, *Upupa*, *Alcedo*, *Cariama*, *Psittacus*, and *Erythropitta*; also found in *Ichthyornis*, Anseriformes, and *Phoenicopterus*).

Two character states were optimized as potential synapomorphies for a clade containing all neoavians except Strisores under the Prum et al. (2015) topology.

- Procoracoid process of coracoid elongate (char. 68: 1 > 2, reversed in Mirandornithes, *Monias*, *Ardeotis*, *Sarothrura*, *Rallus*, *Eurypyga*, *Gavia*, *Phoebastria*, *Leucocarbo*, Pelecaniformes, *Urocolius*, *Trogon*, Pici, Accitrimorphae, *Nestor*, and Passeriformes; also found in *Gansus*, *Ichthyornis*, *Chauna*, and Letornithes).
- Radial tendinal sulcus well defined (char. 148: 0 > 1, reversed in *Corythaeola*, *Psophia*, *Leptoptilos*, *Eudocimus*, *Phoenicopterus*, Charadrii, *Turnix*, *Pandion*, *Merops*, *Bucco*, *Nestor*, and *Neopelma*; also found in *Ichthyornis*, *Chauna*, and Letornithes).

Contrasting with other phylogenomic studies, S. Wu et al. (2024) recovered the split at the root of Neoaves between Telluraves and a clade containing all other neoavians, referred to as Aquaterraves. However, no unambiguous synapomorphies were found for Aquaterraves using the present dataset.

It is perhaps noteworthy that a greater number of characters in our dataset potentially exclude Mirandornithes from other neoavians compared to those supporting competing topologies. However, three of these characters are also absent in *Caprimulgus*, a representative of Caprimulgidae, which is likely the extant sister group to the rest of Strisores (Prum et al., 2015; Chen et al., 2019; White and Braun, 2019; Kuhl et al., 2021; Stiller et al., 2024; S. Wu et al., 2024). As such, they do not offer unambiguous support against Strisores representing the extant sister group to other neoavians. Conversely, both of the characters potentially excluding Strisores from other neoavians are absent in at least some Mirandornithes and thus do not provide evidence against Mirandornithes as the extant sister group of other neoavians. The characters potentially excluding Mirandornithes and Columbimorphae from other neoavians are also found in multiple non-neoavian taxa and some mirandornitheans, raising the possibility that they are instead plesiomorphic for crown birds.

Two character states were optimized as potential synapomorphies for a clade uniting Mirandornithes and Columbimorphae to the exclusion of other neoavians under the Jarvis et al. (2014) topology.

- Humeral shaft straight (char. 125: 0 > 1, reversed in *Monias*; also found in Apodiformes, *Opisthocomus*, Charadriiformes, *Phaethon*, Procellariimorphae, *Fregata*, *Leucocarbo*, *Urocolius*, *Psilopogon*, *Psittacus*, *Neopelma*, and Passeri).
- Scapulotricipital impression on ulna deep (char. 157: 0 > 1, also found in Strisores, *Rallus*, Charadriiformes, *Phaethon*, *Gavia*, Procellariiformes, Suliformes, *Eudocimus*, *Scopus*, *Pandion*, Strigiformes, *Trogon*, *Bucco*, *Psilopogon*, and Psittacopasseres).

If Mirandornithes is not the extant sister group to other neoavians (either by itself or alongside Columbimorphae), one alternative hypothesis for its affinities is a position close to Charadriiformes (Prum et al., 2015), members of which also share both of the traits above. One of these character states was optimized as a potential synapomorphy for a clade uniting Mirandornithes and Charadriiformes to the exclusion of other neoavians under the Prum et al. (2015) topology.

- Humeral shaft straight (char. 125: 0 > 1, reversed in *Burhinus*; also found in *Pterocles*, *Columba*, Apodiformes, *Opisthocomus*, *Phaethon*, Procellariimorphae, *Fregata*, *Leucocarbo*, *Urocolius*, *Psilopogon*, *Psittacus*, *Neopelma*, and Passeri).

In the results of S. Wu et al. (2024), Mirandornithes is recovered as the sister group to a clade uniting *Opisthocomus* and Phaethoquornithes. Three character states were optimized as potential synapomorphies supporting this relationship.

- Keel low, less than two-thirds total height of sternum (char. 47: 1 > 0, reversed in *Eudocimus*; also found in *Chauna*, *Anseranas*, *Tapera*, Vanescaves, *Psophia*, Charadrii, *Limosa*, and Telluraves).
- Olecranon fossa on humerus shallow (char. 127: 1 > 0, reversed in Phaethontimorphae, Procellariimorphae, Suliformes, and *Scopus* + *Pelecanus*; also found in non-neognath birds, *Anseranas*, *Monias*, *Ardeotis*, *Nyctibius*, Grues, and *Alca*).
- Extensor process on carpometacarpus surpasses distal articular facet for alular digit by half the width of the facet or less (char. 182: 3 > 2, reversed in *Eurypyga*, Procellariiformes, and Pelecanimorphae; also found in non-neognath birds, *Monias*, *Corythaeola*, *Charadrius*, Bucerotiformes, *Nestor*, and *Climacteris*).

All three of these character states are also found in non-neognath birds, raising the possibility that they are plesiomorphic for Neoaves. Character support from the present dataset therefore does not strongly favor any of the aforementioned alternatives for the affinities of Mirandornithes. However, reanalyses of the Prum et al. (2015) dataset by Braun and Kimball (2021) suggest that the close relationship between Mirandornithes and Charadriiformes may be an artifact driven by data-type effects. Furthermore, Mirarab et al. (2024) presented evidence that the proposed sister-group relationship between Mirandornithes and Columbimorphae is likewise a probable analytical artifact caused by a period of sequence polymorphism early in the evolutionary history of Neoaves. Our identification of multiple potential synapomorphies excluding Mirandornithes from a clade containing all or most other neoavians may thus add to a growing consensus for a sister-group relationship between Mirandornithes and the remainder of Neoaves as the most likely divergence at the root of crown-group Neoaves (Jarvis et al., 2014; Reddy et al., 2017; Braun and Kimball, 2021; Kuhl et al., 2021; Gatesy and Springer, 2022; Stiller et al., 2024).

Both Jarvis et al. (2014) and Prum et al. (2015) recovered a clade within Neoaves that consists of *Opisthocomus*, Gruiformes, Charadriiformes, Phaethoquornithes, and Telluraves while excluding Columbimorphae, Otidimorphae, and Strisores. Under the Prum et al. (2015) topology, this clade also includes Mirandornithes (see previous paragraph for discussion on the phylogenetic position of Mirandornithes). Four character states were optimized as potential synapomorphies of this clade using the present dataset, though only one was inferred as such under both topologies.

- Costal margin 25–75% of total sternum length (char. 35: 0 > 1, reversed in Mirandornithes [if considered members of this clade], *Sarothrura*, *Burhinus*, *Rostratula*, *Turnix*, *Tigrisoma*, *Ninox*, *Urocolius*, Eucavitaves, and Eupasseres; also found in *Ichthyornis*, Anseriformes, *Corythaeola*, and *Streptoprocne*); J14, P15.
- Five or more articular facets on costal margin of sternum (char. 36: 1 > 2, reversed in *Tigrisoma*, *Turnix*, Strigiformes, *Urocolius*, Eucavitaves, and Tyranni; also found in *Gansus*, *Ichthyornis*, Anseriformes, *Podargus*, Apodiformes, *Corythaeola*, and *Monias*); P15.
- Dorsal pneumotricipital fossa narrower than ventral pneumotricipital fossa (char. 112: 1 > 0, reversed in *Sterna*, *Trogon*, *Alcedo*, and *Acanthisitta*; also found in *Rollulus*); P15.
- Ventral epicondyle on humerus equal or distal to ventral condyle (char. 130: 0 > 1, reversed in *Balearica*, *Podica*, *Rallus*, *Burhinus*, *Turnix*, *Sterna*, *Phaethon*, *Oceanites*, *Leptoptilos*, Suloidea, *Eudocimus*, *Coragyps*, *Pandion*, *Tyto*, and *Nestor*; also found in Galliformes, *Phoenicopterus*, Pteroclimesites, *Tapera*, and *Streptoprocne*); J14.

Characters 35 (1) and 130 (1) were also optimized as potential synapomorphies for a similar neoavian clade recovered by Stiller et al. (2024), which excludes Mirandornithes, Columbimorphae, and Otidimorphae, but not Strisores. Under this topology, these character states experienced reversals in Strisores.

In contrast, Kuhl et al. (2021) recovered a clade consisting of Gruiformes, Charadriiformes, Otidimorphae, Columbimorphae, *Opisthocomus*, and Strisores. However, no unambiguous synapomorphies were optimized for this clade using the present dataset.

*Opisthocomus*, Gruiformes, Charadriiformes, Phaethoquornithes, and Strisores form an exclusive clade called Elementaves in the results of Stiller et al. (2024). One character state was optimized as a potential synapomorphy for this clade.

- Pronounced ventral curvature of scapula (char. 92: 0 > 1, reversed in *Podargus*, *Aegotheles*, *Streptoprocne*, *Balearica*, Charadriiformes, *Phaethon*, *Gavia*, *Phoebastria*, and Pelecanes; also found in *Coragyps*, *Pandion*, *Tyto*, *Merops*, *Alcedo*, *Psilopogon*, *Nestor*, *Acanthisitta*, and Tyranni).

Within Elementaves, Stiller et al. (2024) recovered Phaethoquornithes and Strisores as extant sister taxa. Four character states were optimized as potential synapomorphies for this clade.

- Hypocleideum less than 5% the total length of the furcula (char. 9: 1 > 0, reversed in *Streptoprocne* and *Oceanites*; also found in *Podilymbus*, Scolopaci, *Alca*, *Coragyps*, *Bucorvus*, *Alcedo*, *Cariama*, and *Menura*).
- Internal lip of coracoid expanded in medial portion (char. 87: 0 > 1, reversed in *Gavia*, *Pagodroma*, and Pelecanes; also found in *Phoenicopterus*, *Corythaeola*, *Ardeotis*, *Tapera*, *Balearica*, *Elanus*, *Leptosomus*, *Bucorvus*, and *Cariama*).
- Transverse sulcus on humerus intermediate in length (char. 115: 0 > 1, reversed in *Podargus*, *Eurypyga*, *Spheniscus*, and *Phoebastria*; also found in *Anseranas*, *Balearica*, *Charadrius*, *Coragyps*, *Elanus*, *Tyto*, *Coracias*, and Psittaciformes).
- Scapulotricipital impression on ulna deep (char. 157: 0 > 1, reversed in Daedalornithes, *Spheniscus*, *Leptoptilos*, *Pelecanus*, and *Tigrisoma*; also found in Mirandornithes, Columbimorphae, *Rallus*, Charadriiformes, *Pandion*, Strigiformes, *Trogon*, *Bucco*, *Psilopogon*, and Psittacopasseres).

A sister-group relationship between Gruiformes and Charadriiformes, forming a clade called Cursorimorphae, was found by Jarvis et al. (2014), Kuhl et al. (2021), Stiller et al. (2024), and S. Wu et al. (2024). No character states in the present dataset were optimized as unambiguous synapomorphies of this clade under the Jarvis et al. (2014), Kuhl et al. (2021), or Stiller et al. (2024) topologies, though a single potential synapomorphy was recovered under the S. Wu et al. (2024) topology.

- Minor metacarpal prominently narrows distally in caudal view (char. 180: 0 > 1, reversed in *Psophia*; also found in *Ichthyornis*, Anseres, Mirandornithes, Columbimorphae, *Aegotheles*, Phaethontimorphae, Pelecanimorphae, *Leptosomus*, *Coracias*, Accipitrimorphae, Strigiformes, and Australaves).

A close relationship between members of Gruiformes and Charadriiformes has long been hypothesized based on anatomical data, especially cranial characteristics such as the presence of occipital fontanelles, supraorbital salt glands, and schizorhinal nostrils (e.g., Beddard, 1898; Olson, 1979; Olson, 1985; Livezey and Zusi, 2007), but support for this hypothesis from the pectoral girdle and forelimb skeleton seems to be limited.

Cursorimorphae was recovered as the extant sister group to Strisores by S. Wu et al. (2024), forming a clade called Litusilvanae. One character state was optimized as a potential synapomorphy for this clade.

- Dorsoventral orientation of acrocoracoid process essentially coplanar with main craniocaudal axis of coracoid (char. 62: 0 > 1, reversed in Grues, *Rallus*, *Limosa*, and *Alca*; also found in Galliformes, *Pterocles*, *Columba*, *Urocolius*, Eucavitaves, Tyranni, and *Climacteris*).

An alternative phylogenetic position for Charadriiformes close to Mirandornithes and Phaethoquornithes as part of an “extended waterbird clade” (i.e. Aequorlitornithes) was recovered by Prum et al. (2015). Four character states were optimized as potential synapomorphies of Aequorlitornithes using the present dataset.

- Pointed acromial processes on furcula (char. 5: 0 > 1, reversed in *Podilymbus*, *Phaethon*, and Pelecanimorphae; also found in *Ichthyornis*, Anseriformes, *Nyctibius*, Apodiformes, *Pterocles*, *Sarothrura*, *Coragyps*, *Pandion*, *Urocolius*, and *Acanthisitta*).
- Mound-shaped tuberosity on lateral surface of scapula (char. 94: 0 > 1, reversed in *Podilymbus*, *Turnix*, *Sterna*, *Phaethon*, Procellariimorphae, *Scopus*, and *Tigrisoma*; also found in *Ardeotis* and Bucerotiformes).
- Scapulotricipital impression on ulna deep (char. 157: 0 > 1, reversed in *Charadrius*, *Eurypyga*, *Spheniscus*, *Leptoptilos*, *Pelecanus*, and *Tigrisoma*; also found in Columbimorphae, Strisores, *Rallus*, *Pandion*, Strigiformes, *Trogon*, *Bucco*, *Psilopogon*, and Psittacopasseres).
- Minor metacarpal weakly bowed (char. 174: 1 > 0, reversed in *Turnix*, *Eurypyga*, *Oceanites*, *Leptoptilos*, Suloidea, and *Pelecanus* + *Tigrisoma*; also found in *Ichthyornis*, *Dendrocygna*, Apodiformes, Ralloidea, *Pandion*, Picocoraciades, and Psittacopasseres).

It is perhaps noteworthy that half of these character states (chars. 5: 1 and 174: 0) are also found in *Ichthyornis* and anseriforms, raising the possibility that adaptation towards an aquatic lifestyle readily drives their convergent evolutionary acquisition. This would also be consistent with the findings of Braun and Kimball (2021), which suggest that the recovery of Aequorlitornithes is an artifact of data-type effects (see also previous discussion on the phylogenetic position of Mirandornithes).

A single character state was optimized as a potential synapomorphy of a group containing members of Aequorlitornithes, *Opsithocomus*, and Telluraves to the exclusion of other neoavians, another clade recovered by Prum et al. (2015).

- Keel low, less than two-thirds total height of sternum (char. 47: 1 > 0, reversed in *Rostratula*, Lari, *Eudocimus*, *Upupa*, Psittaciformes, and *Menura*; also found in *Chauna*, *Anseranas*, *Tapera*, Vanescaves, and *Psophia*).

Jarvis et al. (2014) and Kuhl et al. (2021), on the other hand, recovered a sister-group relationship between Phaethoquornithes and Telluraves, but no unambiguous synapomorphies were optimized for this clade using the present dataset.

#### Affinities of Musophagiformes, Otidiformes, and Cuculiformes

Another point of contention regarding neoavian phylogeny pertains to the affinities of turacos (Musophagiformes), bustards (Otidiformes), and cuckoos (Cuculiformes). In the results of Jarvis et al. (2014), Prum et al. (2015), Stiller et al. (2024), S. Wu et al. (2024), and the combined retroelement and sequence-based gene tree of Gatesy and Springer (2022), these three groups form a clade to the exclusion of other neoavians, known as Otidimorphae.

However, in some other analyses, either Cuculiformes (Kuhl et al., 2021) or Musophagiformes (Reddy et al., 2017; the non-coding tree of Braun and Kimball, 2021) has been found to be only distantly related to the other two lineages. Three character states were optimized as potential synapomorphies of a clade uniting all three groups using the present dataset, though only one was inferred as such under all four molecular reference topologies supporting the monophyly of Otidimorphae.

- Internal lip of coracoid expanded in medial portion (char. 87: 0 > 1, also found in *Phoenicopterus*, Strisores, *Balearica*, Phaethontimorphae, *Spheniscus*, *Phoebastria*, *Leptoptilos*, *Leucocarbo*, *Scopus* + *Pelecanus*, *Elanus*, *Leptosomus*, *Bucorvus*, and *Cariama*); S24.
- Bicipital crest on humerus inconspicuous (char. 117: 1 > 0, also found in *Chauna*, *Anseranas*, *Alectura*, *Opisthocomus*, *Balearica*, *Eurypyga*, *Leptoptilos*, *Fregata*, *Scopus*, *Tigrisoma*, *Tyto*, *Leptosomus*, *Bucorvus*, *Cariama*, and *Micrastur*); J14, P15, S24, W24.
- Ventral ramus of pisiform subequal in cranial extent to dorsal ramus or longer (char. 166: 0 > 1, also found in *Anseranas*, Galliformes, Mirandornithes, *Opisthocomus*, *Rostratula*, *Gavia*, *Oceanites* + *Pagodroma*, *Leptoptilos*, *Eudocimus*, *Ninox*, *Urocolius*, *Upupa*, Picocoraciades, and Australaves); P15.

In general, these characters may be suggestive of close affinities among Musophagiformes, Otidiformes, and Cuculiformes, and a close relationship between at least Musophagiformes and Cuculiformes has long been hypothesized based on comparative anatomy (e.g., Pycraft, 1903; Mayr and Clarke, 2003; Livezey and Zusi, 2007; Mayr, 2011a). However, all of these character states are also found in clades that have been recovered as potential close relatives of these three groups by recent phylogenomic studies, including members of Gruiformes and Aequornithes (Reddy et al., 2017; the non-coding tree of Braun and Kimball, 2021). Consequently, the monophyly of a group uniting Musophagiformes, Otidiformes, and Cuculiformes remains to be further tested by future studies.

Possible comparisons to fossil forms are limited, given that the fossil record of Musophagiformes, Otidiformes, and Cuculiformes is scant (Mayr, 2017b; Mayr, 2022a), though the Eocene *Foro* is known from a nearly complete skeleton and has been identified as a potential stem-musophagid (Field and Hsiang, 2018), and similarities to otidids have been noted in the enigmatic Eocene *Perplexicervix* (Mayr, 2010d; Mayr et al., 2023a). A small bicipital crest is found in *Perplexicervix* (Mayr et al., 2023a), whereas *Foro* has been noted to have a more prominent bicipital crest on the humerus than in extant musophagiforms and cuculiforms, approaching the condition in accipitriforms (Olson, 1992). If *Foro* is a stem-musophagid, and musophagiforms indeed form an exclusive clade with cuculiforms and/or otidiforms, it may represent an apomorphic reversal of a reduced bicipital crest within this clade.

An alternative topology in which Cuculiformes were recovered as the extant sister group of pigeons (Columbiformes) within Columbimorphae was found by Kuhl et al. (2021). Four character states were optimized as potential synapomorphies of this clade using the present dataset.

- Ventral supracondylar tubercle on humerus expanded (char. 141: 0 > 1, also found in *Ichthyornis*, *Chauna*, Galliformes, *Limosa*, *Turnix*, *Gavia*, *Oceanites*, *Fregata*, *Leucocarbo*, Coraciimorphae, and *Climacteris*).
- Carpal tubercle on ulna projects more than half the craniocaudal width of dorsal and ventral condyles (char. 163: 0 > 1, also found in *Ortalis*, *Caprimulgus*, *Florisuga*, *Opisthocomus*, *Psophia*, Charadrii, Scolopaci, *Sterna*, *Phaethon*, *Gavia*, Procellariiformes, *Fregata*, *Urocolius*, *Trogon*, *Upupa*, *Coracias*, *Psilopogon*, and Passeriformes).
- Prominent ligamental groove on ventral ramus of pisiform (char. 168: 0 > 1, also found in *Ichthyornis*, *Dendrocygna*, *Nyctibius*, *Streptoprocne*, Grues, Ralloidea, *Charadrius*, Scolopaci, *Sterna*, Phaethoquornithes, *Pandion*, Strigiformes, *Upupa*, *Coracias*, *Alcedo*, *Psittacus*, *Erythropitta*, and Passeri).
- Ridge linking ventral rim of carpal trochlea and pisiform process forms sharp drop-off relative to ventral surface of extensor process (char. 192: 0 > 1, also found in *Anseranas*, Galliformes, *Florisuga*, *Opisthocomus*, Grues, Heliornithes, *Turnix*, *Eurypyga*, *Leucocarbo*, *Eudocimus*, *Tigrisoma*, *Coragyps*, Strigiformes, *Urocolius*, *Psittacus*, and Tyranni).

Two additional character states present in *Tapera* were optimized as synapomorphies of Columbimorphae under this topology (see previous discussion on potential synapomorphies of Columbimorphae).

Even if Musophagiformes, Otidiformes, and Cuculiformes represent each other’s closest living relatives, recent studies have found conflicting results regarding the internal topology of this clade. In Jarvis et al. (2014), Musophagiformes and Otidiformes were found forming a clade to the exclusion of Cuculiformes. A sister-group relationship between these two clades was also recovered by Kuhl et al. (2021), Luo et al. (2023), and S. Wu et al. (2024), and this grouping was named Musophagotides by Sangster et al. (2022). Seven character states were optimized as potential synapomorphies of Musophagotides using the present dataset, four of which were inferred as such all three molecular reference topologies supporting this group.

- Sternal keel maintains thickness ventrally (char. 46: 0 > 1, also found in *Podilymbus*, *Monias*, *Opisthocomus*, *Psophia*, *Balearica*, *Phaethon*, *Gavia*, *Spheniscus*, *Pagodroma*, *Fregata*, *Leucocarbo*, *Pelecanus*, *Pandion* + *Elanus*, *Leptosomus*, *Alcedo*, *Jynx*, and *Menura*); J14, K21, W24.
- Caudolateral trabeculae on sternum turn medially (char. 53: 1 > 0, also found in *Gansus*, *Eudromia*, *Ortalis*, *Podilymbus*, Columbimorphae, *Charadrius*, Lari, *Gavia*, *Spheniscus*, *Pelecanus*, *Coragyps*, *Bucorvus*, and *Bucco*); J14.
- Line linking acrocoracoid process and medial angle of coracoid forms angle with lateral and medial extremes of sternal facet that is markedly greater than 90–100° (char. 73: 1 > 0, also found in *Anseranas*, *Sula*, *Pelecanus*, *Bucorvus*, and *Psittacus*); J14, K21, W24.
- Bicipital crest on humerus inconspicuous (char. 117: 1 > 0, also found in *Chauna*, *Anseranas*, *Alectura*, *Tapera*, *Opisthocomus*, *Balearica*, *Eurypyga*, *Leptoptilos*, *Fregata*, *Scopus*, *Tigrisoma*, *Tyto*, *Leptosomus*, *Bucorvus*, *Cariama*, and *Micrastur*); K21.
- Deltopectoral crest well developed, extending at least one third the total length of the humerus (char. 122: 1 > 0, also found in *Gansus*, *Ichthyornis*, *Nyctibius*, *Podargus*, *Spheniscus*, *Leptoptilos*, *Leucocarbo*, *Eudocimus*, *Scopus*, Accipitrimorphae, Strigiformes, Cavitaves, *Micrastur*, and Passeri); J14, K21, W24.
- Humeroulnar trochlea on ulna absent or shallow (char. 154: 1 > 0, also found in *Ichthyornis*, Galloanserae, Columbimorphae, *Podargus*, *Aegotheles*, *Opisthocomus*, *Podica*, *Charadrius*, *Limosa*, *Turnix*, *Phaethon*, *Spheniscus*, *Leptoptilos*, *Fregata*, *Tigrisoma*, *Pandion*, *Leptosomus*, Bucerotiformes, *Coracias*, and *Alcedo*); J14.
- Impression of m. brachialis on ulna deep (char. 155: 0 > 1, also found in *Gansus*, *Ichthyornis*, *Eudromia*, *Chauna*, *Dendrocygna*, *Ortalis*, *Phoenicopterus*, *Aegotheles*, *Streptoprocne*, *Balearica*, *Podica*, *Charadrius*, *Limosa*, *Sterna*, *Phaethon*, *Phoebastria*, Suloidea, *Pelecanus*, Accipitrimorphae, Strigiformes, Cavitaves, *Psittacus*, *Acanthisitta*, and Passeri); J14, K21, W24.

Most of these characters cannot be assessed from published descriptions of *Foro*, though as noted previously it differs from extant musophagids in having a more prominent bicipital crest on the humerus. However, *Foro* does exhibit a well-developed deltopectoral crest (Olson, 1992). An elongated deltopectoral crest also appears to be present in *Perplexicervix* (Mayr, 2010d), but the sternal and medial margins of coracoids tentatively referred to this genus do not form a wide angle (Fig. 8 in Mayr et al., 2023a).

Prum et al. (2015) and Stiller et al. (2024) instead recovered a clade uniting Otidiformes and Cuculiformes to the exclusion of Musophagiformes, for which five character states were optimized as potential synapomorphies using the present dataset under both molecular topologies.

- Pneumatic foramen in median sulcus of sternum immediately caudal to cranial margin (char. 29: 0 > 1, also found in *Ichthyornis*, *Dendrocygna*, *Alectura*, *Ortalis*, *Phoenicopterus*, *Pterocles*, Strisores, *Opisthocomus*, Grues, *Podica*, *Phaethon*, Pelecanimorphae, *Coragyps*, Strigiformes, Picocoraciades, *Micrastur*, Psittaciformes, *Neopelma*, and *Menura*); P15, S24.
- Keel apex pointed in lateral view (char. 43: 0 > 1, also found in *Alectura*, *Podilymbus*, *Podargus*, *Aegotheles*, *Balearica*, Ralloidea, *Charadrius*, *Rostratula*, *Turnix*, *Alca*, *Leptoptilos*, Pelecaniformes, *Leptosomus*, *Bucorvus*, *Coracias*, *Alcedo*, *Acanthisitta*, and *Erythropitta*); P15, S24.
- Coracoid rounded and relatively thick at ventromedial margin of supracoracoid sulcus (char. 69: 1 > 0, also found in non-neoavian birds, Mirandornithes, *Pterocles*, *Aegotheles*, *Streptoprocne*, Gruiformes, Aequornithes, *Bucorvus*, *Nestor*, and *Acanthisitta*); P15, S24.
- Lateral process of coracoid sharply hooked (char. 80: 0 > 1, also found in *Gansus*, *Ichthyornis*, *Anseranas*, *Rollulus*, Strisores, Grues, *Burhinus*, *Sterna*, *Phaethon*, Procellariiformes, *Leptoptilos*, *Fregata*, *Leucocarbo*, *Scopus* + *Pelecanus*, *Pandion*, *Trogon*, *Bucorvus*, *Coracias*, *Bucco*, and *Psittacus*); P15, S24.
- Radial bicipital tubercle prominent (char. 146: 0 > 1, also found in *Aegotheles*, *Florisuga*, Grues, *Charadrius*, *Phoebastria*, *Leucocarbo*, *Scopus*, *Tigrisoma*, *Elanus*, *Tyto*, *Leptosomus*, Bucerotiformes, *Merops*, Piciformes, and Passeriformes); P15, S24.

In the absence of increased taxon sampling and more completely known fossil representatives of Musophagiformes, Otidiformes, and Cuculiformes to further clarify the polarity and distribution of these potential synapomorphies, skeletal characters from the pectoral girdle and forelimb do not seem to clearly favor any of the recent alternative hypotheses regarding the interrelationships of these clades.

With respect to the phylogenetic relationships of Musophagiformes, Otidiformes, and Cuculiformes to other neoavians, Jarvis et al. (2014) recovered Otidimorphae as the extant sister group to Strisores. Four character states were optimized as potential synapomorphies uniting Otidimorphae and Strisores to the exclusion of other neoavians using the present dataset.

- Pneumatic foramen in median sulcus of sternum immediately caudal to cranial margin (char. 29: 0 > 1, reversed in *Corythaeola* and *Nyctibius*; also found in *Ichthyornis*, *Dendrocygna*, *Alectura*, *Ortalis*, *Phoenicopterus*, *Pterocles*, *Opisthocomus*, Grues, *Podica*, *Phaethon*, Pelecanimorphae, *Coragyps*, Strigiformes, Picocoraciades, *Micrastur*, Psittaciformes, *Neopelma*, and *Menura*).
- Lateral process of coracoid strongly hooked (char. 80: 0 > 1, reversed in *Corythaeola* and Daedalornithes; also found in *Gansus*, *Ichthyornis*, *Anseranas*, *Rollulus*, Grues, *Burhinus*, *Sterna*, *Phaethon*, Procellariiformes, *Leptoptilos*, *Fregata*, *Leucocarbo*, *Scopus* + *Pelecanus*, *Pandion*, *Trogon*, *Bucorvus*, *Coracias*, *Bucco*, and *Psittacus*).
- Internal lip of coracoid expanded in medial portion (char. 87: 0 > 1, also found in *Phoenicopterus*, *Balearica*, Phaethontimorphae, *Spheniscus*, *Phoebastria*, *Leptoptilos*, *Leucocarbo*, *Scopus* + *Pelecanus*, *Elanus*, *Leptosomus*, *Bucorvus*, and *Cariama*).
- Ventral collateral ligamental tubercle on ulna strongly developed (char. 158: 0 > 1, reversed in *Ardeotis*; also found in *Ichthyornis*, *Ortalis*, *Psophia*, Charadriiformes, Procellariiformes, *Fregata*, *Pelecanus*, *Elanus*, Strigiformes, Coraciimorphae, *Nestor*, and Passeriformes).

In contrast, Prum et al. (2015), Kuhl et al. (2021), Stiller et al. (2024), and S. Wu et al. (2024) recovered Otidimorphae as most closely related to Columbimorphae (with Cuculiformes being nested within the latter group in the results of Kuhl et al., 2021), forming a clade called Columbaves. Five character states were optimized as potential synapomorphies of Columbaves, one of which was inferred as such all four molecular reference topologies supporting this group.

- Pneumatic pores along cranial margin of dorsal sternum (char. 32: 0 > 1, reversed in *Pterocles* and *Corythaeola*; also found in Anseriformes, *Ortalis*, *Phoenicopterus*, *Tapera*, *Caprimulgus*, Apodiformes, *Opisthocomus*, Gruoidea, *Podica*, *Phaethon*, *Leptoptilos*, *Sula*, *Eudocimus*, *Scopus*, *Pelecanus*, *Pandion*, Strigiformes, *Trogon*, *Coracias*, *Merops*, and Australaves); W24.
- Caudolateral trabeculae on sternum turn medially (char. 53: 1 > 0, reversed in *Tapera*; also found in *Gansus*, *Eudromia*, *Ortalis*, *Podilymbus*, *Charadrius*, Lari, *Gavia*, *Spheniscus*, *Pelecanus*, *Coragyps*, *Bucorvus*, and *Bucco*); P15, K21.
- Lateral extent of lateral process on coracoid half the width of sternal facet or greater (char. 79: 0 > 1, reversed in *Ardeotis*; also found in *Caprimulgus*, *Podargus*, *Phoebastria*, *Pagodroma*, *Fregata*, *Sula*, *Scopus* + *Pelecanus*, *Charadrius*, Scolopaci + Lari, *Elanus*, *Tyto*, *Leptosomus*, *Upupa*, *Coracias* + *Merops*, and *Bucco*); P15, S24, W24.
- Quill knobs on ulna strongly developed (char. 161: 0 > 1, reversed in *Pterocles*; also found in *Anseranas*, *Podilymbus*, *Caprimulgus*, *Aramus*, *Charadrius*, Scolopaci + Lari, *Oceanites*, Pelecanimorphae, Accipitrimorphae, Picocoraciades, and Passeriformes); P15, K21, S24, W24.
- Cranial projection on distal end of major metacarpal weak (char. 197: 1 > 0, reversed in *Ardeotis*; also found in *Gansus*, *Ortalis* + *Rollulus*, *Opisthocomus*, *Psophia*, *Aramus*, Heliornithes, *Rostratula*, *Turnix*, *Spheniscus*, *Tigrisoma*, Strigiformes, *Upupa*, *Merops*, *Bucco*, *Cariama*, and Passeriformes); J14, S24.

Three of the above character states can also be found in *Caprimulgus*, and therefore do not necessarily contradict the possibility of an Otidimorphae + Strisores clade. One of these characters, the presence of prominent quill knobs, is lacking in *Foro* (Olson, 1992).

#### Interrelationships Within Telluraves

Within Telluraves, Jarvis et al. (2014), Kuhl et al. (2021), and Stiller et al. (2024) recovered a clade containing Accipitrimorphae, Strigiformes, and Coraciimorphae, forming a group called Afroaves. Seven character states were optimized as potential synapomorphies of Afroaves, of which two were inferred as such under all three topologies.

- Keel apex on sternum caudal to main body of sternum (char. 41: 1 > 0, reversed in Cavitaves; also found in *Eudromia*, Galliformes, *Monias*, *Opisthocomus*, *Aramus*, *Sarothrura*, *Rallus*, *Charadrius*, Scolopaci, *Turnix*, *Pagodroma*, *Fregata*, *Micrastur*, and *Acanthisitta*); J14, K21.
- Acrocoracoid process on coracoid straight (char. 63: 1 > 0, reversed in Cavitaves; also found in *Gansus*, Anseriformes, *Ortalis*, *Phoenicopterus*, *Monias*, *Ardeotis*, Strisores, *Opisthocomus*, Gruiformes, *Gavia*, *Pagodroma*, Pelecanimorphae, *Micrastur*, and *Acanthisitta*); K21.
- Impression for acrocoracohumeral ligament on coracoid deep (char. 64: 0 > 1, reversed in *Leptosomus*, *Bucorvus*, and *Bucco*; also found in *Eudromia*, *Chauna*, *Dendrocygna*, Mirandornithes, *Streptoprocne*, *Rallus*, *Rostratula*, *Turnix*, *Pagodroma*, *Leptoptilos*, Suloidea, *Eudocimus*, *Pelecanus*, and Psittacopasseres); J14, K21, S24.
- Deltopectoral crest well developed, extending at least one third the total length of the humerus (char. 122: 1 > 0, reversed in *Urocolius* and Coraciiformes; also found in *Gansus*, *Ichthyornis*, *Corythaeola*, *Ardeotis*, *Nyctibius*, *Podargus*, *Spheniscus*, *Leptoptilos*, *Leucocarbo*, *Eudocimus*, *Scopus*, *Micrastur*, and Passeri); J14, K21.
- Scar of m. flexor carpi ulnaris on flexor process of humerus forms single large scar (char. 142: 2 > 1, reversed in *Pandion*, *Leptosomus*, and *Psilopogon*; also found in *Eudromia*, Galliformes, *Tapera*, *Streptoprocne*, *Opisthocomus*, *Psophia*, *Aramus*, *Rallus*, *Eurypyga*, *Acanthisitta*, and *Erythropitta*); J14, K21, S24.
- Radius craniocaudally curved (char. 144: 0 > 1, reversed in *Elanus*, *Urocolius*, and *Coracias*; also found in Anseriformes, *Phoenicopterus*, *Ardeotis*, Strisores, Grues, Charadrii, Feraequornithes, *Micrastur*, Tyranni, and *Climacteris*); J14, K21.
- Impression of m. brachialis on ulna deep (char. 155: 0 > 1, reversed in *Urocolius*, *Alcedo*, and Pici; also found in *Gansus*, *Ichthyornis*, *Eudromia*, *Chauna*, *Dendrocygna*, *Ortalis*, *Phoenicopterus*, *Corythaeola*, *Ardeotis*, *Aegotheles*, *Streptoprocne*, *Balearica*, *Podica*, *Charadrius*, *Limosa*, *Sterna*, *Phaethon*, *Phoebastria*, Suloidea, *Pelecanus*, *Psittacus*, *Acanthisitta*, and Passeri); J14, K21.

Prum et al. (2015) instead found a Strigiformes + Coraciimorphae clade as sister to Australaves, forming a group called Eutelluraves. Two character states were optimized as potential synapomorphies of Eutelluraves.

- Humeral articular facet on coracoid flat or convex (char. 67: 0 > 1, reversed in Bucerotiformes, Psittaciformes, *Acanthisitta*, and *Neopelma*; also found in *Ortalis* + *Rollulus*, *Phoenicopterus*, *Monias*, *Columba*, *Corythaeola*, *Ardeotis*, Strisores, *Psophia*, *Rostratula*, Lari, *Eurypyga*, *Pagodroma*, and *Elanus*).
- Prominent caudal groove on caudal surface of minor metacarpal distal to synostosis with major metacarpal (char. 181: 0 > 2, reversed in *Bucorvus*, *Bucco*, and Psittaciformes; also found in *Ortalis* + *Rollulus*, *Tapera*, and *Podica*).

The Strigiformes + Coraciimorphae clade was recovered by Jarvis et al. (2014), Prum et al. (2015), and Kuhl et al. (2021). Two character states were optimized as potential synapomorphies of this clade, both of which were inferred as such under all three molecular topologies.

- Four facets for sternal ribs on costal margin of sternum (char. 36: 2 > 1, reversed in *Leptosomus*, *Upupa*, *Coracias*, and Pici; also found in *Eudromia*, *Alectura*, *Ortalis*, *Pterocles*, *Columba*, *Ardeotis*, *Caprimulgus*, *Nyctibius*, *Aegotheles*, and Tyranni); J14, P15, K21.
- Ventral collateral ligamental tubercle on ulna strongly developed (char. 158: 0 > 1, reversed in *Bucorvus* and *Merops*; also found in *Ichthyornis*, *Ortalis*, *Corythaeola*, *Tapera*, Strisores, *Psophia*, Charadriiformes, Procellariiformes, *Fregata*, *Pelecanus*, *Elanus*, *Nestor*, and Passeriformes); J14, P15, K21.

These characters are difficult to evaluate in potential stem-strigiform specimens (Mourer-Chauviré, 1987; Peters, 1992; Mayr et al., 2020c), though *Ypresiglaux* exhibits five costal processes on the sternum instead of four (Mayr and Kitchener, 2023d), which may suggest that a reduction to four occurred independently in strigiforms and coraciimorphs. Sandcoleids (Pl. I in Houde and Olson, 1992) and *Waltonavis* (Fig. 9i in Mayr and Kitchener, 2023e) appear to have a strongly developed ventral collateral ligamental tubercle on the ulna.

Stiller et al. (2024) and S. Wu et al. (2024) instead recovered Accipitrimorphae and Strigiformes as sister groups, forming a clade called Hieraves. Six character states were optimized as potential synapomorphies of this clade, though only one was inferred as such under both topologies.

- Keel apex on sternum caudal to main body of sternum (char. 41: 1 > 0, also found in *Eudromia*, Galliformes, *Monias*, *Opisthocomus*, *Aramus*, *Sarothrura*, *Rallus*, *Charadrius*, Scolopaci, *Turnix*, *Pagodroma*, *Fregata*, *Urocolius*, *Bucco*, *Micrastur*, and *Acanthisitta*); W24.
- Acrocoracoid process on coracoid straight (char. 63: 1 > 0, also found in *Gansus*, Anseriformes, *Ortalis*, *Phoenicopterus*, *Monias*, *Ardeotis*, Strisores, *Opisthocomus*, Gruiformes, *Gavia*, *Pagodroma*, Pelecanimorphae, *Urocolius*, *Psilopogon*, *Micrastur*, and *Acanthisitta*); W24.
- Supracoracoid nerve foramen in coracoid present (char. 70: 0 > 1, also found in *Ichthyornis*, *Anseranas*, *Phoenicopterus*, *Corythaeola*, Daedalornithes, Gruiformes, Charadrii, *Alca*, *Sterna*, *Phaethon*, Procellariiformes, *Leptoptilos*, *Eudocimus*, *Pelecanus*, *Leptosomus*, and *Micrastur*); S24, W24.
- Deltopectoral crest well developed, extending at least one third the total length of the humerus (char. 122: 1 > 0, also found in *Gansus*, *Ichthyornis*, *Corythaeola*, *Ardeotis*, *Nyctibius*, *Podargus*, *Spheniscus*, *Leptoptilos*, *Leucocarbo*, *Eudocimus*, *Scopus*, Cavitaves, *Micrastur*, and Passeri); W24.
- Radius craniocaudally curved (char. 144: 0 > 1, reversed in *Elanus*; also found in Anseriformes, *Phoenicopterus*, *Ardeotis*, Strisores, Grues, Charadrii, Feraequornithes, Cavitaves, *Micrastur*, Tyranni, and *Climacteris*); W24.
- Tendinal sulcus on carpometacarpus wraps around cranial surface of bone (char. 172: 0 > 1, reversed in *Pandion*; also found in *Nyctibius*, *Florisuga*, *Balearica*, Picocoraciades, *Micrastur*, and Passeriformes); S24.

A supracoracoid nerve foramen is present in *Ypresiglaux* (Mayr and Kitchener, 2023d) and the stem-strigiform *Primoptynx* (Mayr et al., 2020c), but has also been reported in sandcoleids (Houde and Olson, 1992; Ksepka et al., 2017; Mayr, 2018), *Ypresicolius* (Mayr and Kitchener, 2024g) and total-group leptosomids (Weidig, 2006; Mayr, 2008c; Mayr and Kitchener, 2023e), and may therefore be ancestral for a more inclusive clade.

Whereas Stiller et al. (2024) placed Hieraves within Afroaves, S. Wu et al. (2024) recovered it as the sister group to Australaves. Three character states were optimized as potential synapomorphies of this clade.

- Costal margin 25–75% of total sternum length (char. 35: 0 > 1, reversed in *Ninox* and Eupasseres; also found in *Ichthyornis*, Anseriformes, *Corythaeola*, *Streptoprocne*, and *Leptosomus*).
- Minor metacarpal prominently narrows distally in caudal view (char. 180: 0 > 1, also found in *Ichthyornis*, Anseres, Mirandornithes, Columbimorphae, *Aegotheles*, Gruiformes, Charadriiformes, Phaethontimorphae, Pelecanimorphae, *Leptosomus*, and *Coracias*).
- Prominent tubercle on minor metacarpal distal to proximal synostosis of metacarpals (char. 186: 0 > 1, reversed in *Pandion* and Psittacopasseres; also found in *Ichthyornis*, *Eudromia*, *Rollulus*, *Psophia*, *Burhinus*, *Turnix*, *Sterna*, *Phaethon*, *Leptosomus*, and *Coracias*).

Overall, a greater number of potential synapomorphies in this dataset support the Afroaves topology than the Eutelluraves or the Hieraves + Australaves topologies, though the presence of several potential afroavian synapomorphies in certain australavians (especially the falconiform *Micrastur*) raises the possibility that these characters may be telluravian symplesiomorphies. Similarly, a seemingly large number of characters supports Hieraves, but all of these traits are also found in members of Coraciimorphae and Australaves, rendering their inferences as hieravian synapomorphies questionable.

#### Affinities of *Opisthocomus*

One of the most contentious subjects in avian phylogenetics concerns the position of *Opisthocomus*. Jarvis et al. (2014) and Stiller et al. (2024) recovered it as the extant sister taxon of Cursorimorphae. Three character states were optimized as potential synapomorphies of this clade, all of which were inferred as such under both molecular topologies.

- Keel apex on sternum caudal to main body of sternum (char. 41: 1 > 0, reversed in *Psophia*, *Balearica*, *Podica*, *Burhinus*, *Alca*, and *Sterna*; also found in *Eudromia*, Galliformes, *Monias*, *Pagodroma*, *Fregata*, Accipitrimorphae, Strigiformes, *Urocolius*, *Bucco*, *Micrastur*, and *Acanthisitta*); J14, S24.
- Impression of m. sternocoracoidei on coracoid deep (char. 76: 0 > 1, reversed in *Psophia*, *Podica*, *Burhinus*, Scolopaci, and *Alca*; also found in *Gansus*, *Ichthyornis*, *Chauna*, *Phoenicopterus*, *Aegotheles*, *Spheniscus*, *Phoebastria*, *Coragyps*, *Coracias*, and *Cariama*); J14, S24.
- Tendinal sulcus on carpometacarpus barely perceptible (char. 170: 1 > 0, reversed in Gruoidea, Scolopaci, and *Sterna*; also found in *Podilymbus*, *Gavia*, *Spheniscus*, *Leucocarbo*, and *Urocolius*); J14, S24.

In contrast, Prum et al. (2015) recovered a clade uniting *Opisthocomus* and Telluraves to the exclusion of other neoavians, called Inopinaves. One character state was optimized as a potential synapomorphy of Inopinaves.

- Keel apex on sternum caudal to main body of sternum (char. 41: 1 > 0, reversed in Cavitaves, Psittaciformes, and Eupasseres; also found in *Eudromia*, Galliformes, *Monias*, *Aramus*, *Sarothrura*, *Rallus*, *Charadrius*, Scolopaci, *Turnix*, *Pagodroma*, and *Fregata*).

Kuhl et al. (2021) recovered *Opisthocomus* as the extant sister group of Strisores. Four character states were optimized as potential synapomorphies of a clade uniting *Opisthocomus* and Strisores to the exclusion of other neoavians.

- Acrocoracoid process on coracoid straight (char. 63: 1 > 0, reversed in *Aegotheles* and *Streptoprocne*; also found in *Gansus*, Anseriformes, *Ortalis*, *Phoenicopterus*, *Monias*, *Ardeotis*, Gruiformes, *Gavia*, *Pagodroma*, Pelecanimorphae, Accipitrimorphae, Strigiformes, *Urocolius*, *Micrastur*, and *Acanthisitta*).
- Pronounced ventral curvature of scapula (char. 92: 0 > 1, reversed in Letornithes; also found in Gruiformes, *Eurypyga*, *Spheniscus*, *Oceanites* + *Pagodroma*, *Leptoptilos*, *Tigrisoma*, *Coragyps*, *Pandion*, *Tyto*, *Merops*, *Alcedo*, *Psilopogon*, *Nestor*, *Acanthisitta*, and Tyranni).
- Apex of dorsal cotylar process of ulna approximately coplanar with dorsal surface of ulnar main body (char. 150: 0 > 1, reversed in *Aegotheles* and *Streptoprocne*; also found in *Ichthyornis*, Anseriformes, *Podilymbus*, *Corythaeola*, Charadriiformes, *Urocolius*, Eucavitaves, *Nestor*, and Passeriformes).
- Pneumatic foramen in infratrochlear fossa on carpometacarpus absent (char. 191: 1 > 0, reversed in *Streptoprocne*; also found in Mirandornithes, Pteroclimesites, *Rallus*, Scolopaci + Lari, Procellariimorphae, *Leucocarbo*, *Eudocimus*, *Tigrisoma*, *Ninox*, *Urocolius*, *Merops*, *Jynx*, *Pandion*, *Nestor*, and Passeriformes).

S. Wu et al. (2024) recovered *Opisthocomus* as the extant sister group of Phaethoquornithes. One character state was optimized as a potential synapomorphy of a clade uniting *Opisthocomus* and Phaethoquornithes to the exclusion of other neoavians.

- Costal margin 25–75% of total sternum length (char. 35: 0 > 1, reversed in *Tigrisoma*; also found in *Ichthyornis*, Anseriformes, *Corythaeola*, *Streptoprocne*, Gruiformes, *Charadrius*, *Limosa*, *Alca* + *Sterna*, Accipitrimorphae, *Tyto*, *Leptosomus*, and Australaves).

Well corroborated fossils of stem-opisthocomids are mostly based on fairly incomplete remains (Mayr et al., 2011; Mayr and De Pietri, 2014), though a straight acrocoracoid process can be seen in *Namibiavis* and *Hoazinavis* (Mayr et al., 2011), and a ventrally curved scapula is evidenced in *Protoazin* (Mayr and De Pietri, 2014). The pectoral girdle and forelimb skeleton provides limited support for preferring any of the three aforementioned alternative hypotheses for the phylogenetic position of *Opisthocomus*, given that essentially all of the potential synapomorphies listed above are found in members of more than one group that has been proposed to be its extant sister taxon. For example, the lone character state optimized as a potential synapomorphy of *Opisthocomus* + Telluraves is also present in members of Gruiformes and Charadriiformes, whereas the character states recovered as potential synapomorphies of *Opisthocomus* + Strisores or *Opisthocomus* + Phaethoquornithes are also present in some members of Gruiformes, Charadriiformes, or Telluraves. However, the presence of a barely perceptible tendinal sulcus on the carpometacarpus as a feature possibly linking *Opisthocomus* to Gruiformes and Charadriiformes is intriguing, considering that this character is otherwise primarily found in specialized diving taxa within Mirandornithes and Aequornithes.

In general, pectoral and forelimb skeletal characters appear to provide little unambiguous support for selecting among alternative molecular topologies. However, qualitative assessment of potential synapomorphies using the current dataset may weakly favor Mirandornithes as the extant sister group to all other neoavians (as opposed to being most closely related to Columbimorphae or Charadriiformes) and Accipitrimorphae as being more closely related to Strigiformes and Coraciimorphae than to Australaves (forming Afroaves, as opposed to being the extant sister group to all other telluravians). Although congruence between molecular and morphological sources of data can help increase confidence in specific phylogenetic hypotheses (Mayr, 2008d; Lee and Camens, 2009; Mayr, 2011a; Giribet, 2015; Lee and Palci, 2015), the conclusions here must still be treated with caution. Homoplasy is evidently pervasive in the avian pectoral girdle and forelimb skeleton, and furthermore, the present study did not identify potential synapomorphies for at least one clade (Eufalconimorphae) strongly supported by phylogenomic studies.

That the morphological characters compiled in the present study generally do not strongly favor specific molecular topologies is also evidenced by the fact that no significant differences in RHI values resulted from mapping these characters onto alternative molecular topologies. Tree-distance metrics relative to topologies derived from analyzing the morphological dataset also did not differ appreciably among the molecular topologies examined here. Of the three molecular topologies studied, tree-distance metrics indicate that, on average, the Kuhl et al. (2021) topology exhibits the least similarity to most of the topologies derived from analyzing the morphological dataset (though the Jarvis et al., 2014 and S. Wu et al., 2024 topologies produce slightly longer tree length when enforced under the present dataset). The Stiller et al. (2024) topology exhibits the lowest average RF distances and, along with the Prum et al. (2015) topology, also produces the shortest tree length, highest RI and CI, and lowest RHI (when ordered characters are considered) relative to the other molecular topologies. Stiller et al. (2024) reported that their phylogenomic topology was more congruent with the taxonomic distribution of nine continuous anatomical traits than that of Prum et al. (2015). The relatively high congruence of the Stiller et al. (2024) topology with morphological characters is upheld by our findings, despite extensive homoplasy in the present dataset.

In terms of RF distances, the results of analyzing the morphological dataset tended to exhibit the greatest similarity to molecular trees when constrained to the consensus tree of Braun and Kimball (2021). Although these results lead to the self-evident conclusion that increasing topological constraints informed by molecular topologies lead to greater congruence between morphological and molecular trees, they also highlight the challenge of identifying phylogenetic signal in avian pectoral and forelimb skeletal characters. This difficulty may partly be due to rampant homoplasy, but a mosaic distribution of morphological characters might also reflect the existence of a developmental zone of variability (*sensu* Bever et al., 2011) in the early evolutionary history of birds. Accentuated effects of these phenomena would furthermore be consistent with recent hypotheses that the early radiation of neoavians was characterized by rapid, near-simultaneous diversification (Jarvis et al., 2014; Prum et al., 2015; Suh, 2016; Houde et al., 2019; Kimball et al., 2019; Braun and Kimball, 2021; Brocklehurst and Field, 2024; Stiller et al., 2024), incomplete lineage sorting (Suh et al., 2015; Houde et al., 2020), and introgression (Braun and Kimball, 2021; Gatesy and Springer, 2022) among major lineages, and might not be best approximated by a strictly bifurcating branching pattern.

### Utility of Pectoral and Forelimb Osteological Characters in Avian Phylogenetics

The widespread distribution of most pectoral and forelimb characters optimized as synapomorphies of major avian clades, the limited congruence between results derived from unconstrained analyses of the morphological data and those of phylogenomic analyses, and heatmap visualizations of the phylogenetic distribution of morphological characters suggest that homoplasy is prevalent across the avian pectoral girdle and forelimb skeleton. Low phylogenetic signal in this anatomical region is further evidenced by generally low statistical support for deep nodes recovered by analyses of the present dataset (Figs. 8–15). This is consistent with studies that have concluded that body size and functional specializations, particularly those related to locomotion, exert substantial control on the skeletal morphology of the avian pectoral girdle and forelimb (Stegmann, 1963; Stegmann, 1964; Hui, 2002; Nudds et al., 2007; Habib and Ruff, 2008; Simons, 2010; Wang et al., 2011; Close and Rayfield, 2012; Voeten et al., 2018; Serrano et al., 2020; Karoullas and Nudds, 2021; Lowi-Merri et al., 2021; Mayr, 2021a; Mayr et al., 2021; Watanabe et al., 2021; Holmes et al., 2022; Smith et al., 2022; Akeda and Fujiwara, 2023; Eliason et al., 2023; Gündemir et al., 2023; Haidr, 2023; Lowi-Merri et al., 2023; Shatkovska and Ghazali, 2023; De Mendoza et al., 2024; Orkney and Hedrick, 2024; Segesdi et al., 2024). This trend may be especially evident in the results of our unconstrained analyses, which recovered, for example, one group containing the burst-flying *Eudromias*, Galliformes, *Monias*, and *Turnix*, and another uniting the diving *Podilymbus*, *Alca*, *Gavia*, and *Spheniscus*. Such observations furthermore align with previously reported evidence for strong evolutionary constraints on the morphological disparity (Wang and Zhou, 2023), complexity (Brinkworth et al., 2023), and modularity (Orkney and Hedrick, 2024) of the forelimb skeleton in birds and their close extinct relatives (though see Navalón et al., 2022 regarding complex intra-skeletal and intra-clade variability in the establishment of these evolutionary patterns). Considerable variation with little obvious correlation with either phylogeny or function has furthermore been previously reported in the sternal morphology of passeriforms (Heimerdinger and Ames, 1967; Webster, 1992). Our results also broadly parallel those of Ericson and Qu (2024), who found evidence of pervasive homoplasy and few unambiguous synapomorphies characterizing deep avian divergences captured in the morphological dataset of Livezey and Zusi (2006), especially among postcranial osteological characters.

These findings echo previous recommendations for caution in assessing the phylogenetic placement of avian or avian-like fossil specimens known only from isolated pectoral and forelimb elements, especially those potentially belonging to stem-group lineages for which morphological variation is poorly understood (Longrich et al., 2011; Mohr et al., 2021). Nonetheless, even if subject to homoplastic gains and losses, some of the characters identified in the present study seem to represent plausible synapomorphies of clades that have hitherto been difficult to characterize morphologically (such as Phaethontimorphae and Telluraves), and are thus potentially informative for understanding morphological evolution across Neornithes and identifying fossil representatives of these groups. Therefore, the optimization of thoroughly vetted anatomical characters onto molecular phylogenetic topologies may prove fruitful for diagnosing recently recognized avian clades, as has been previously suggested (Mayr, 2011a; Mayr, 2014c; Steell et al., 2023a). Among the individual elements of the pectoral girdle and forelimb skeleton, the humerus and carpometacarpus may be relatively reliable for assessing phylogenetic relationships according to our heatmap visualizations (though see Steell et al., 2023a regarding widespread homoplasy in carpometacarpal characters within Passeriformes), potentially increasing confidence in the identification of fossil bird taxa based on these bones.

However, the absence of fossil crown bird taxa in the present dataset should be kept in mind as a notable limitation. Fossils provide the only direct evidence of transitional ancestral morphologies that have been lost or modified in extant organisms, and as a result have the potential to dramatically affect inferences about phylogenetic topology and character polarity (e.g., Gauthier et al., 1988; Donoghue et al., 1989; Mayr, 2007a; Mayr and Knopf, 2007; Mayr, 2008d; Mayr, 2014c; Hsiang et al., 2015; Puttick, 2016; Chen et al., 2019; Field et al. 2020; Mongiardino Koch and Parry, 2020; Mongiardino Koch et al., 2021; Asher and Smith, 2022; Beck et al., 2022; Kuo et al., 2023; Phillips et al., 2023). This effect is particularly noticeable in the present study regarding the optimization of pectoral and forelimb synapomorphies for crown birds and neognaths. The presence of two burst-flying groups (Tinamidae and Galliformes) among the extant outgroups to Neoaves likely leads to the reconstruction of some adaptations related to burst flight as ancestral to crown birds, even though these features are lacking in potential stem-paleognaths and early stem-galliforms (see previous discussion on potential synapomorphies of Neornithes and Neognathae). Our heatmap visualizations further indicate that crown galliforms (especially the phasianid *Rollulus*) do not closely resemble crownward stem-birds in their overall pectoral girdle and forelimb osteology (Fig. 20). The fact that our equal weights analysis without topological constraints recovered a clade containing *Chauna*, *Dendrocygna*, and *Phoenicopterus* as the sister taxon to all remaining crown birds might suggest that the pectoral girdle and forelimb skeleton of Anseriformes and Phoenicopteriformes are more reminiscent of plesiomorphic states in the avian crown group, recalling the recent conclusion that elements of the palate in extant anseriforms may approximate the crown bird ancestral condition (Benito et al., 2022b). As noted by previous authors (Ericson, 1997; Ericson, 1999; Zelenkov, 2021; Houde et al., 2023), the bones comprising the pectoral girdle and forelimb apparatus in putative stem-anseriforms share noticeable similarities with those of putative stem-mirandornitheans and some Cretaceous avialans. Careful reconsideration of Mesozoic fossils that have been assigned to total-group Anseriformes based on elements from these anatomical regions (e.g., Hope, 2002; Kurochkin et al., 2002; De Pietri et al., 2016; Novas et al., 2019; Acosta Hospitaleche et al., 2023) may therefore be warranted.

Whether the addition of characters from other anatomical regions and the incorporation of information from a representative sample of fossil avian taxa will increase congruence between the results of morphological and molecular phylogenetic analyses for birds remains to be seen, especially given that recent research has indicated that different anatomical regions may favor different phylogenetic topologies in macroevolutionary studies of some vertebrate taxa (Benson, 2012; Mounce et al., 2016; Sansom and Wills, 2017; Sansom et al., 2017; Li et al., 2020; Callender-Crowe and Sansom, 2022; Ericson and Qu, 2024). Although large-scale morphological phylogenetic datasets have been criticized on the grounds that they may contain relatively little phylogenetic signal (Mayr, 2008a; Yu et al., 2021), increased dataset size is an inevitable outcome of expanding taxonomic and character scope (Laing et al., 2018). Even if incongruent with phylogenomic studies when analyzed on their own, large morphological datasets may yet prove useful for identifying synapomorphies of major clades, placing fossils into phylogenetic context, and studying rates of character evolution.

On top of increasing taxon and character sampling, various methodological techniques have been suggested to improve the accuracy of morphological phylogenetic investigation. These include Bayesian analyses (Wright and Hillis, 2014; O’Reilly et al., 2016; Puttick et al., 2017; O’Reilly et al., 2018; Puttick et al., 2019; Smith, 2019; Keating et al., 2020; Vernygora et al., 2020; Barbosa et al., 2024), especially the implementation of tip dating (Mongiardino Koch et al., 2021; Barido-Sottani et al., 2023; Mongiardino Koch et al., 2023; Lian et al., 2024); implied weights parsimony analyses (Goloboff et al., 2018; Smith, 2019; Rio and Mannion, 2021); the use of continuous characters (Parins-Fukuchi, 2018; Rio and Mannion, 2021); the selective removal of homoplasy-prone characters (Dávalos et al., 2014; Zou and Zhang, 2016); the inclusion of hypothetical ancestral morphologies as distinct operational taxonomic units (Asher et al., 2019); and correcting for ecological signal in morphological datasets (Phillips et al., 2023). However, the value of some of these techniques remains contested, and probably no single method on its own will emerge as a panacea for increasing congruence between morphological and molecular phylogenies (Dávalos et al., 2012; Sookias, 2020; Brady and Springer, 2021; Asher and Smith, 2022; Keating et al., 2023; Phillips et al., 2023; Holvast et al., 2024). In the present study, implied weights parsimony and especially Bayesian analyses produced trees slightly more congruent with molecular topologies than equal weights parsimony analysis under unconstrained topologies. Notably, our unconstrained Bayesian analysis (Analysis 3) likely represents the first morphological phylogenetic analysis to recover the monophyly of Heliornithes without topological constraints or a combined morphological–molecular dataset (compare Livezey, 1998; Musser and Cracraft, 2019; Musser et al., 2019). However, under partial topological constraints, our implied weights parsimony and Bayesian analyses did not consistently recover results substantially closer to molecular topologies compared to equal weights parsimony analyses. Irrespective of dataset size and method choice, clear character construction and repeatability of character scoring is essential to all morphological phylogenetic analyses (Simões et al., 2017; Sookias, 2020; Khakurel et al., 2024).

## Conclusion

The present study represents part of an initiative to produce a comprehensive morphological phylogenetic dataset for crown-group birds. By analyzing a novel dataset that extensively samples characters from the pectoral girdle and forelimb skeleton of a phylogenetically diverse set of extant birds, we found that this anatomical region does appear to exhibit substantial signal conflict with molecular sources of data, perhaps in part as a result of homoplasy driven by functional convergence. However, optimizing patterns of character acquisition onto molecular topologies recovered plausible synapomorphies for several clades for which morphological support had been largely unrecognized. Future work sourcing characters from other anatomical regions, incorporating key fossil taxa, and experimenting with additional phylogenetic methods may further clarify controversial aspects of avian systematics and clarify patterns of morphological evolution during the early evolutionary history of crown birds.

## Acknowledgements

We would like to thank Juan Benito Moreno for assistance with scoring characters for *Ichthyornis*, Matt Wills, Gabe Bever, Rob Sansom, and John D’Angelo for helpful discussion, and the Willi Hennig Society for making TNT a freely available software. This project was partly funded by an Allison R. “Pete” Palmer Student Research Award to A.C. by the Paleontological Society. D.J.F. is supported by UK Research and Innovation Future Leaders Fellowship MR/S032177/1. Additional funding for the project was provided by the European Research Council Starting Grant: TEMPO (ERC-2015-STG-677774) to R.B.J.B. For the purpose of open access, the authors have applied a Creative Commons Attribution (CC BY) license to any Author Accepted Manuscript version arising.

