## Appendix S1 for "Towards a comprehensive anatomical matrix for crown birds: phylogenetic insights from the pectoral girdle and forelimb skeleton"

**Appendix S1: Character List and Descriptions**

Characters coded with ordered states are indicated by an asterisk (*). References do not represent an exhaustive list of studies in which these characters have been used.

1. Furcula, overall form of shafts in cranial or caudal view (Fig. S1; Ericson, 1997: 44; Worthy and Lee, 2008: 123; Ksepka, 2009: 29; Worthy et al., 2017: 114; Musser and Clarke, 2020: 416):

0 – U-shaped, with essentially parallel clavicular shafts

1 – V-shaped, with clavicular shafts that diverge omally

Furcula could not be located in *Nyctibius* specimen; scored based on photographs of *Nyctibius jamaicensis* USNM 557530.

<https://panamabiota.org/stri/imagelib/imgdetails.php?imgid=4922>

<https://panamabiota.org/stri/imagelib/imgdetails.php?imgid=4923>

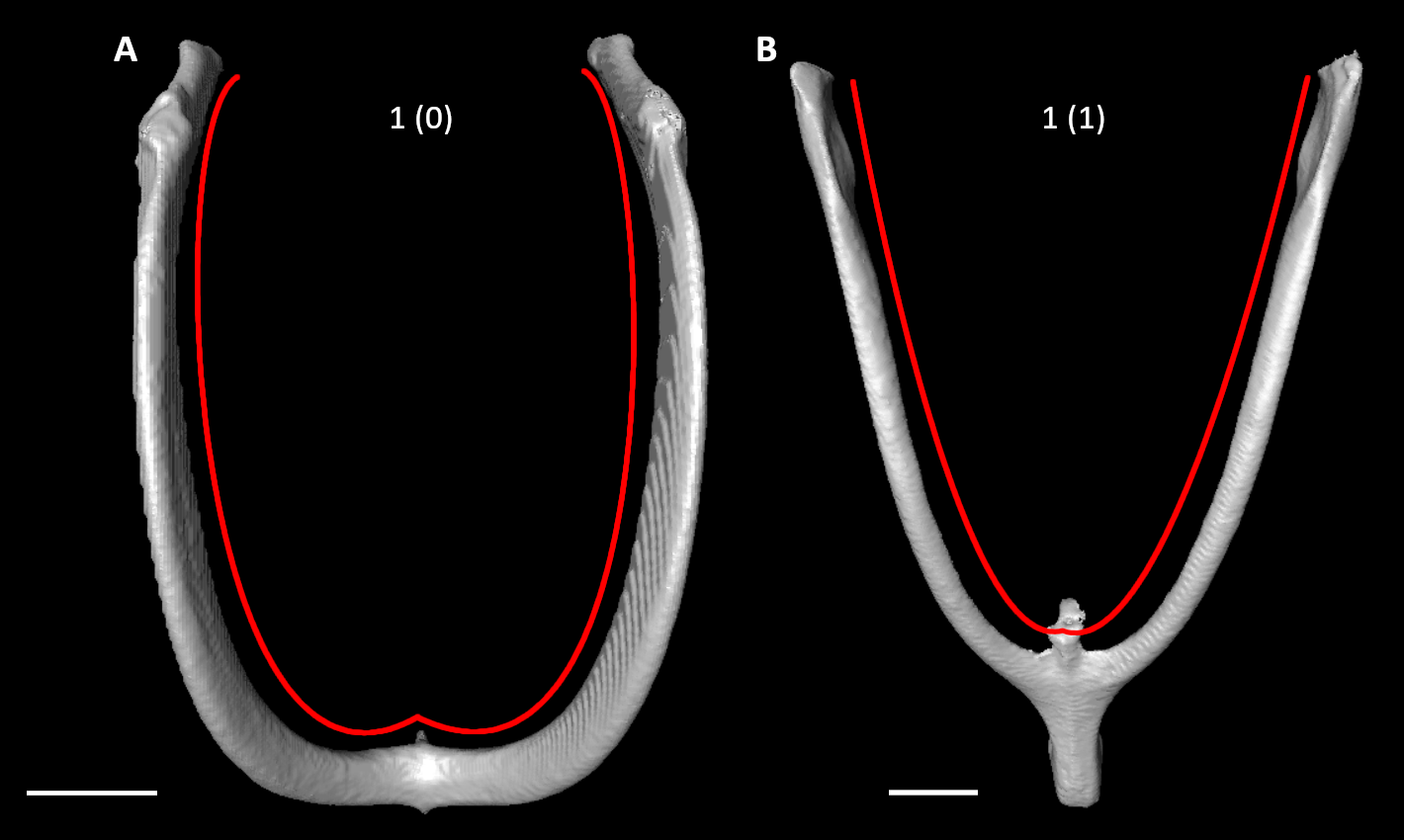

**Figure S1.** Furculae of *Limosa lapponica* (**A**) and *Podica senegalensis* (**B**) in cranial view, illustrating alternative states for character 1 (complete character description in text). Red lines indicate approximate curvature of furcular shafts. Scale bars = 5 mm.

2. Furcula, craniocaudal curvature in lateral view, exclusive of omal extremities (Fig. S2; Livezey and Zusi, 2006: 1233; Smith, 2010: 147; Musser and Clarke, 2020: 423):

0 – weakly to moderately convex cranially

1 – strongly curved and convex, approaching subcircular form

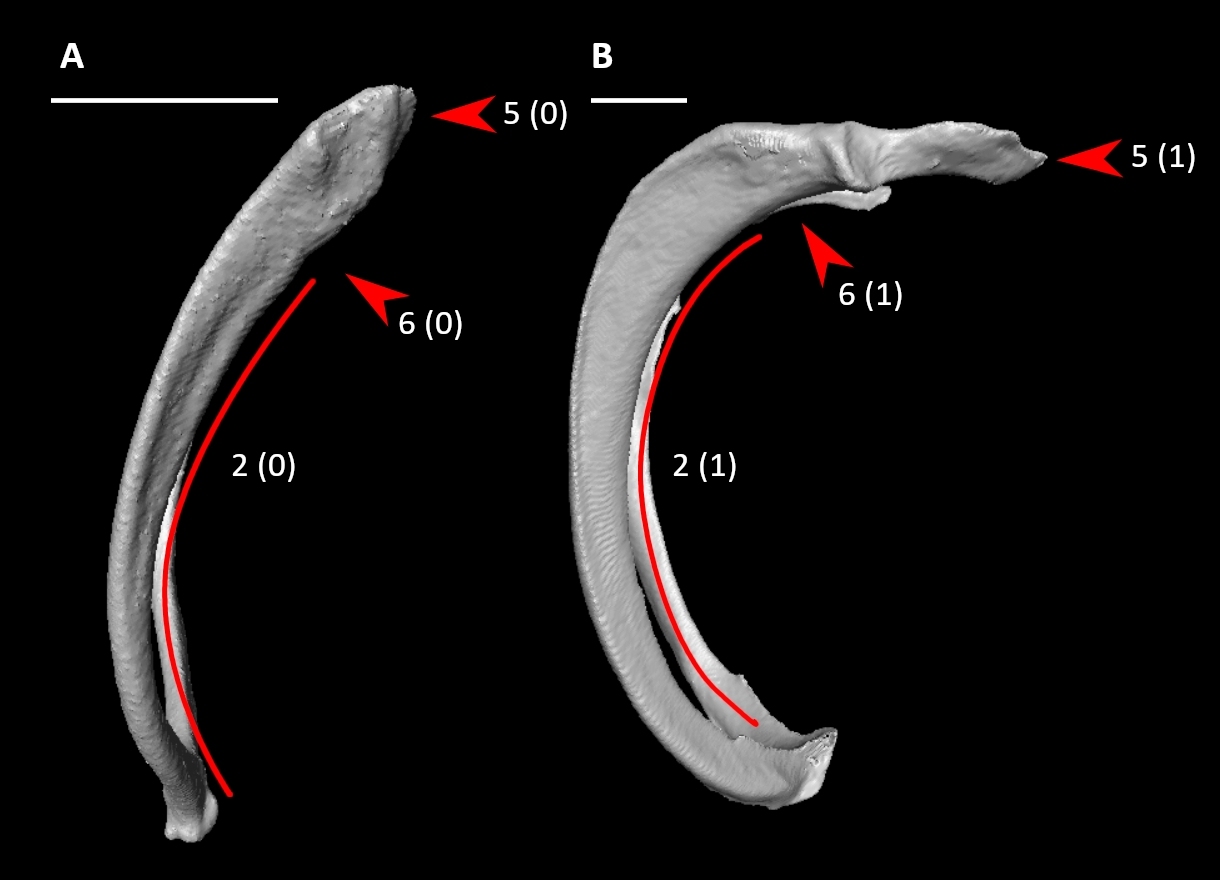

**Figure S2.** Furculae of *Aegotheles cristatus* (**A**) and *Alca torda* (**B**) in left (**A**) or right (**B**, mirrored) lateral view, illustrating alternative states for characters 2, 5, and 6 (complete character descriptions in text). Red lines indicate approximate craniocaudal curvature of furcular shafts. Arrows for character 5 indicate the tip of the acromial process, whereas arrows for character 6 indicate the base of the acromial process. Scale bars = 5 mm.

3. Furcula, protruding articular facets for acrocoracoids (Fig. S3; Mayr and Clarke, 2003: 62; Mayr, 2010: 25; Ksepka et al., 2013: 43):

0 – absent or weak

1 – prominent, may extend laterally or cranially

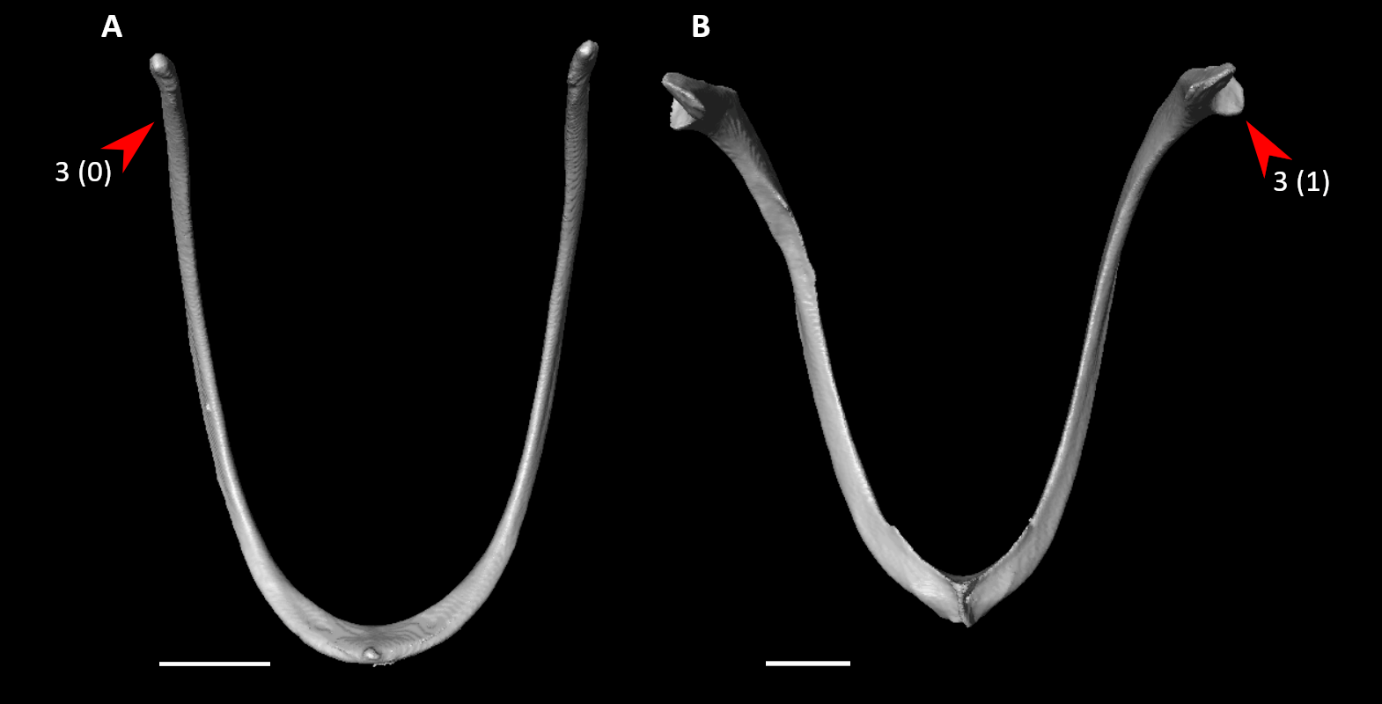

**Figure S3.** Furculae of *Podilymbus podiceps* (**A**) and *Alca torda* (**B**) in caudal view, illustrating alternative states for character 3 (complete character description in text). Arrows indicate articular facets for the acrocoracoid, or approximate homologous site for the facets. Scale bars = 5 mm.

4. Furcula, fusion at midline (Fig. S4; Field and Hsiang, 2018: 153):

0 – unfused

1 – fused

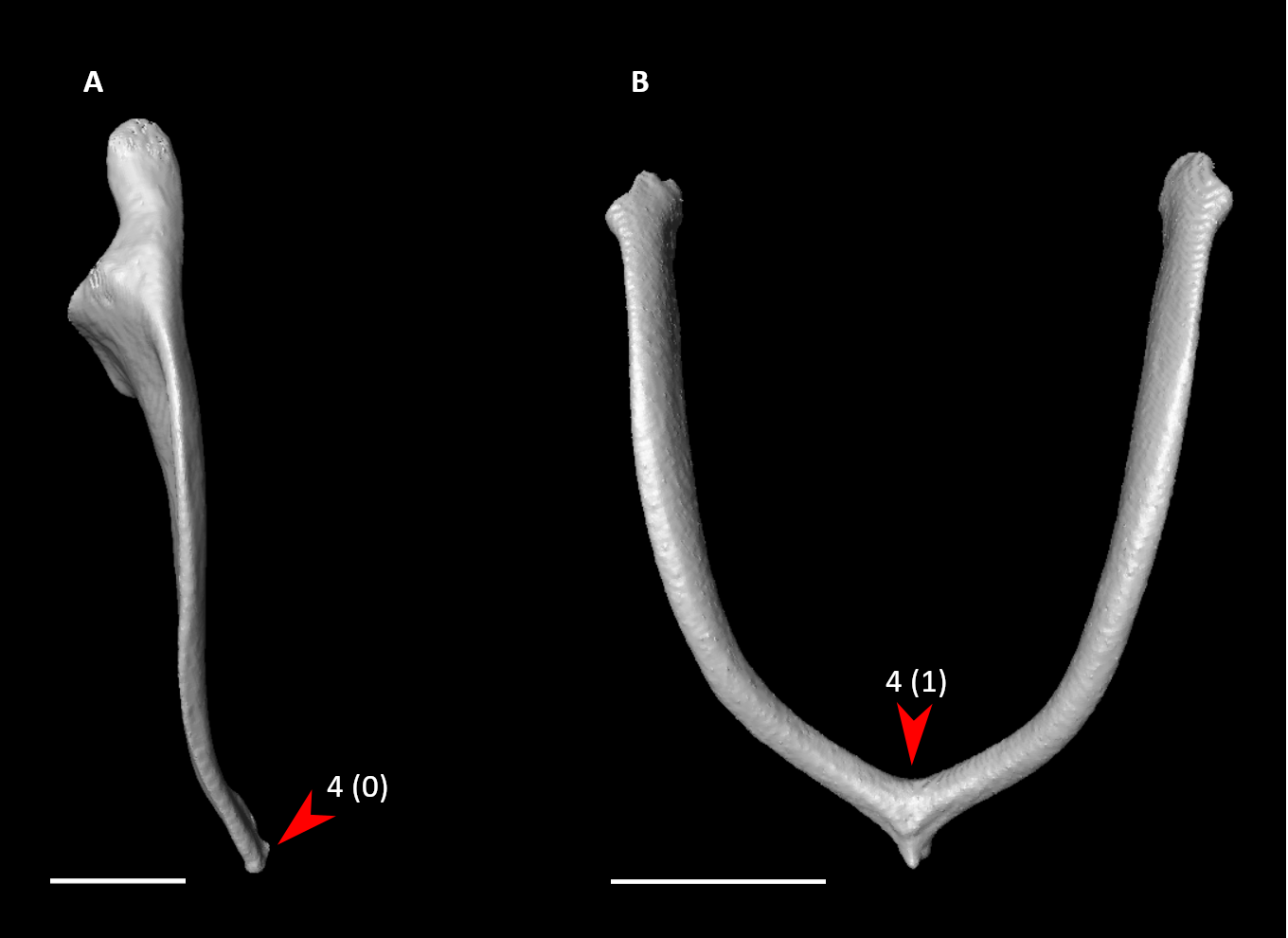

**Figure S4.** Clavicular shaft of *Corythaeola cristata* (**A**, right element) and furcula of *Aegotheles cristatus* (**B**) in cranial view, illustrating alternative states for character 4 (complete character description in text). Arrows indicate site of interclavicular articulation. Scale bars = 5 mm.

5. Furcula, acromial processes, omal extremities (Fig. S2; Ericson, 1997: 45; Worthy et al., 2017: 110):

0 – rounded

1 – pointed

6. Furcula, acromial processes, craniocaudal orientation (Fig. S2; Musser and Clarke, 2020: 430):

0 – slightly caudally inflected

1 – strongly caudally inflected, forming angle of approximately 90 degrees relative to clavicular shafts

7. Furcula, interclavicle dorsal process (Fig. S5; Livezey, 1998: 170; Musser and Clarke, 2020: 421):

0 – absent

1 – present

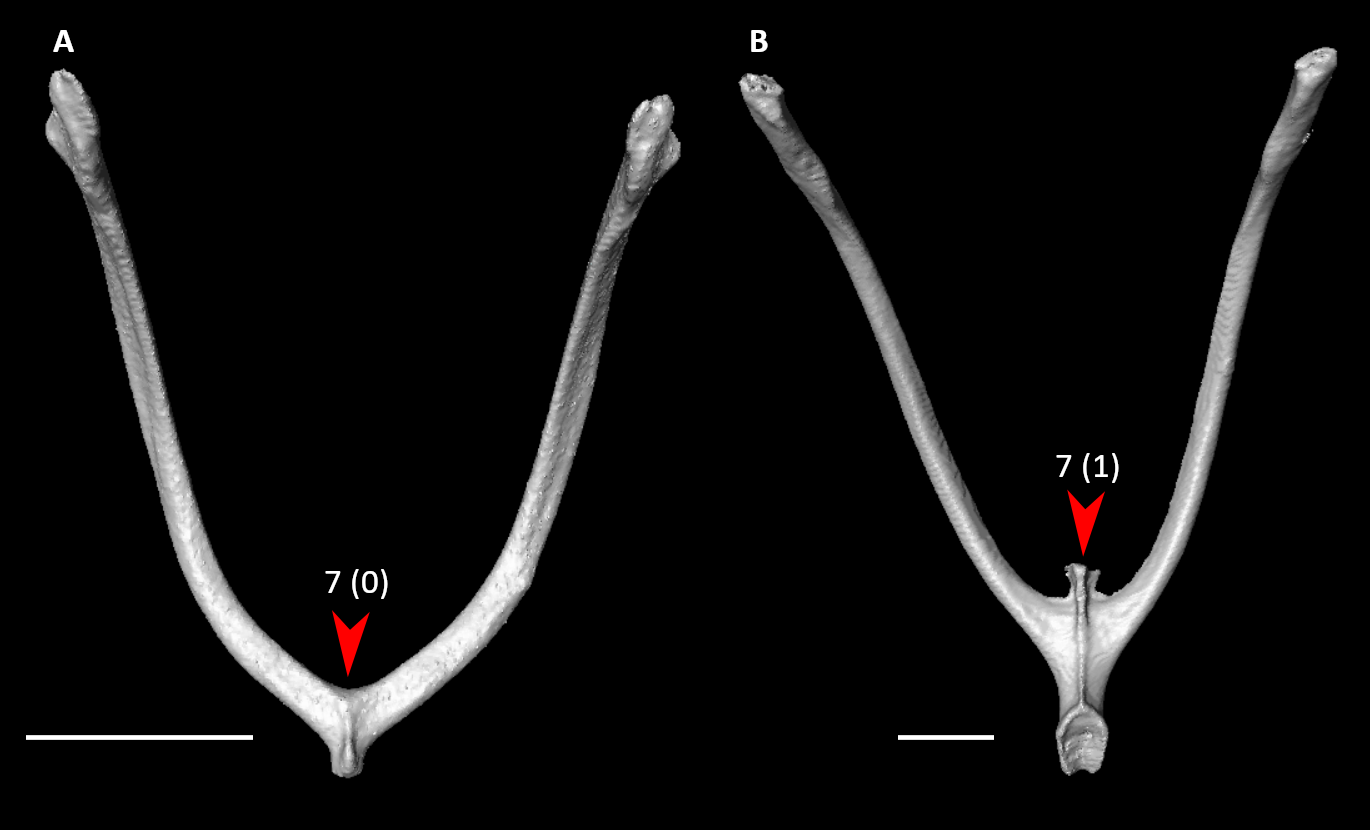

**Figure S5.** Furculae of *Aegotheles cristatus* (**A**) and *Podica senegalensis* (**B**) in caudal view, illustrating alternative states for character 7 (complete character description in text). Arrows indicate the interclavicle process or approximate homologous site. Scale bars = 5 mm.

8. Furcula, hypocleideum (Fig. S6; Musser and Clarke, 2020: 417):

0 – absent

1 – present

Broken in *Pagodroma* specimen; scored based on condition in other procellariiforms.

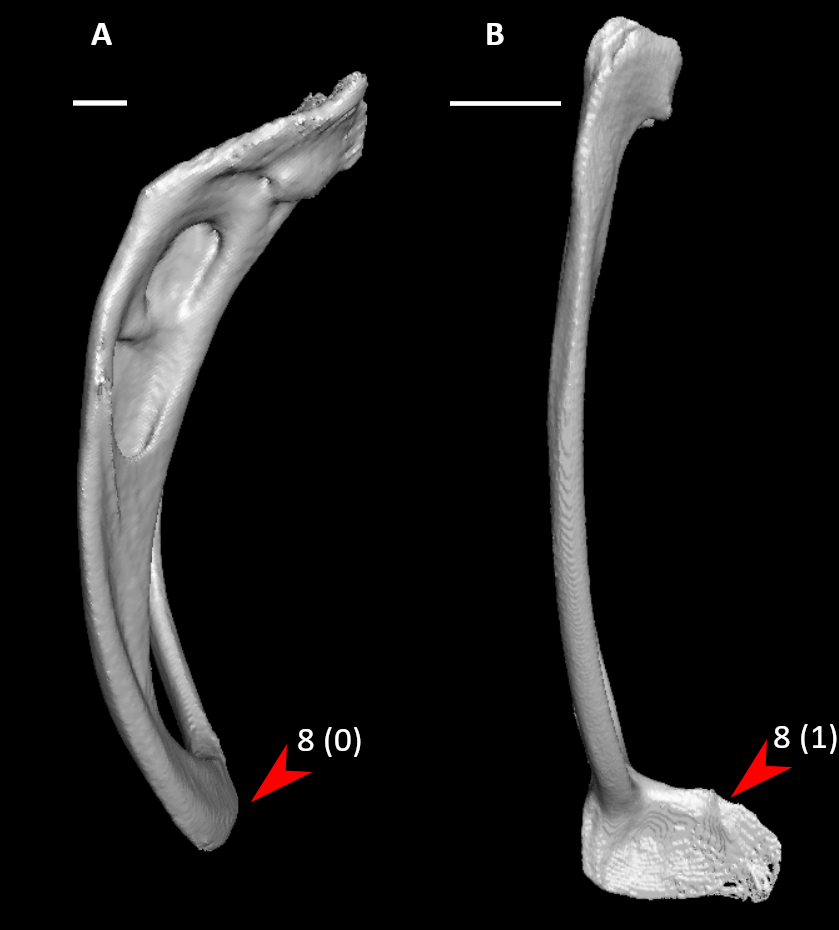

**Figure S6.** Furculae of *Balearica pavonina* (**A**) and *Rollulus rouloul* (**B**) in right lateral view (mirrored), illustrating alternative states for character 8 (complete character description in text). Arrows indicate the hypocleideum or approximate homologous site. Scale bars = 5 mm.

*9. Furcula, hypocleideum length measured in lateral view (Fig. S7):

0 – less than 5% total length of the furcula

1 – between 5% and 10% total length of the furcula

2 – greater than 10% total length of the furcula

Scored as inapplicable in *Fregata*, in which the hypocleideum is fused to the sternum.

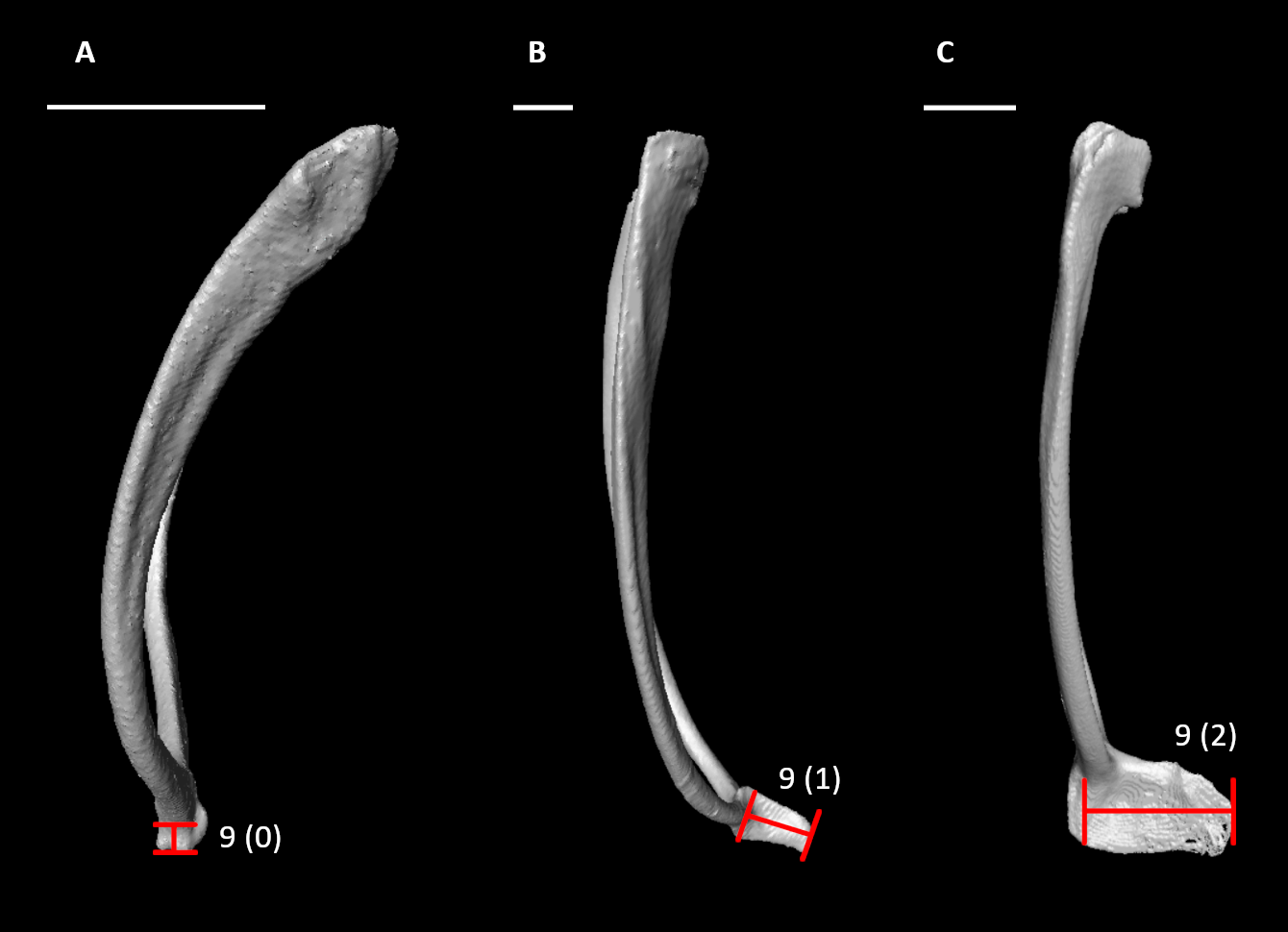

**Figure S7.** Furculae of *Aegotheles cristatus* (**A**), *Alectura lathami* (**B**), and *Rollulus rouloul* (**C**) in left (**A, B**) or right (**C**, mirrored) lateral view, illustrating alternative states for character 9 (complete character description in text). Red lines span the length of the hypocleideum. Scale bars = 5 mm.

*10. Furcula, hypocleideum, orientation of primary axis in lateral view (Fig. S8):

0 – dorsocaudally deflected from clavicular shafts

1 – continues curvature of clavicular shafts

2 – ventrocranially deflected from clavicular shafts

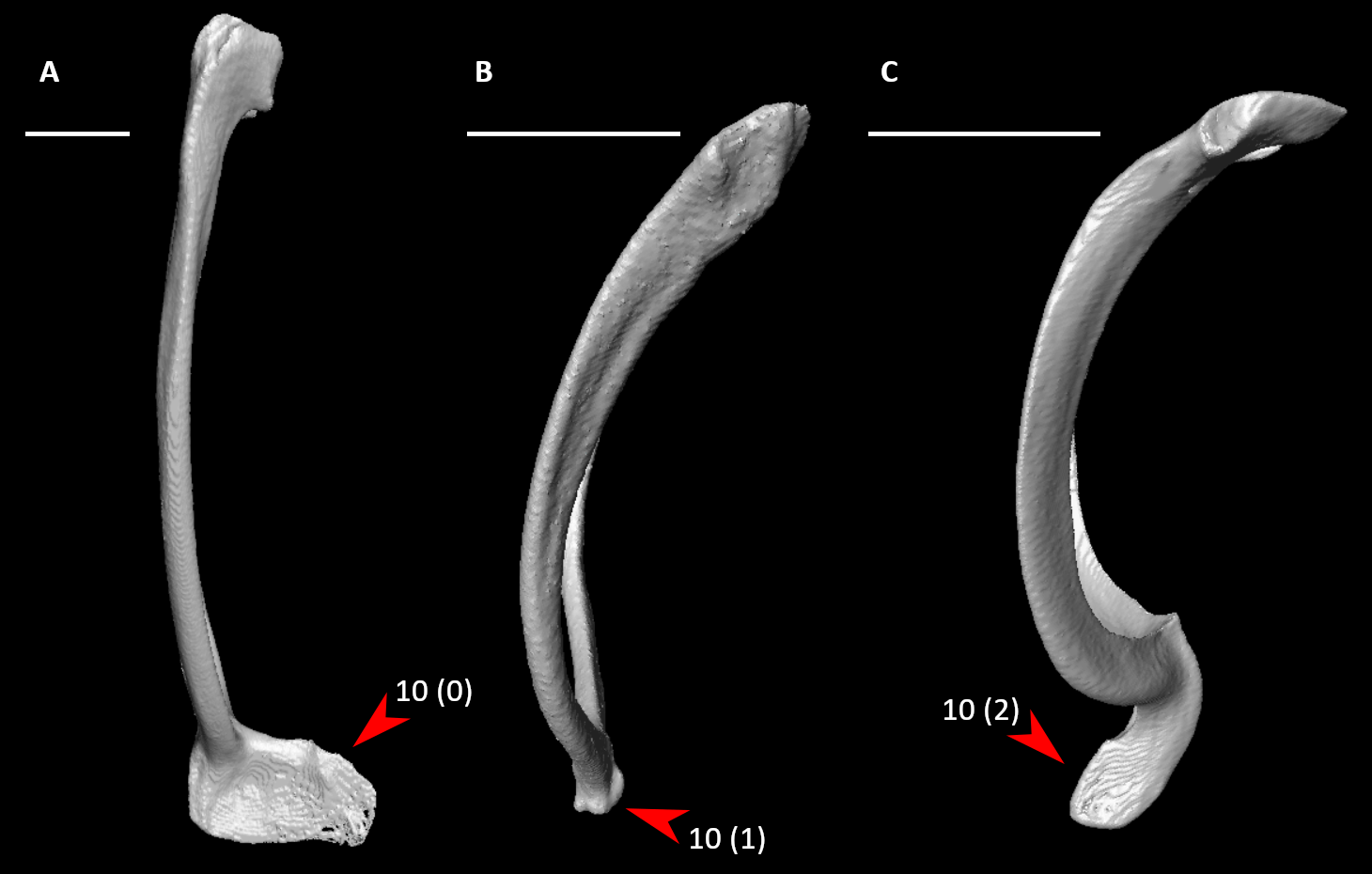

**Figure S8.** Furculae of *Rollulus rouloul* (**A**), *Aegotheles cristatus* (**B**), and *Oceanites oceanicus* (**C**) in right (**A**, mirrored) or left (**B, C**) lateral view, illustrating alternative states for character 10 (complete character description in text). Arrows indicate the hypocleideum. Scale bars = 5 mm.

11. Furcula, symphysis (Fig. S9; Musser and Clarke, 2020: 427):

0 – craniocaudally compressed

1 – dorsoventrally compressed

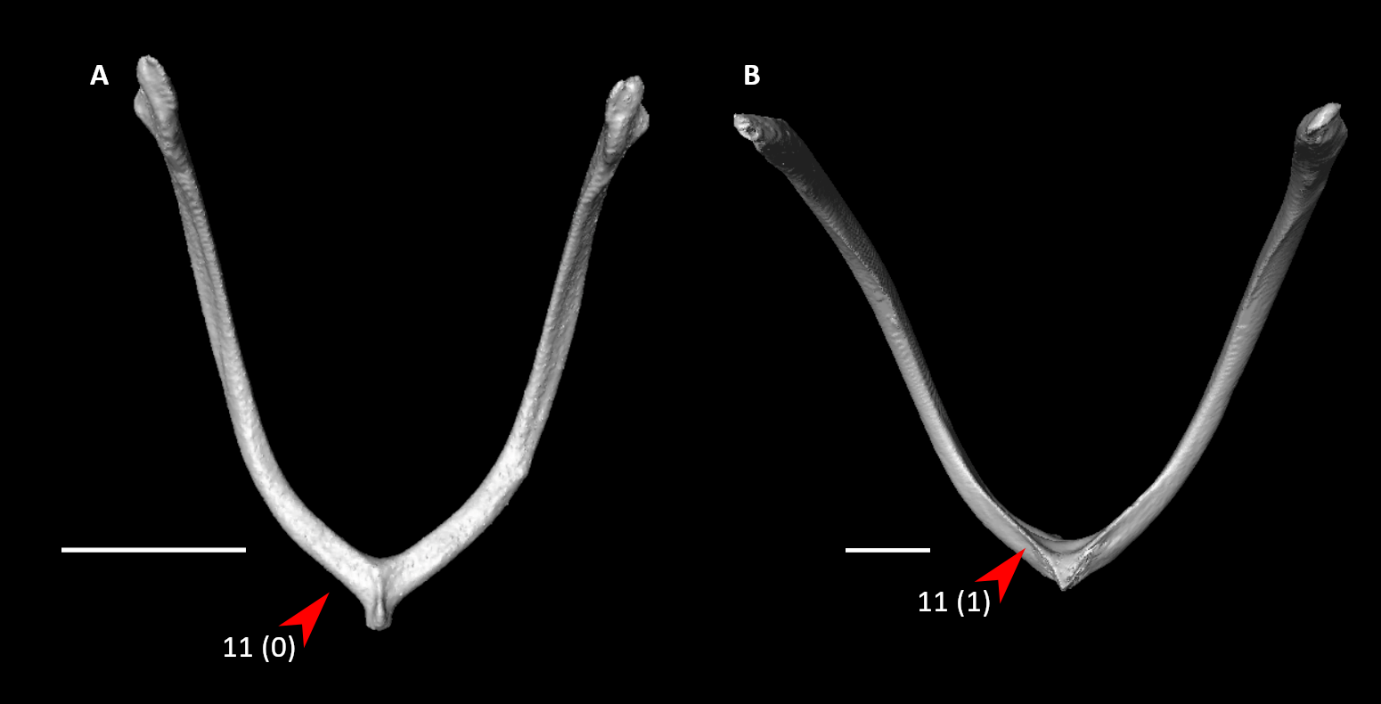

**Figure S9.** Furculae of *Aegotheles cristatus* (**A**) and *Phaethon lepturus* (**B**) in caudal view, illustrating alternative states for character 11 (complete character description in text). Arrows indicate the furcular symphysis. Scale bars = 5 mm.

12. Furcula, clavicular shafts (Fig. S10; Musser and Clarke, 2020: 426):

0 – craniocaudally compressed

1 – mediolaterally compressed

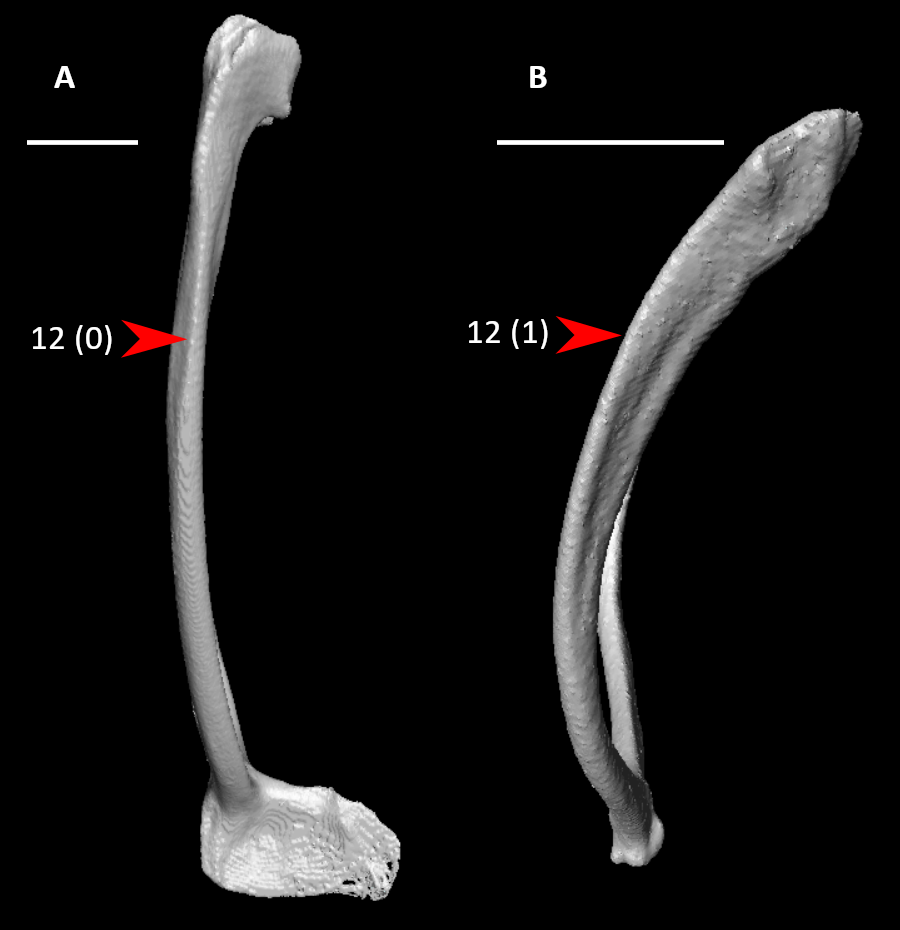

**Figure S10.** Furculae of *Rollulus rouloul* (**A**) and *Aegotheles cristatus* (**B**) in right (**A**, mirrored) or left (**B**) lateral view, illustrating alternative states for character 12 (complete character description in text). Arrows indicate the furcular shaft. Scale bars = 5 mm.

13. Furcula, clavicular shafts, pneumatic foramina on lateral surfaces (Fig. S11; Livezey, 1986: 105; Livezey, 1996: 35; Worthy and Lee, 2008: 122; Worthy et al., 2017: 113; Musser and Clarke, 2020: 424):

0 – absent

1 – present

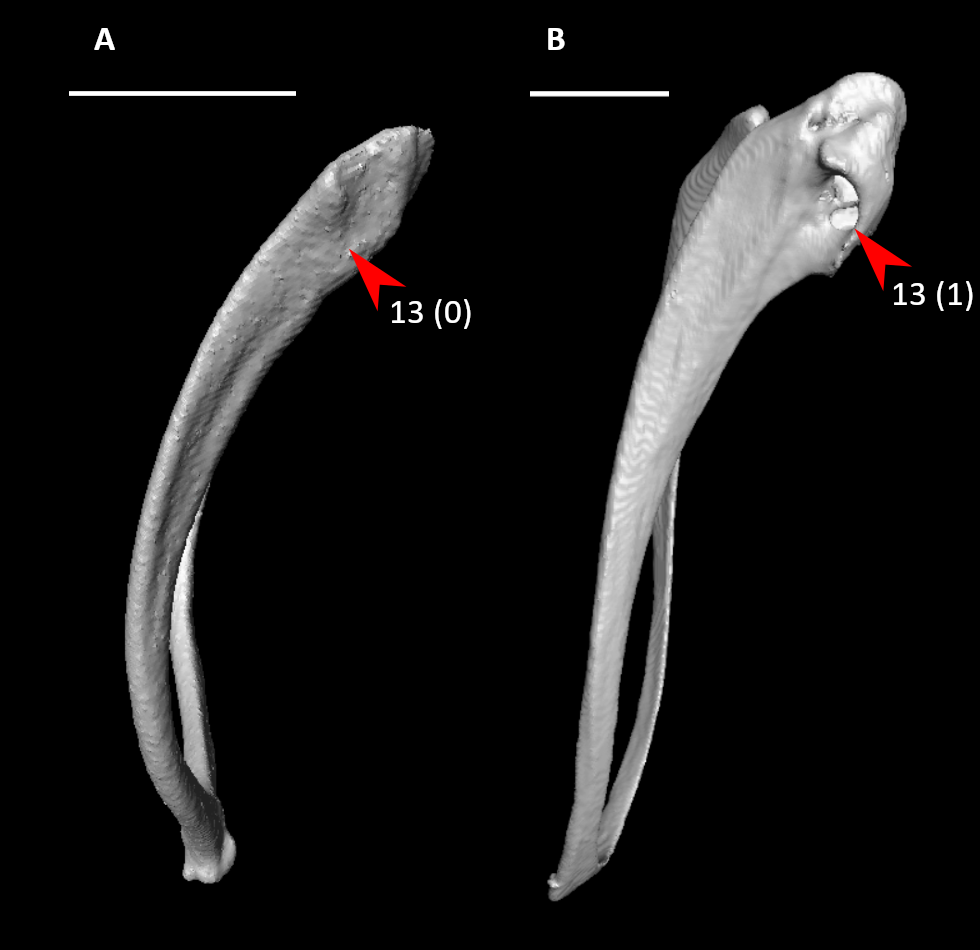

**Figure S11.** Furculae of *Aegotheles cristatus* (**A**) and *Ninox novaeseelandiae* (**B**) in left (**A**) or right (**B**, mirrored) lateral view, illustrating alternative states for character 13 (complete character description in text). Arrows indicate the pneumatic foramen on the lateral surface of the furcular shaft, or approximate homologous site to the foramen. Scale bars = 5 mm.

14. Sternum, external spine (Fig. S12; Musser and Clarke, 2020: 294):

0 – absent

1 – present

Scored as inapplicable in *Opisthocomus*, as furcula fuses to sternum at site where external spine would be.

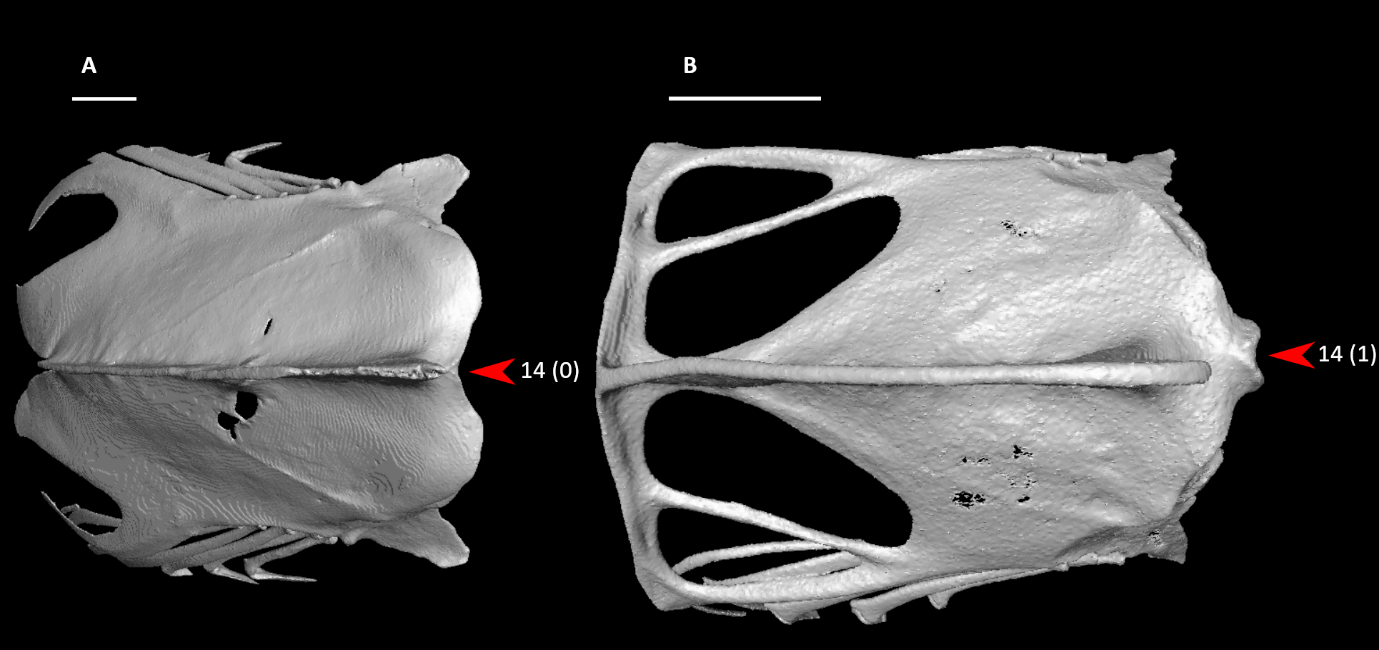

**Figure S12.** Sterna of *Podilymbus podiceps* (**A**) and *Aegotheles cristatus* (**B**) in cranioventral (**A**) or ventral (**B**) view, illustrating alternative states for character 14 (complete character description in text). Arrows indicate the external spine, or approximate homologous site. Scale bars = 5 mm.

*15. Sternum, external spine length (Fig. S13):

0 – less than 5% total sternum length

1 – between 5% and 10% total sternum length

2 – between 10% and 20% total sternum length

3 – greater than 20% total sternum length

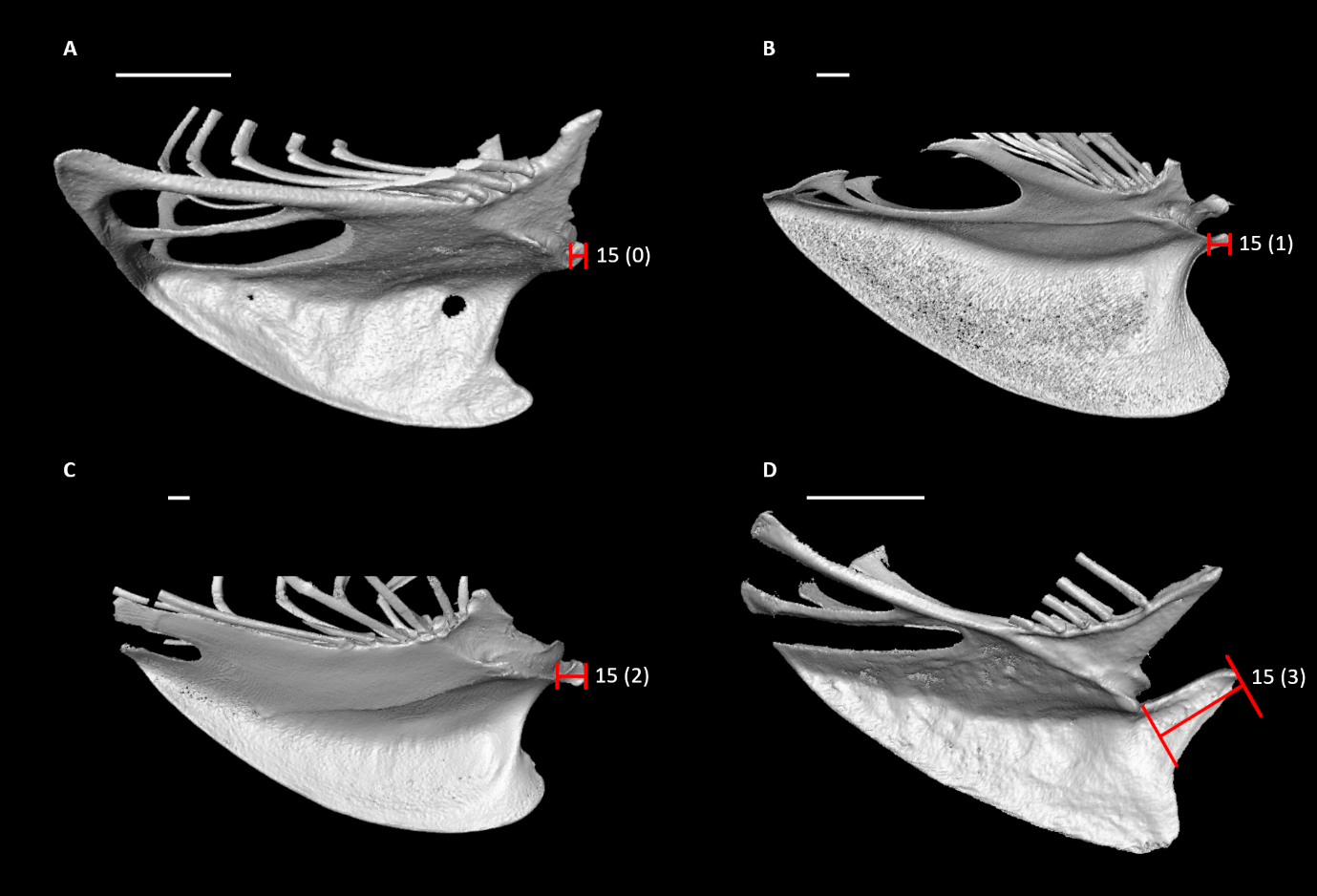

**Figure S13.** Sterna of *Aegotheles cristatus* (**A**), *Columba livia* (**B**), *Phoenicopterus ruber* (**C**), and *Jynx torquilla* (**D**) in right (**A, B, C**) or left (**D**, mirrored) lateral view, illustrating alternative states for character 15 (complete character description in text). Red lines span the length of the external spine. Scale bars = 5 mm.

16. Sternum, external spine in lateral view (Fig. S14):

0 – not tapered or weakly tapered

1 – distinctly tapered, forming sharp point

Scored inapplicable in *Psilopogon* due to fusion with the sternal keel.

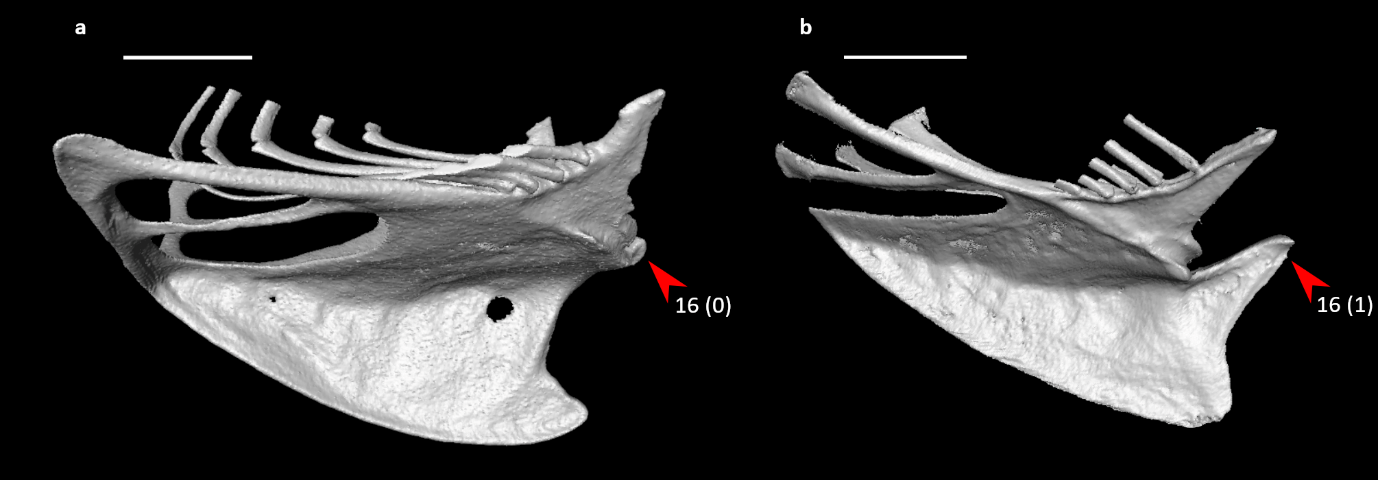

**Figure S14.** Sterna of *Aegotheles cristatus* (**A**) and *Jynx torquilla* (**B**) in right (**A**) or left (**B**, mirrored) lateral view, illustrating alternative states for character 16 (complete character description in text). Arrows indicate the tip of the external spine. Scale bars = 5 mm.

*17. Sternum, external spine in lateral view (Fig. S15; Ksepka et al., 2019: 39):

0 – dorsally oriented

1 – cranially oriented

2 – ventrally oriented

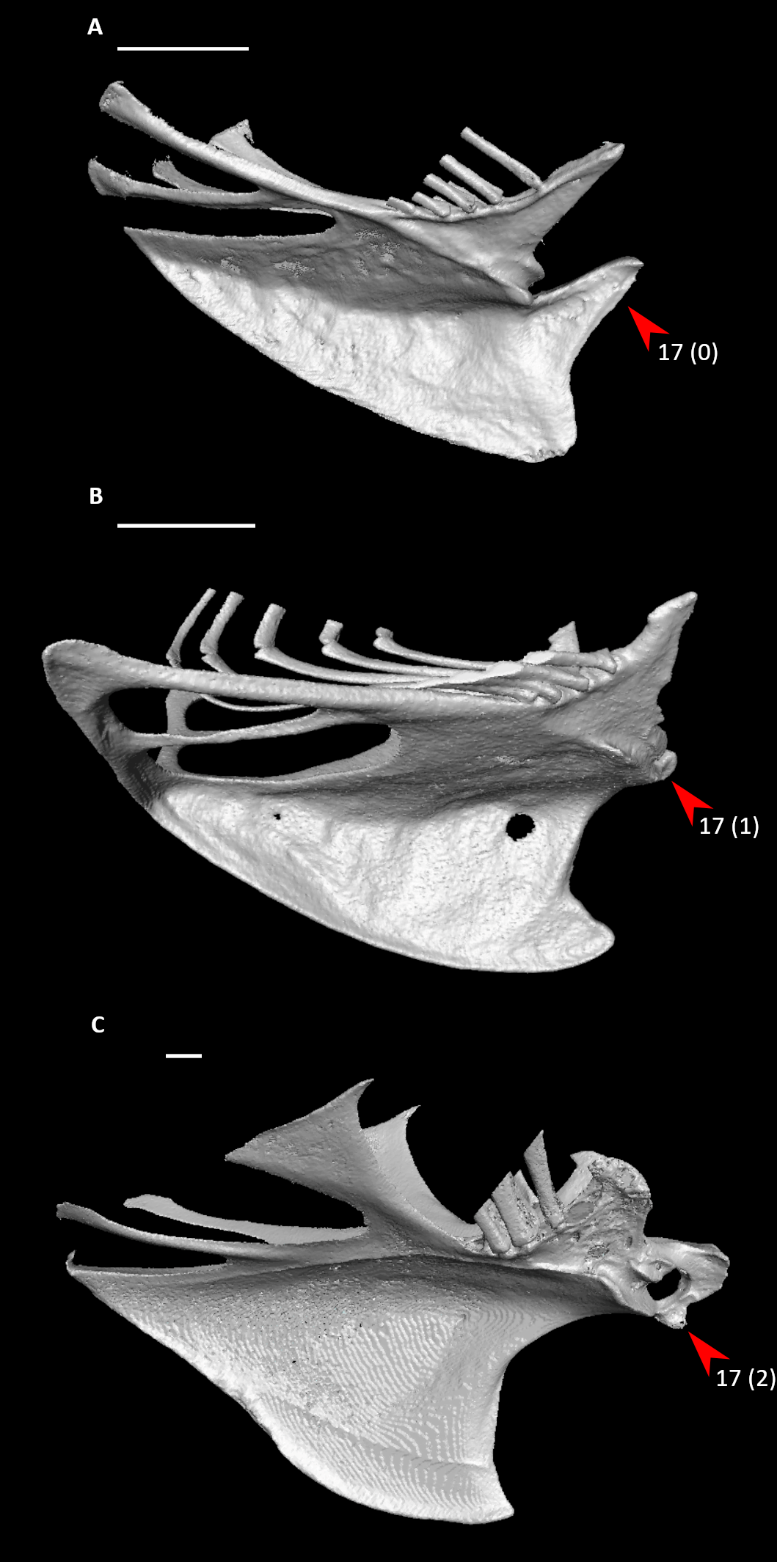

**Figure S15.** Sterna of *Jynx torquilla* (**A**), *Aegotheles cristatus* (**B**), and *Alectura lathami* (**C**) in left (**A, C**, mirrored) or right (**B**) lateral view, illustrating alternative states for character 17 (complete character description in text). Arrows indicate the external spine. Scale bars = 5 mm.

18. Sternum, external spine in dorsal or ventral view (Fig. S16; Mayr, 2003: 4; Ksepka et al., 2013: 52; Ksepka et al., 2019: 38):

0 – single

1 – bifurcated

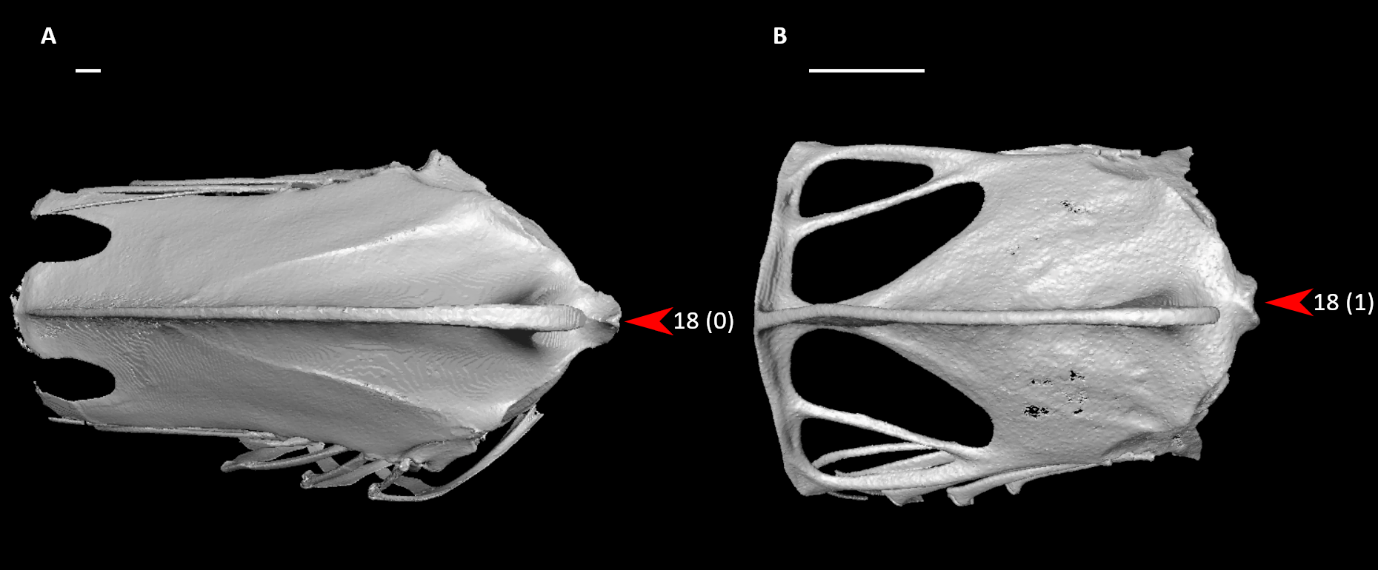

**Figure S16.** Sterna of *Phoenicopterus ruber* (**A**) and *Aegotheles cristatus* (**B**) in ventral view, illustrating alternative states for character 18 (complete character description in text). Arrows indicate the tip of the external spine. Scale bars = 5 mm.

19. Sternum, external spine, tongue-shaped ventral projection (Fig. S17; Ksepka et al., 2019: 40):

0 – absent

1 – present

Scored inapplicable in *Psilopogon* due to fusion with the sternal keel.

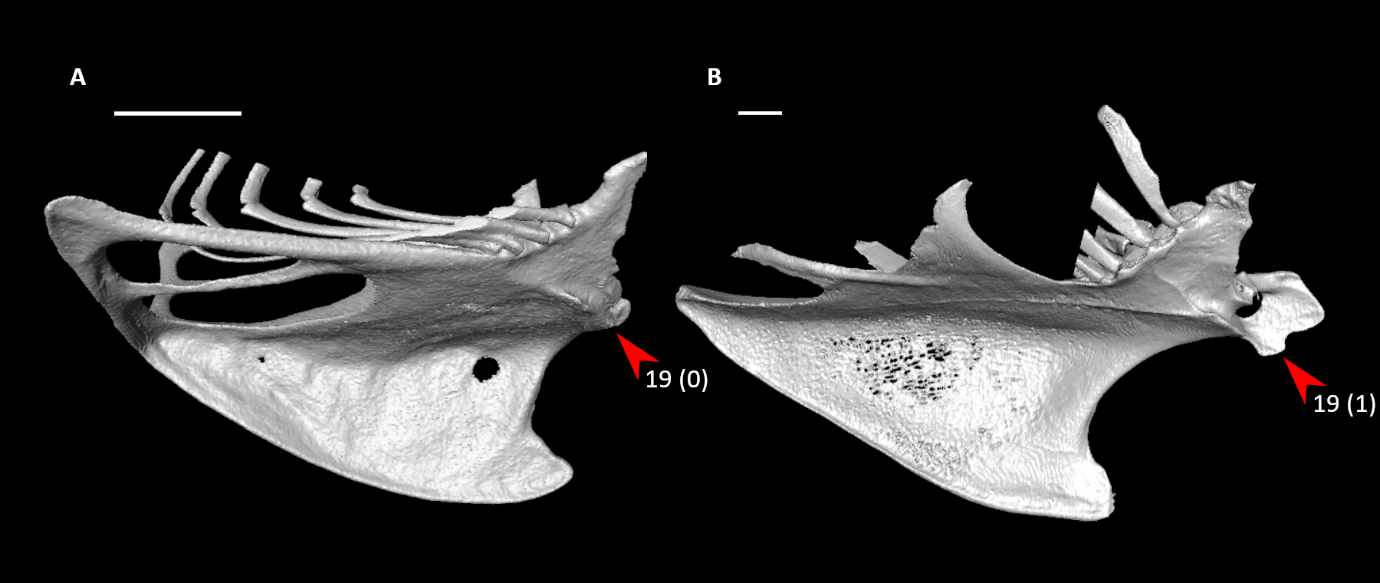

**Figure S17.** Sterna of *Aegotheles cristatus* (**A**) and *Ortalis ruficauda* (**B**) in right (**A**) or left (**B**, mirrored) lateral view, illustrating alternative states for character 19 (complete character description in text). Arrows indicate the ventral margin of the external spine. Scale bars = 5 mm.

20. Sternum, internal spine (Fig. S18; Livezey, 1986: 82; Livezey, 1996: 38; Worthy and Lee, 2008: 33; Ksepka, 2009: 33; Worthy et al., 2017: 83; Ksepka et al., 2019: 36; Musser and Clarke, 2020: 299):

0 – absent

1 – present

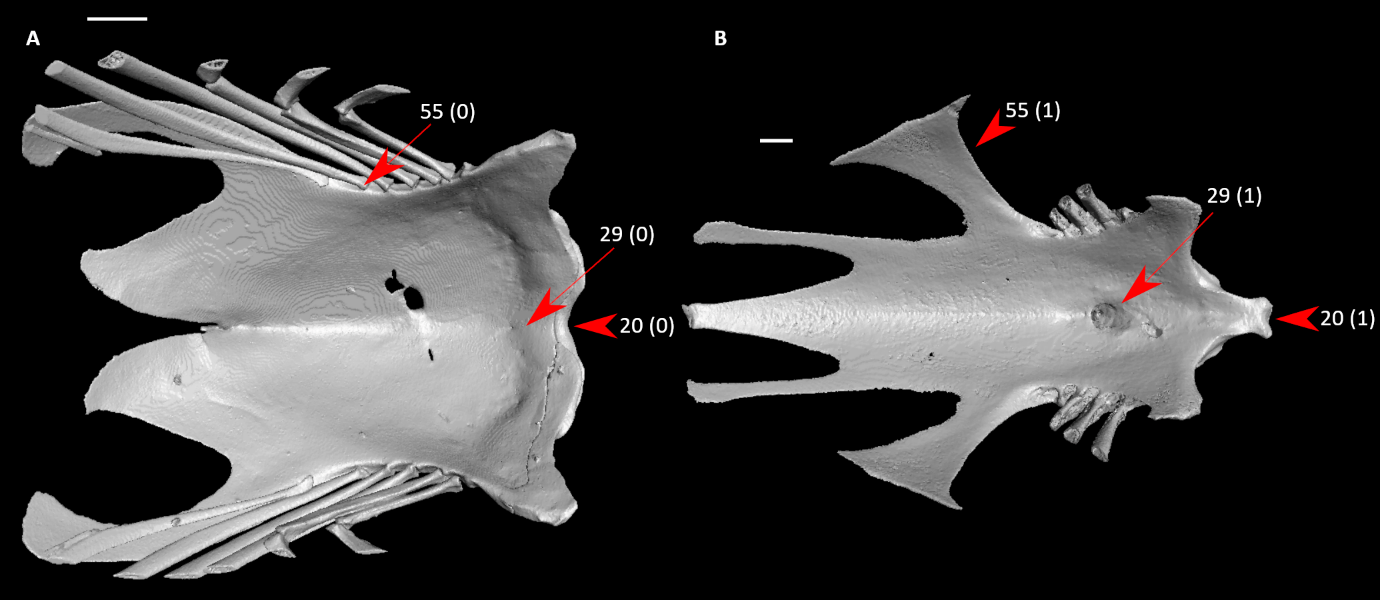

**Figure S18.** Sterna of *Podilymbus podiceps* (**A**) and *Alectura lathami* (**B**) in dorsal view, illustrating alternative states for characters 20, 29, and 55 (complete character descriptions in text). Arrows for character 20 indicate the internal spine or approximate homologous site; arrows for character 29 indicate the pneumatic foramen in the median sulcus immediately caudal to the cranial margin, or approximate homologous site to the foramen; and arrows for character 55 indicate the lateral trabecula or approximate homologous site. Scale bars = 5 mm.

21. Sternum, ossified connection between external and internal spines (Fig. S19; Worthy et al., 2017: 85; Musser and Clarke, 2020: 300):

0 – absent

1 – present

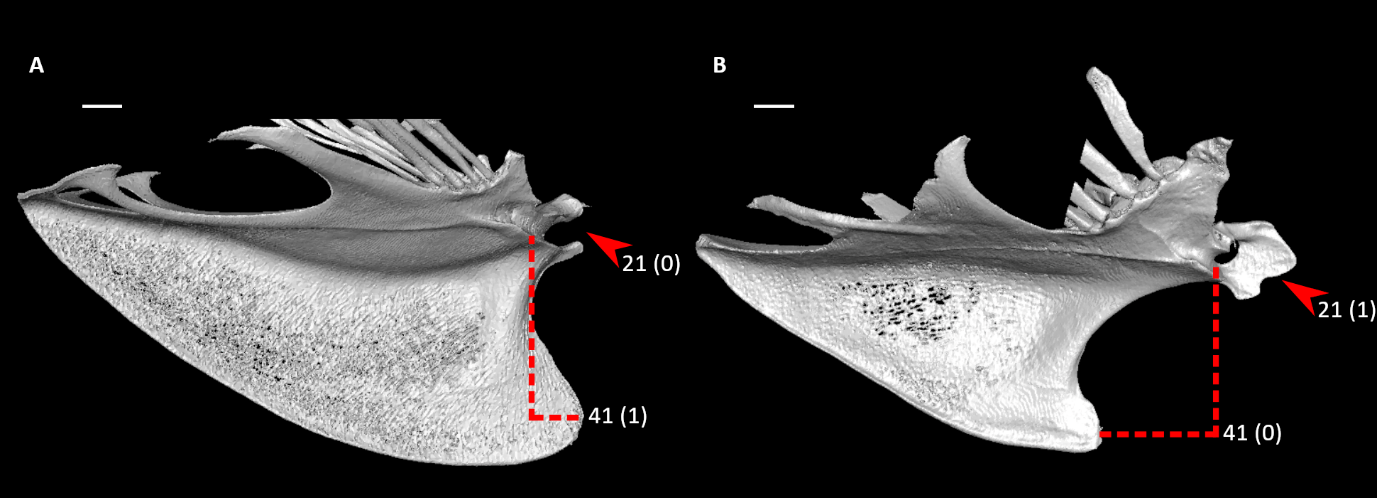

**Figure S19.** Sterna of *Columba livia* (**A**) and *Alectura lathami* (**B**) in right (**A**) or left (**B**, mirrored) lateral view, illustrating alternative states for characters 21 and 41 (complete character descriptions in text). Arrows indicate the region between the internal and external spines. Dotted lines indicate the cranial extent of the keel apex relative to that of the main body. Scale bars = 5 mm.

22. Sternum, cranial margin, coracoid sulcus, dorsal lip, pair of cranially extending flanges on cranial margin (Fig. S20; Musser and Cracraft, 2019: 161; Musser and Clarke, 2020: 348):

0 – absent

1 – present

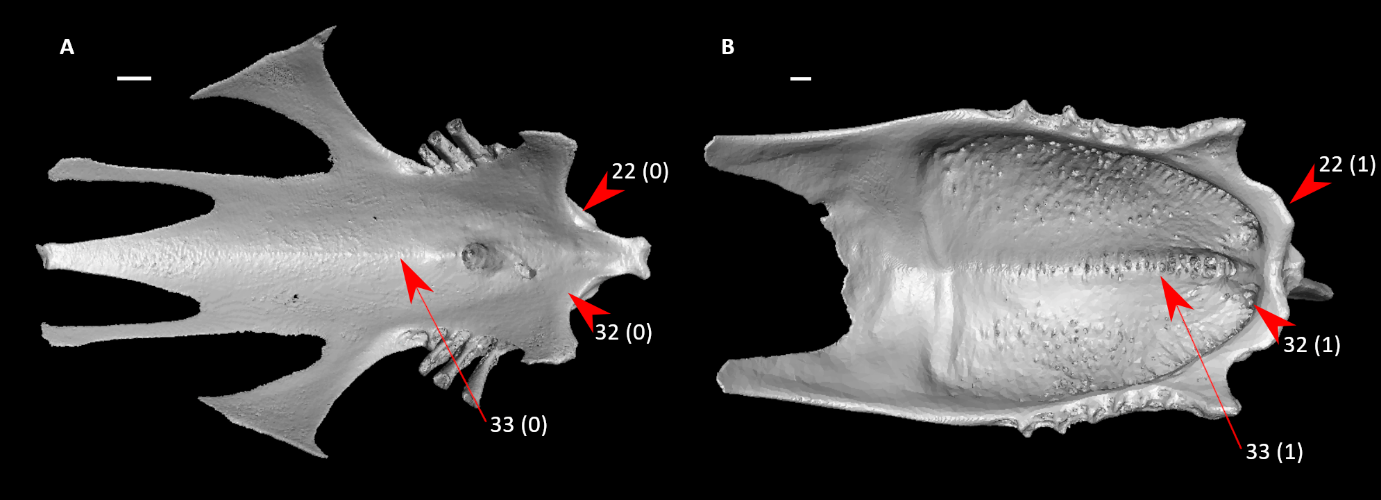

**Figure S20.** Sterna of *Alectura lathami* (**A**) and *Chauna chavaria* (**B**) in dorsal view, illustrating alternative states for characters 22, 32, and 33 (complete character descriptions in text). Arrows for character 22 indicate the cranial margin of the sternum, arrows for character 32 indicate the region immediately caudal to the cranial margin and its pneumatic pores (when present), and arrows for character 33 indicate the median sulcus and its pneumatic pores (when present). Scale bars = 5 mm.

23. Sternum, cranial margin, coracoid sulcus, ventral lip, external labial tubercle (Fig. S21):

0 – absent or weak

1 – prominent, forming sharply pointed projection

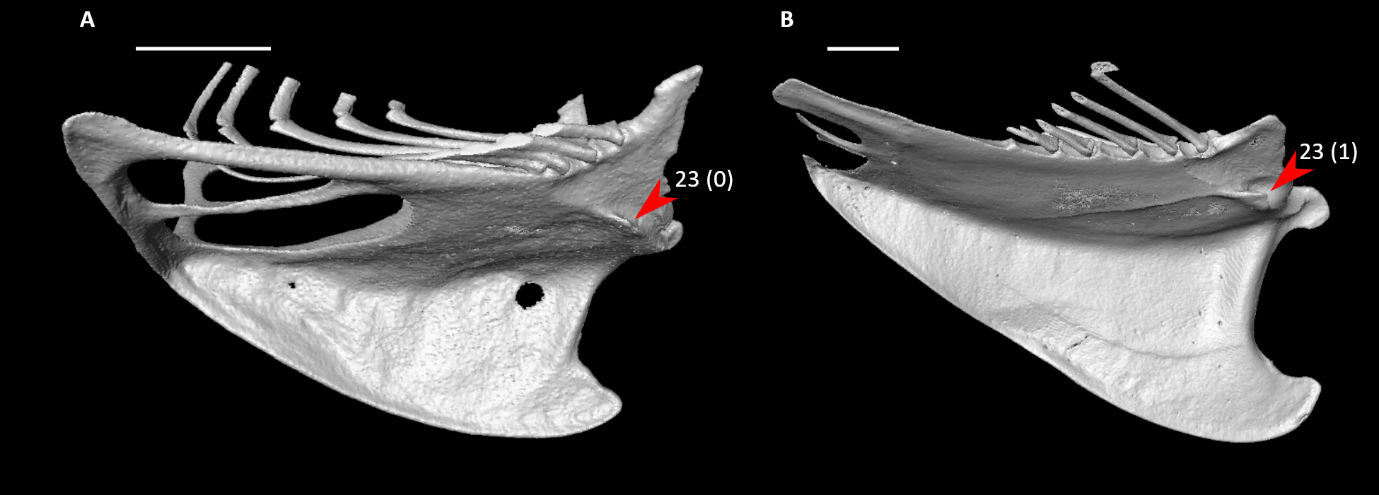

**Figure S21.** Sterna of *Aegotheles cristatus* (**A**) and *Sterna hirundo* (**B**) in right lateral view, illustrating alternative states for character 23 (complete character description in text). Arrows indicate the external labial tubercle. Scale bars = 5 mm.

*24. Sternum, cranial margin, coracoid sulcus, ventral lip, angle of lateral portion relative to midline of sternum in ventral view (Fig. S22; Livezey and Zusi, 2006: 1128; Smith, 2010: 116):

0 – angle extremely low, ventral lip and midline nearly parallel to each other

1 – angle intermediate, greater than 15 degrees, less than 60 degrees

2 – angle extremely high, approaching 90 degrees

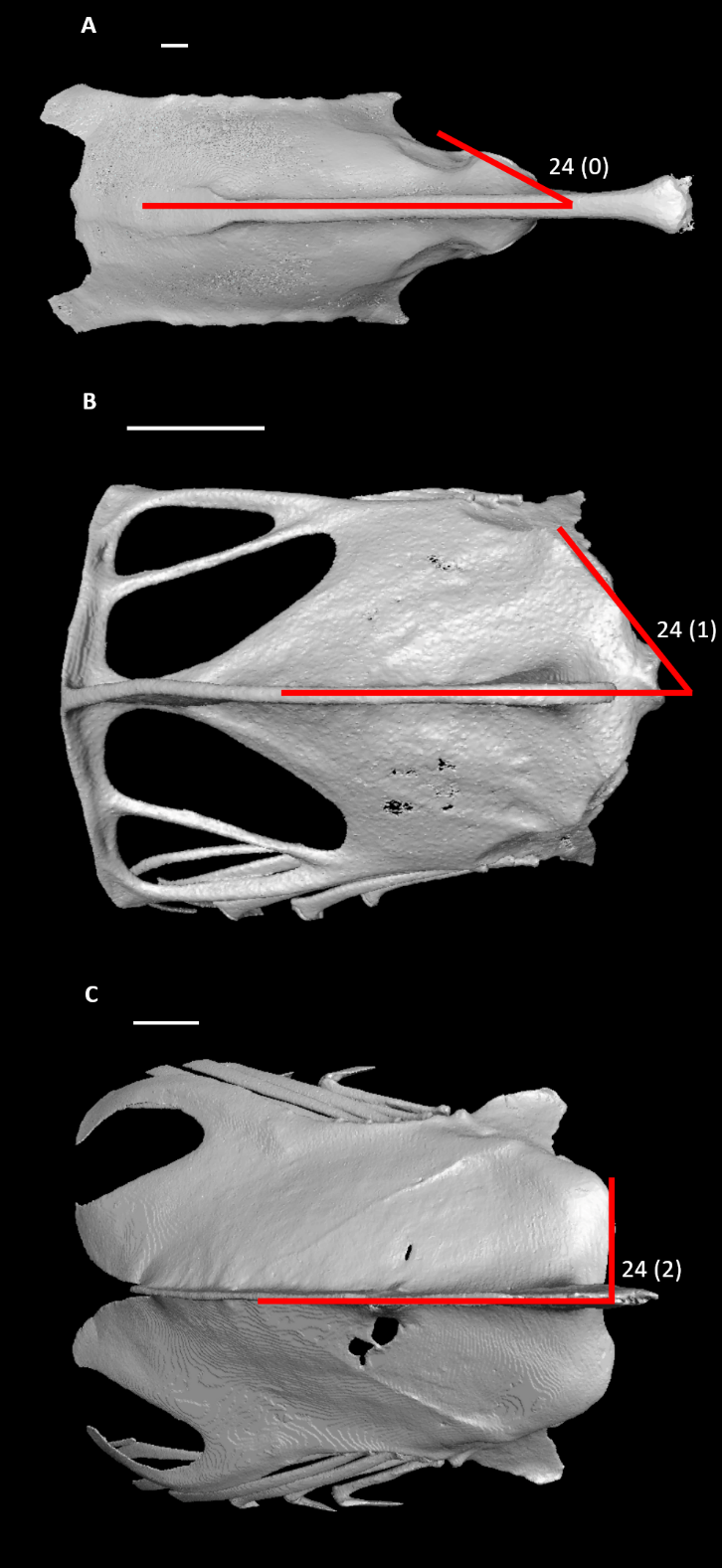

**Figure S22.** Sterna of *Sula dactylatra* (**A**) *Aegotheles cristatus* (**B**), and *Podilymbus podiceps* (**C**) in ventral view, illustrating alternative states for character 24 (complete character description in text). Red lines indicate the angle between the ventral lip and midline of the sternum. Scale bars = 5 mm.

25. Sternum, cranial margin, coracoid sulci extent across midline (Fig. 23; Musser et al., 2019: 53; Musser and Clarke, 2020: 309):

0 – sulci do not overlap over midline

1 – sulci extend over midline and cross

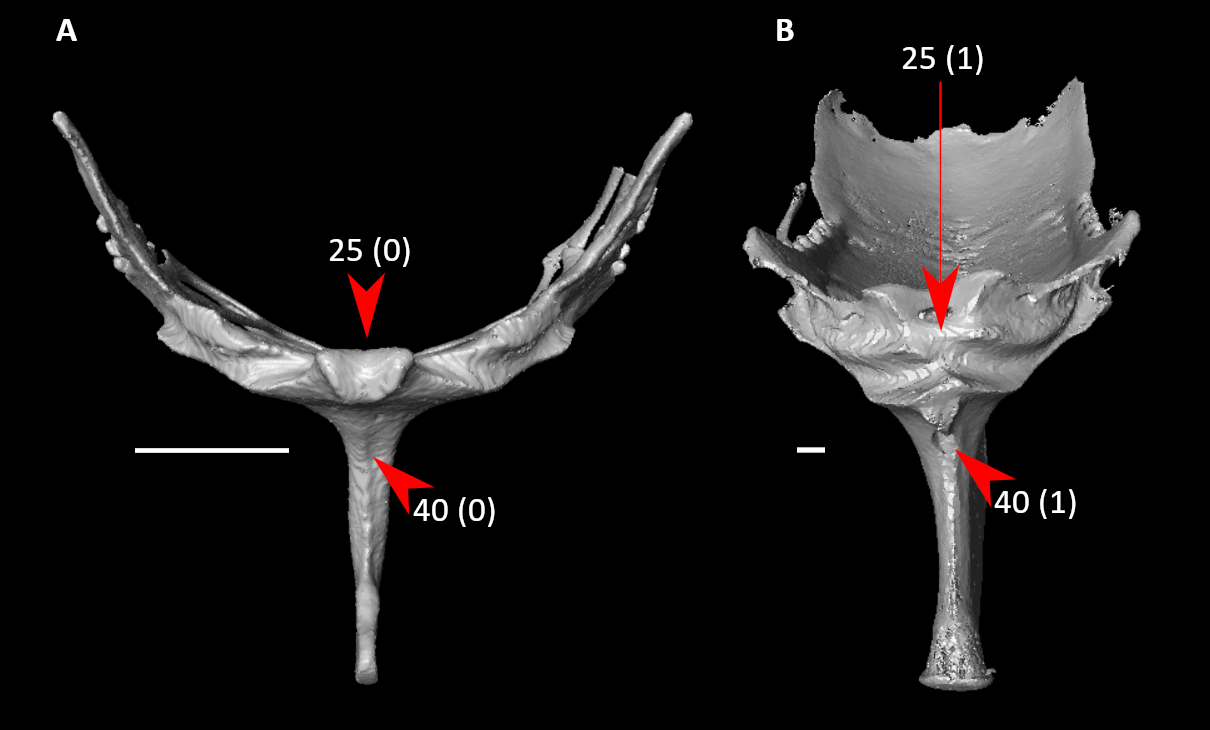

**Figure S23.** Sterna of *Aegotheles cristatus* (**A**) and *Balearica pavonina* (**B**) in cranial view, illustrating alternative states for characters 25 and 40 (complete character descriptions in text). Arrows for character 25 indicate the midline of the cranial margin of the sternum and overlap of coracoid sulci (when present); whereas arrows for character 40 indicate the pneumatic foramen on the keel, or approximate homologous site to the foramen. Scale bars = 5 mm.

26. Sternum, cranial margin, coracoid sulci, pneumatic foramen (Fig. 24):

0 – absent

1 – present

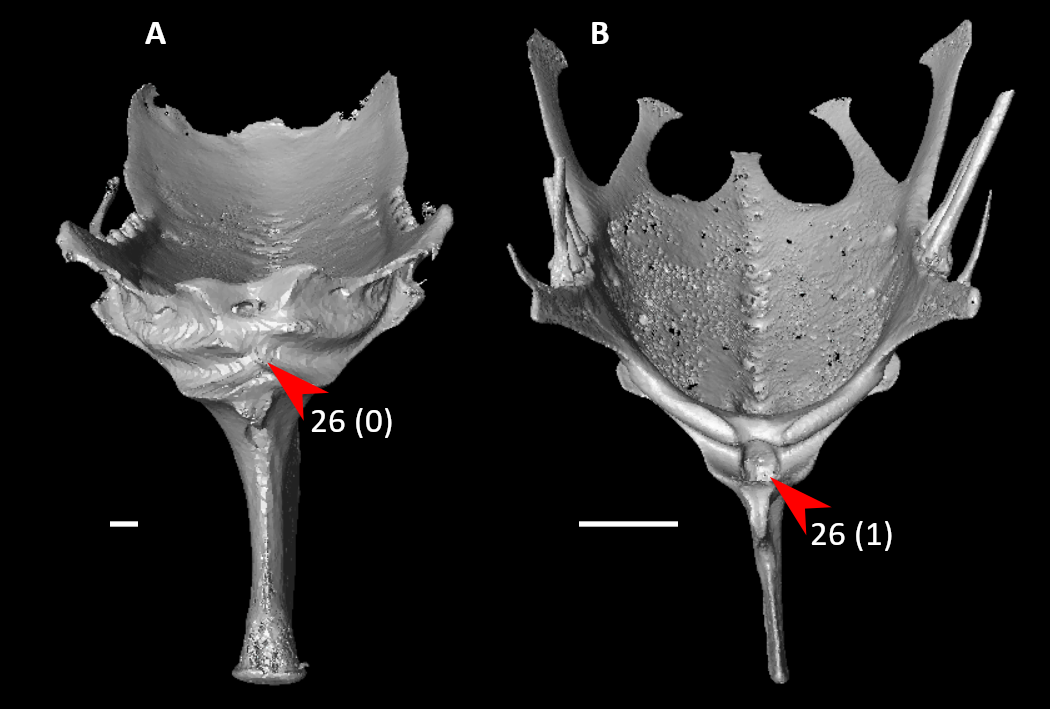

**Figure S24.** Sterna of *Balearica pavonina* (**A**) and *Coracias benghalensis* (**B**) in cranial view, illustrating alternative states for character 26 (complete character description in text). Arrows indicate the pneumatic foramen between the coracoid sulci, or approximate homologous site to the foramen. Scale bars = 5 mm.

27. Sternum, cranial margin, angle of craniolateral process (Fig. S25; Livezey and Zusi, 2006: 1149; Musser and Cracraft, 2019: 165; Musser and Clarke, 2020: 311):

0 – extending laterally

1 – extending cranially

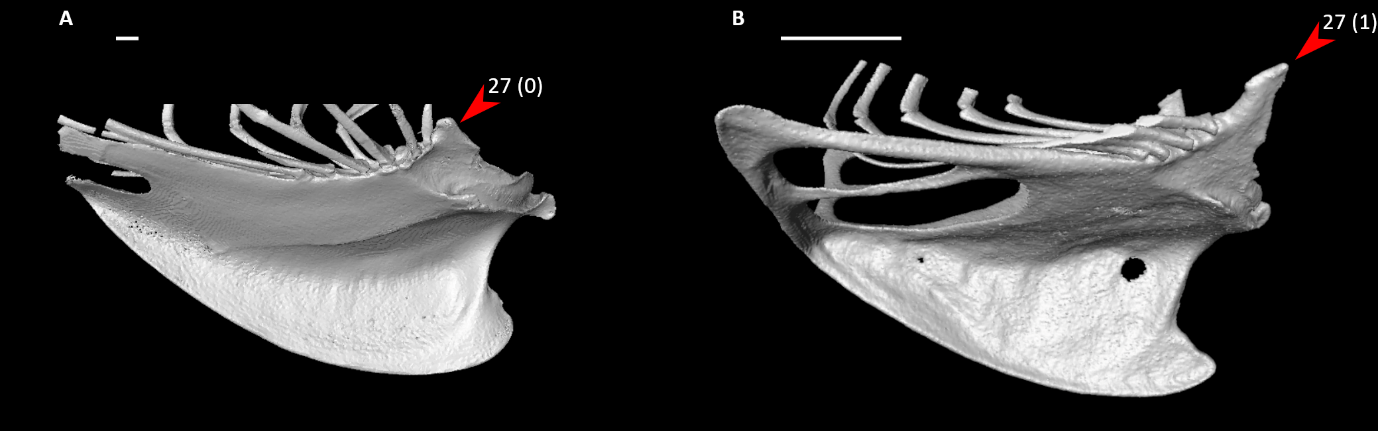

**Figure S25.** Sterna of *Phoenicopterus ruber* (**A**) and *Aegotheles cristatus* (**B**) in right lateral view, illustrating alternative states for character 27 (complete character description in text). Arrows indicate the cranial process. Scale bars = 5 mm.

28. Sternum, dorsal surface, median sulcus (Fig. S26; Livezey and Zusi, 2006: 1111; Musser and Cracraft, 2019: 170):

0 – absent

1 – present

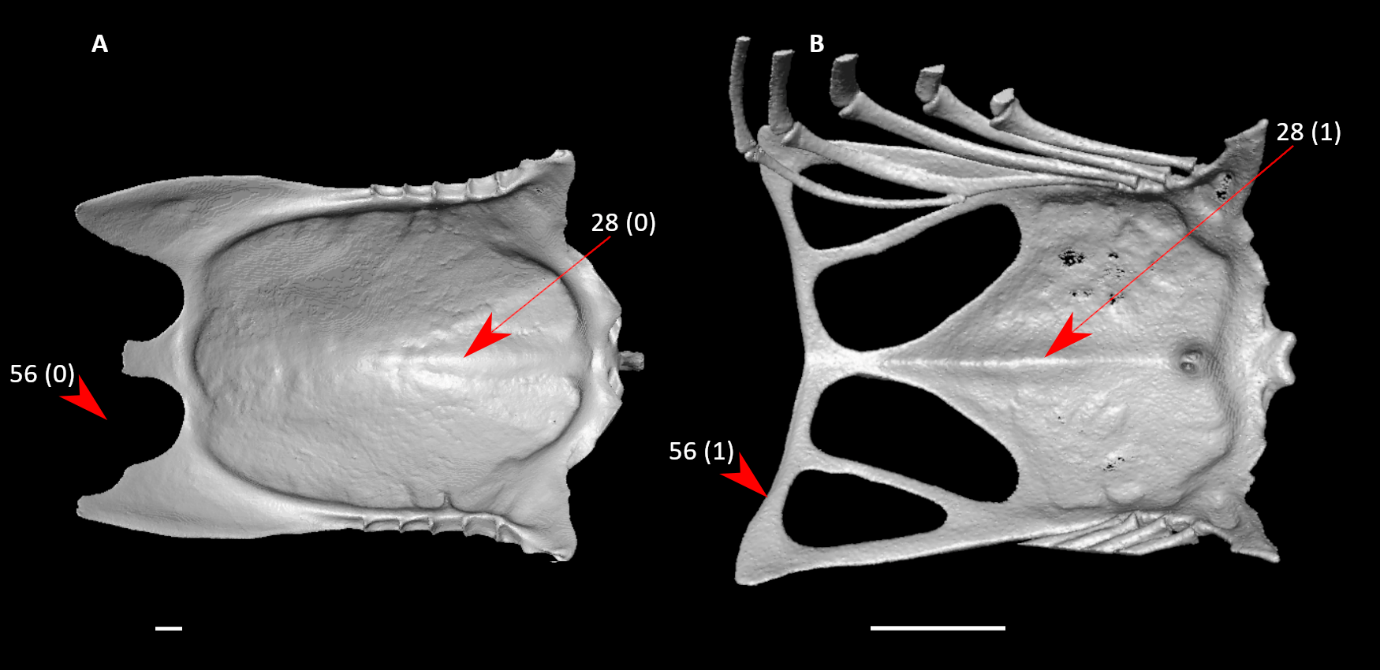

**Figure S26.** Sterna of *Leucocarbo atriceps* (**A**) and *Aegotheles cristatus* (**B**) in dorsal view, illustrating alternative states for characters 28 and 56 (complete character descriptions in text). Arrows for character 28 indicate the median sulcus or approximate homologous site, whereas arrows for character 56 indicate the caudal margin of the lateral incisure or fenestra. Scale bars = 5 mm.

29. Sternum, dorsal surface, pneumatic foramen in median sulcus immediately caudal to cranial margin, exclusive of pneumatic pores (Fig. S18):

0 – absent

1 – present

30. Sternum, dorsal surface, bony lamina immediately caudal to cranial margin (Fig. 27; Livezey and Zusi, 2006: 1108; Musser and Cracraft, 2019: 167; Musser and Clarke, 2020: 315):

0 – absent

1 – single lamina present

2 – two laminae present

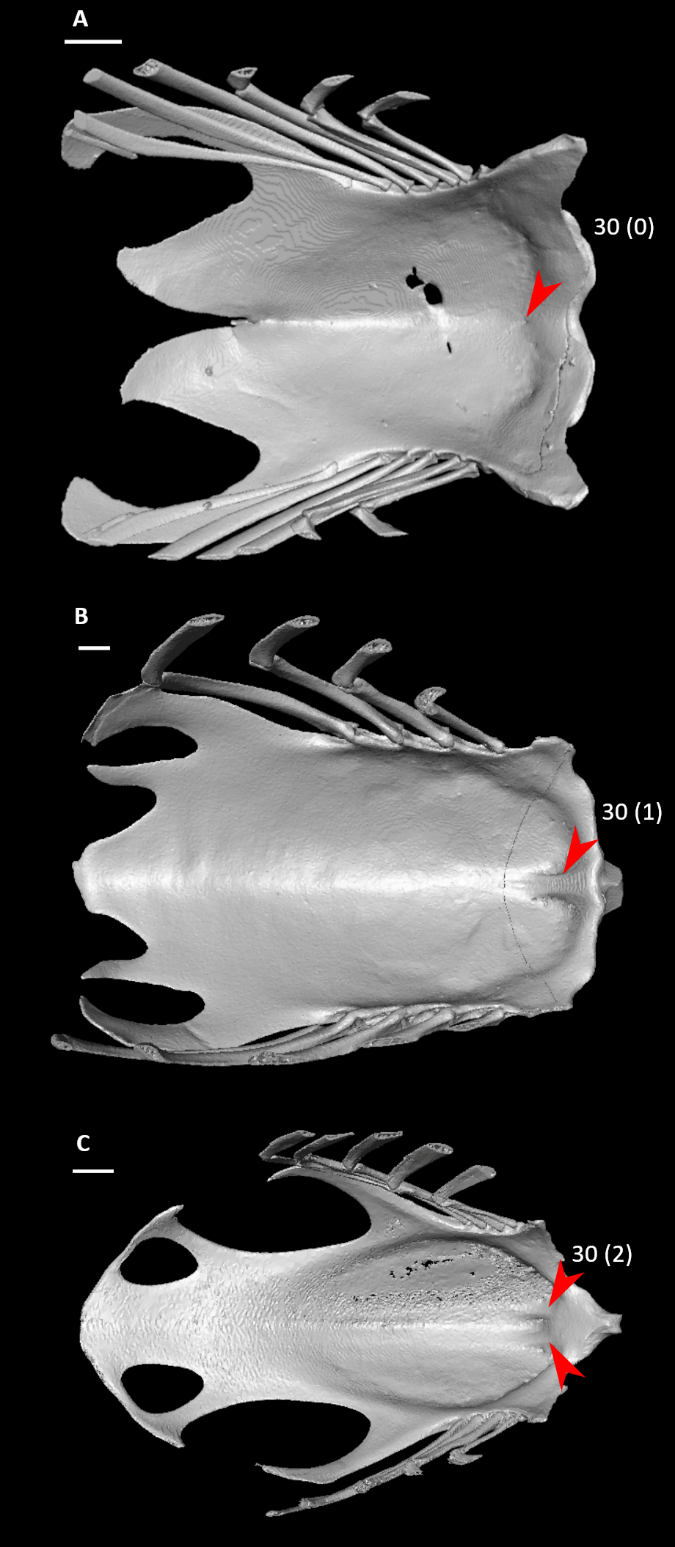

**Figure S27.** Sterna of *Podilymbus podiceps* (**A**), *Coragyps atratus* (**B**), and *Columba livia* (**C**) in dorsal view, illustrating alternative states for character 30 (complete character description in text). Arrows indicate the lamina(e) immediately caudal to the cranial margin, or approximate homologous site to the laminae. Scale bars = 5 mm.

31. Sternum, dorsal surface, site of medial sulcus and/or pneumatic foramen (Fig. S28; Livezey and Zusi, 2006: 1109; Musser and Cracraft, 2019: 168; Musser and Clarke, 2020: 316):

0 – beginning within margin of or immediately caudal to coracoid pillar

1 – significantly caudal to coracoid pillar, approximately caudal to base of keel

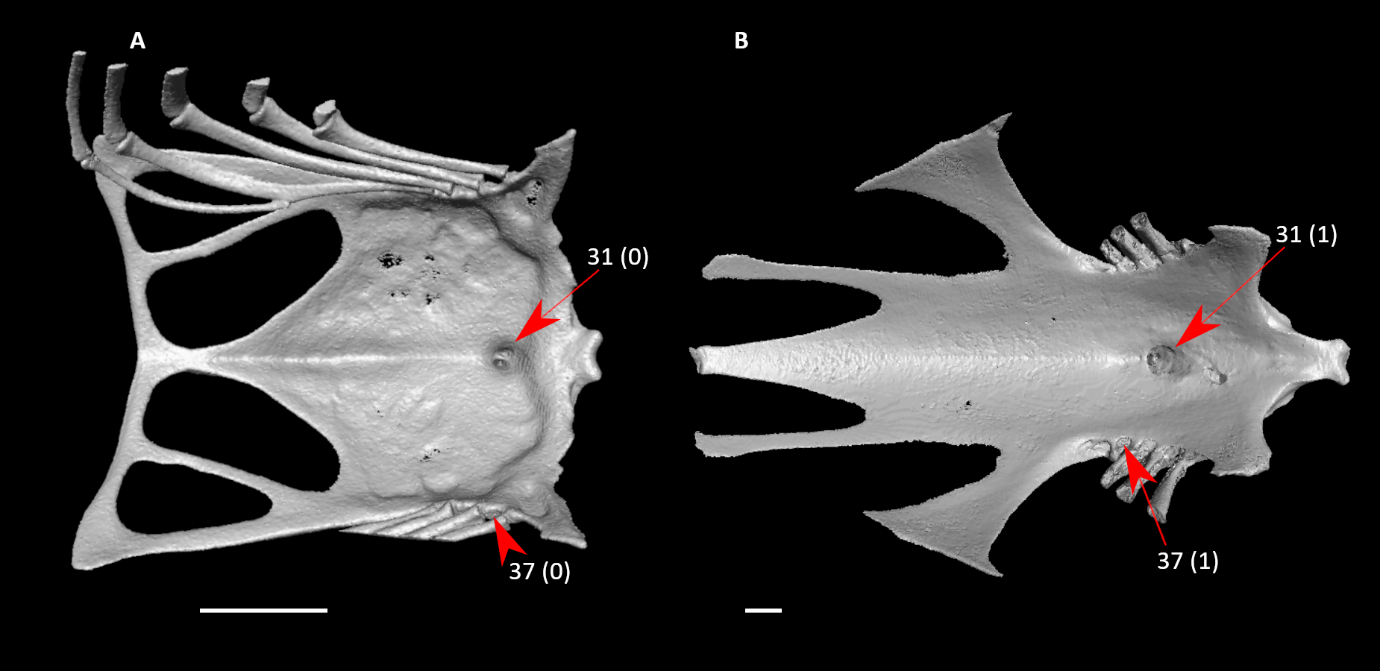

**Figure S28.** Sterna of *Aegotheles cristatus* (**A**) and *Alectura lathami* (**B**) in dorsal view, illustrating alternative states for characters 31 and 37 (complete character descriptions in text). Arrows for character 31 indicate the pneumatic foramen in the median sulcus immediately caudal to the cranial margin, whereas arrows for character 37 indicate intercostal incisure and its pneumatic pores (when present). Scale bars = 5 mm.

32. Sternum, dorsal surface, additional pneumatic pores, exclusive of those included within pneumatic foramen, cranial margin (Fig. S20):

0 – absent

1 – present

33. Sternum, dorsal surface, additional pneumatic pores, exclusive of those included within pneumatic foramen, median sulcus (Fig. S20; Livezey and Zusi, 2006: 1110; Musser and Cracraft, 2019: 169; Musser and Clarke, 2020: 317):

0 – absent

1 – present

34. Sternum, ventrolateral sulcus (longitudinal trough on ventral surface immediately medial to costal processes) (Fig. S29; Livezey, 1998: 153; Musser and Cracraft, 2019: 171; Musser and Clarke, 2020: 318):

0 – absent or so shallow as to be indistinct

1 – distinct, typically for length of costal margin

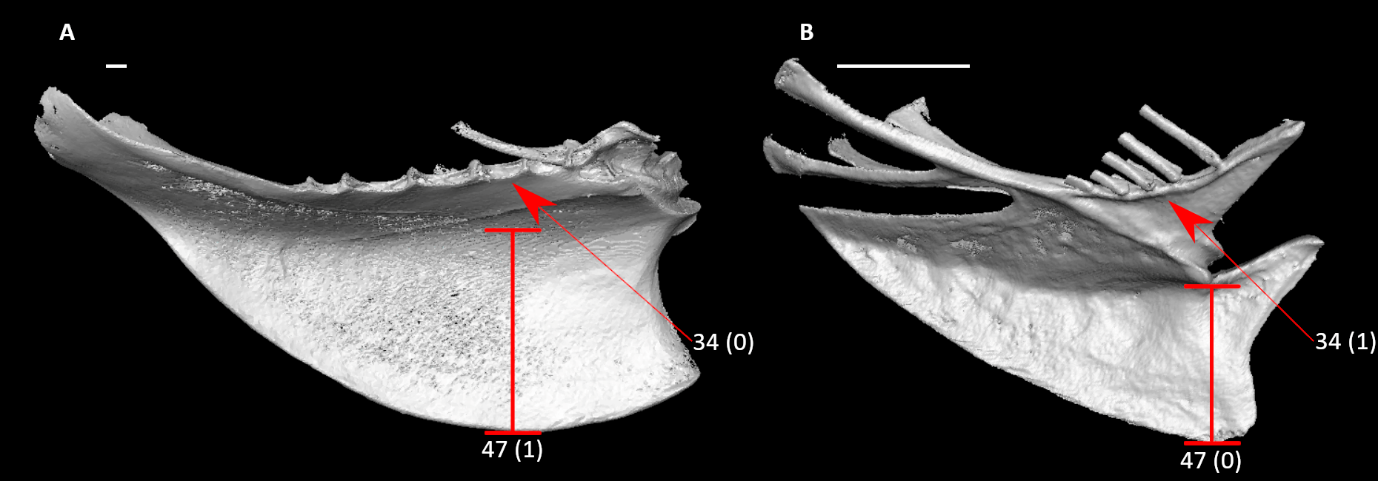

**Figure S29.** Sterna of *Balearica pavonina* (**A**) and *Jynx torquilla* (**B**) in right (**A**) or left (**B**, mirrored) lateral view, illustrating alternative states for characters 34 and 47 (complete character descriptions in text). Arrows indicate the ventrolateral sulcus. Red lines span the depth of the keel. Scale bars = 5 mm.

*35. Sternum, costal margin, craniocaudal length relative to that of entire sternum along medial axis (Fig. S30; Livezey and Zusi, 2006: 1114; Musser and Cracraft, 2019: 172; Musser and Clarke, 2020: 319):

0 – less than 25% total sternal length

1 – between 25-75% total sternal length

2 – greater than 75% total sternal length

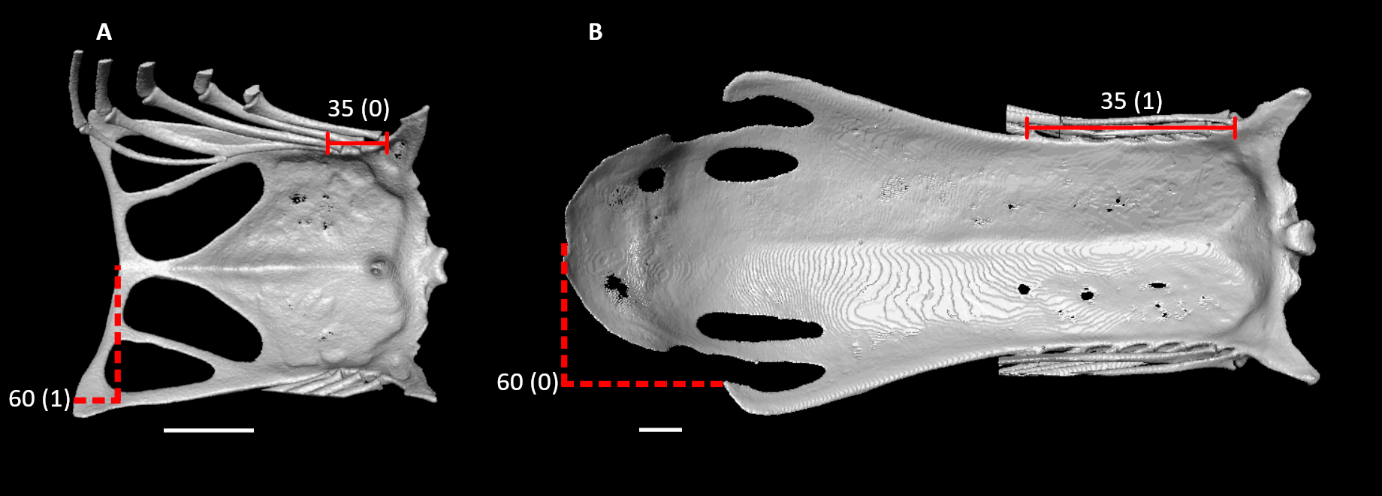

**Figure S30.** Sterna of *Aegotheles cristatus* (**A**) and *Alca torda* (**B**) in dorsal view, illustrating alternative states for characters 35 and 60 (complete character descriptions in text). Solid red lines span the length of the costal margin. Dotted lines indicate the caudal extent of the median trabecula relative to that of the caudolateral trabecula. Scale bars = 5 mm.

*36. Sternum, costal margin, number of articular facets for sternal ribs (Fig. S31; Mayr and Clarke, 2003: 71; Musser and Cracraft, 2019: 173; Musser and Clarke, 2020: 320):

0 – three

1 – four

2 – five

3 – six

4 – seven

5 – eight

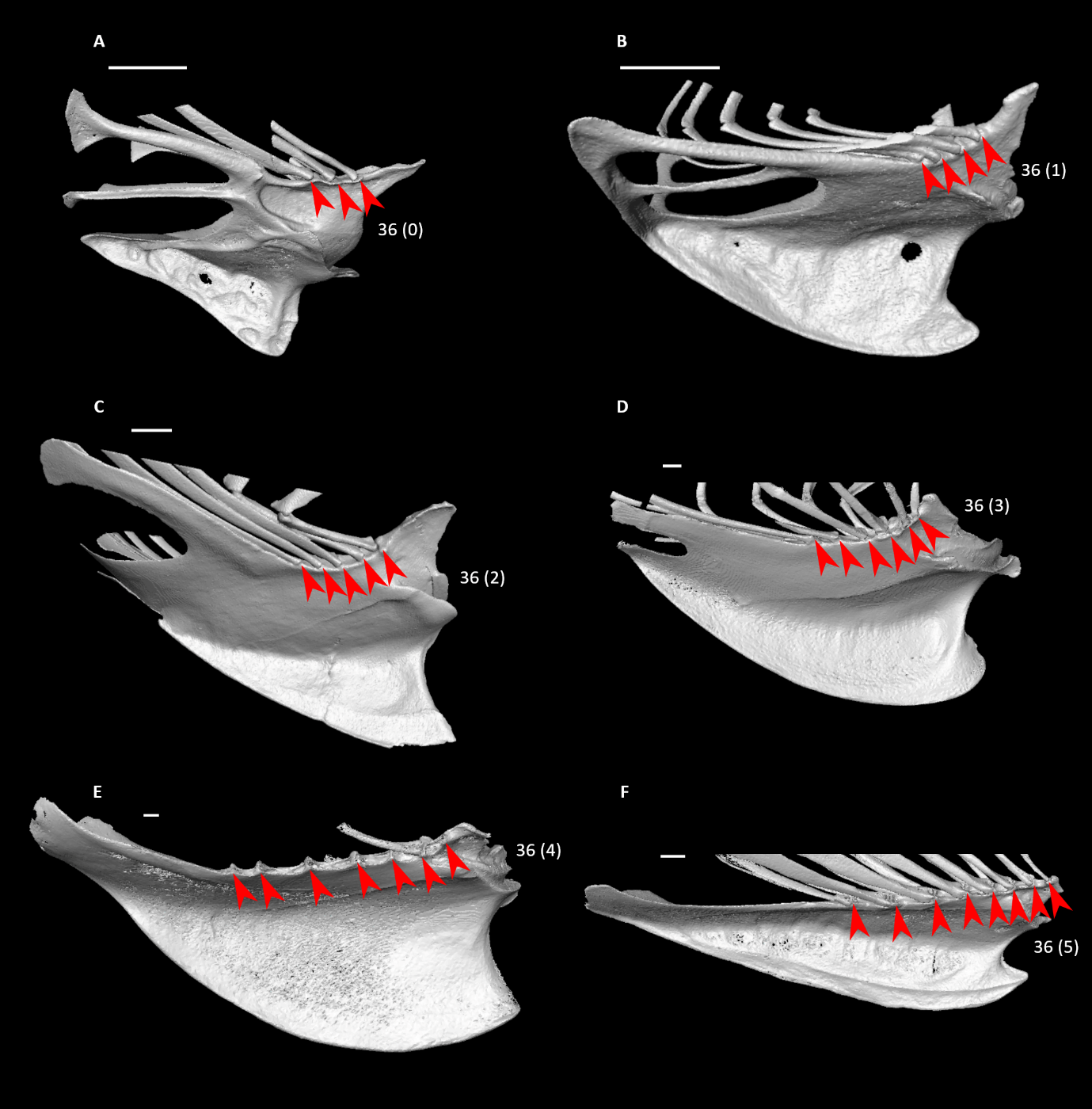

**Figure S31.** Sterna of *Bucco capensis* (**A**), *Aegotheles cristatus* (**B**), *Podilymbus podiceps* (**C**), *Phoenicopterus ruber* (**D**), *Balearica pavonina* (**E**), and *Psophia crepitans* (**F**) in right (**A, B, D, E, F**) or left (**C**, mirrored) lateral view, illustrating alternative states for character 36 (complete character description in text). Arrows indicate articular facets for sternal ribs. Scale bars = 5 mm.

37. Sternum, costal margin, intercostal incisures, pneumatic pores (Fig. S28; Livezey and Zusi, 2006: 1115; Musser and Cracraft, 2019: 174):

0 – absent

1 – present

38. Sternum, keel:

0 – absent or weak

1 – prominent

39. Sternum, keel, cranial margin, keel sulcus (Fig. S32; Livezey and Zusi, 2006: 1213; Musser and Cracraft, 2019: 177; Musser and Clarke, 2020: 326):

0 – absent

1 – present

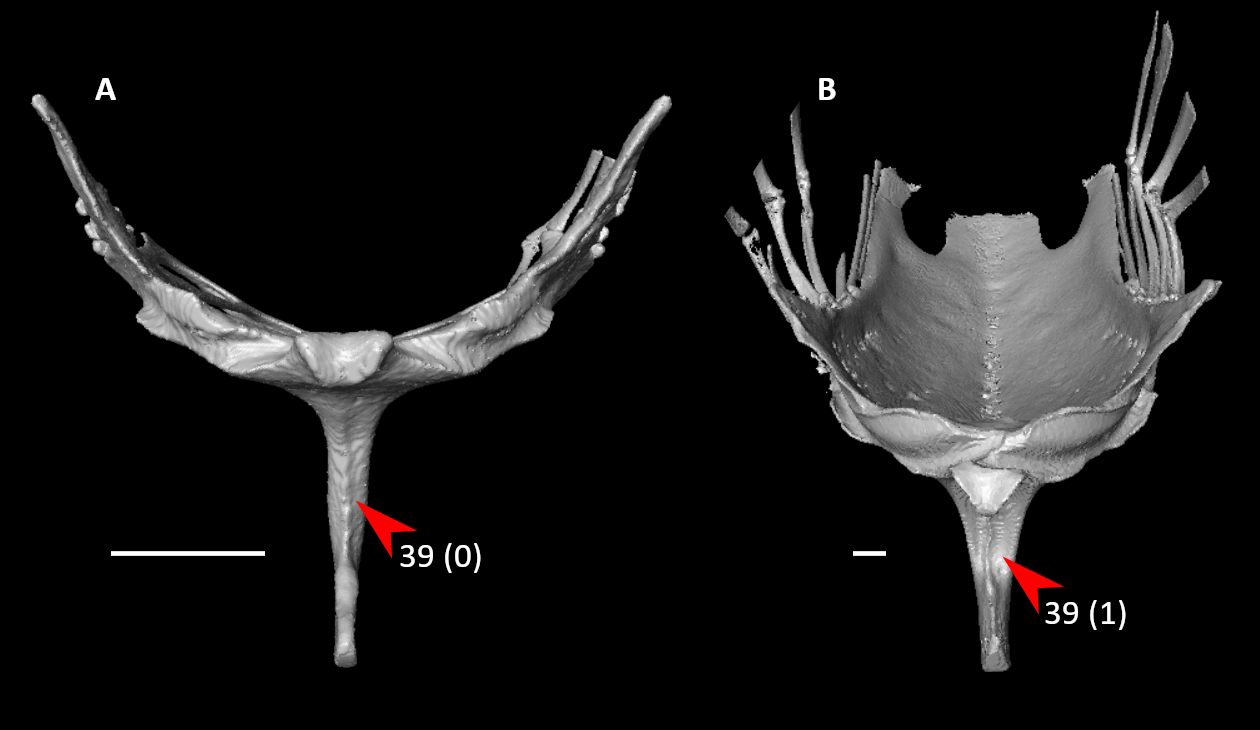

**Figure S32.** Sterna of *Aegotheles cristatus* (**A**) and *Phoenicopterus ruber* (**B**) in cranial view, illustrating alternative states for character 39 (complete character description in text). Arrows indicate the keel sulcus or approximate homologous site. Scale bars = 5 mm.

40. Sternum, keel, cranial margin, keel sulcus or homologous site, pneumatic foramen (Fig. S23; Livezey and Zusi, 2006: 1218; Musser and Cracraft, 2019: 178; Musser and Clarke, 2020: 327):

0 – absent

1 – present

*41. Sternum, keel, site of apex relative to cranial end of sternal main body (excluding external spine) (Fig. S19; Livezey and Zusi, 2006: 1198; Musser and Cracraft, 2019: 179; Musser and Clarke, 2020: 329):

0 – caudal to sternal main body

1 – cranial to sternal main body

42. Sternum, keel, recurved cranial margin (Fig. S33; Musser and Clarke, 2020: 335):

0 – absent

1 – present

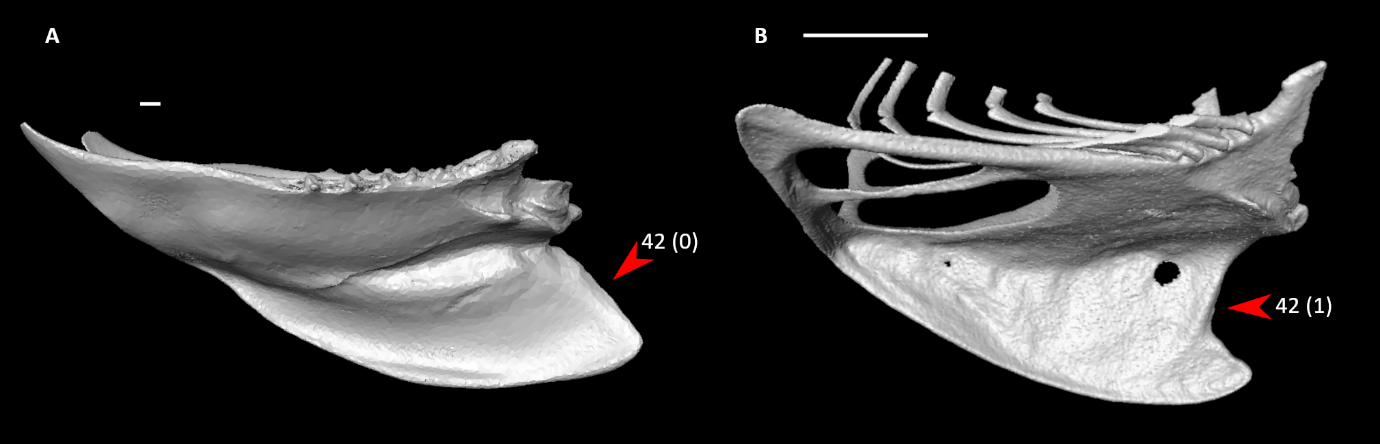

**Figure S33.** Sterna of *Chauna chavaria* (**A**) and *Aegotheles cristatus* (**B**) in right lateral view, illustrating alternative states for character 42 (complete character description in text). Arrows indicate the cranial margin of the sternal keel. Scale bars = 5 mm.

43. Sternum, keel, apex shape in lateral view (Fig. S34; Musser and Clarke, 2020: 334):

0 – rounded

1 – pointed

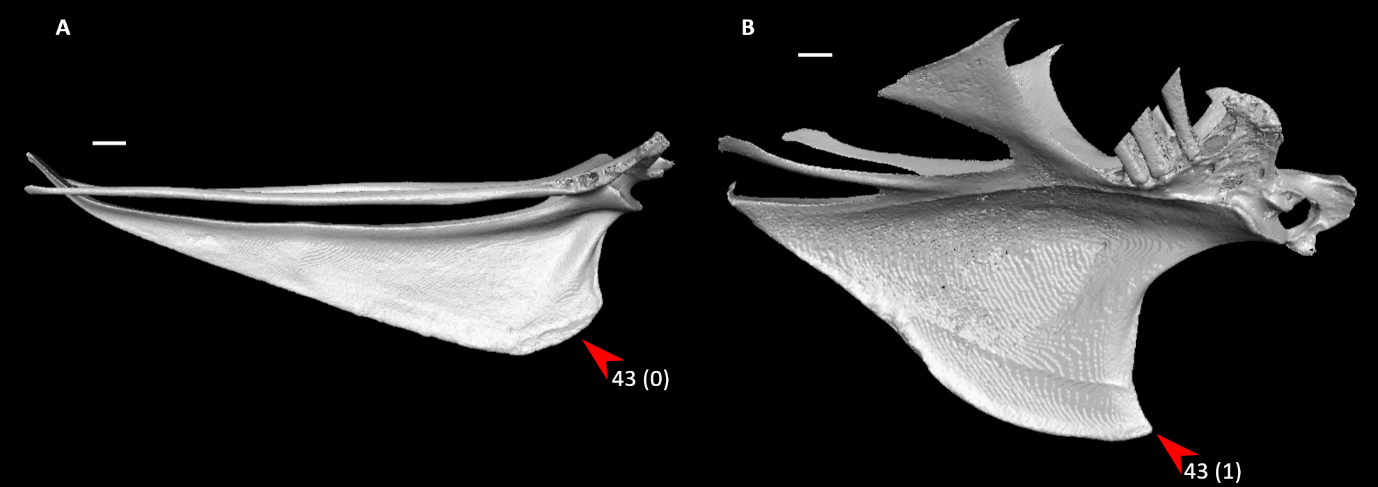

**Figure S34.** Sterna of *Eudromia elegans* (**A**) and *Alectura lathami* (**B**) in right (**A**) or left (**B**, mirrored) lateral view, illustrating alternative states for character 43 (complete character description in text). Arrows indicate the keel apex. Scale bars = 5 mm.

44. Sternum, keel, articular facet for furcula (Fig. S35; Livezey and Zusi, 2006: 1200; Musser and Cracraft, 2019: 180; Musser and Clarke, 2020: 330):

0 – absent, furcula does not articulate directly with sternum

1 – present, incisure or concavity

2 – keel fused to furcula with no visible articular facet

Scored as unknown for *Ichthyornis*. Scored as inapplicable in *Opisthocomus*, in which the furcula is fused to the sternum but not to the sternal keel.

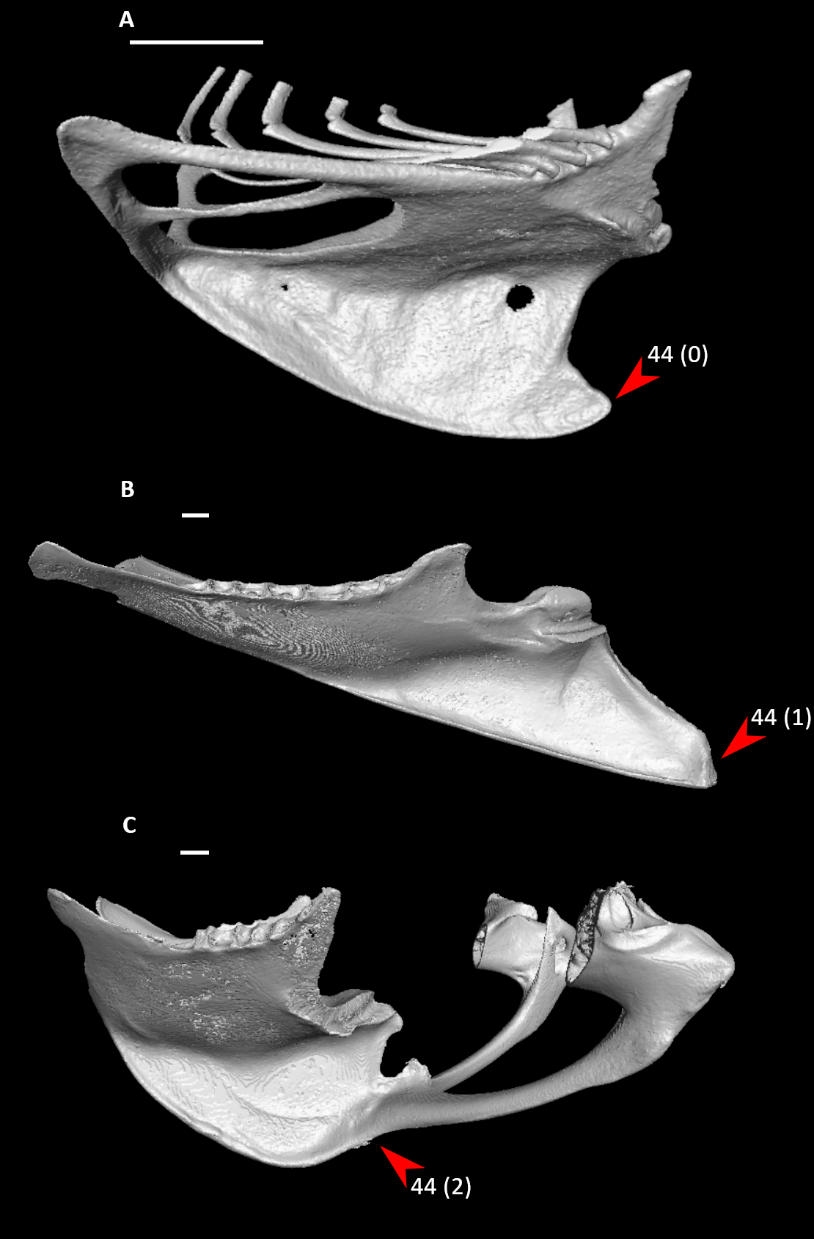

**Figure S35.** Sterna of *Aegotheles cristatus* (**A**), *Sula dactylatra* (**B**), and *Fregata aquila* (**C**) in right (**A, B**) or left (**C**, mirrored) lateral view, illustrating alternative states for character 44 (complete character description in text). Arrows indicate the keel apex and its articulation point (when present) with the furcula. Scale bars = 5 mm.

45. Sternum, keel, position of articular facet for furcula relative to apex of keel (Fig. S36; Livezey and Zusi, 2006: 1319; Smith, 2010: 137):

0 – approximately at apex

1 – caudal to apex

2 – cranial to apex

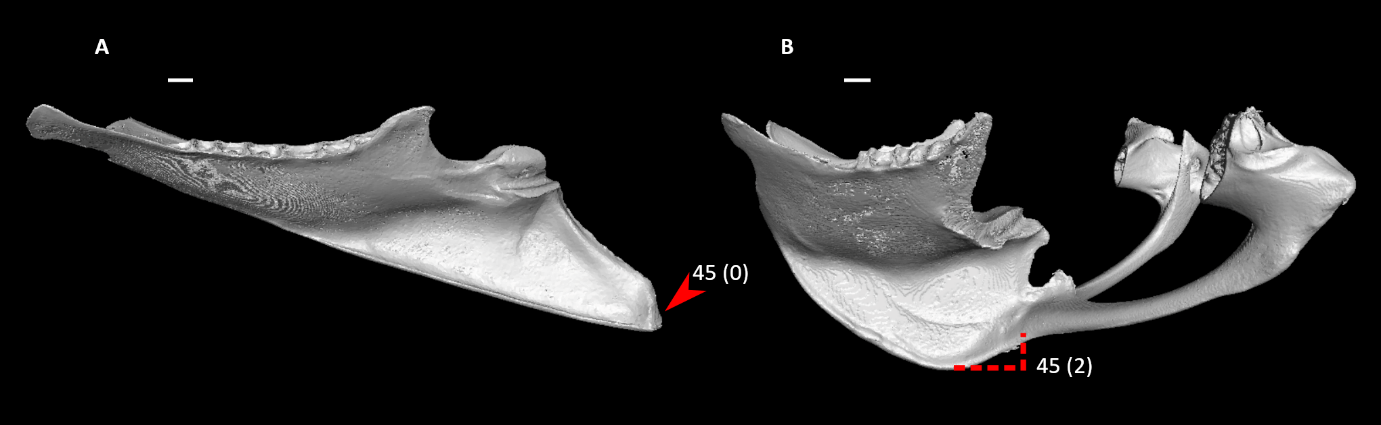

**Figure S36.** Sterna of *Sula dactylatra* (**A**) and *Fregata aquila* (**B**) in right (**A**) or left (**B**, mirrored) lateral view, illustrating alternative states for character 45 (complete character description in text). Arrow indicates the keel apex in *Sula*, whereas dotted lines indicate the position of the keel apex relative to the articulation with the furcula in *Fregata*. Scale bars = 5 mm.

46. Sternum, keel, cranial margin, degree of thickening in cranial view (Fig. S37; Musser and Cracraft, 2019: 181; Musser and Clarke, 2020: 332):

0 – keel becomes thinner ventrally

1 – keel maintains thickness ventrally (excluding articular facet for furcula)

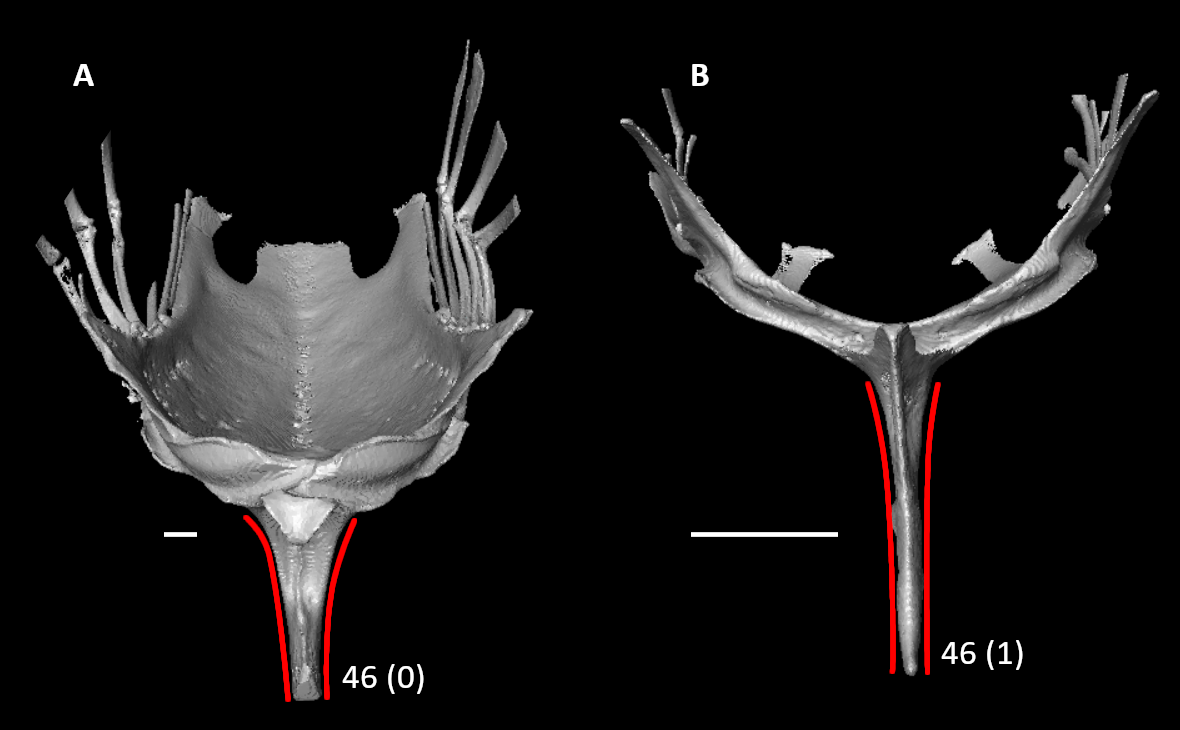

**Figure S37.** Sterna of *Phoenicopterus ruber* (**A**) and *Alcedo atthis* (**B**) in cranial view, illustrating alternative states for character 46 (complete character description in text). Red lines indicate the approximate lateral contours of the sternal keel. Scale bars = 5 mm.

47. Sternum, keel, depth of keel relative to total height of sternum, the latter as measured in a straight line from maximal depth of keel to the dorsal extent of the costal margin at its midpoint with the keel positioned vertically (Fig. S29; Livezey and Zusi, 2006: 1199; Musser and Cracraft, 2019: 182; Musser and Clarke, 2020: 333):

0 – keel low, at or less than two-thirds the total height of sternum

1 – keel prominent, greater than two-thirds the total height of sternum

48. Sternum, keel, caudal extent (Fig. S38; Mayr, 2011: 26):

0 – extends to termination of median trabecula

1 – ends distinctly before termination of median trabecula

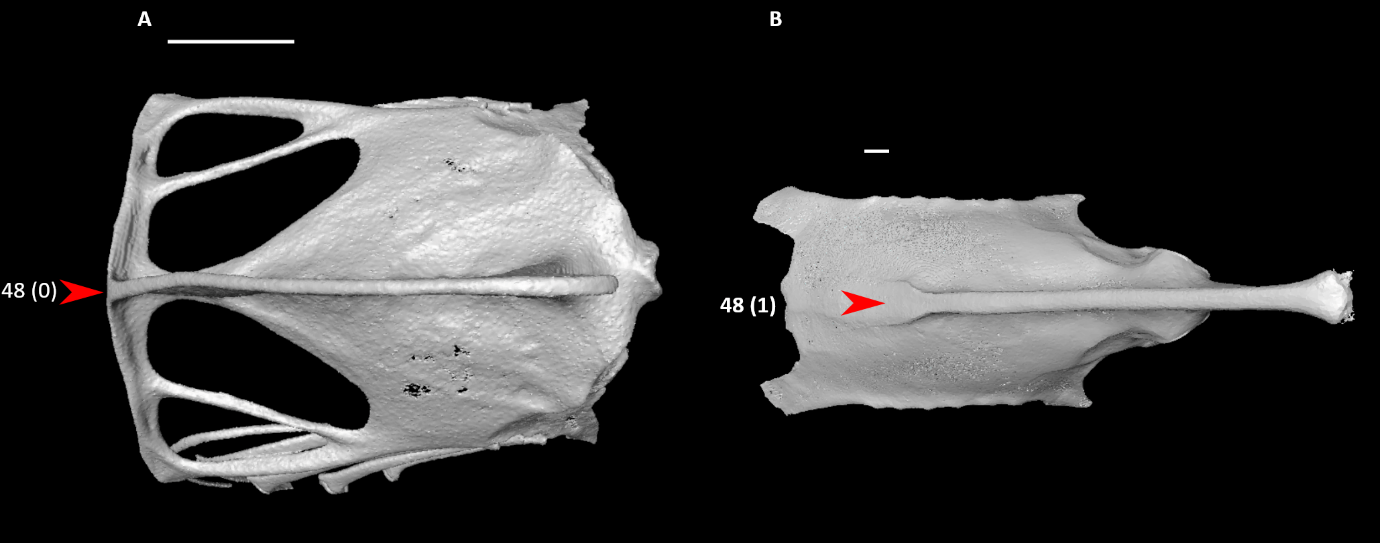

**Figure S38.** Sterna of *Aegotheles cristatus* (**A**) and *Sula dactylatra* (**B**) in ventral view, illustrating alternative states for character 48 (complete character description in text). Arrows indicate the caudal extent of the sternal keel. Scale bars = 5 mm.

49. Sternum, caudal margin, fenestrae and incisurae (Fig. S39; Mayr and Clarke, 2003: 73; Ksepka, 2009: 40; Worthy et al., 2017: 87; Musser and Clarke, 2020: 340):

0 – absent

1 – present as a single pair of fenestrae or incisurae

2 – present as two pairs of fenestrae or incisurae

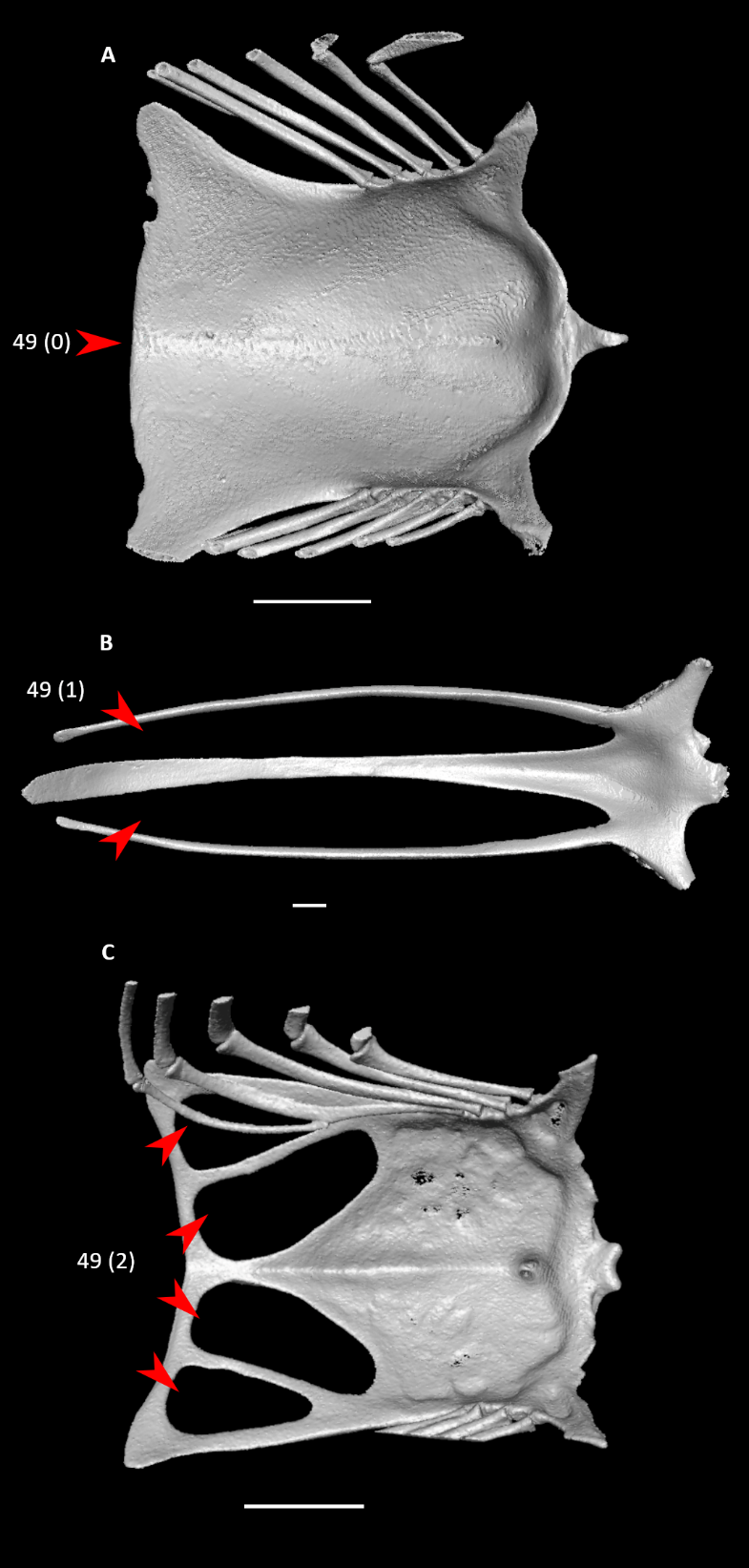

**Figure S39.** Sterna of *Oceanites oceanicus* (**A**), *Eudromia elegans* (**B**), and *Aegotheles cristatus* (**C**) in dorsal view, illustrating alternative states for character 49 (complete character description in text). Arrows indicate the caudal region of the sternum and the caudal incisures or fenestrae (when present). Scale bars = 5 mm.

*50. Sternum, caudal margin, cranial extent of lateral fenestrae or incisurae (Fig. S40; Livezey and Zusi, 2006: 1183; Musser and Cracraft, 2019: 184; Musser and Clarke, 2020: 341):

0 – short, length of incisurae or fenestrae less than one third craniocaudal length of sternal main body

1 – intermediate, length of incisurae or fenestrae between one third and two thirds of craniocaudal length of sternal main body

2 – long, length of incisurae or fenestrae greater than two thirds craniocaudal length of sternal main body, approaching caudal termination of costal processes

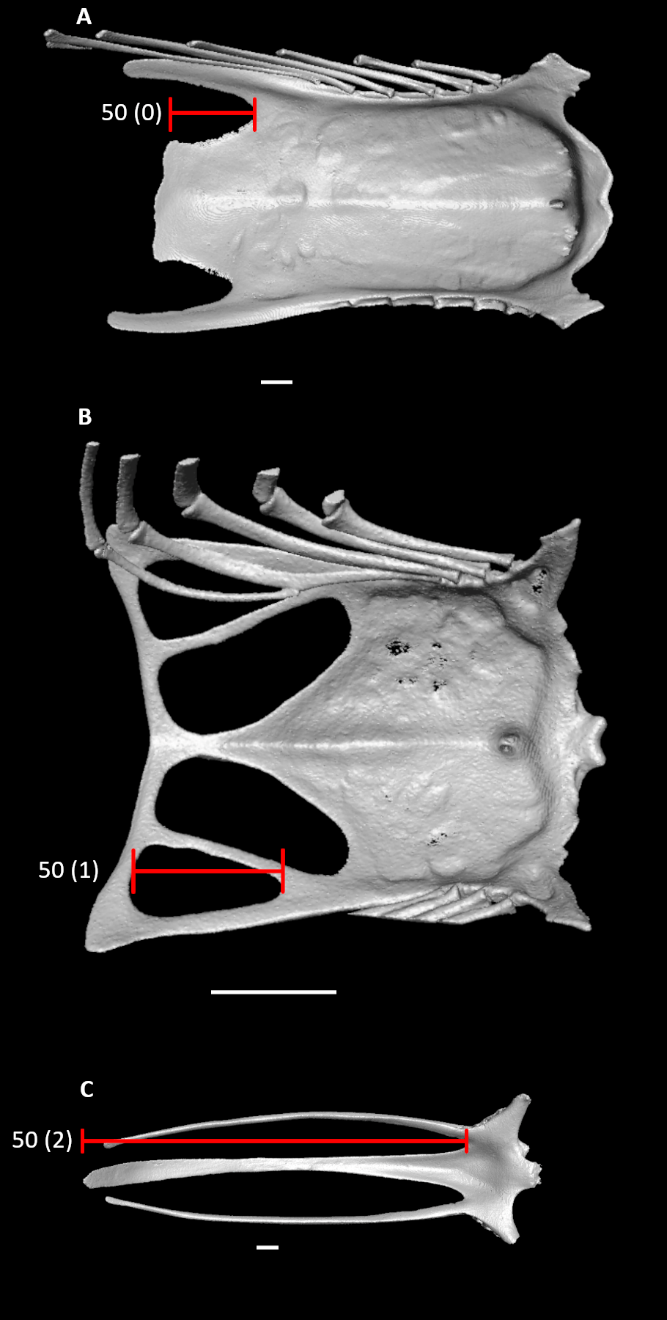

**Figure S40.** Sterna of *Dendrocygna bicolor* (**A**), *Aegotheles cristatus* (**B**), and *Eudromia elegans* (**C**) in dorsal view, illustrating alternative states for character 50 (complete character description in text). Red lines span the length of the lateral incisures or fenestrae. Scale bars = 5 mm.

51. Sternum, caudal margin, cranial extent of medial fenestrae or incisurae (Fig. S41; Livezey and Zusi, 2006: 1188; Musser and Cracraft, 2019: 188; Musser and Clarke, 2020: 354):

0 – short, length of incisurae or fenestrae less than one third craniocaudal length of sternal main body

1 – long, length of incisurae or fenestrae between one third and two thirds of craniocaudal length of sternal main body

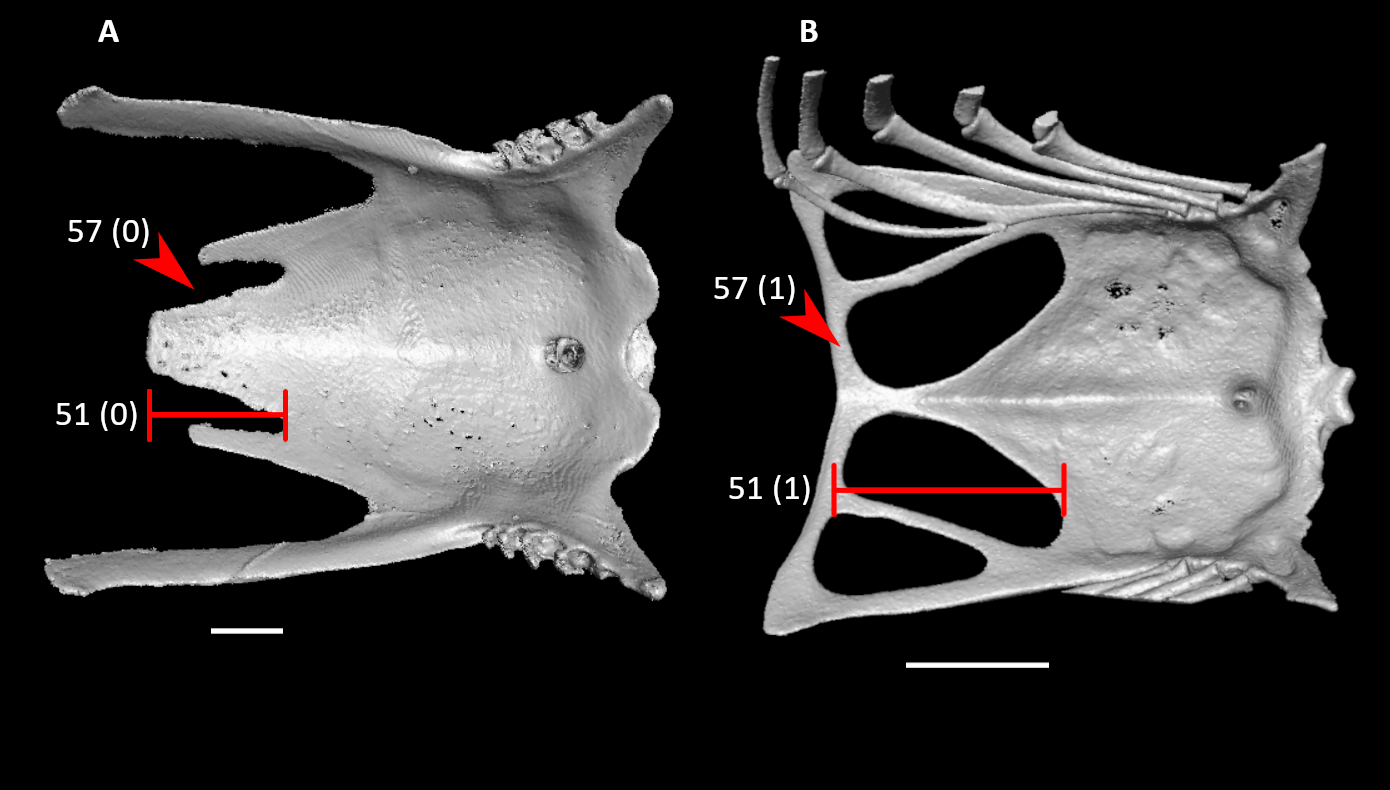

**Figure S41.** Sterna of *Podargus strigoides* (**A**) and *Aegotheles cristatus* (**B**) in dorsal view, illustrating alternative states for characters 51 and 57 (complete character descriptions in text). Red lines span the length of the medial incisures or fenestrae. Arrows indicate the caudal margin of the medial incisure or fenestra. Scale bars = 5 mm.

52. Sternum, caudal margin, caudolateral trabecula (if present), transverse expansion (Fig. S42; Livezey and Zusi, 2006: 1185; Musser and Cracraft, 2019: 186; Musser and Clarke, 2020: 345):

0 – absent or slight

1 – present

Broken in *Monias* specimen, scored based on *Mesitornis*.

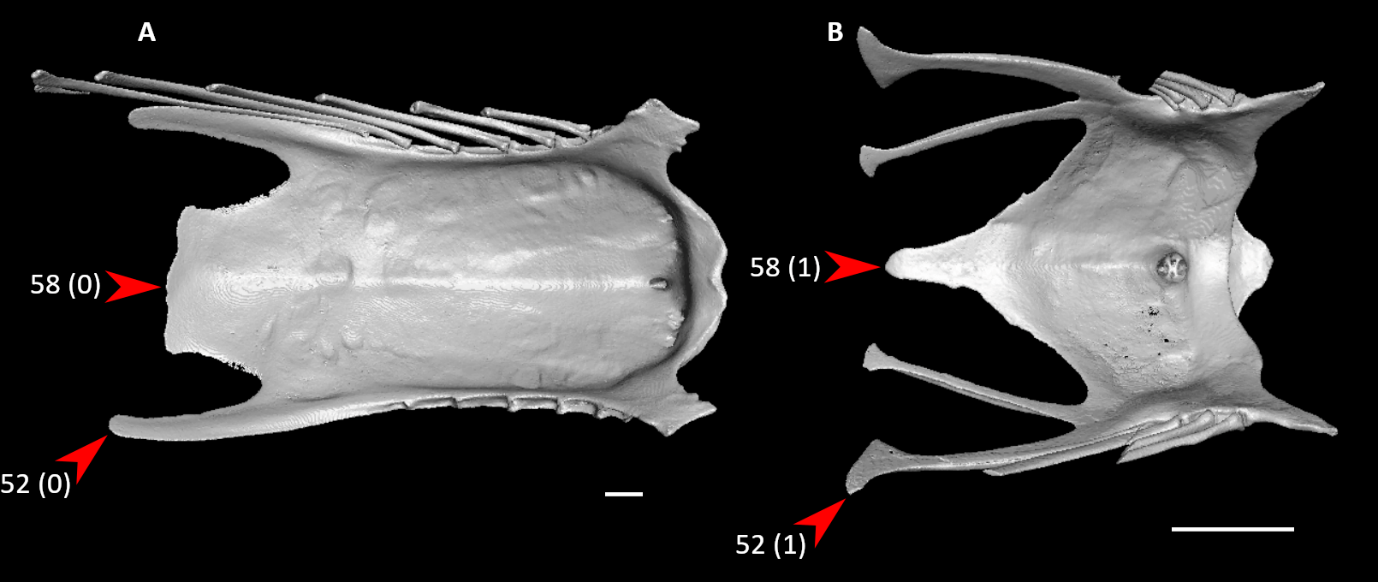

**Figure S42.** Sterna of *Dendrocygna bicolor* (**A**) and *Bucco capensis* (**B**) in dorsal view, illustrating alternative states for characters 52 and 58 (complete character descriptions in text). Arrows for character 52 indicate the tip of the caudolateral trabecula, whereas arrows for character 58 indicate the tip of the median trabecula. Scale bars = 5 mm.

*53. Sternum, caudal margin, caudolateral trabecula, orientation of main axis (Musser and Clarke, 2020: 346):

0 – turns medially

1 – essentially straight

2 – turns laterally

Broken in *Monias* specimen, scored based on *Mesitornis*.

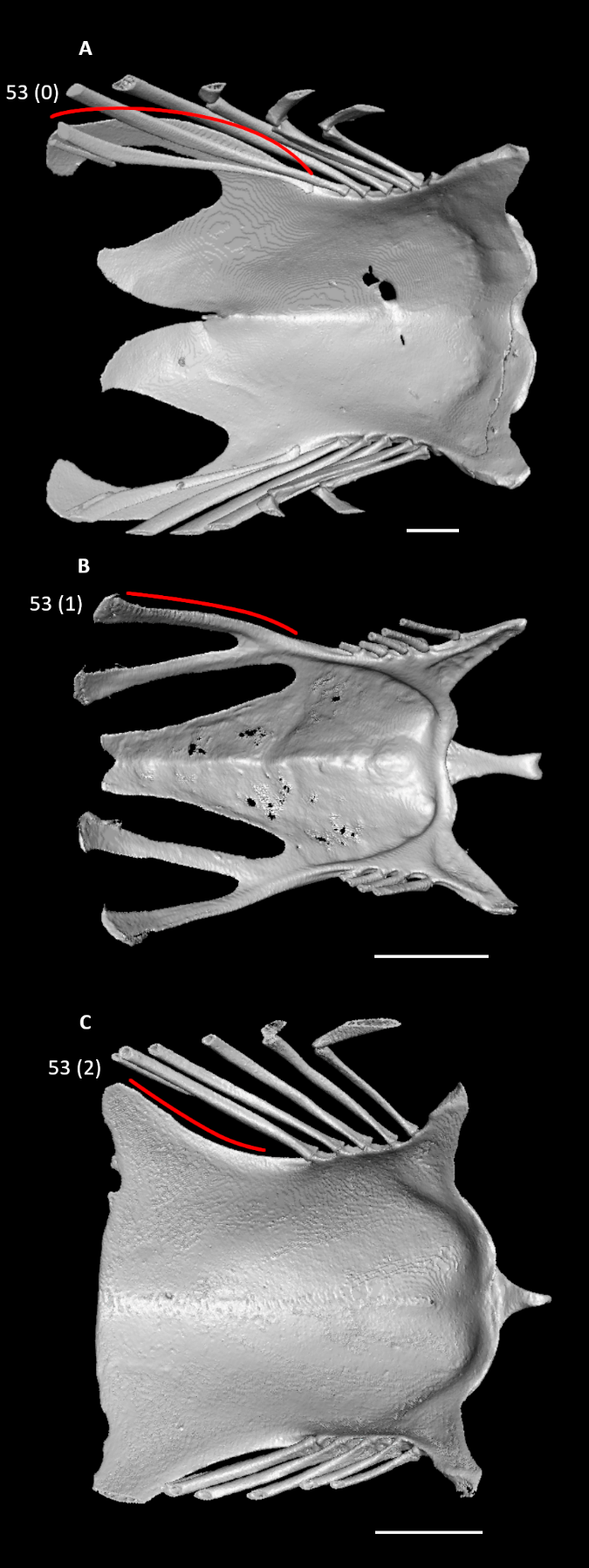

**Figure S43.** Sterna of *Podilymbus podiceps* (**A**), *Jynx torquilla* (**B**), and *Oceanites oceanicus* (**C**) in dorsal view, illustrating alternative states for character 53 (complete character description in text). Red lines indicate the approximate curvature of the caudolateral trabecula. Scale bars = 5 mm.

*54. Sternum, caudal margin, intermediate trabecula, orientation of main axis (Fig. S44; Musser and Clarke, 2020: 363):

0 – turns medially

1 – essentially straight

2 – turns laterally

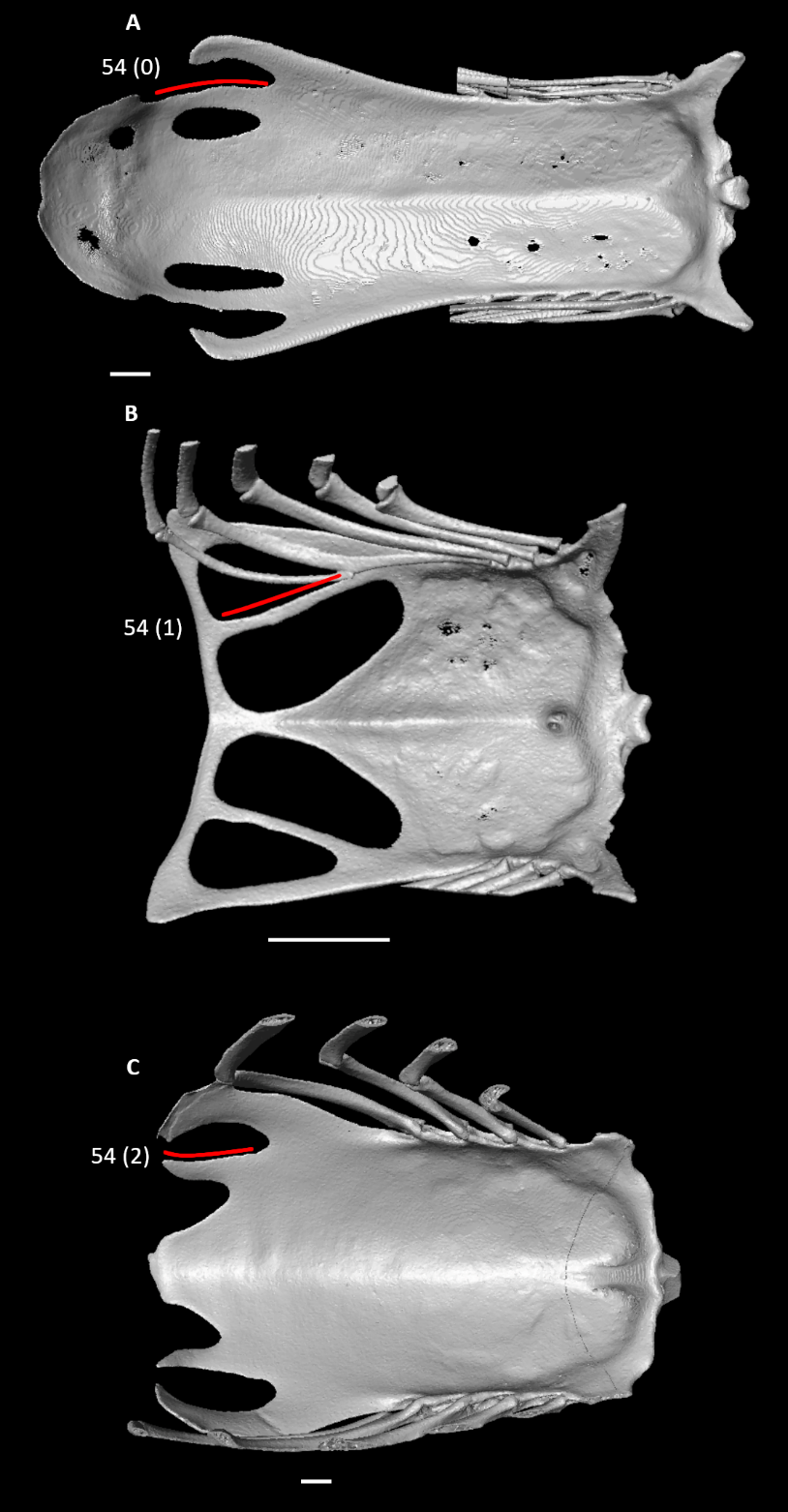

**Figure S44.** Sterna of *Alca torda* (**A**), *Aegotheles cristatus* (**B**), and *Coragyps atratus* (**C**) in dorsal view, illustrating alternative states for character 54 (complete character description in text). Red lines indicate the approximate curvature of the intermediate trabecula. Scale bars = 5 mm.

55. Sternum, lateral trabecula (Fig. S18; Musser and Clarke, 2020: 344):

0 – absent

1 – present

56. Sternum, caudal margin, ossified distal connection between caudolateral and intermediate or median trabeculae (Fig. S26; Musser and Clarke, 2020: 357):

0 – absent

1 – present

57. Sternum, caudal margin, ossified distal connection between intermediate and median trabeculae (Fig. S41; Musser and Clarke, 2020: 358):

0 – absent

1 – present

Characters 57–60 scored inapplicable for *Chauna*, which lacks a distinct median trabecula.

58. Sternum, caudal margin, median trabecula, form of margin and caudal termination (Fig. S42; Livezey and Zusi, 2006: 1191; Musser and Cracraft, 2019: 189; Musser and Clarke, 2020: 355):

0 – not tapered or weakly tapered

1 – distinctly tapered

Scored based on Fig. 2 in Mayr (2004a) for *Podica* (broken in specimen).

59. Sternum, caudal margin, median trabecula, shape of margin and caudal termination (Fig. S45; Livezey and Zusi, 2006: 1191; Musser and Cracraft, 2019: 189; Musser and Clarke, 2020: 356):

0 – rounded

1 – squared off

2 – V-shaped slit

Scored based on Fig. 2 in Mayr (2004a) for *Podica* (broken in specimen).

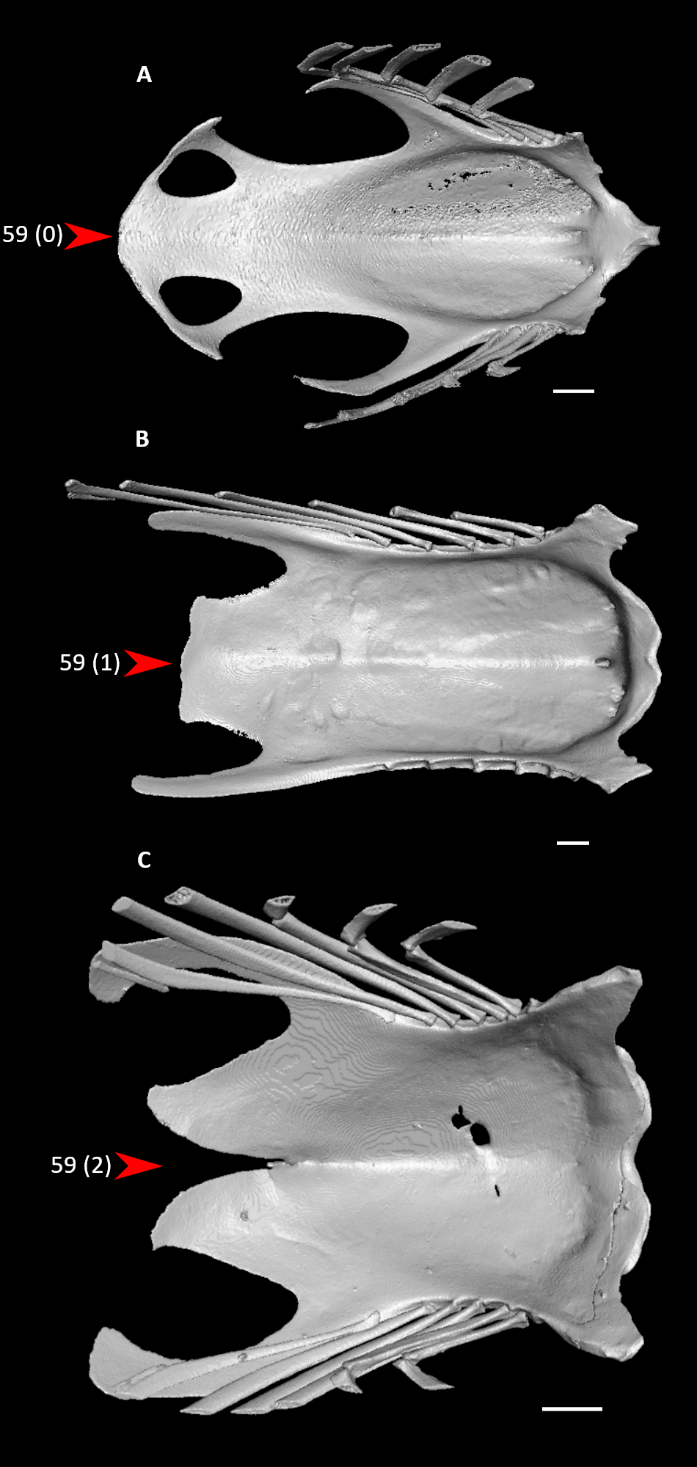

**Figure S45.** Sterna of *Columba livia* (**A**), *Dendrocygna bicolor* (**B**), and *Podilymbus podiceps* (**C**) in dorsal view, illustrating alternative states for character 59 (complete character description in text). Arrows indicate the tip of the median trabecula. Scale bars = 5 mm.

60. Sternum, caudal margin, caudal extents of caudolateral and median trabeculae (Fig. S30; Livezey and Zusi, 2006: 1192; Musser and Cracraft, 2019: 190; Musser and Clarke, 2020: 360):

0 – median > caudolateral

1 – caudolateral > median

Broken in *Monias* specimen, scored based on *Mesitornis*.

61. Proportions of scapula and coracoid (Fig. S46; Ksepka et al., 2019: 75):

0 – coracoid longer than scapula

1 – scapula longer than coracoid

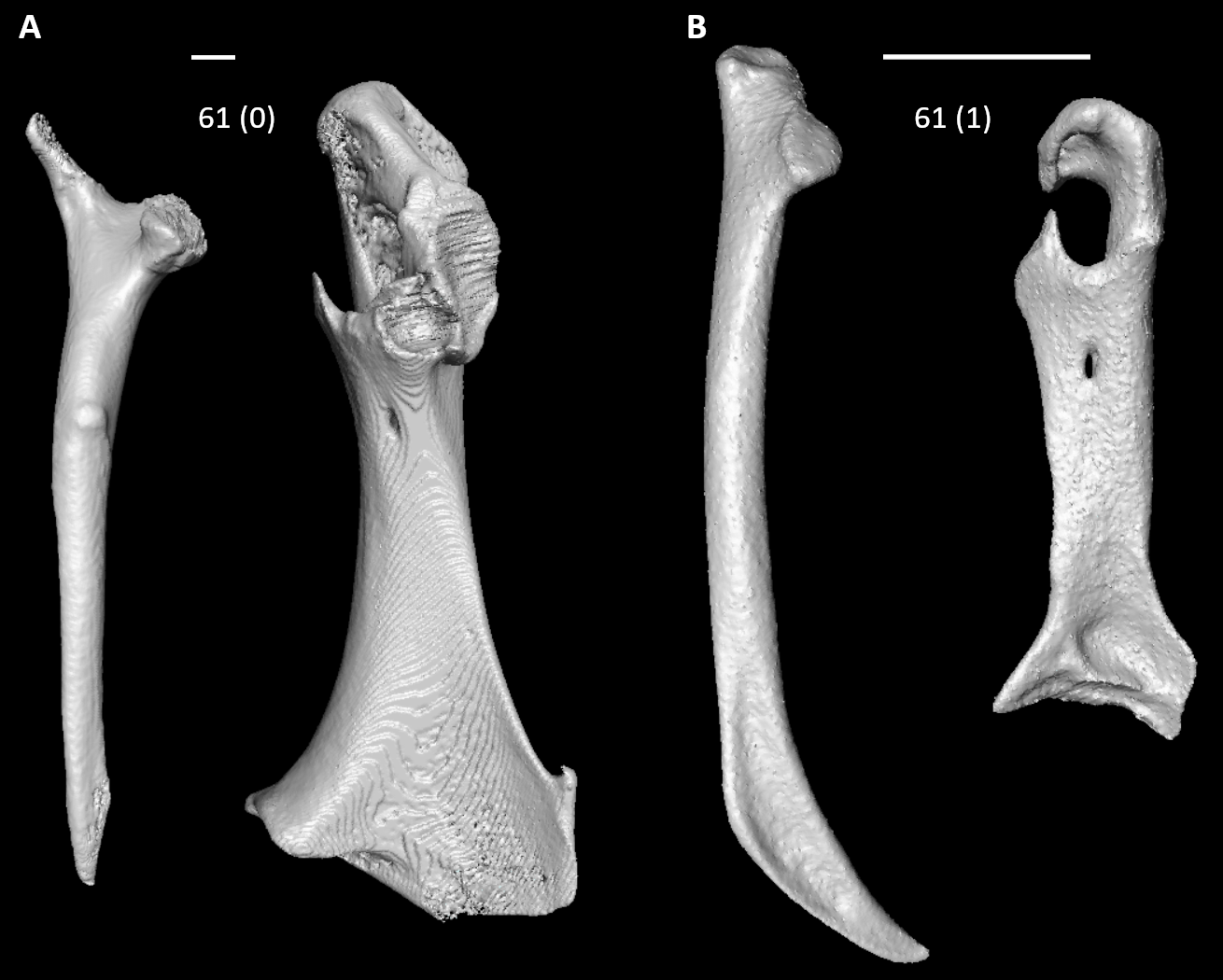

**Figure S46.** Scapulae (left) and coracoids (right) of *Pelecanus occidentalis* (**A**, left elements, mirrored) and *Aegotheles cristatus* (**B**, right elements) showing the relative lengths of the two elements for each species, illustrating alternative states for character 61 (complete character description in text). Scapulae are shown in lateral view, whereas coracoids are in dorsal view. Scale bars = 5 mm.

62. Coracoid, acrocoracoid process, principal dorsoventral orientation relative to major craniocaudal axis of coracoid in lateral view (Fig. S47; Livezey and Zusi, 2006: 1267; Musser and Cracraft, 2019: 193; Musser and Clarke, 2020: 375):

0 – distinctly ventral

1 – essentially coplanar

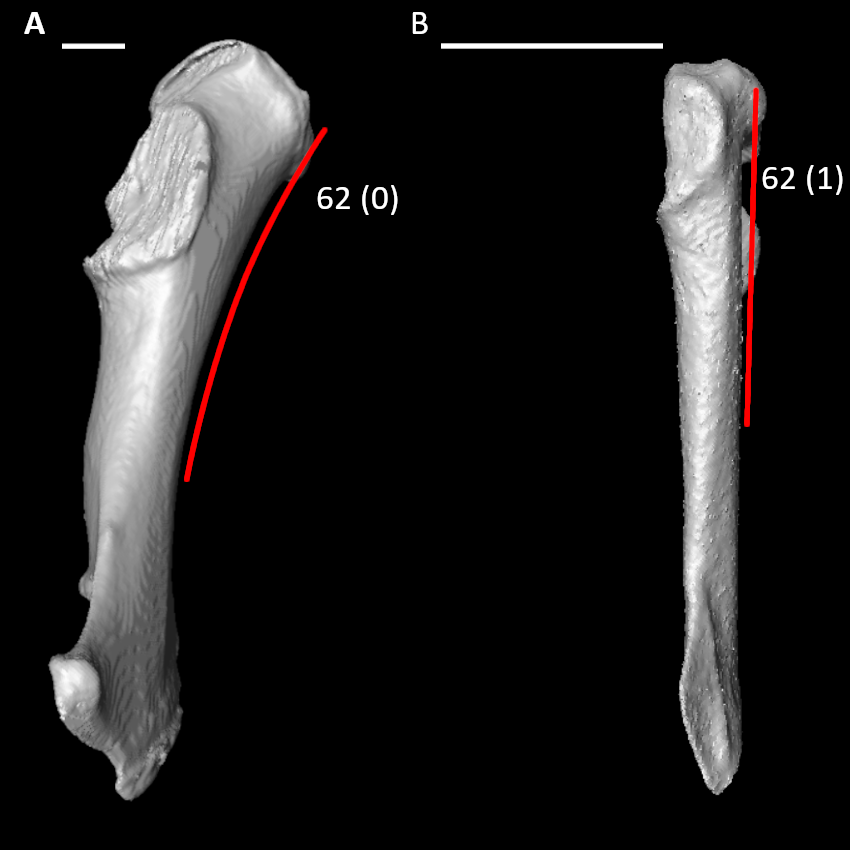

**Figure S47.** Coracoids of *Phoenicopterus ruber* (**A**, left element, mirrored) and *Aegotheles cristatus* (**B**, right element) in lateral view, illustrating alternative states for character 62 (complete character description in text). Red lines indicate approximate ventral curvature of the acrocoracoid process. Scale bars = 5 mm.

63. Coracoid, acrocoracoid process, medial curvature (Fig. S48; Livezey and Zusi, 2006: 1268; Smith, 2010: 167; Musser and Cracraft, 2019: 194):

0 – straight

1 – pronounced curvature, forming distinct hook

Scored straight in *Opisthocomus*.

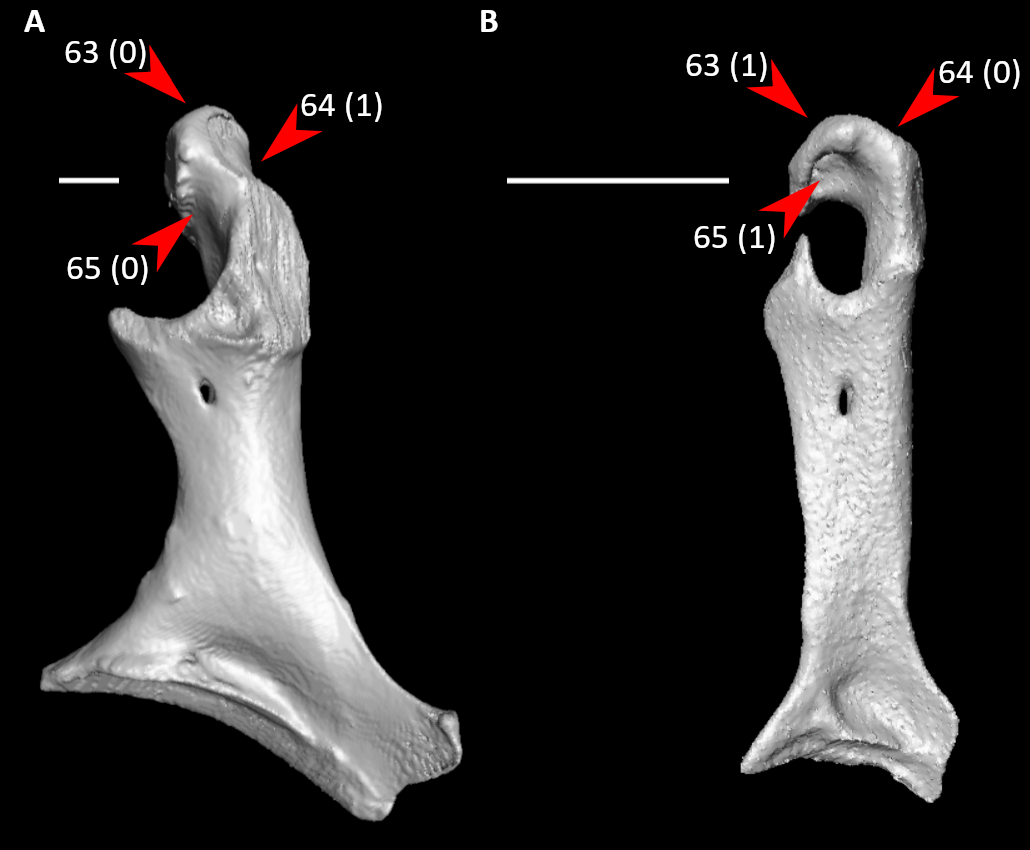

**Figure S48.** Coracoids of *Phoenicopterus ruber* (**A**, left element, mirrored) and *Aegotheles cristatus* (**B**, right element) in dorsal view, illustrating alternative states for characters 63, 64, and 65 (complete character descriptions in text). Arrows for character 63 indicate the tip of the acrocoracoid process, arrows for character 64 indicate the impression for the acrocoracohumeral ligament, and arrows for character 65 indicate the infra-acrocoracoid recess. Scale bars = 5 mm.

64. Coracoid, acrocoracoid process, impression for acrocoracohumeral ligament (Fig. S48):

0 – absent or shallow

1 – deep

65. Coracoid, infra-acrocoracoid recess (Fig. S48; Musser and Clarke, 2020: 374):

0 – shallow or absent

1 – deep

*66. Coracoid, dorsal view, scapular cotyle (Fig. S49; Ericson, 1997: 40; Livezey and Zusi, 2006: 1281; Ksepka, 2009: 47; Worthy et al., 2017: 95; Ksepka et al., 2019: 45; Musser and Clarke, 2020: 391):

0 – deep and cup-like

1 – shallow concavity

2 – nearly flat

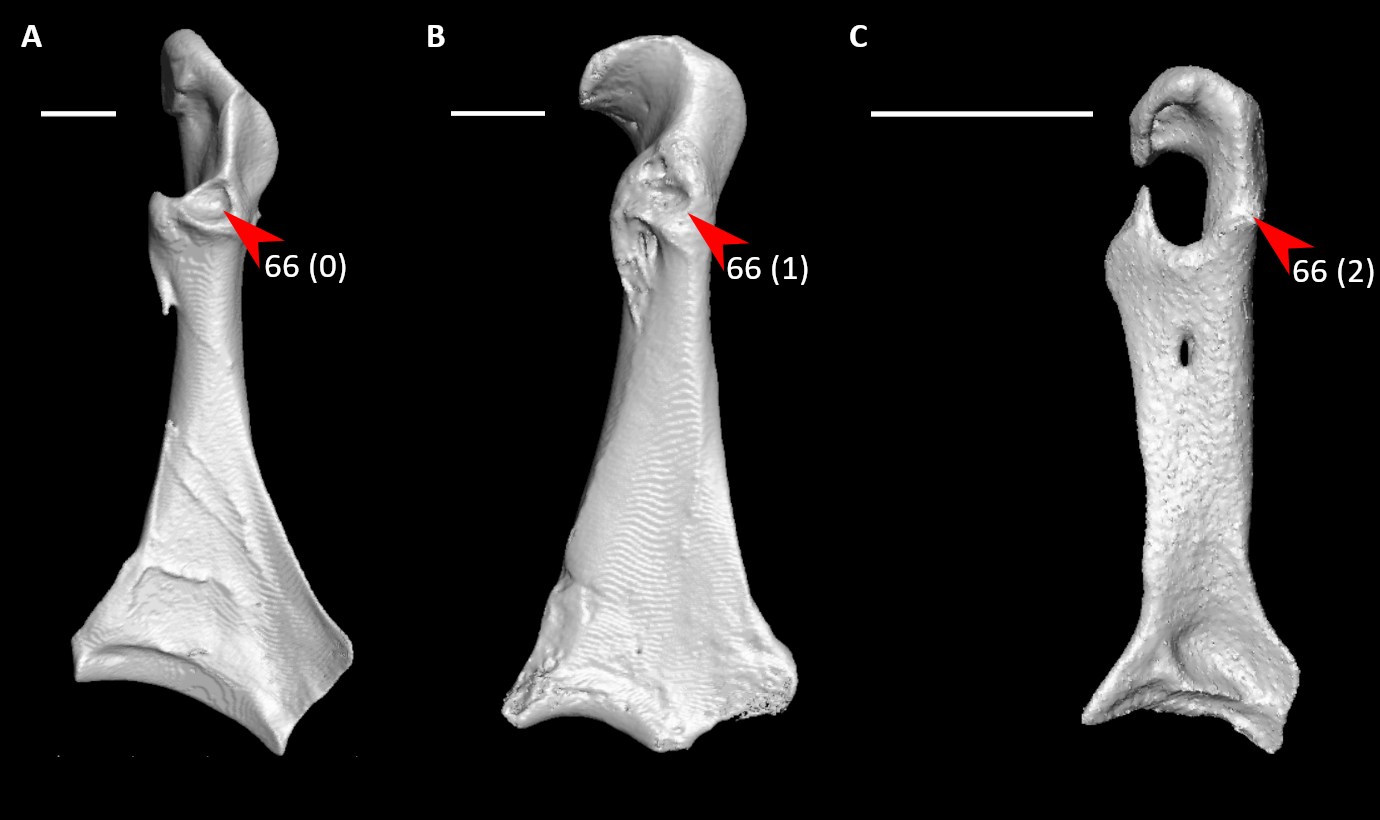

**Figure S49.** Coracoids of *Dendrocygna bicolor* (**A**, left element, mirrored), *Eudromia elegans* (**B**, left element, mirrored), and *Aegotheles cristatus* (**C**, right element) in dorsal view, illustrating alternative states for character 66 (complete character description in text). Arrows indicate the scapular cotyle. Scale bars = 5 mm.

67. Coracoid, dorsal view, articular facet for humerus (Fig. S50; Worthy et al., 2017: 100):

0 – concave

1 – flat or convex

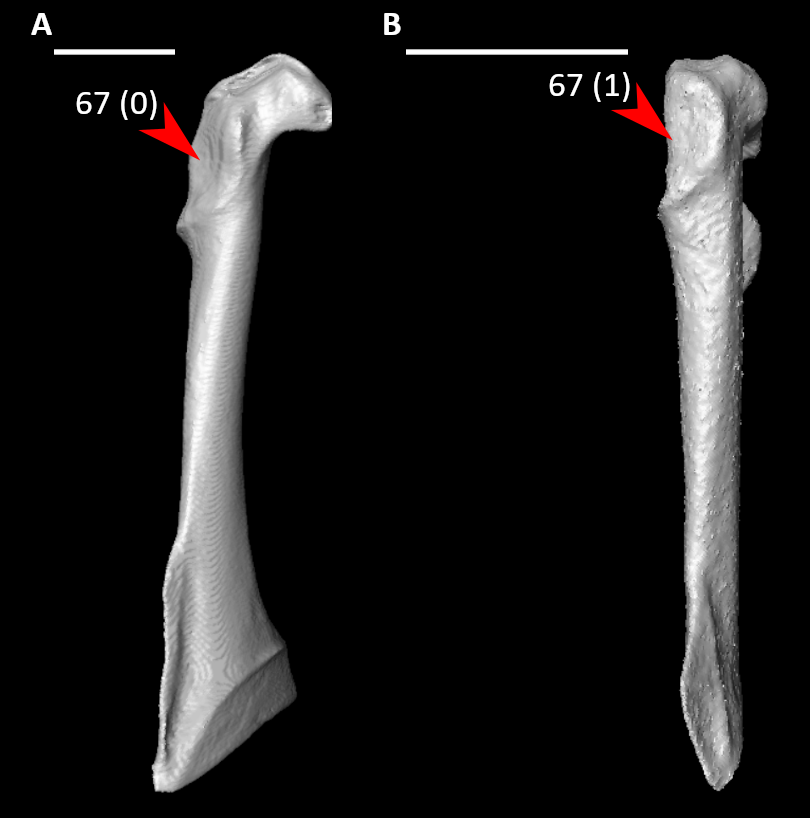

**Figure S50.** Coracoids of *Podilymbus podiceps* (**A**, right element) and *Aegotheles cristatus* (**B**, right element) in lateral view, illustrating alternative states for character 67 (complete character description in text). Arrows indicate the articular facet for the humerus. Scale bars = 5 mm.

*68. Coracoid, dorsal view, procoracoid process form (Fig. S51; Livezey and Zusi, 2006: 1283; Musser and Cracraft, 2019: 197; Musser and Clarke, 2020: 382):

0 – shallow cotyle or crest

1 – moderately prominent, a typically curved tubercle

2 – an elongate process, terminal end distinctly extending medially or dorsally to main body of coracoid

3 – markedly curved, creating circular connection with acrocoracoid process

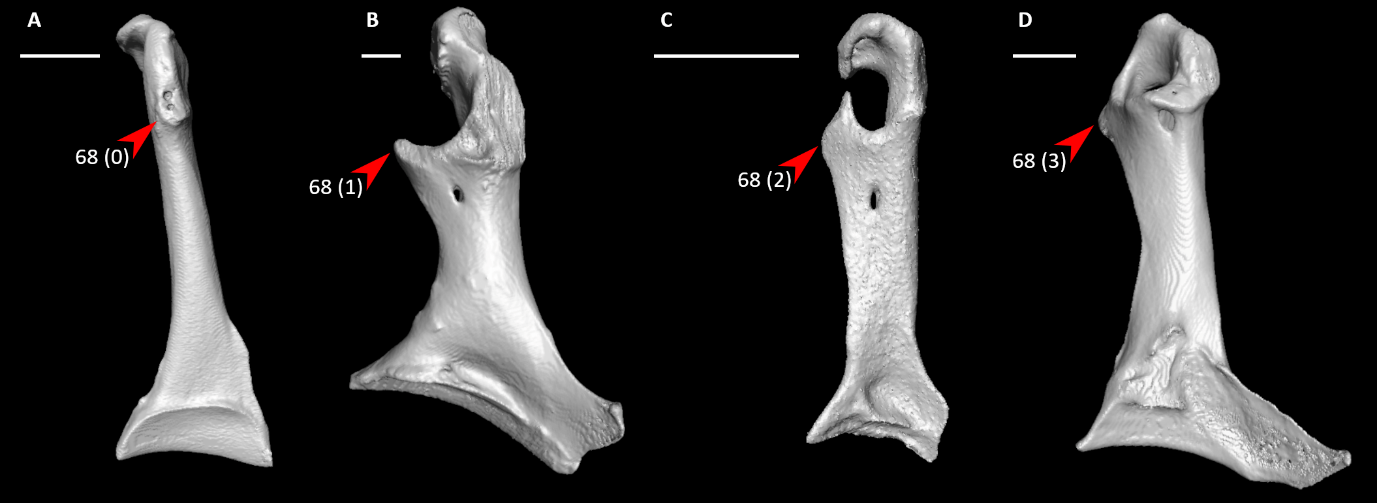

**Figure S51.** Coracoids of *Podilymbus podiceps* (**A**, right element), *Phoenicopterus ruber* (**B**, left element, mirrored), *Aegotheles cristatus* (**C**, right element), and *Corythaeola cristata* (**D**, left element, mirrored) in dorsal view, illustrating alternative states for character 68 (complete character description in text). Arrows indicate the procoracoid process. Scale bars = 5 mm.

69. Coracoid, omal end, main body of coracoid at ventromedial margin of supracoracoid sulcus (Fig. S52; Worthy et al., 2017: 103):

0 – rounded, relatively thick

1 – compressed, keeled

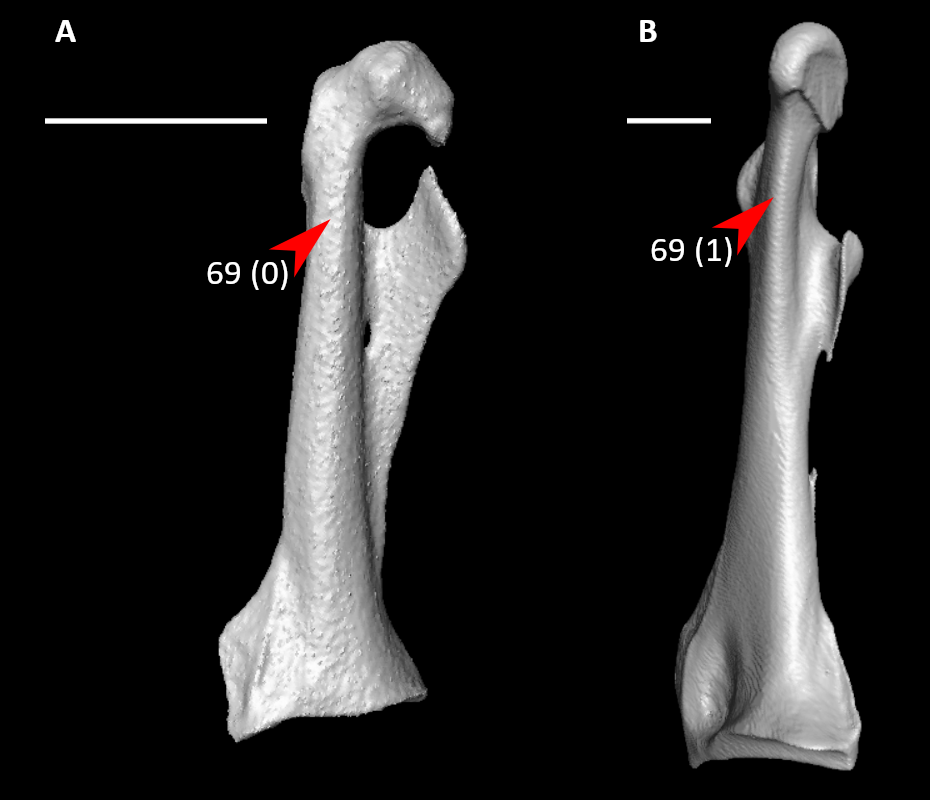

**Figure S52.** Coracoids of *Aegotheles cristatus* (**A**, right element) and *Dendrocygna bicolor* (**B**, left element, mirrored) in ventromedial view, illustrating alternative states for character 69 (complete character description in text). Arrows indicate the coracoid main body at the ventromedial margin of supracoracoid sulcus. Scale bars = 5 mm.

70. Coracoid, shaft, supracoracoid nerve foramen (Fig. S53; Mayr and Clarke, 2003: 65; Musser and Clarke, 2020: 393)

0 – absent

1 – present

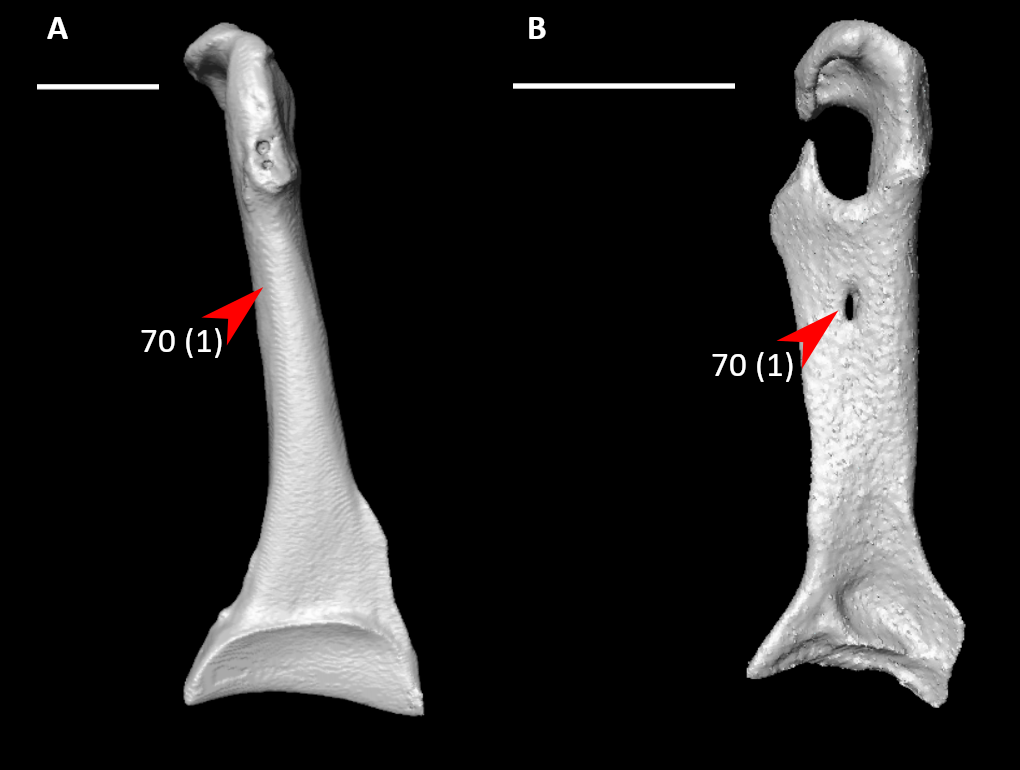

**Figure S53.** Coracoids of *Podilymbus podiceps* (**A**, right element) and *Aegotheles cristatus* (**B**, right element) in dorsal view, illustrating alternative states for character 70 (complete character description in text). Arrows indicate the supracoracoid nerve foramen or approximate homologous site. Scale bars = 5 mm.

71. Coracoid, shaft, supracoracoid nerve foramen (Fig. S54; Livezey and Zusi, 2006: 1286; Musser and Cracraft, 2019: 198):

0 – circular

1 – craniocaudally elongate

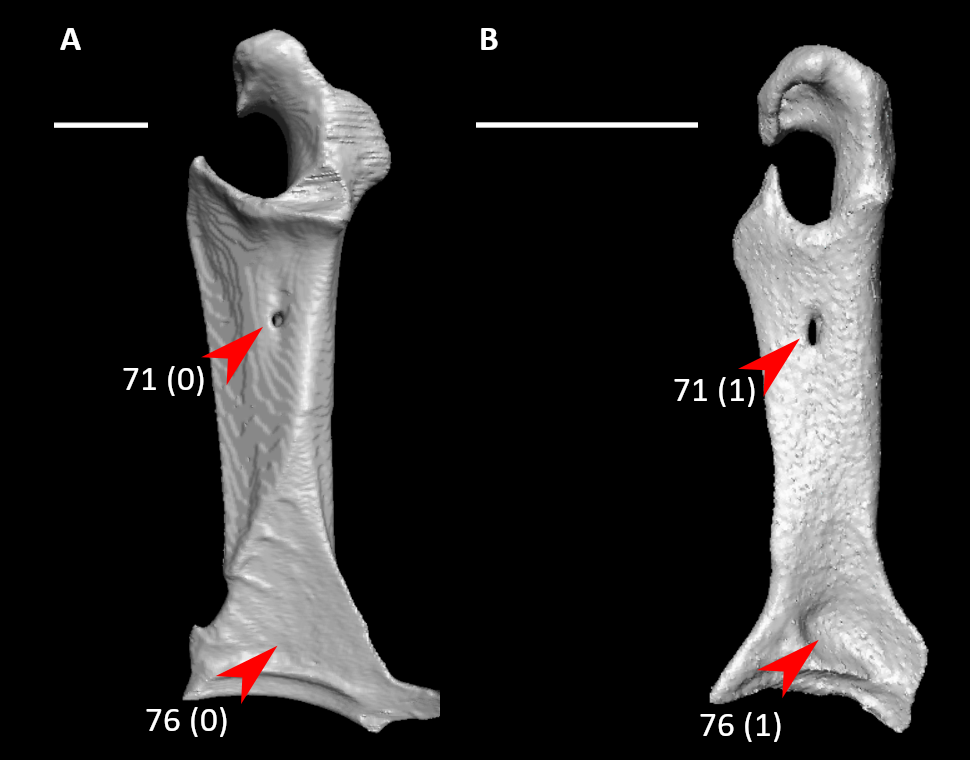

**Figure S54.** Coracoids of *Podica senegalensis* (**A**, left element, mirrored) and *Aegotheles cristatus* (**B**, right element) in dorsal view, illustrating alternative states for characters 71 and 76 (complete character descriptions in text). Arrows for character 71 indicate the supracoracoid nerve foramen, whereas arrows for character 76 indicate the impression of m. sternocoracoidei. Scale bars = 5 mm.

*72. Coracoid, shaft, length relative to width of articular facet for sternum (Fig. S55; Livezey and Zusi, 2006: 1292; Musser and Cracraft, 2019: 199; Musser and Clarke, 2020: 395):

0 – short, length less than two times the width

1 – moderately short, length between two and three times the width

2 – moderately elongate, length between three and four times the width

3 – very elongate, length greater than four times the width

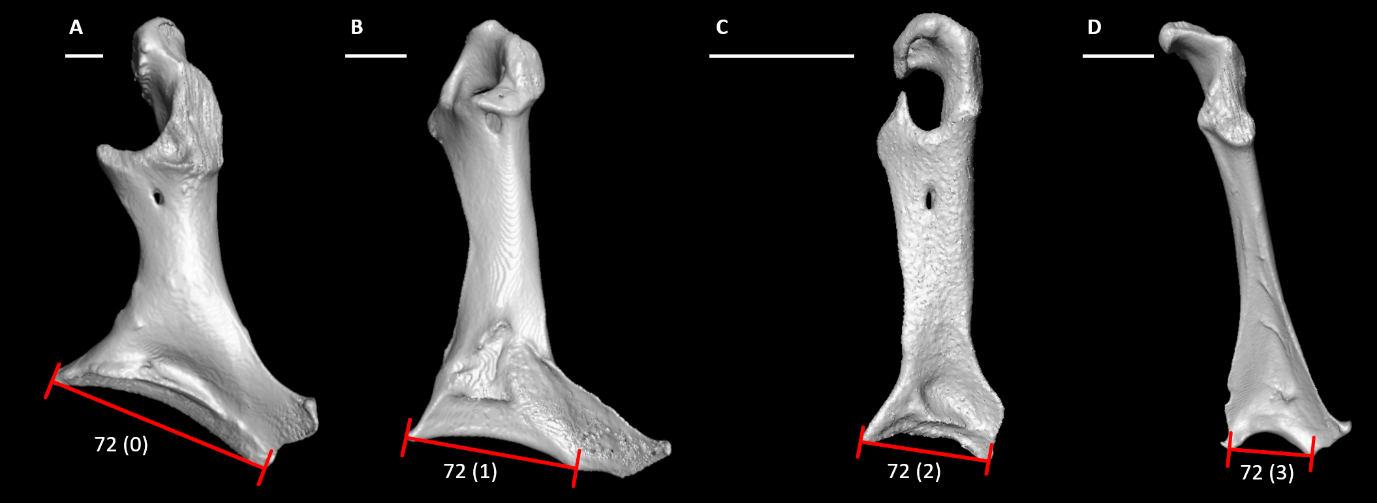

**Figure S55.** Coracoids of *Phoenicopterus ruber* (**A**, left element, mirrored), *Corythaeola cristata* (**B**, left element, mirrored), *Aegotheles cristatus* (**C**, right element), and *Rollulus rouloul* (**D**, right element) in dorsal view, illustrating alternative states for character 72 (complete character description in text). Red lines span the width of the articular facet for the sternum. Scale bars = 5 mm.

73. Coracoid, shaft, line linking acrocoracoid process and medial angle forming angle with line linking lateral and medial extremes of sternal facet (Fig. S56; Worthy and Lee, 2008: 50; Worthy et al., 2017: 108):

0 – markedly greater than 90-100 degrees

1 – approximates 90-100 degrees

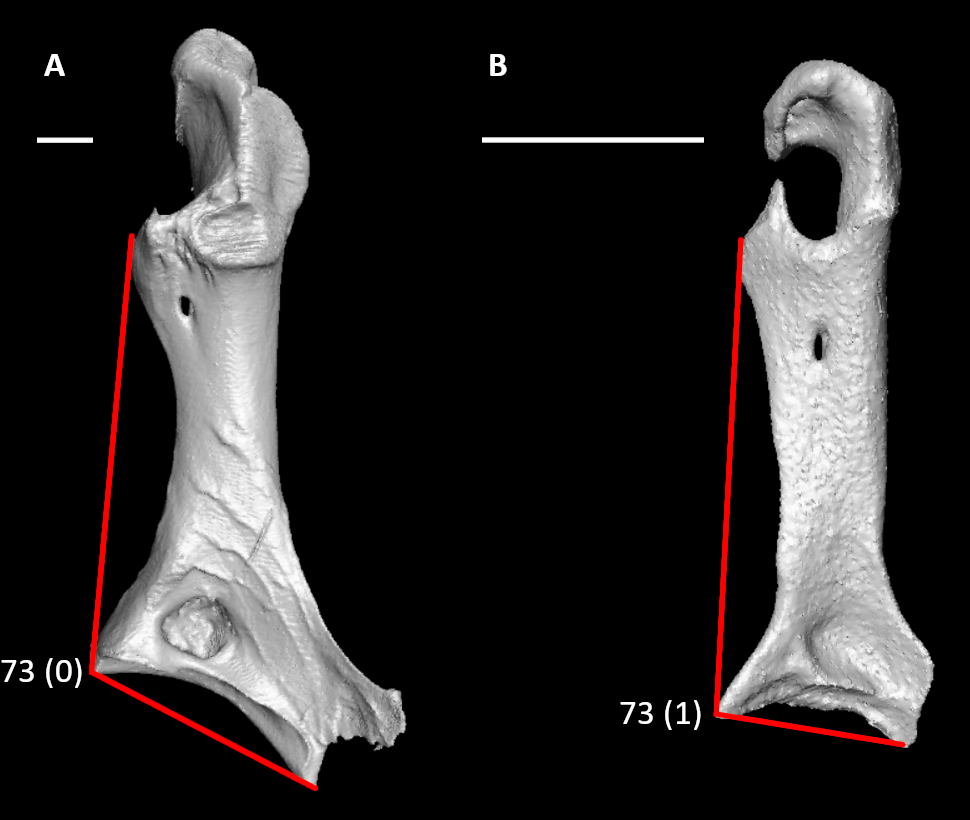

**Figure S56.** Coracoids of *Anseranas semipalmata* (**A**, left element, mirrored) and *Aegotheles cristatus* (**B**, right element) in dorsal view, illustrating alternative states for character 73 (complete character description in text). Red lines indicate the angle between the line linking the acrocoracoid process and medial angle with the line linking the lateral and medial extremes of the sternal facet. Scale bars = 5 mm.

74. Coracoid, shaft, several muscle scars diagonally traversing dorsal surface (Fig. S57; Ericson, 1997: 38; Worthy et al., 2017: 105):

0 – absent

1 – present

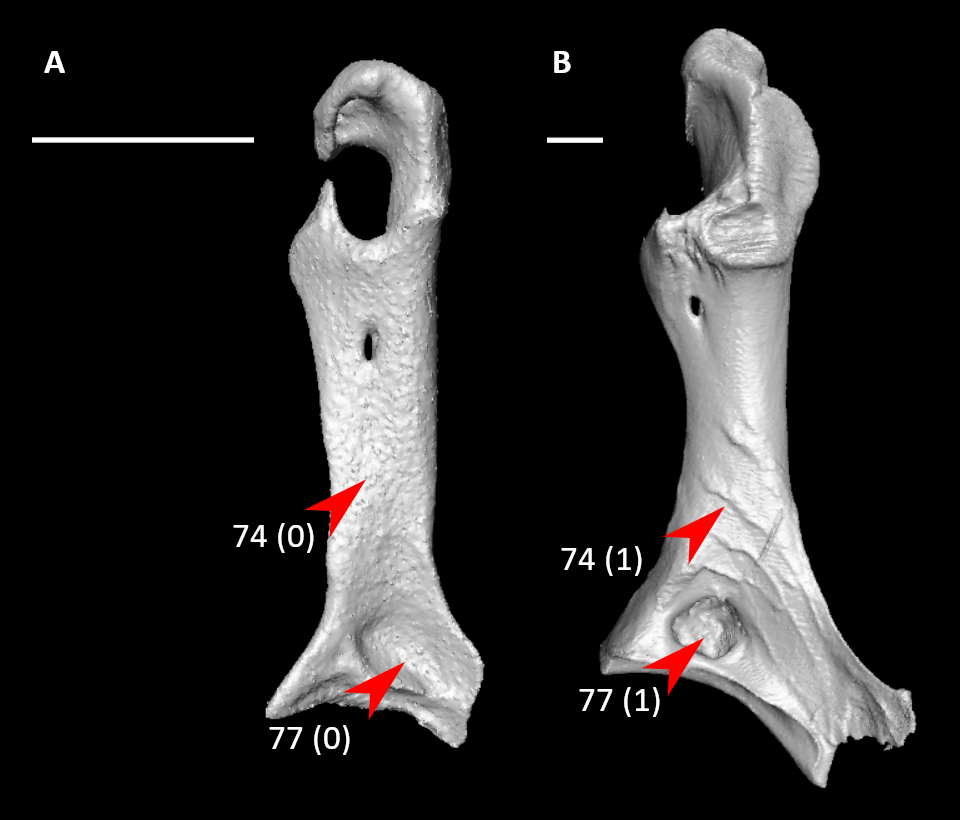

**Figure S57.** Coracoids of *Aegotheles cristatus* (**A**, right element) and *Anseranas semipalmata* (**B**, left element, mirrored) in dorsal view, illustrating alternative states for characters 74 and 77 (complete character descriptions in text). Arrows for character 74 indicate the coracoid shaft and diagonal muscle scars (when present), whereas arrows for characters 77 indicate the impression of m. sternocoracoidei and its pneumatic foramen (when present). Scale bars = 5 mm.

75. Coracoid, sternal end, notch in medial margin (Fig. S58; Mayr, 2015: 17):

0 – absent

1 – present

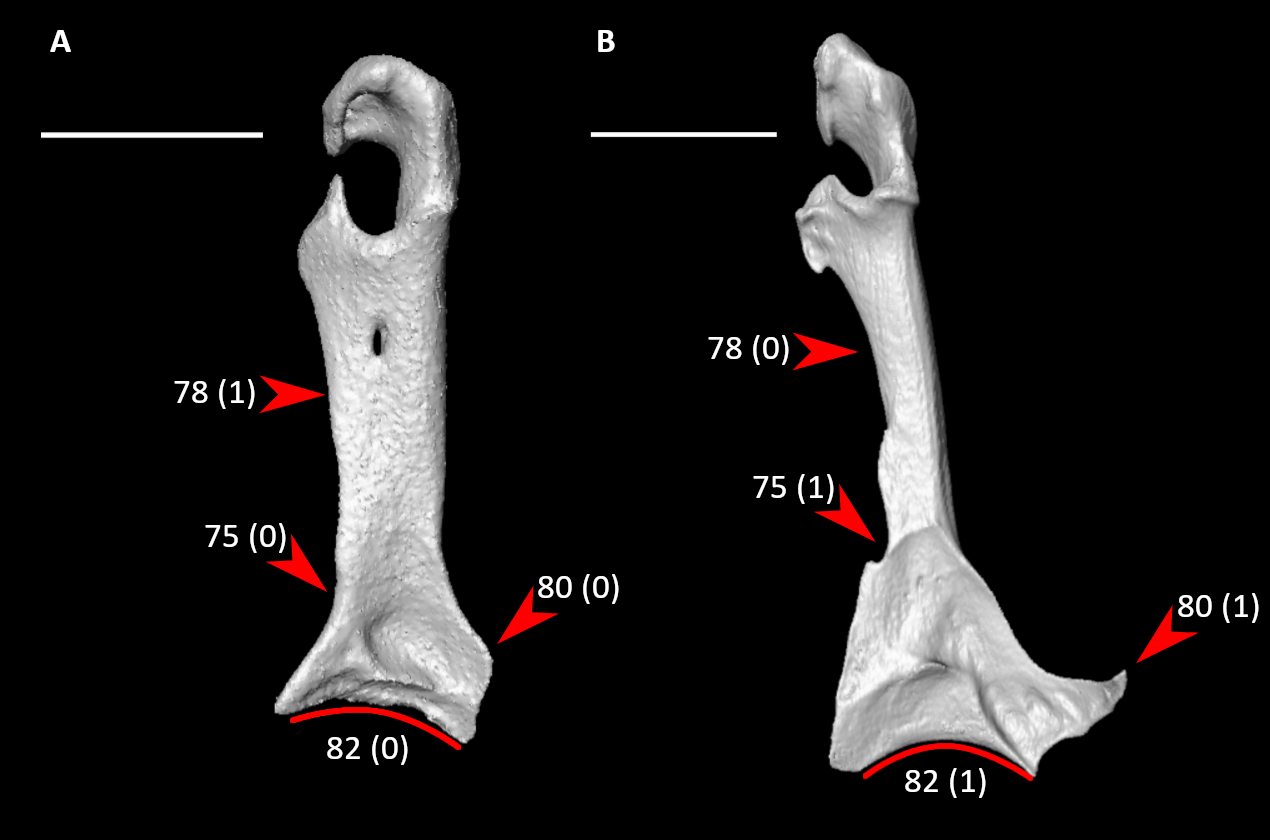

**Figure S58.** Coracoids of *Aegotheles cristatus* (**A**, right element) and *Bucco capensis* (**B**, right element) in dorsal view, illustrating alternative states for characters 75, 78, 80, and 82 (complete character descriptions in text). Arrows for character 75 indicate the medial margin of the coracoid and its notch (when present), arrows for character 78 indicate the cranial extension of the medial crest or approximate homologous site, and arrows for character 80 indicate the lateral process. Red lines indicate approximate cranial curvature of the articular facet with the sternum. Scale bars = 5 mm.

76. Coracoid, sternal end, impression of m. sternocoracoidei on dorsal surface of sternal extremity (Fig. S54; Livezey and Zusi, 2006: 1294; Musser and Cracraft, 2019: 200; Musser and Clarke, 2020: 396):

0 – shallow or difficult to discern

1 – deep

Scored shallow in *Elanus*, but there is a depression near the sternal end of the impression.

77. Coracoid, sternal end, dorsal surface, pneumatic foramen in impression of m. sternocoracoidei (Fig. S57; Mayr and Clarke, 2003: 67; Worthy et al., 2017: 104; Musser and Cracraft, 2019: 201; Musser and Clarke, 2020: 398):

0 – small or not visible

1 – large, ovoid

78. Coracoid, sternal end, medial crest (Fig. S58; Livezey, 1998: 194; Musser and Cracraft, 2019: 202):

0 – slight or absent

1 – present, continued cranially by procoracoid crest

79. Coracoid, sternal end, extent of lateral process (Fig. S59; Musser and Clarke, 2020: 405):

0 – lateral extent measured from coracoid shaft less than half the width of sternal facet

1 – lateral extent measured from coracoid shaft half the width of sternal facet or greater

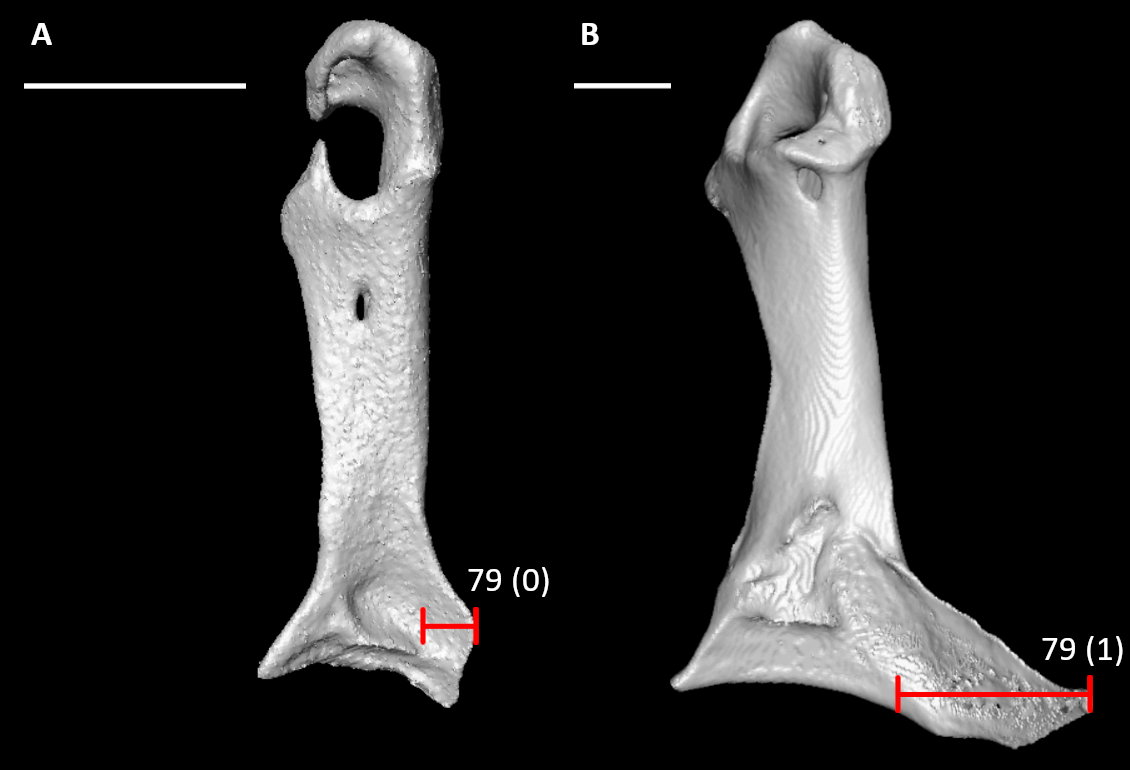

**Figure S59.** Coracoids of *Aegotheles cristatus* (**A**, right element) and *Corythaeola cristata* (**B**, left element, mirrored) in dorsal view, illustrating alternative states for character 79 (complete character description in text). Red lines span the width of the lateral process. Scale bars = 5 mm.

80. Coracoid, sternal end, lateral process shape (Fig. S58; Mayr, 2015: 19; Musser and Clarke, 2020: 407):

0 – unhooked or weakly hooked

1 – sharply hooked, such that tip forms a sharp, cranially-directed point

81. Coracoid, sternal end, articular facet for sternum, cranial extent of cranial margin of external lip relative to that of internal lip (Fig. S60; Livezey and Zusi, 2006: 1314; Musser and Cracraft, 2019: 205; Musser and Clarke, 2020: 414):

0 – former distinctly caudal to latter, producing internally (dorsally) angled facet

1 – former approximately equal to latter, producing facet of dorsoventrally equal expanse

2 – former significantly cranial to the latter, producing externally (ventrally) angled facet

Scored as inapplicable in Apodiformes, in which facet is concave, and *Opisthocomus*, in which coracoids are fused to sternum.

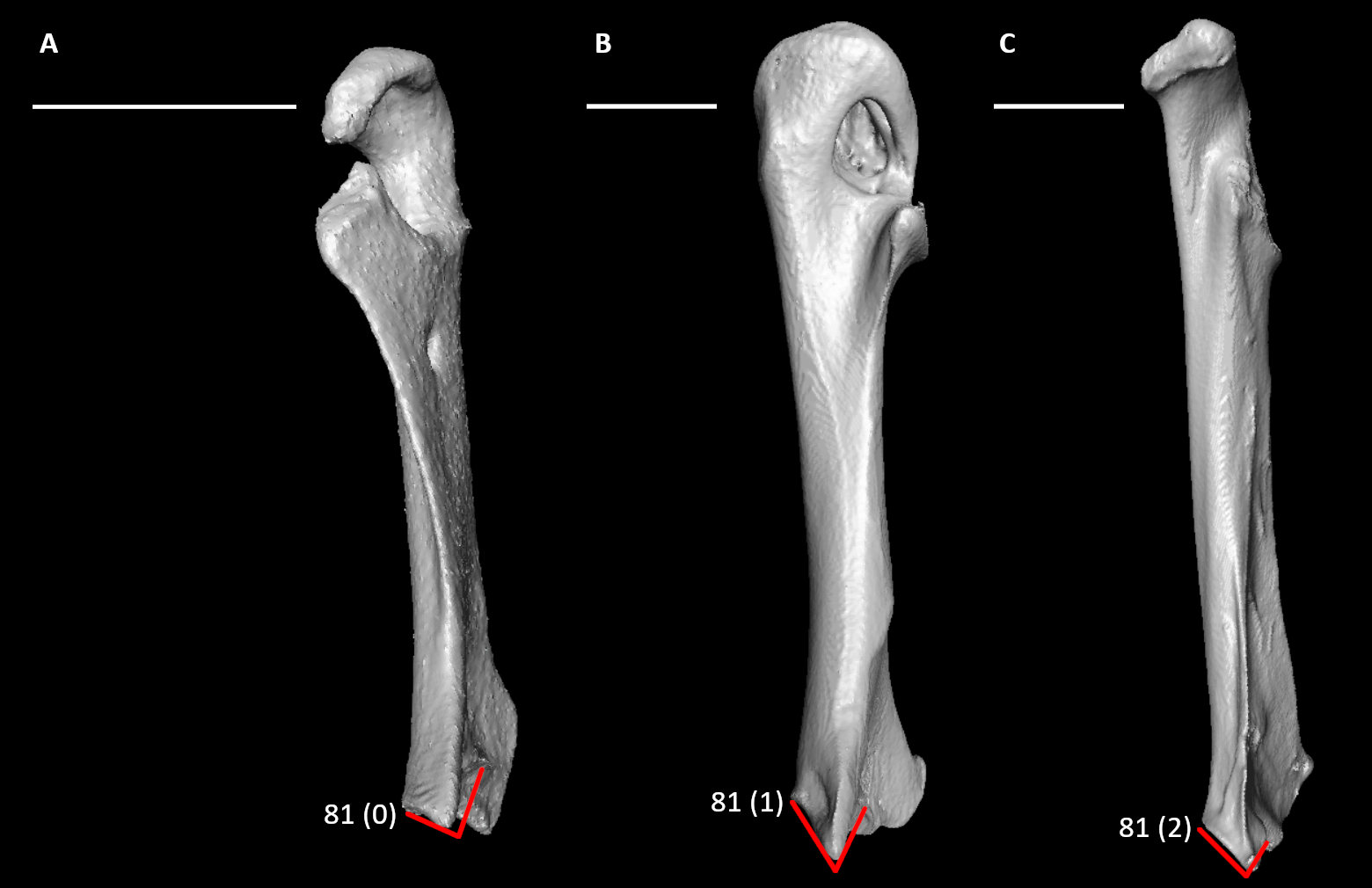

**Figure S60.** Coracoids of *Aegotheles cristatus* (**A**, right element), *Scopus umbretta* (**B**, right element), and *Rollulus rouloul* (**C**, right element) in medial view, illustrating alternative states for character 81 (complete character description in text). Red lines indicate approximate angle and extent of external and internal lips of the articulation facet with the sternum. Scale bars = 5 mm.

82. Coracoid, sternal end, cranial curvature of articular facet in dorsal view (Fig. S58):

0 – straight or slight

1 – strong curvature

Scored as inapplicable in *Opisthocomus*, in which coracoids are fused to sternum.

83. Coracoid, sternal end, ventral curvature of articular facet in sternal view (Fig. S61; Musser and Clarke, 2020: 400):

0 – straight or slight

1 – strong curvature

Scored as inapplicable in *Opisthocomus*, in which coracoids are fused to sternum.

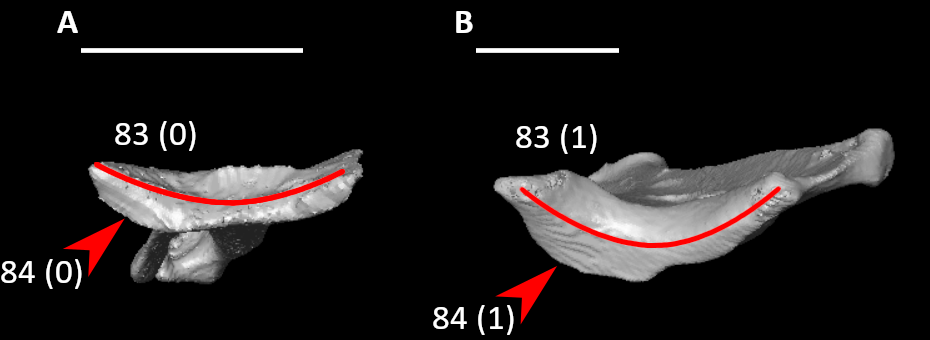

**Figure S61.** Coracoids of *Aegotheles cristatus* (**A**, right element) and *Rollulus rouloul* (**B**, right element) in sternal view, illustrating alternative states for characters 83 and 84 (complete character descriptions in text). Arrows indicate the external lip of the articulation facet with the sternum. Red lines indicate approximate ventral curvature of the articular facet with the sternum. Scale bars = 5 mm.

84. Coracoid, sternal end, external lip in sternal view (Fig. S61):

0 – little dorsoventral expansion

1 – prominently expanded

Scored as inapplicable in *Opisthocomus*, in which coracoids are fused to sternum.

85. Coracoid, sternal end, external lip expansion (Fig. S62):

0 – expanded along entire width

1 – only expanded in medial portion

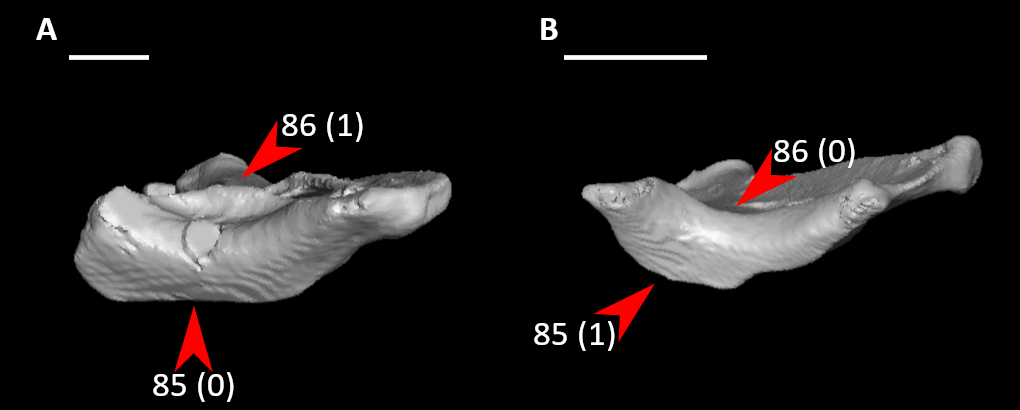

**Figure S62.** Coracoids of *Alectura lathami* (**A**, left element, mirrored) and *Rollulus rouloul* (**B**, right element) in sternal view, illustrating alternative states for characters 85 and 86 (complete character descriptions in text). Arrows for character 85 indicate the external lip of the articulation facet with the sternum, whereas arrows for character 86 indicate the internal lip. Scale bars = 5 mm.

86. Coracoid, sternal end, internal lip in sternal view (Fig. S62):

0 – little dorsoventral expansion

1 – prominently expanded

Scored as inapplicable in *Opisthocomus*, in which coracoids are fused to sternum.

87. Coracoid, sternal end, internal lip expansion (Fig. S63):

0 – expanded along entire width

1 – only expanded in medial portion

2 – only expanded in lateral portion

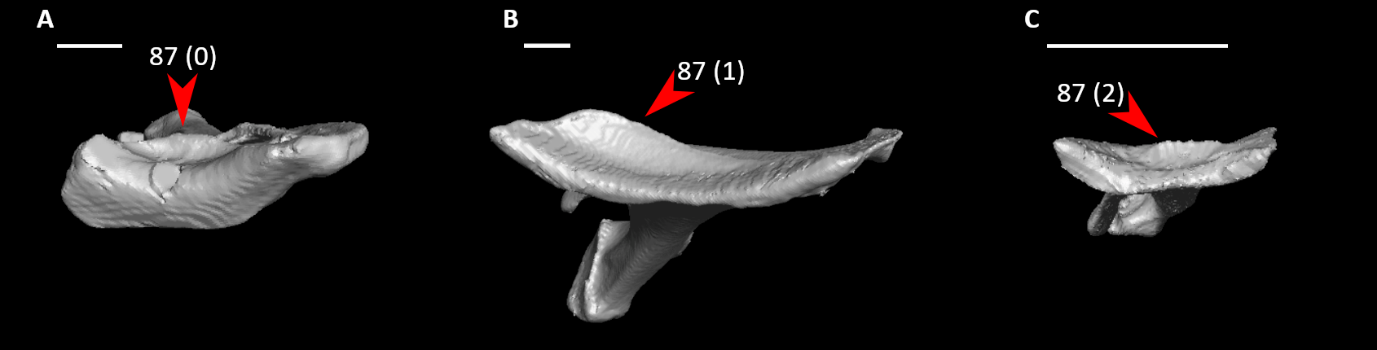

**Figure S63.** Coracoids of *Alectura lathami* (**A**, left element, mirrored), *Phoenicopterus ruber* (B, left element, mirrored), and *Aegotheles cristatus* (**B**, right element) in sternal view, illustrating alternative states for character 87 (complete character description in text). Arrows indicate the internal lip of the articulation facet with the sternum. Scale bars = 5 mm.

88. Scapula, acromion in dorsal view (Fig. S64; Ksepka et al., 2019: 53):

0 – single

1 – bifurcated, with an additional medial process

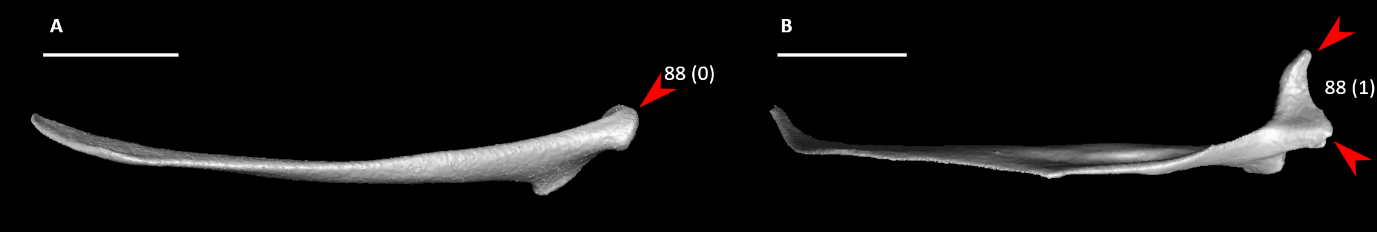

**Figure S64.** Scapulae of *Aegotheles cristatus* (**A**, right element) and *Alcedo atthis* (**B**, right element) in dorsal view, illustrating alternative states for character 88 (complete character description in text). Arrows indicate the tip(s) of the acromion process. Scale bars = 5 mm.

89. Scapula, acromion, cranial extent with scapular neck horizontal (Fig. S65; Livezey, 1986: 109; Livezey, 1996: 38; Ericson, 1997: 48; Livezey and Zusi, 2006: 1245; Worthy and Lee, 2008: 39; Smith, 2010: 155; Worthy et al., 2017: 89):

0 – subequal or caudal to coracoid tubercle

1 – extends distinctly craniad of coracoid tubercle

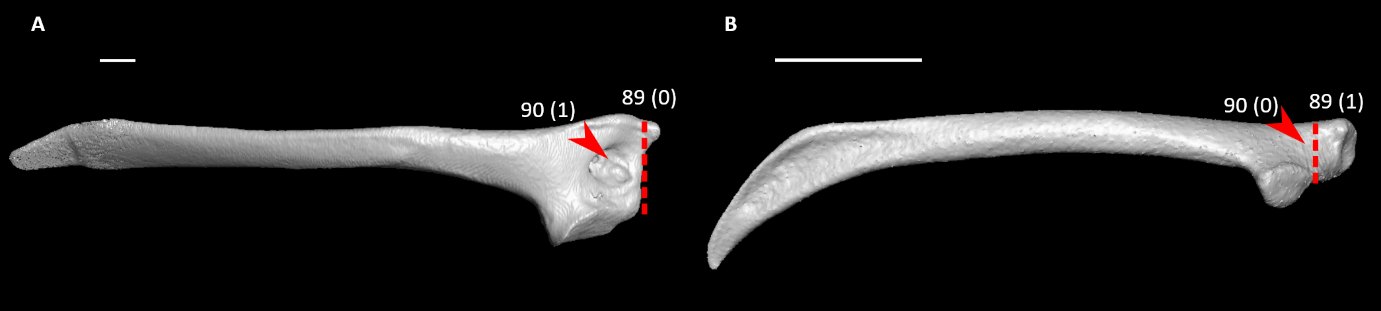

**Figure S65.** Scapulae of *Chauna chavaria* (**A**, left element, mirrored) and *Aegotheles cristatus* (**B**, right element) in lateral view, illustrating alternative states for characters 89 and 90 (complete character descriptions in text). Dotted lines indicate the cranial extent of the coracoid tubercle to show relative cranial projection of the acromion process. Arrows indicate the pneumatic foramen in the cranial end, or approximate homologous site to the foramen. Scale bars = 5 mm.

90. Scapula, cranial end, pneumatic foramen (Livezey, 1986: 111; Livezey, 1996: 37; Worthy and Lee, 2008: 40; Ksepka, 2009: 46; Worthy et al., 2017: 90):

0 – absent

1 – present

91. Scapula, ventral surface under articular facet for humerus, pneumatic fossa (Fig. S66; Worthy et al., 2017: 91):

0 – absent

1 – present

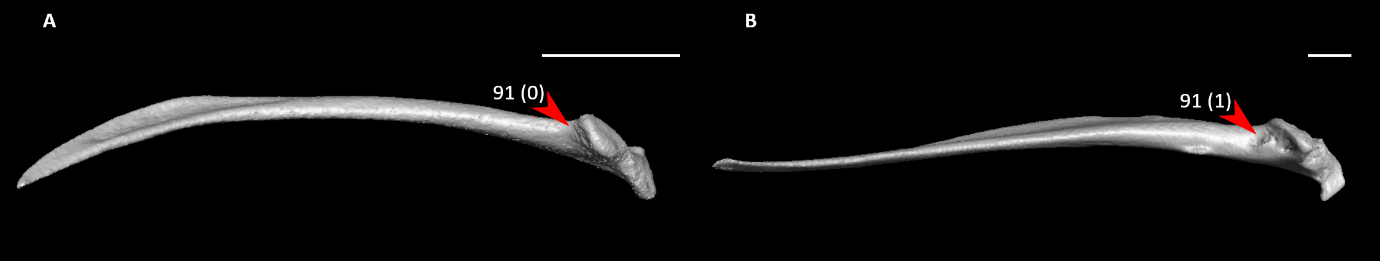

**Figure S66.** Scapulae of *Aegotheles cristatus* (**A**, right element) and *Alectura lathami* (**B**, left element, mirrored) in ventral view, illustrating alternative states for character 91 (complete character description in text). Arrows indicate the ventral surface under articular facet for humerus. Scale bars = 5 mm.

92. Scapula, shaft, monotonic ventral curvature (Fig. S67; Livezey and Zusi, 2006: 1260; Musser and Cracraft, 2019: 207; Musser and Clarke, 2020: 436):

0 – moderate, main body and distal margin of scapula is slightly to moderately convex

1 – pronounced, main body and distal margin of scapula conspicuously convex

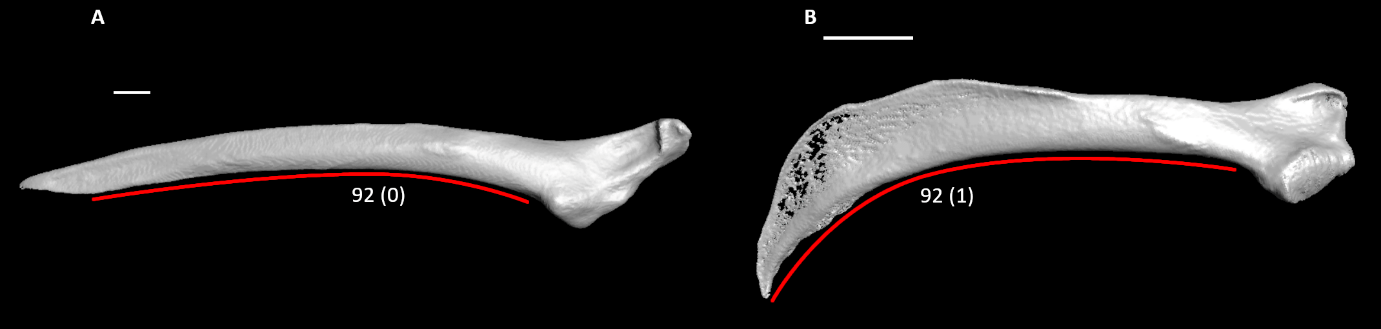

**Figure S67.** Scapulae of *Leucocarbo atriceps* (**A**, left element, mirrored) and *Nyctibius griseus* (**B**, left element, mirrored) in lateral view, illustrating alternative states for character 92 (complete character description in text). Red lines indicate the approximate ventral curvature of the scapular shaft. Scale bars = 5 mm.

93. Scapula, shaft, lateral view, longitudinal concavity (Fig. S68; Livezey and Zusi, 2006: 1257; Musser and Cracraft, 2019: 208; Musser and Clarke, 2020: 437):

0 – absent or weak, essentially planar throughout or shallow concavity limited to cranial and medial portions

1 – prominent, distinctly concave throughout, accented by lateral displacement of dorsal margin of shaft

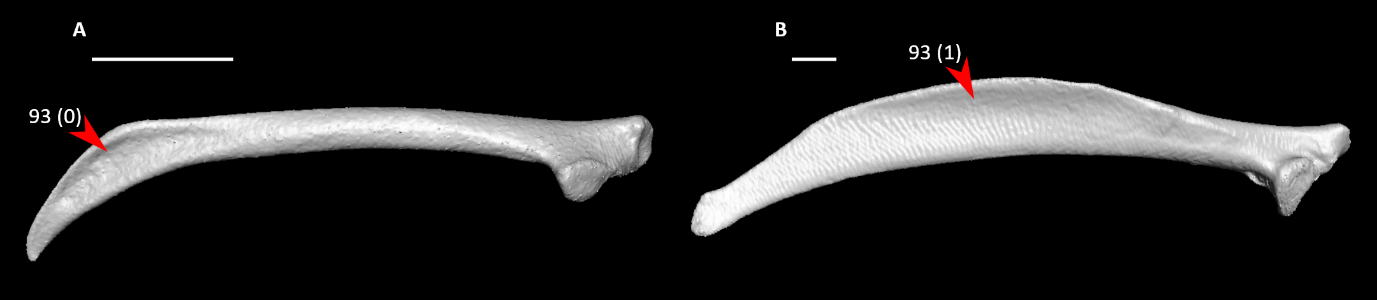

**Figure S68.** Scapulae of *Aegotheles cristatus* (**A**, right element) and *Alectura lathami* (**B**, left element, mirrored) in lateral view, illustrating alternative states for character 93 (complete character description in text). Arrows indicate the longitudinal concavity. Scale bars = 5 mm.

94. Scapula, shaft, lateral view, mound-shaped tuberosity on lateral surface (Fig. S69):

0 – absent

1 – present

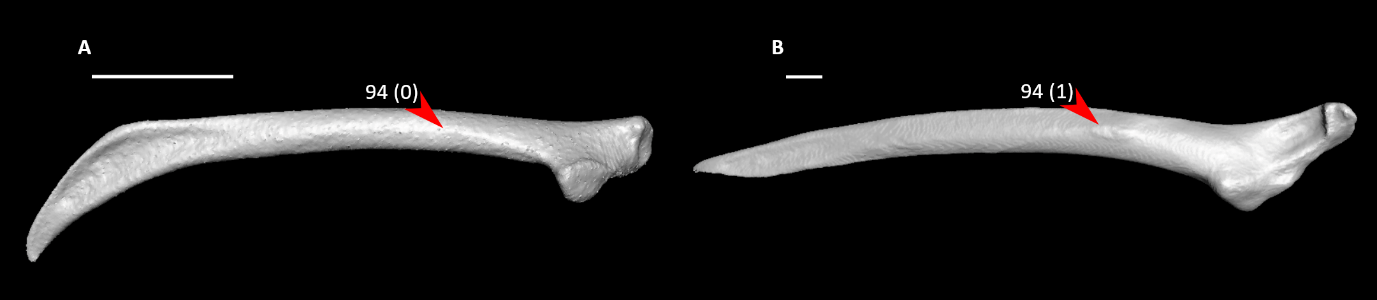

**Figure S69.** Scapulae of *Aegotheles cristatus* (**A**, right element) and *Leucocarbo atriceps* (**B**, left element, mirrored) in lateral view, illustrating alternative states for character 94 (complete character description in text). Arrows indicate the lateral surface and its mound-shaped tuberosity (when present). Scale bars = 5 mm.

95. Scapula, shaft, lateral view, shape of distal portion (Fig. S70; Livezey and Zusi, 2006: 1264; Musser and Cracraft, 2019: 210; Musser and Clarke, 2020: 442):

0 – widened dorsocaudally, spatulate

1 – long, bladelike, compressed dorsocaudally

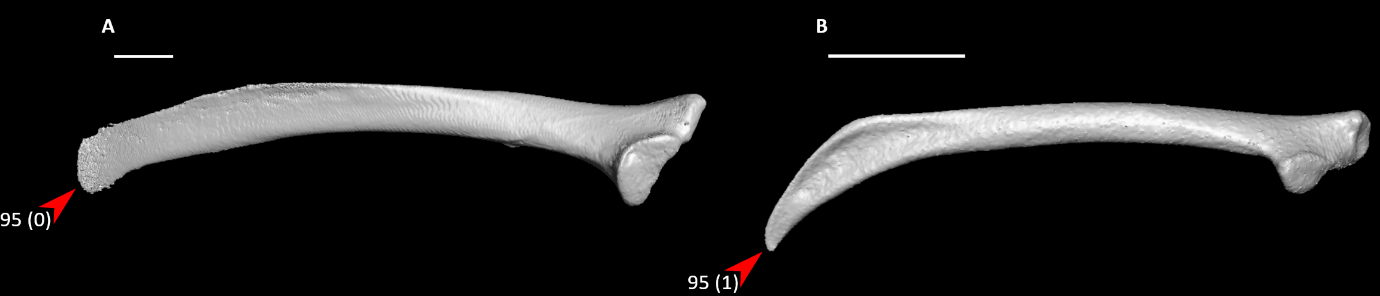

**Figure S70.** Scapulae of *Eudromia elegans* (**A**, left element, mirrored) and *Aegotheles cristatus* (**B**, right element) in lateral view, illustrating alternative states for character 95 (complete character description in text). Arrows indicate the distal tip of the scapula. Scale bars = 5 mm.

96. Scapula, shaft, dorsal view, lateral curvature (Fig. S71):

0 – straight or slight

1 – strong curvature

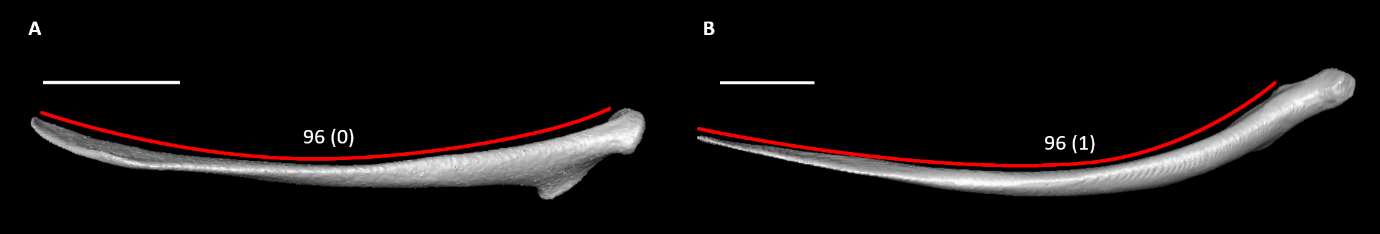

**Figure S71.** Scapulae of *Aegotheles cristatus* (**A**, right element) and *Podilymbus podiceps* (**B**, right element) in dorsal view, illustrating alternative states for character 96 (complete character description in text). Red lines indicate the approximate lateral curvature of the scapular shaft. Scale bars = 5 mm.

*97. Proportions of humerus relative to total length of [humerus + ulna + carpometacarpus] (Appendix S2):

0 – less than 30% total length

1 – between 30-40% total length

2 – greater than 40% total length

Estimated for *Ichthyornis* by cross-scaling different elements among specimens described by Benito et al. (2022).

98. Proportions of ulna relative to total length of [humerus + ulna + carpometacarpus] (Appendix S2):

0 – less than or equal to 40% total length

1 – greater than 40% total length

99. Proportions of carpometacarpus relative to total length of [humerus + ulna + carpometacarpus] (Appendix S2):

0 – less than or equal to 25% total length

1 – greater than 25% total length

100. Humerus, proximal end, capital incisure form (Fig. S72; Livezey and Zusi, 2006: 1358: Musser and Cracraft, 2019: 215; Musser and Clarke, 2020: 448):

0 – shallow

1 – deep and prominent

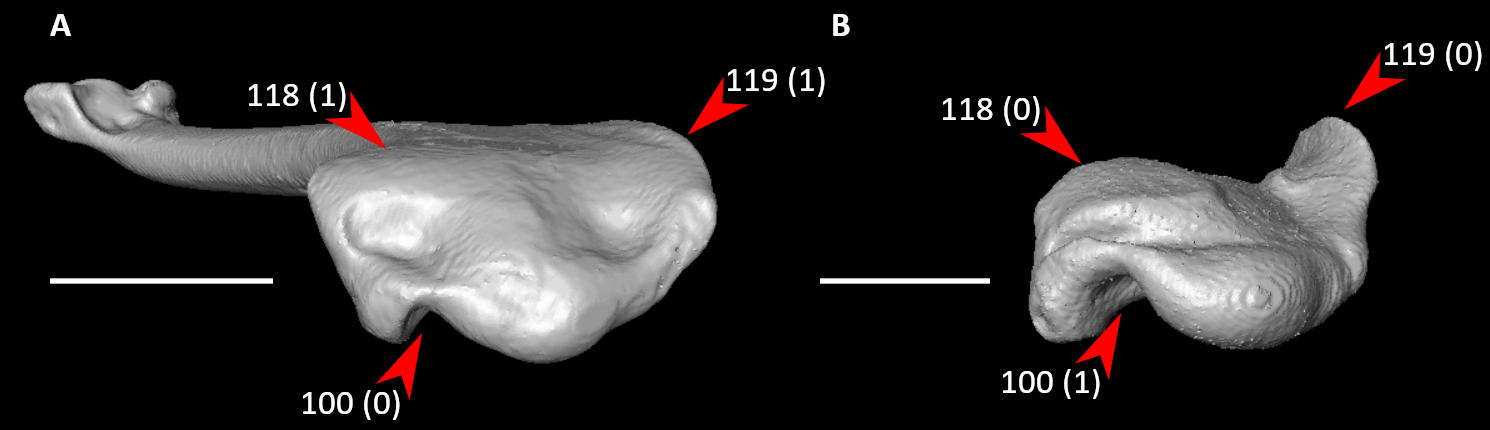

**Figure S72.** Humeri of *Jynx torquilla* (**A**, right element) and *Aegotheles cristatus* (**B**, right element) in proximal view, illustrating alternative states for characters 100, 118, and 119 (complete character descriptions in text). Arrows for character 100 indicate the capital incisure, arrows for character 118 indicate the cranial surface of the bicipital crest, and arrows from character 119 indicate the deltopectoral crest. Scale bars = 5 mm.

101. Humerus, proximal end, capital incisure (Fig. S73; Musser et al., 2019: 63; Musser and Clarke, 2020: 449):

0 – not as follows

1 – enclosed by distal projection of humeral head

2 – closed by transverse ridge

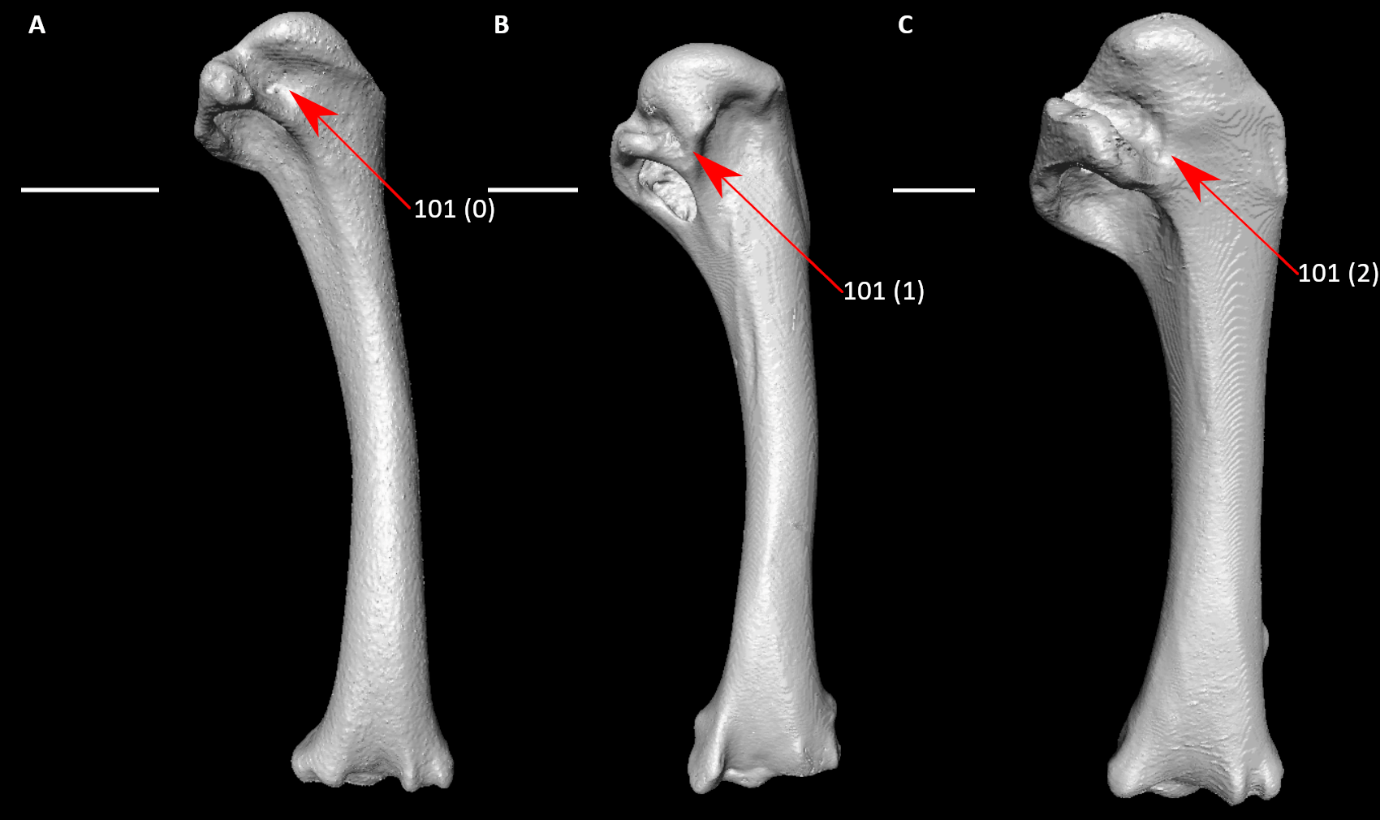

**Figure S73.** Humeri of *Aegotheles cristatus* (**A**, right element), *Rollulus rouloul* (**B**, right element), and *Columba livia* (**C**, left element, mirrored) in caudal view, illustrating alternative states for character 101 (complete character description in text). Arrows indicate the distal end of the capital incisure. Scale bars = 5 mm.

102. Humerus, proximal end, ventral tubercle (Fig. S74; Livezey and Zusi, 2006: 1365; Musser and Cracraft, 2019: 211; Musser and Clarke, 2020: 445):

0 – not very prominent

1 – prominent, projecting well beyond rest of bone caudally

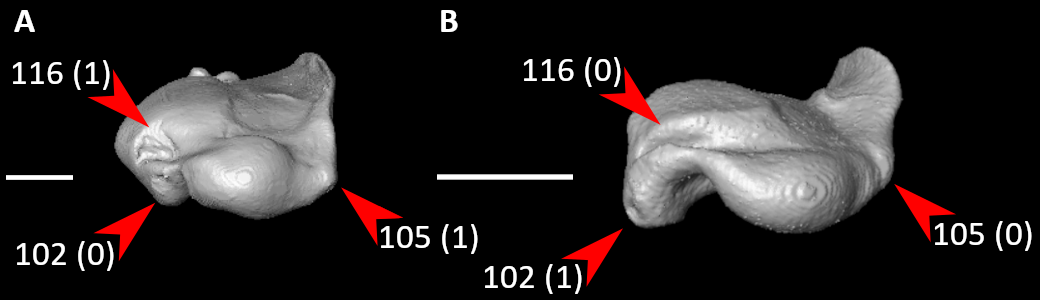

**Figure S74.** Humeri of *Podilymbus podiceps* (**A**, right element) and *Aegotheles cristatus* (**B**, right element) in proximal view, illustrating alternative states for characters 102, 105, and 116 (complete character descriptions in text). Arrows for character 102 indicate the ventral tubercle, arrows for character 105 indicate the dorsal tubercle, and arrows for character 116 indicate the transverse sulcus. Scale bars = 5 mm.

103. Humerus, proximal end, ventral tubercle, proximodistal position relative to pneumotricipital fossa or homologous site (Fig. S75; Livezey and Zusi, 2006: 1366; Musser and Cracraft, 2019: 212):

0 – comparatively cranioproximal, completely exposing pneumotricipital fossa

1 – comparatively caudodistal, largely or completely concealing pneumotricipital fossa

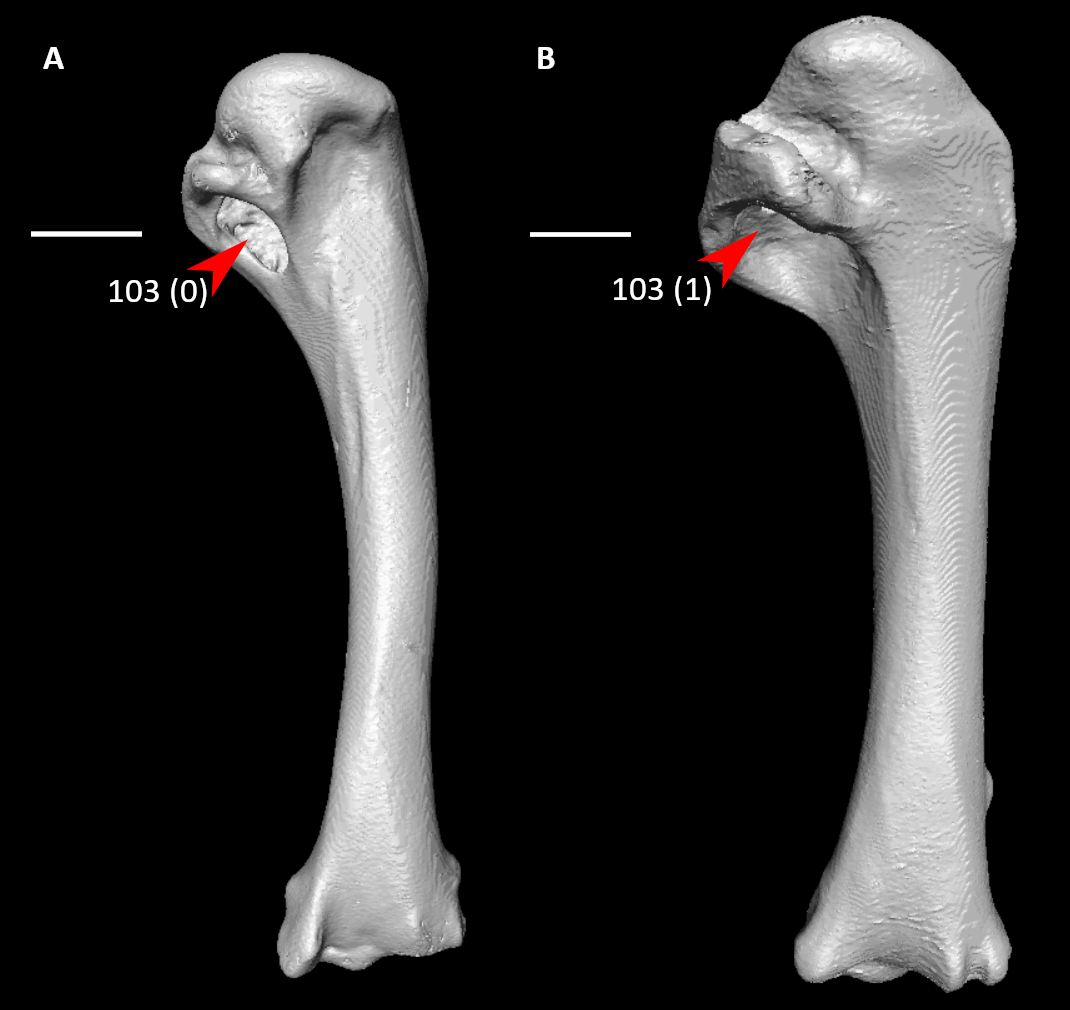

**Figure S75.** Humeri of *Rollulus rouloul* (**A**, right element) and *Columba livia* (**B**, left element, mirrored) in caudal view, illustrating alternative states for character 103 (complete character description in text). Arrows indicate the pneumotricipital fossa. Scale bars = 5 mm.

*104. Humerus, proximal end, ventral tubercle, proximal elevation regardless of pneumotricipital fossa exposure (Musser and Cracraft, 2019: 213; Musser and Clarke, 2020: 446):

0 – inferior to dorsal tubercle

1 – subequal in elevation to that of dorsal tubercle

2 – elevated immediately proximally to dorsal tubercle

3 – elevated well proximal to dorsal tubercle, proximal to humeral head

Possibly taphonomic in *Ichthyornis*.

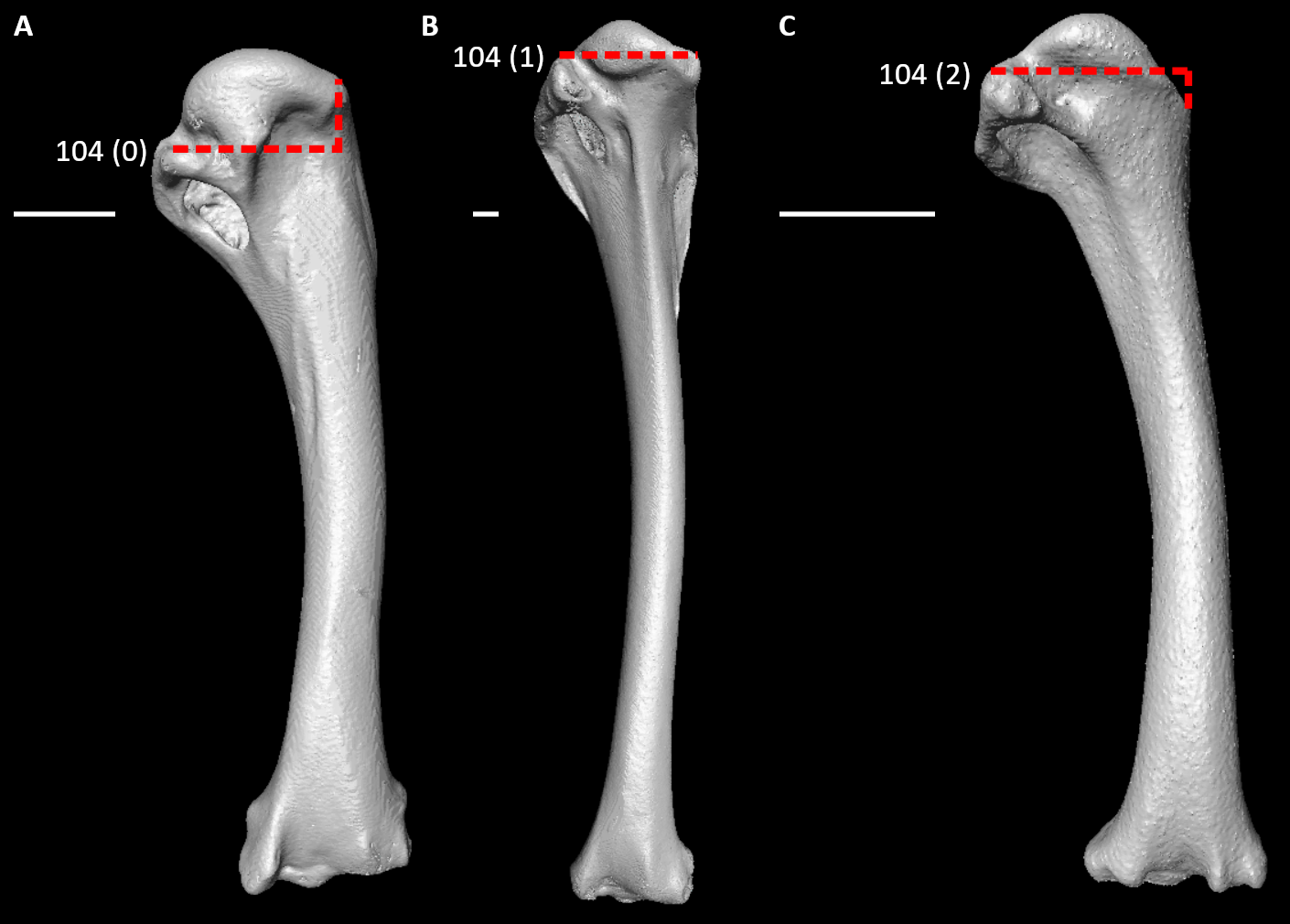

**Figure S76.** Humeri of *Rollulus rouloul* (**A**, right element), *Anseranas semipalmata* (**B**, left element, mirrored), and *Aegotheles cristatus* (**C**, right element) in caudal view, illustrating alternative states for character 104 (complete character description in text). Dotted lines indicate the proximal extent of the ventral tubercle relative to that of the dorsal tubercle. Scale bars = 5 mm.

105. Humerus, proximal end, dorsal tubercle (Fig. S74; Livezey and Zusi, 2006: 1370; Musser and Cracraft, 2019: 214):

0 – smooth and rounded

1 – pointed

106. Humerus, proximal end, caudal view, capital shaft ridge (Fig. 77; Worthy and Lee, 2008: 51; Worthy et al., 2017: 115; creates triangular cross section of Musser and Cracraft, 2019: 232, Musser et al., 2019: 67, and Musser and Clarke, 2020: 467):

0 – absent

1 – present

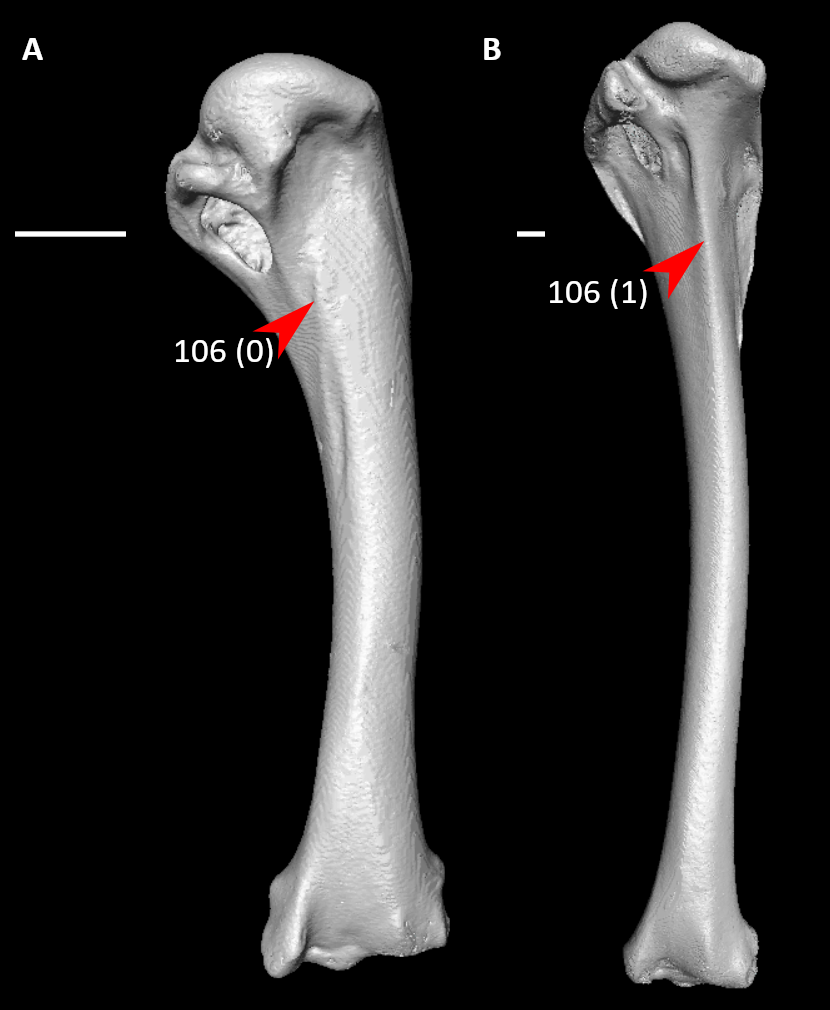

**Figure S77.** Humeri of *Rollulus rouloul* (**A**, right element) and *Anseranas semipalmata* (**B**, left element, mirrored) in caudal view, illustrating alternative states for character 106 (complete character description in text). Arrows indicate the capital shaft ridge or approximate homologous site. Scale bars = 5 mm.

107. Humerus, proximal end, caudal view, capital shaft orientation (Fig. S78; Livezey, 1986: 22; Livezey, 2996: 51; Worthy and Lee, 2008: 52; Worthy et al., 2017: 116):

0 – directed towards humeral head

1 – directed towards zone between head and dorsal tubercle

2 – directed towards dorsal tubercle

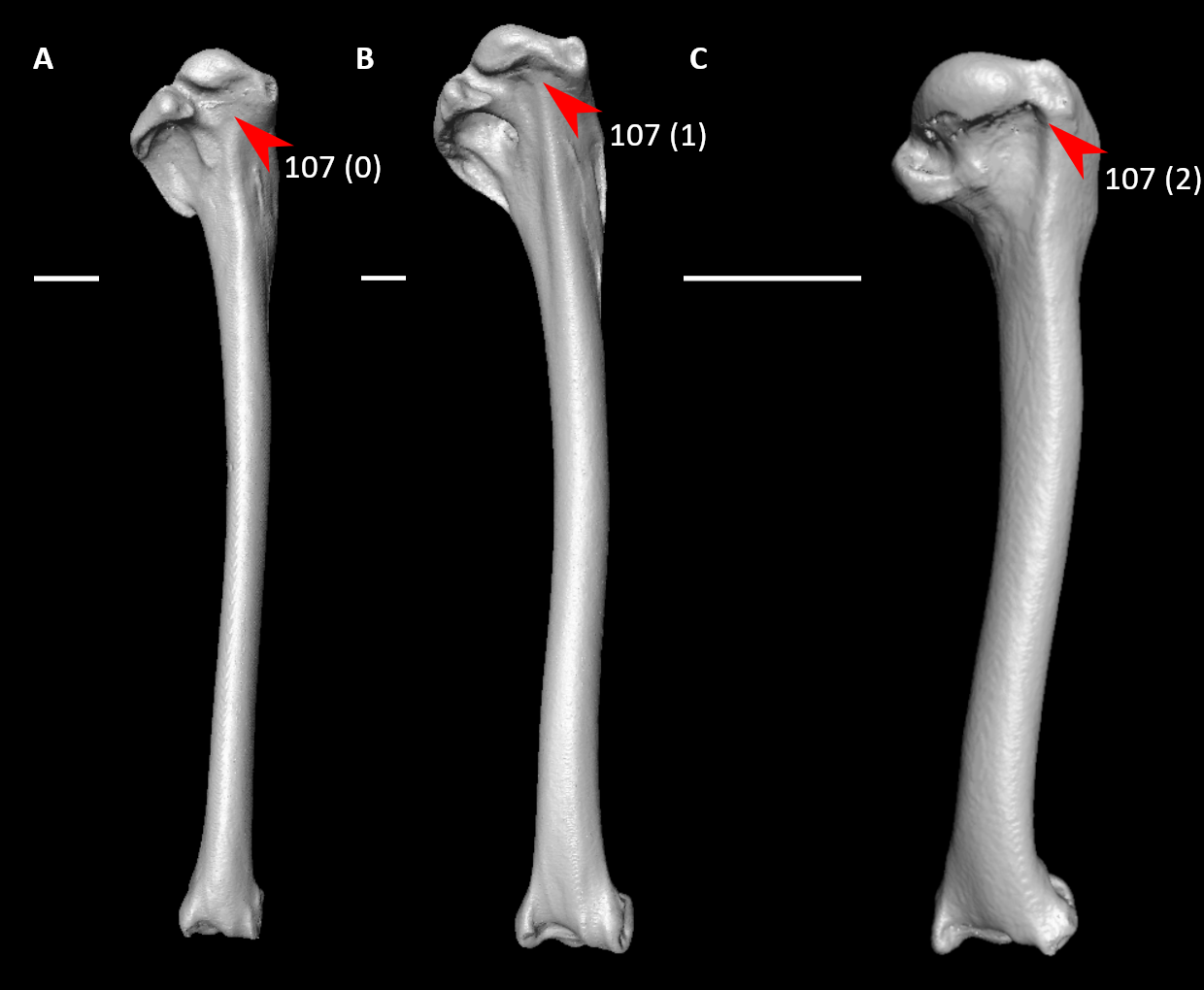

**Figure S78.** Humeri of *Podilymbus podiceps* (**A**, right element), *Dendrocygna bicolor* (**B**, right element), and *Alcedo atthis* (**C**, left element, mirrored) in caudal view, illustrating alternative states for character 107 (complete character description in text). Arrows indicate the proximal end of the capital shaft ridge. Scale bars = 5 mm.

108. Humerus, proximal end, caudal view, marked oval depression at insertion site of m. scapulohumeralis cranialis (Fig. S79; Mayr, 2004b: 41):

0 – absent

1 – present

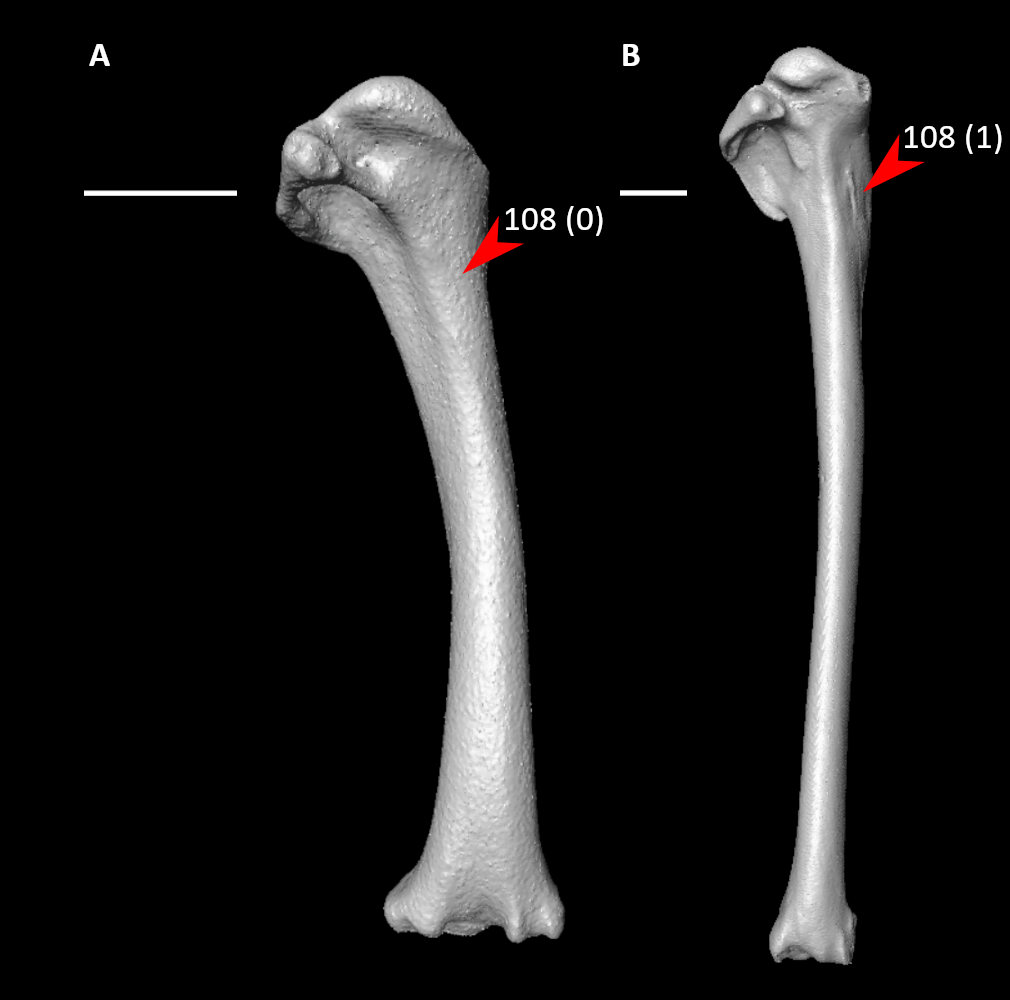

**Figure S79.** Humeri of *Aegotheles cristatus* (**A**, right element) and *Podilymbus podiceps* (**B**, right element) in caudal view, illustrating alternative states for character 108 (complete character description in text). Arrows indicate the depression at insertion site of m. scapulohumeralis cranialis or approximate homologous site. Scale bars = 5 mm.

109. Humerus, proximal end, caudal view, foramen in pneumotricipital fossa (Fig. S80; Mayr and Clarke, 2003: 77; Livezey and Zusi, 2006: 1414; Musser and Cracraft, 2019: 216; Musser and Clarke, 2020: 450):

0 – apneumatic

1 – deep and pneumatic

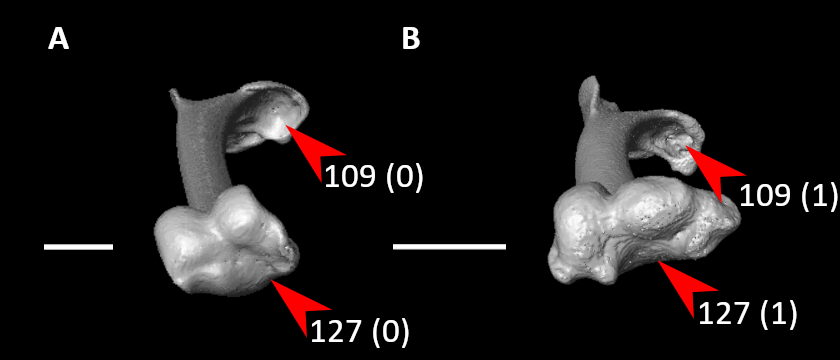

**Figure S80.** Humeri of *Podilymbus podiceps* (**A**, right element) and *Aegotheles cristatus* (**B**, right element) in distal view, illustrating alternative states for characters 109 and 127 (complete character descriptions in text). Arrows for character 109 indicate the pneumotricipital fossa and its pneumatic foramen (when present), whereas arrows for character 127 indicate the olecranon fossa. Scale bars = 5 mm.

110. Humerus, proximal end, caudal view, pneumotricipital fossa, dorsal ramus (Fig. S81; Livezey, 1998: 202; Musser and Cracraft, 2019: 219; Musser and Clarke, 2020: 453):

0 – dorsoventrally narrower than ventral ramus

1 – dorsoventrally equal or deeper than ventral ramus

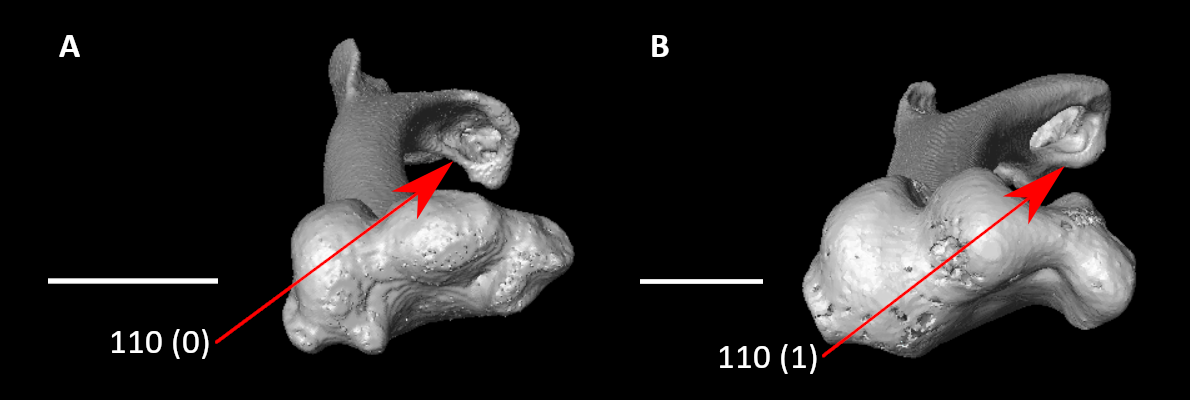

**Figure S81.** Humeri of *Aegotheles cristatus* (**A**, right element) and *Rollulus rouloul* (**B**, right element) in distal view, illustrating alternative states for character 110 (complete character description in text). Arrows indicate the dorsal ramus of the pneumotricipital fossa. Scale bars = 5 mm.

111. Humerus, proximal end, caudal view, second (dorsal) pneumotricipital fossa (Fig. S82; Musser et al., 2019: 64; Musser and Clarke, 2020: 451):

0 – absent

1 – present

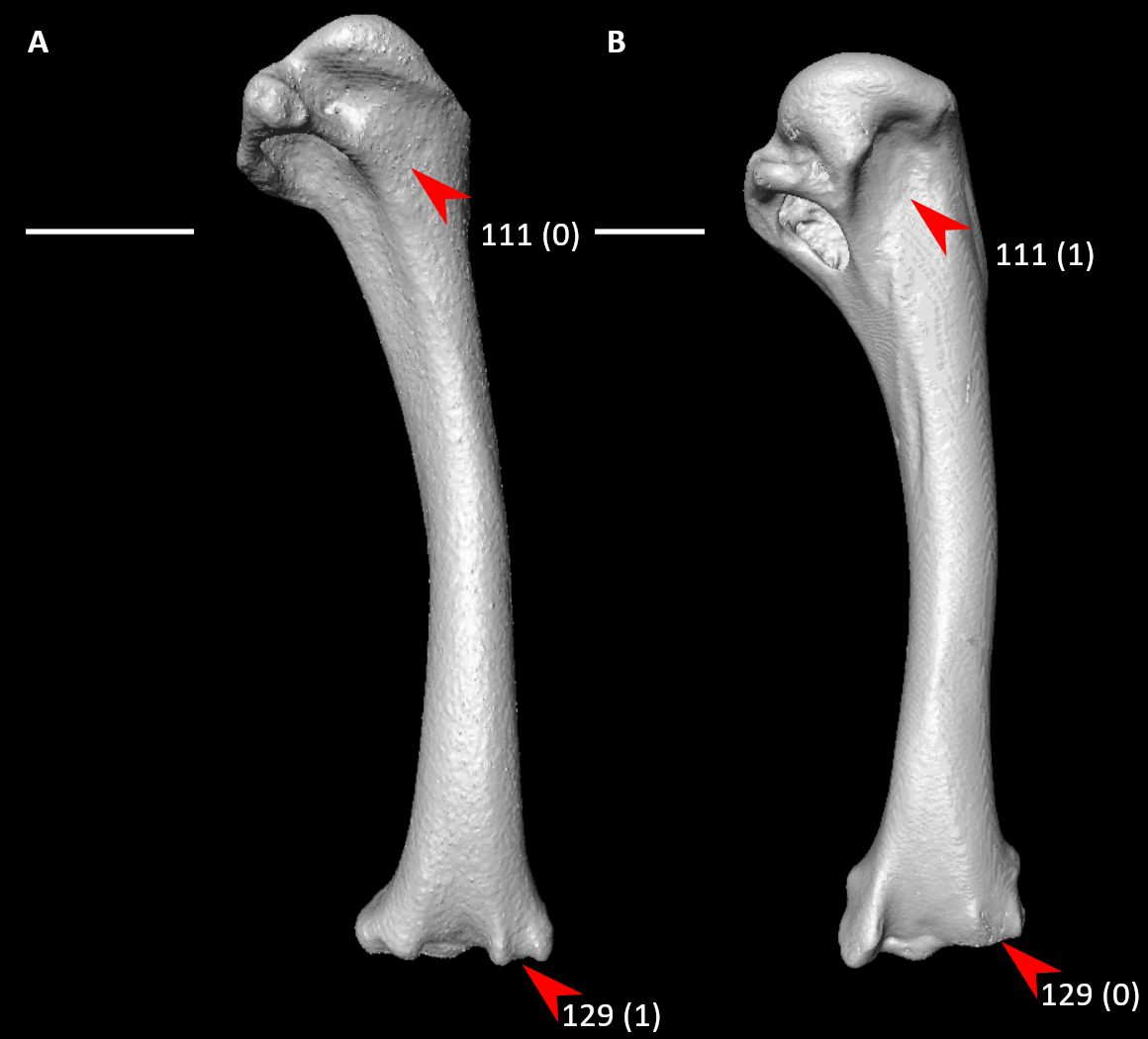

**Figure S82.** Humeri of *Aegotheles cristatus* (**A**, right element) and *Rollulus rouloul* (**B**, right element) in caudal view, illustrating alternative states for characters 111 and 129 (complete character descriptions in text). Arrows for character 111 indicate the dorsal pneumotricipital fossa or approximate homologous site, whereas arrows for character 129 indicate the tendinal sulcus of m. scapulotricipitalis. Scale bars = 5 mm.

112. Humerus, proximal end, caudal view, dorsal pneumotricipital fossa form (Fig. S83; Worthy and Lee, 2008: 53; Ksepka, 2009: 55; Worthy et al., 2017: 118):

0 – narrower than ventral pneumotricipital fossa

1 – wider than ventral pneumotricipital fossa

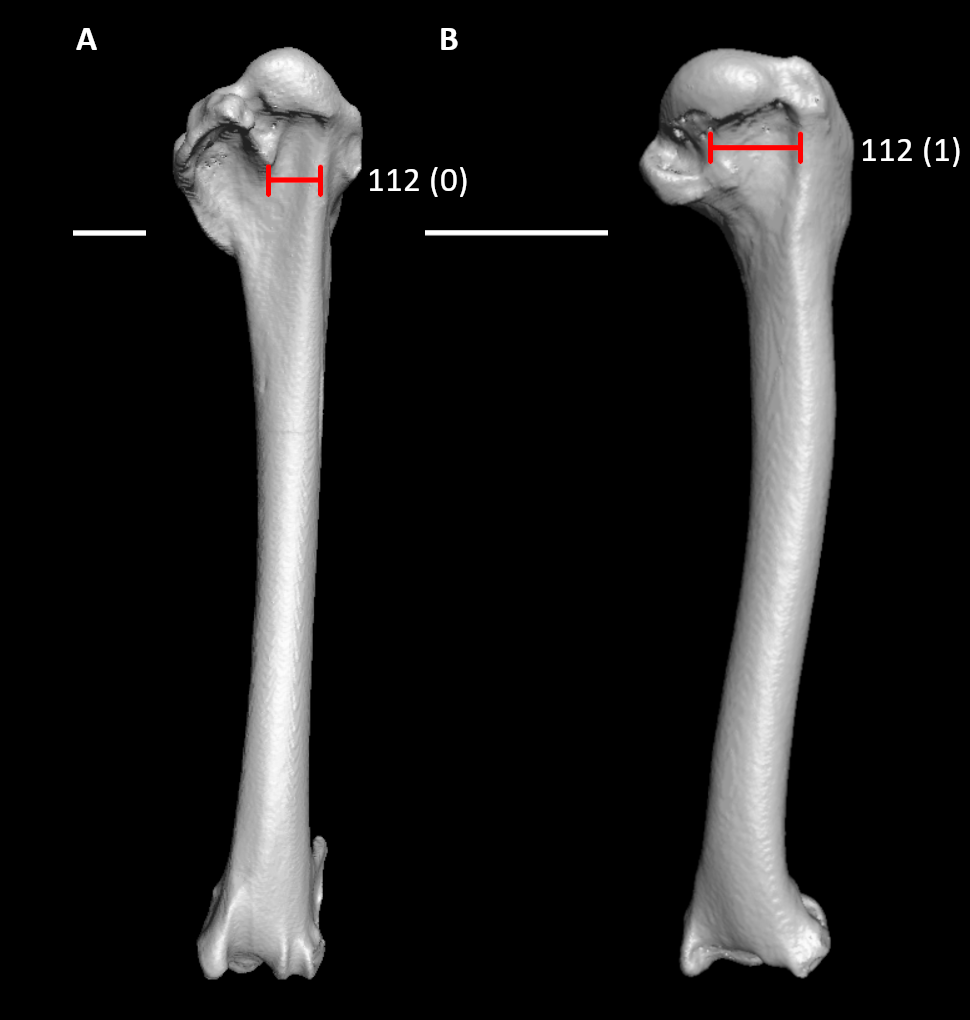

**Figure S83.** Humeri of *Limosa lapponica* (**A**, left element, mirrored) and *Alcedo atthis* (**B**, left element, mirrored) in caudal view, illustrating alternative states for character 1112 (complete character description in text). Red lines span the width of the dorsal pneumotricipital fossa. Scale bars = 5 mm.

113. Humerus, proximal end, cranial view, intertubercular plane, coracobrachial nerve sulcus (Fig. S84; Livezey and Zusi, 2006: 1428; Musser and Cracraft, 2019: 222; Musser and Clarke, 2020: 457):

0 – absent or weak

1 – prominent

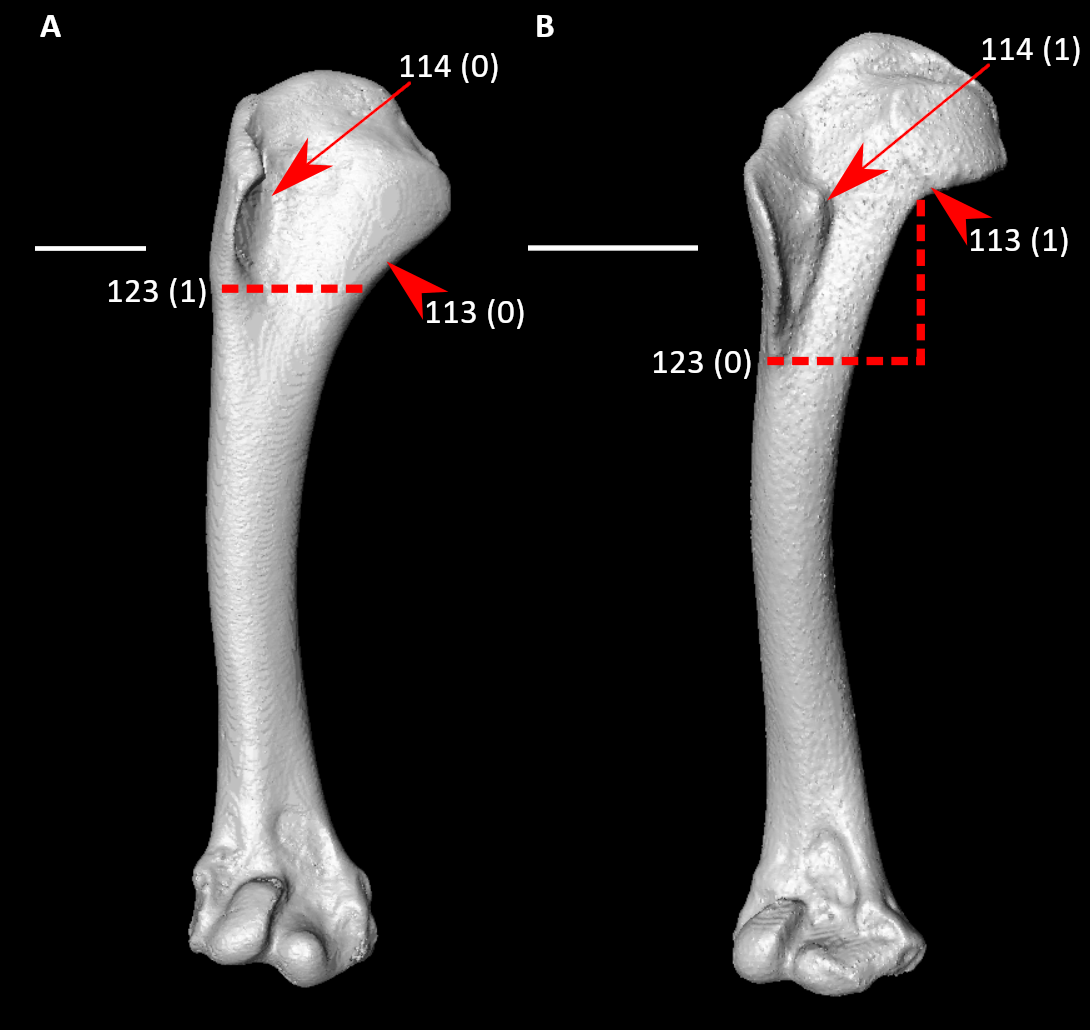

**Figure S84.** Humeri of *Rollulus rouloul* (**A**, right element) and *Aegotheles cristatus* (**B**, right element) in cranial view, illustrating alternative states for characters 113, 114, and 123 (complete character descriptions in text). Arrows for character 113 indicate the coracobrachial nerve sulcus or approximate homologous site, whereas arrows for character 114 indicate the coracobrachial impression or approximate homologous site. Dotted lines indicate the distal extent of the deltopectoral crest relative to that of the bicipital crest. Scale bars = 5 mm.

114. Humerus, proximal end, cranial view, coracobrachial impression (Fig. S84; Mayr, 2011: 31):

0 – absent or weak

1 – prominent, forming sharp edge on dorsal margin of humeral intumescence

*115. Humerus, proximal end, cranial view, dorsoventral extent of transverse ligamental sulcus (Fig. S85; Livezey and Zusi, 2006: 1431; Musser and Cracraft, 2019: 223; Musser and Clarke, 2020: 458):

0 – extremely short, at most suggested by short, shallow depression or dorsally restricted pit

1 – intermediate, typically limited to roughly entire proximal margin of bicipital surface and reaching midpoint of the proximal portion of the humerus cranial surface

2 – long, extends dorsoventrally across proximal portion of humerus to intersect bases of dorsal and ventral tubercles

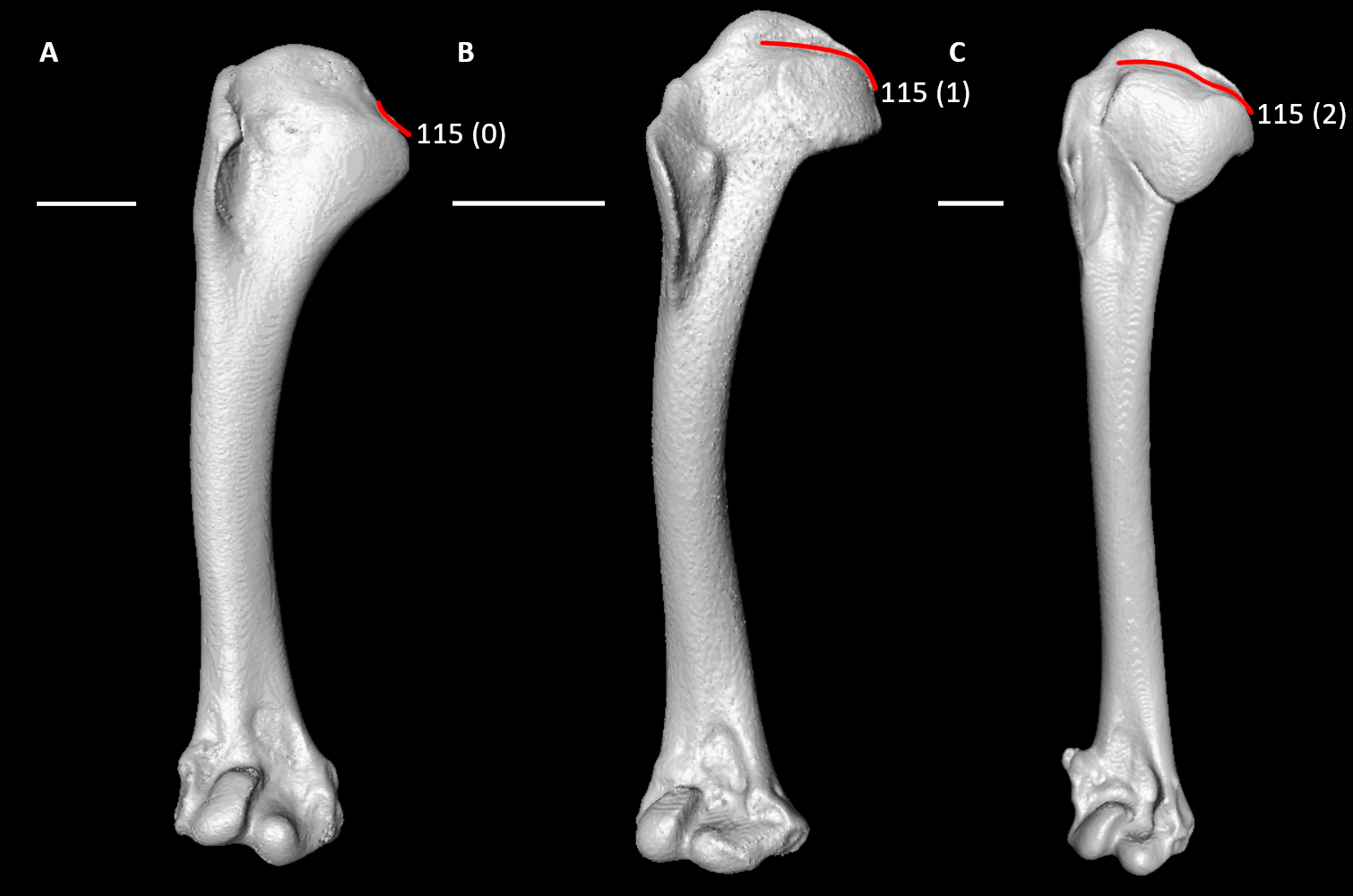

**Figure S85.** Humeri of *Rollulus rouloul* (**A**, right element), *Aegotheles cristatus* (**B**, right element), and *Limosa lapponica* (**C**, left element, mirrored) in cranial view, illustrating alternative states for character 115 (complete character description in text). Red lines indicate the dorsoventral extent of the transverse sulcus. Scale bars = 5 mm.

116. Humerus, proximal end, cranial view, ventral section of transverse ligament sulcus, marked triangular raised subplanar region delimited by pronounced marginal crests enclosing deep ventral pit (Fig. S74; Livezey and Zusi, 2006: 1432; Musser and Cracraft, 2019: 224):

0 – absent

1 – present

117. Humerus, proximal end, bicipital crest (Fig. S86):

0 – inconspicuous

1 – prominent, protruding ventrally far beyond ventral condyle

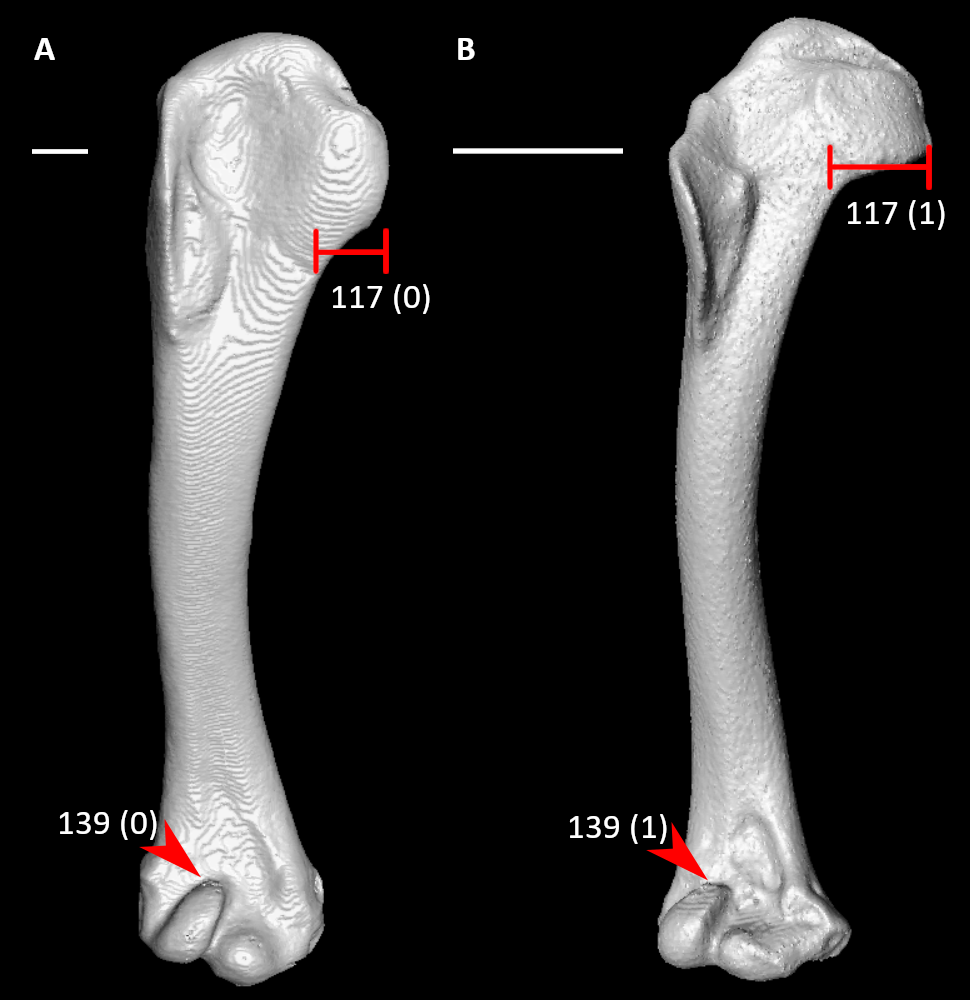

**Figure S86.** Humeri of *Alectura lathami* (**A**, left element, mirrored) and *Aegotheles cristatus* (**B**, right element) in cranial view, illustrating alternative states for characters 117 and 139 (complete character descriptions in text). Red lines span the dorsoventral width of the bicipital crest. Arrows indicate the proximal margin of the dorsal condyle. Scale bars = 5 mm.

118. Humerus, proximal end, bicipital crest, cranial surface in proximal view (Fig. S72; Livezey and Zusi, 2006: 1405; Musser and Cracraft, 2019: 226):

0 – convex

1 – planar or slightly concave

119. Humerus, proximal end, deltopectoral crest orientation (Fig. S72):

0 – primarily flared cranially

1 – primarily flared dorsally

Scored state 1 in *Ichthyornis*; is flared slightly cranially, but less so than in most modern birds.

120. Humerus, proximal end, deltopectoral crest (Fig. S87; Nesbitt and Clarke, 2016: 114):

0 – less than width of shaft

1 – same width as shaft or greater

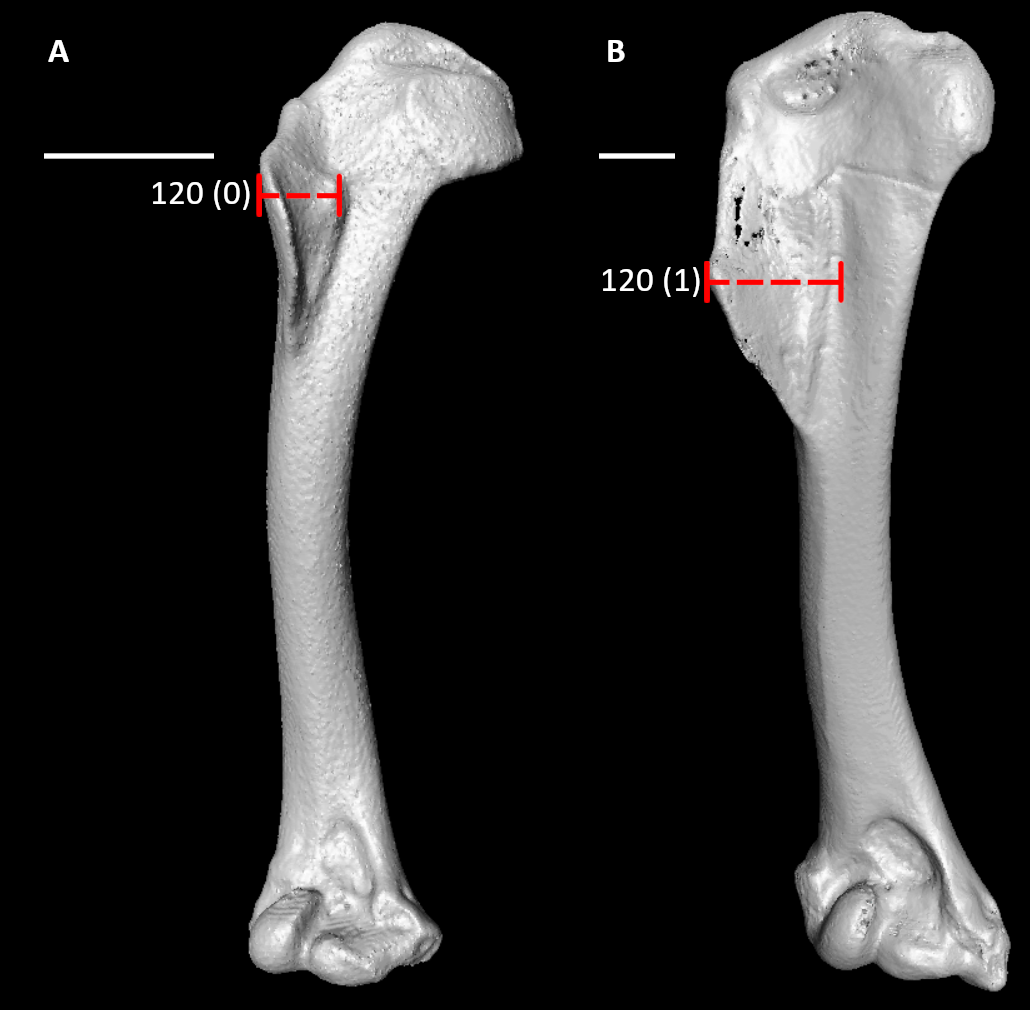

**Figure S87.** Humeri of *Aegotheles cristatus* (**A**, right element) and *Corythaeola cristata* (**B**, left element, mirrored) in cranial view, illustrating alternative states for character 120 (complete character description in text). Dotted lines span the approximate dorsoventral width of the deltopectoral crest. Scale bars = 5 mm.

121. Humerus, proximal end, deltopectoral crest shape (Fig. S88; Livezey and Zusi, 2006: 1374; Musser and Cracraft, 2019: 229; Musser and Clarke, 2020: 464):

0 – rounded dorsally

1 – straight dorsally

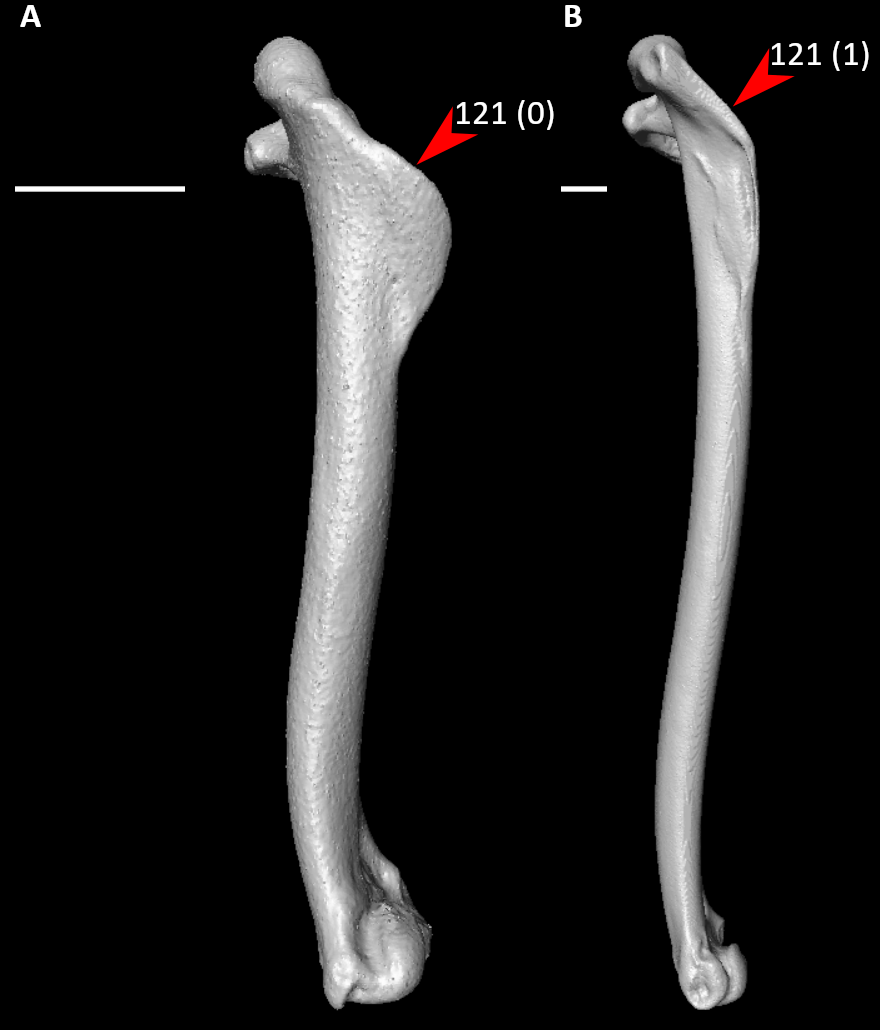

**Figure S88.** Humeri of *Aegotheles cristatus* (**A**, right element) and *Dendrocygna bicolor* (**B**, right element) in dorsal view, illustrating alternative states for character 121 (complete character description in text). Arrows indicate the dorsal margin of the deltopectoral crest. Scale bars = 5 mm.

122. Humerus, proximal end, deltopectoral crest, proximodistal length relative to main body of humerus (Fig. S89; Livezey and Zusi, 2006: 1382; Musser and Cracraft, 2019: 230; Musser and Clarke, 2020: 465):

0 – well developed, extending at least one third length of humerus

1 – small eminence, extending less than one third length of humerus

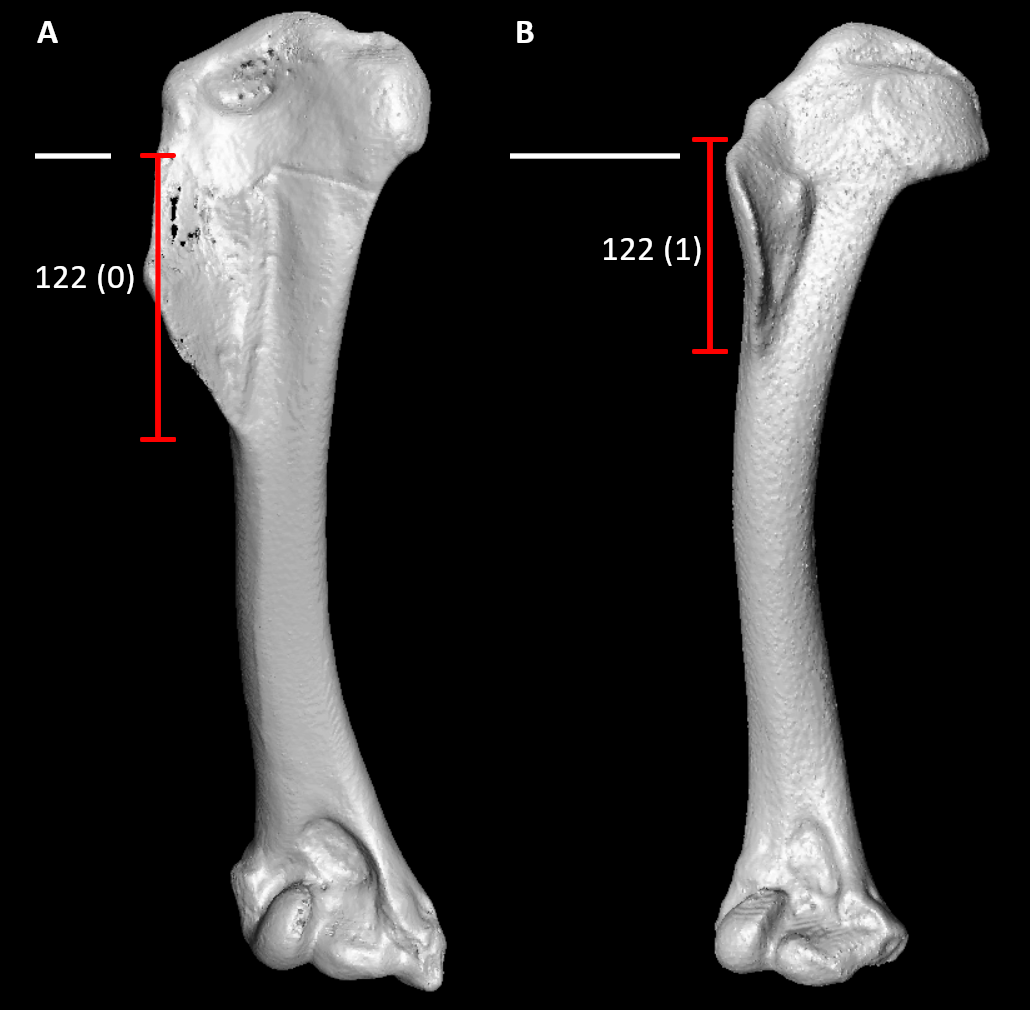

**Figure S89.** Humeri of *Corythaeola cristata* (**A**, left element, mirrored) and *Aegotheles cristatus* (**B**, right element) in cranial view, illustrating alternative states for character 121 (complete character description in text). Red lines span the proximodistal length of the deltopectoral crest. Scale bars = 5 mm.

123. Humerus, proximal end, deltopectoral crest, oblique caudal view, proximodistal extent relative to that of bicipital crest (Fig. S84; Livezey and Zusi, 2006: 1383; Musser and Cracraft, 2019: 231):

0 – terminates distally to bicipital crest

1 – approximately subequal to bicipital crest

124. Humerus, proximal end, deltopectoral crest, caudal view (Fig. S90; Mayr, 2011: 32):

0 – concave

1 – convex

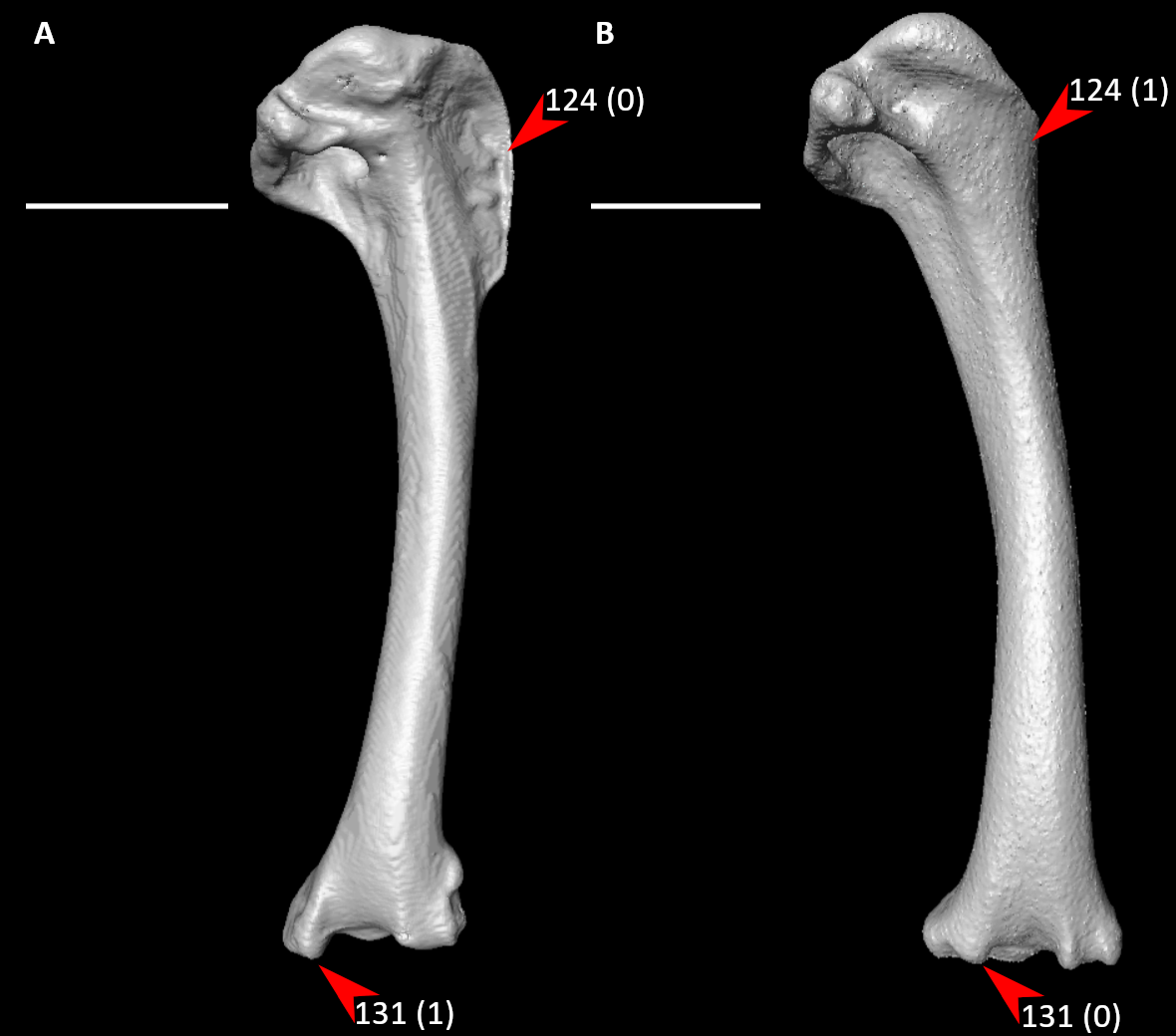

**Figure S90.** Humeri of *Aegotheles cristatus* (**A**, right element) and *Rollulus rouloul* (**B**, right element) in caudal view, illustrating alternative states for characters 124 and 131 (complete character descriptions in text). Arrows for character 124 indicate the caudal surface of the deltopectoral crest, whereas character 131 indicate the flexor process. Scale bars = 5 mm.

125. Humerus, shaft, dorsoventral curvature distal to deltopectoral crest (Fig. S91; Ksepka et al., 2019: 58):

0 – curved

1 – straight

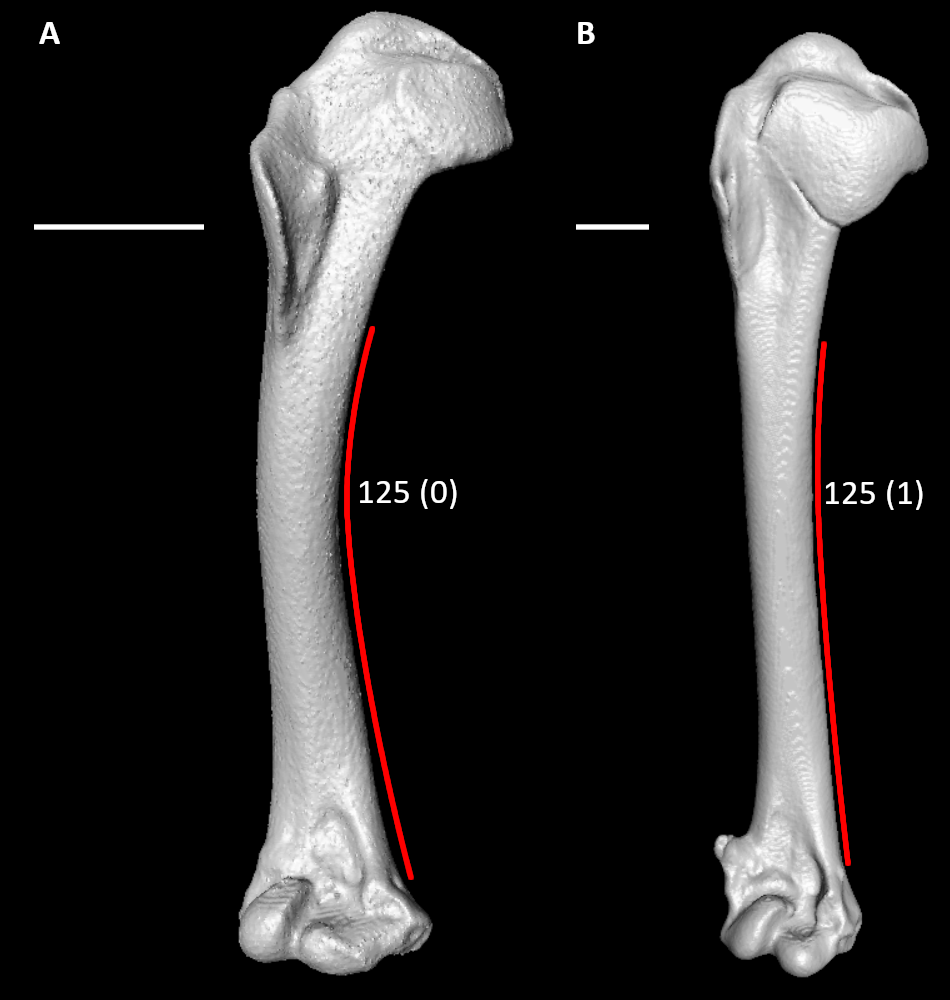

**Figure S91.** Humeri of *Aegotheles cristatus* (**A**, right element) and *Limosa lapponica* (**B**, left element, mirrored) in cranial view, illustrating alternative states for character 125 (complete character description in text). Red lines indicate approximate curvature of the humeral shaft. Scale bars = 5 mm.

126. Humerus, shaft width (Fig. S92; Worthy and Lee, 2008: 61; Worthy et al., 2017: 135):

0 – approximately constant throughout length, essentially parallel sides in caudal or cranial view

1 – narrows distally (at least 10% reduction on mid-length width), narrowest point in distal third

2 – narrowest near midshaft point

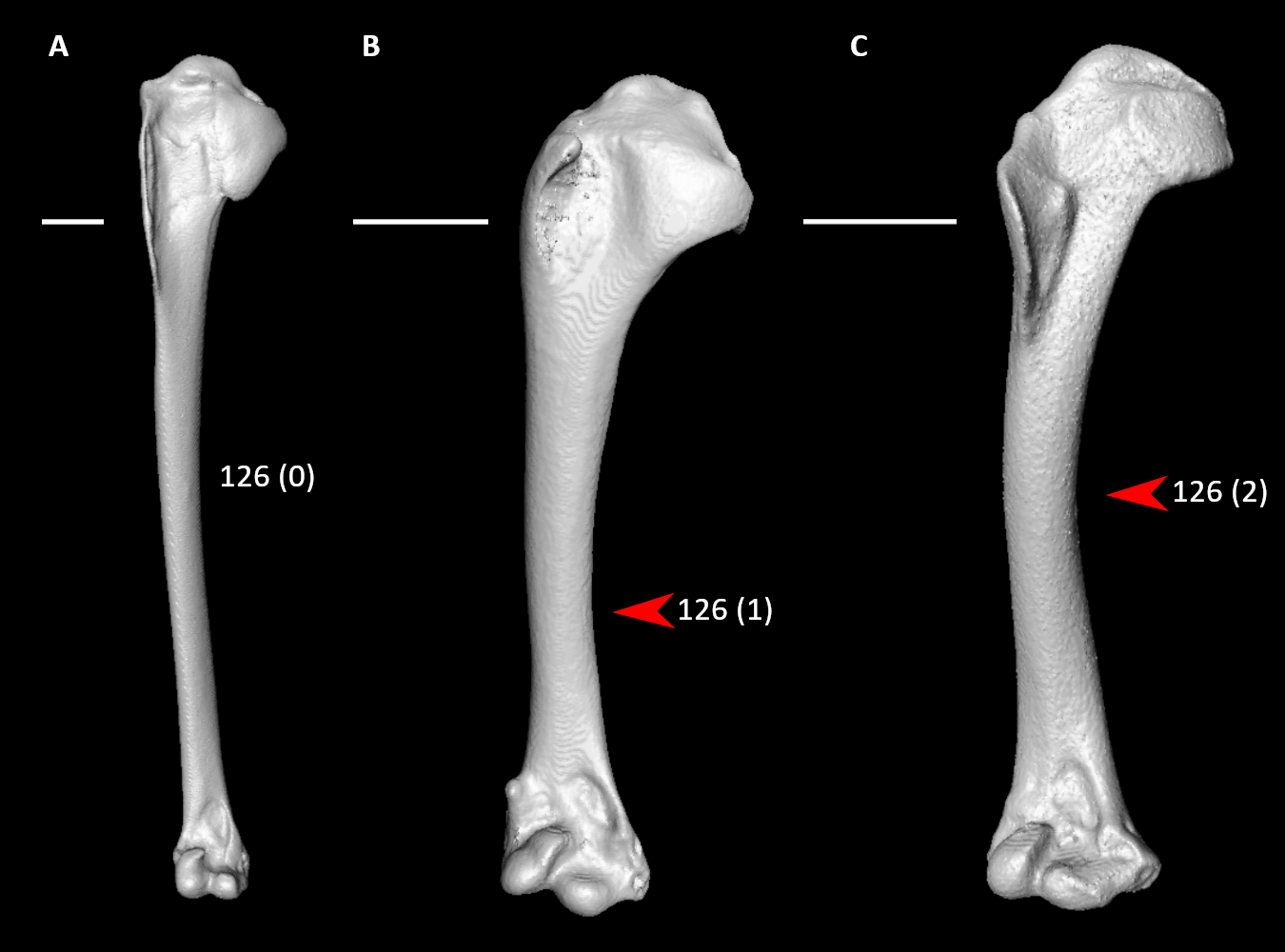

**Figure S92.** Humeri of *Podilymbus podiceps* (**A**, right element), *Turnix varius* (**B**, left element, mirrored), and *Aegotheles cristatus* (**C**, right element) in cranial view, illustrating alternative states for character 126 (complete character description in text). Arrows indicate narrowest point of the humeral shaft. Scale bars = 5 mm.

127. Humerus, distal end, caudal view, olecranon fossa (Fig. S80; Livezey and Zusi, 2006: 1482; Musser and Cracraft, 2019: 236; Musser and Clarke, 2020: 469):

0 – shallow

1 – deep, creating flattened region of distal humerus

128. Humerus, distal end, caudal view, tendinal sulcus of m. scapulotricipitalis (Fig. S93; Mayr and Clarke, 2003: 81):

0 – absent

1 – present

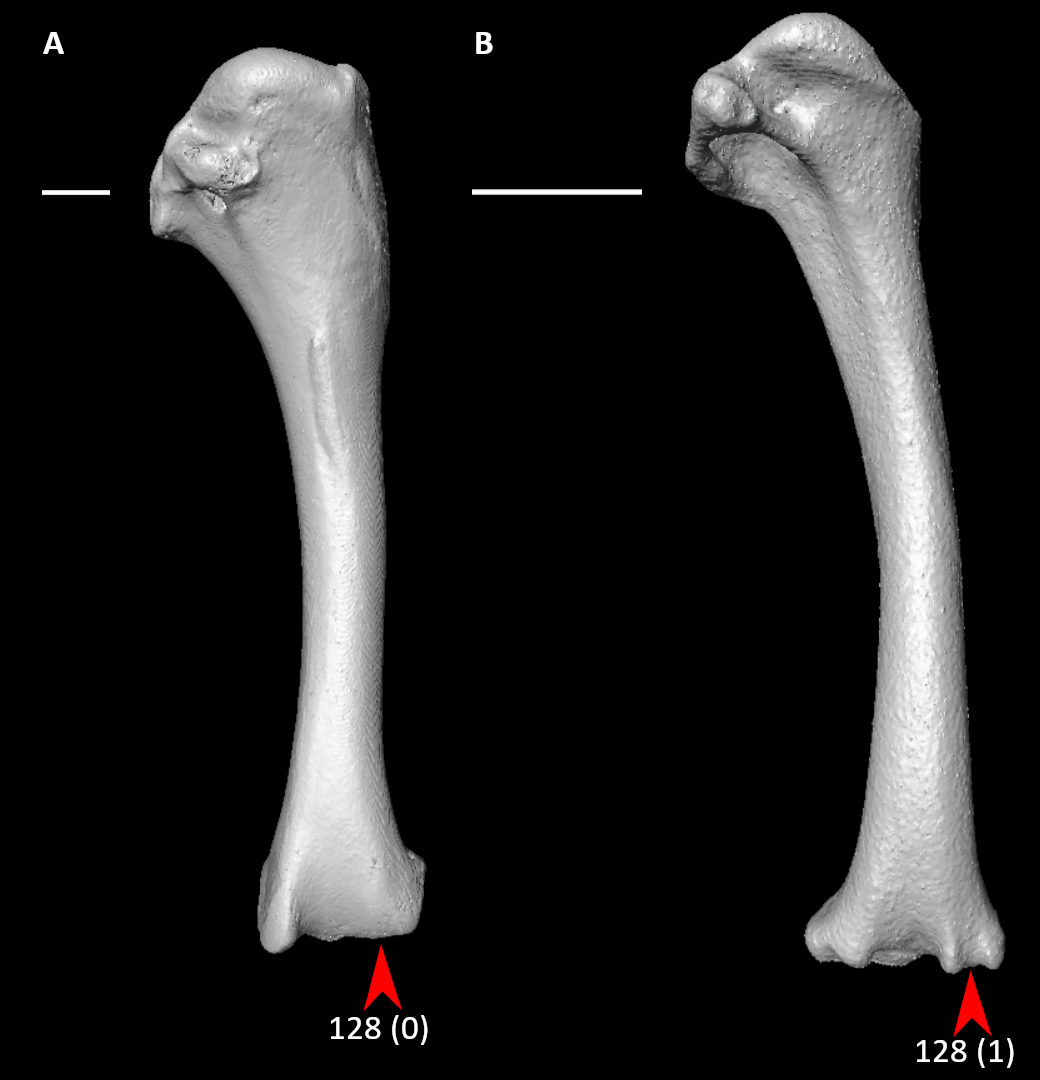

**Figure S93.** Humeri of *Eudromia elegans* (**A**, right element) and *Aegotheles cristatus* (**B**, right element) in caudal view, illustrating alternative states for character 128 (complete character description in text). Arrows indicate the tendinal sulcus of m. scapulotricipitalis or approximate homologous site. Scale bars = 5 mm.

129. Humerus, distal end, caudal view, tendinal sulcus of m. scapulotricipitalis (Fig. S82; Livezey and Zusi, 2006: 1488; Musser and Cracraft, 2019: 237; Musser and Clarke, 2020: 472):

0 – shallow, inconspicuous

1 – deep

130. Humerus, distal end, caudal view, proximal site of ventral epicondyle (entepicondyle) relative to ventral condyle (Fig. S94; Livezey and Zusi, 2006: 1475; Musser and Cracraft, 2019: 238):

0 – former proximal to latter

1 – former approximately equal or distal to latter

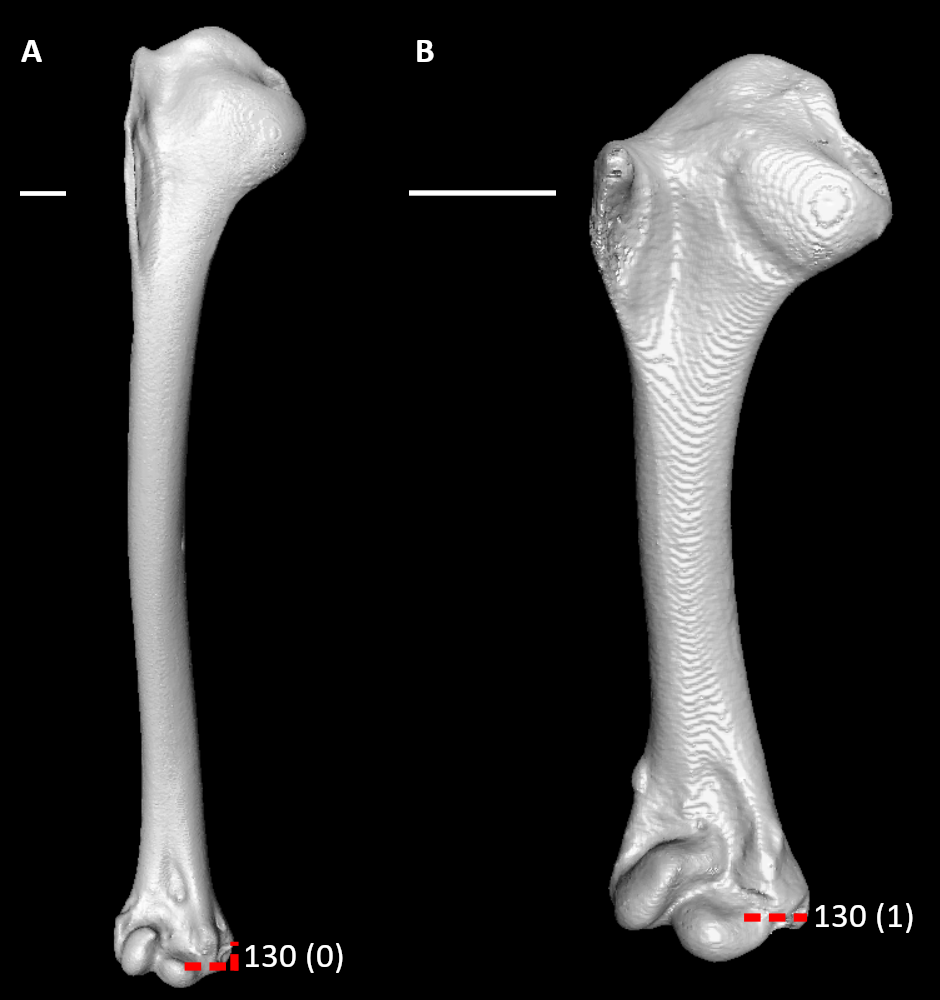

**Figure S94.** Humeri of *Dendrocygna bicolor* (**A**, right element) and *Pterocles quadricinctus* (**B**, left element, mirrored) in cranial view, illustrating alternative states for character 130 (complete character description in text). Dotted lines indicate proximal extent of the ventral epicondyle relative to that of the ventral condyle. Scale bars = 5 mm.

131. Humerus, distal end, caudal view, flexor process (Fig. S90; Ksepka et al., 2019: 64; Musser and Clarke, 2020: 471):

0 – weakly developed

1 – strongly projected, protruding distally beyond rest of bone

132. Humerus, distal end, cranial view, brachial fossa (Fig. S95):

0 – absent

1 – present

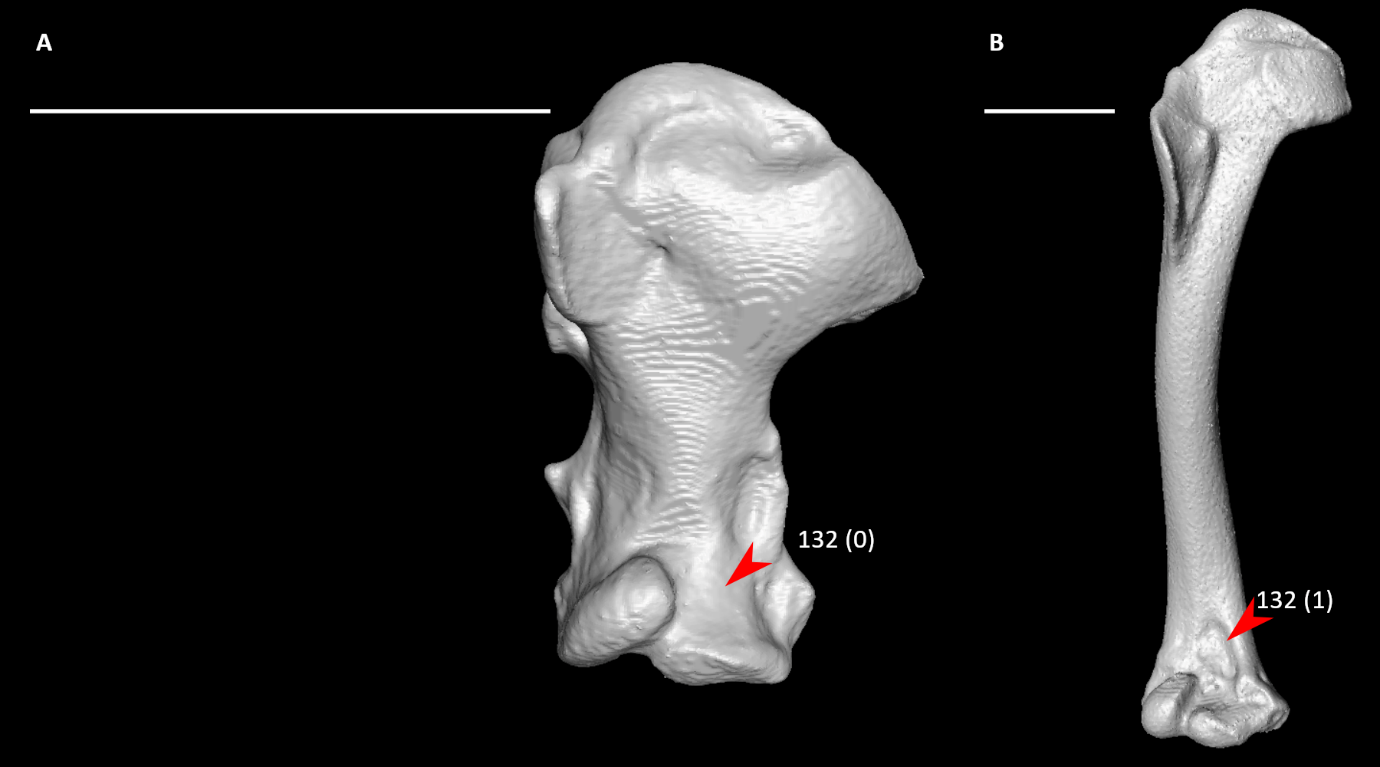

**Figure S95.** Humeri of *Florisuga mellivora* (**A**, left element, mirrored) and *Aegotheles cristatus* (**B**, right element) in cranial view, illustrating alternative states for character 132 (complete character description in text). Arrows indicate the brachial fossa or approximate homologous site. Scale bars = 5 mm.

133. Humerus, distal end, cranial view, dorsoventral position of brachial fossa relative to medial axis of humerus (Fig. S96; Livezey and Zusi, 2006: 1460; Musser and Cracraft, 2019: 239; Musser and Clarke, 2020: 473):

0 – ventral

1 – medial

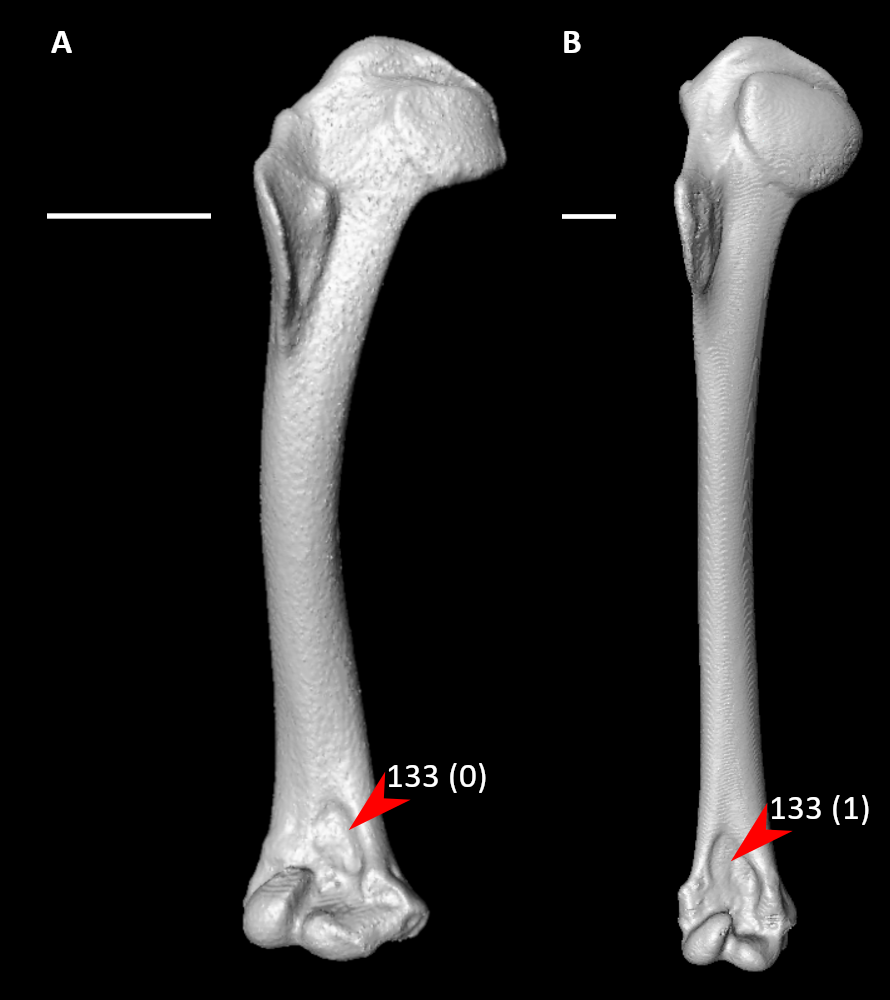

**Figure S96.** Humeri of *Aegotheles cristatus* (**A**, right element) and *Phaethon lepturus* (**B**, right element) in cranial view, illustrating alternative states for character 133 (complete character description in text). Arrows indicate the brachial fossa. Scale bars = 5 mm.

134. Humerus, distal end, cranial view, depth of brachial fossa (Fig. S97; Mayr and Clarke, 2003: 80; Livezey and Zusi, 2006: 1456; Musser and Cracraft, 2019: 240; Musser and Clarke, 2020: 474):

0 – shallow

1 – deep, distal portion of humerus tends to be especially depressed

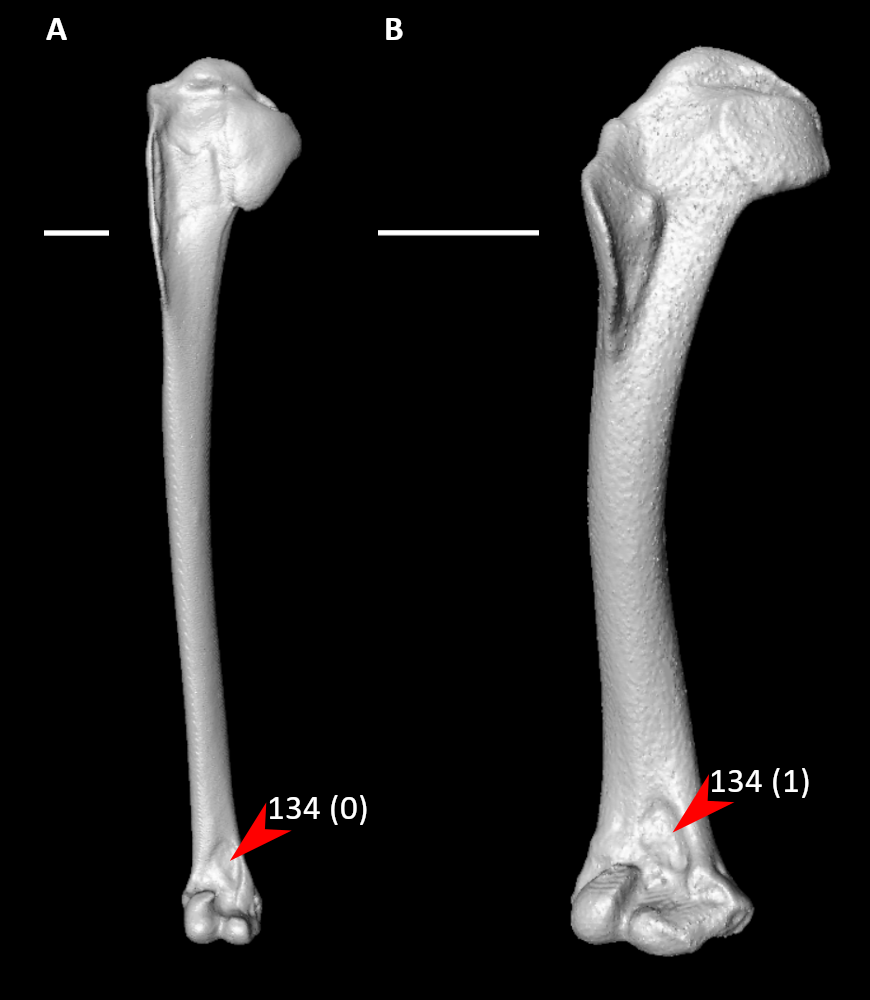

**Figure S97.** Humeri of *Podilymbus podiceps* (**A**, right element) and *Aegotheles cristatus* (**B**, right element) in cranial view, illustrating alternative states for character 134 (complete character description in text). Arrows indicate the brachial fossa. Scale bars = 5 mm.

135. Humerus, distal end, cranial view, dorsal supracondylar process (Fig. S98; Ksepka et al., 2013: 63):

0 – proximally placed

1 – distally placed

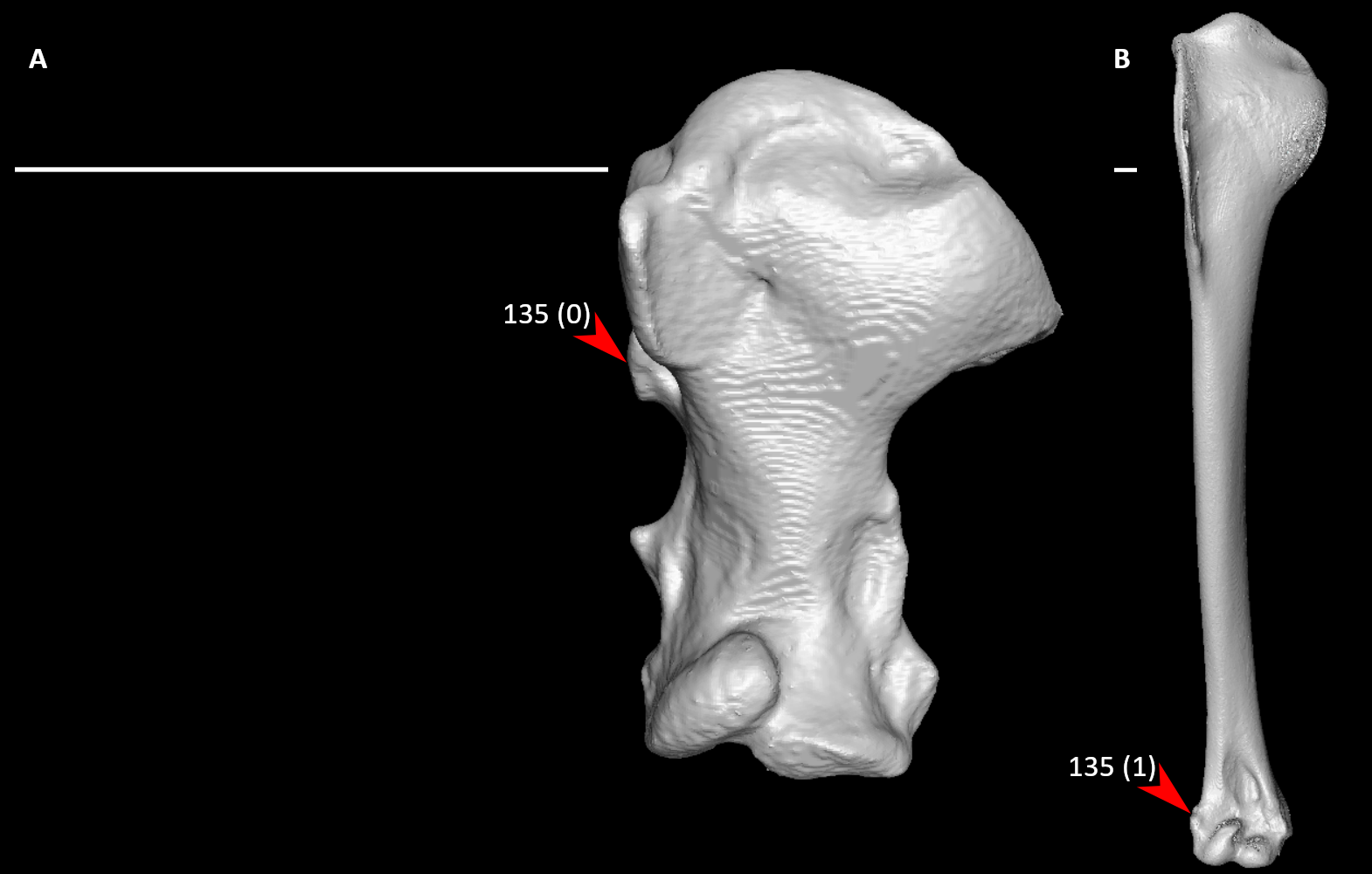

**Figure S98.** Humeri of *Florisuga mellivora* (**A**, left element, mirrored) and *Phoenicopterus ruber* (**B**, right element) in cranial view, illustrating alternative states for character 135 (complete character description in text). Arrows indicate the dorsal supracondylar process. Scale bars = 5 mm.

136. Humerus, distal end, cranial view, dorsal supracondylar process (Fig. S99; Livezey and Zusi, 2006: 1467; Ksepka et al., 2013: 62; Musser and Cracraft, 2019: 241; Musser and Clarke, 2020: 470):

0 – extremely small

1 – moderately large tubercle

2 – prominent, subtriangular process oriented dorsoproximally

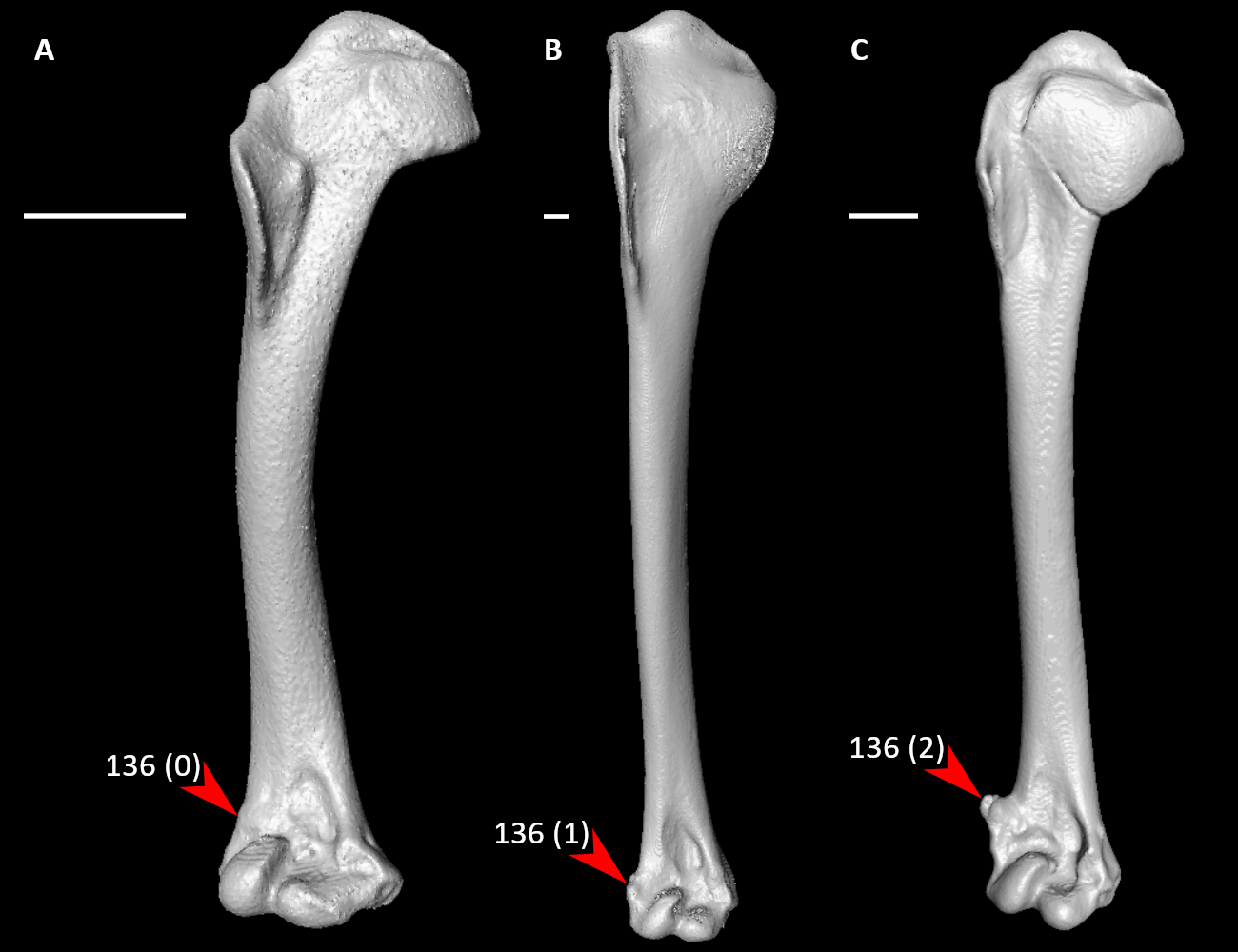

**Figure S99.** Humeri of *Aegotheles cristatus* (**A**, right element), *Phoenicopterus ruber* (**B**, right element), and *Limosa lapponica* (**C**, left element, mirrored) in cranial view, illustrating alternative states for character 136 (complete character description in text). Arrows indicate the dorsal supracondylar process. Scale bars = 5 mm.

137. Humerus, distal end, cranial view, dorsal supracondylar process (Fig. S100; Ksepka et al., 2019: 62):

0 – single

1 – bifurcated

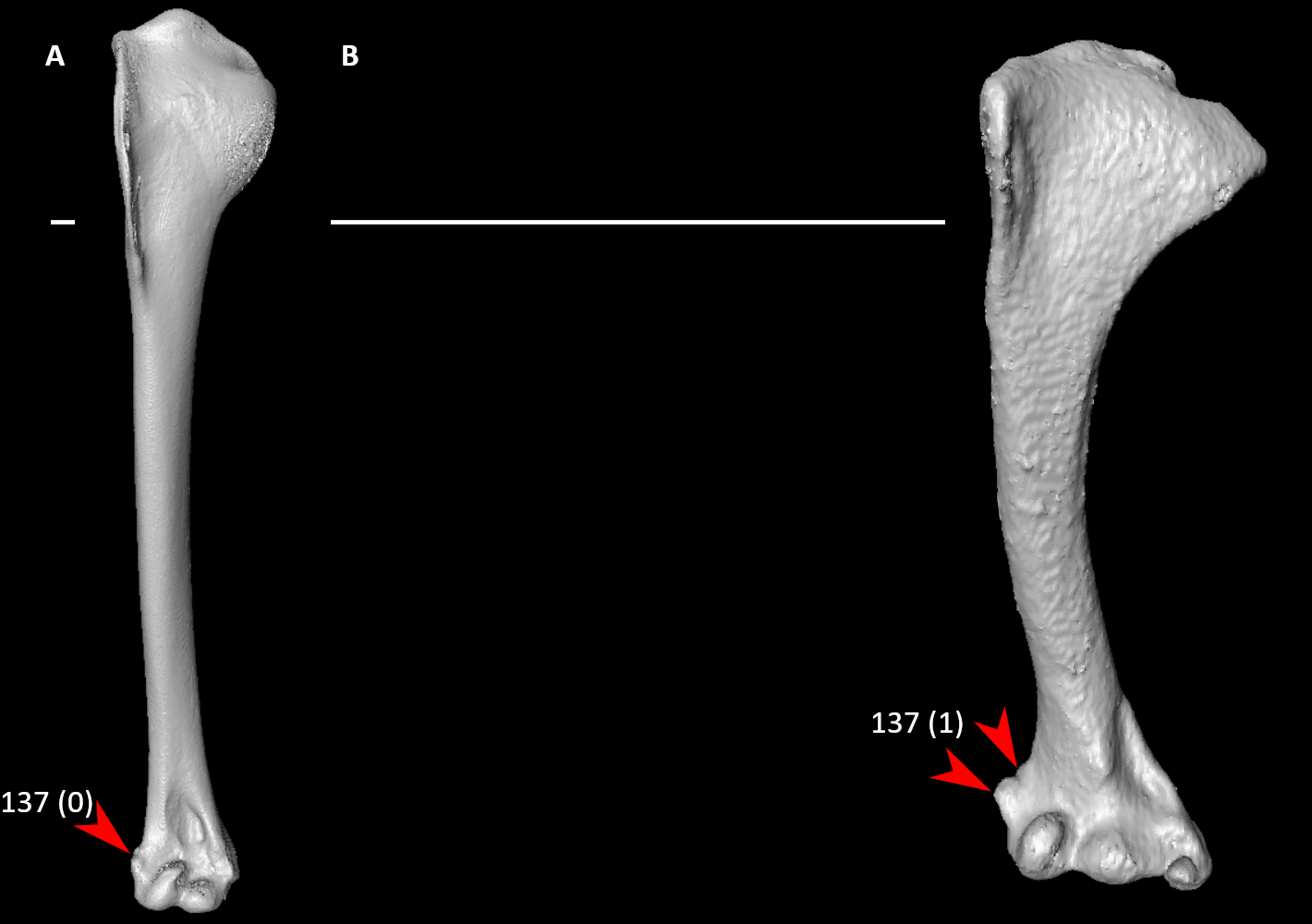

**Figure S100.** Humeri of *Phoenicopterus ruber* (**A**, right element) and *Acanthisitta chloris* (**B**, right element) in cranial view, illustrating alternative states for character 137 (complete character description in text). Arrows indicate the tip(s) of the dorsal supracondylar process. Scale bars = 5 mm.

138. Humerus, distal end, cranial view, dorsal epicondyle (Fig. S101; Livezey and Zusi, 2006: 1461; Musser and Cracraft, 2019: 242):

0 – absent or virtually coplanar with dorsal surface of humerus

1 – prominent

**Figure S101.** Humeri of *Podilymbus podiceps* (**A**, right element) and *Alectura lathami* (**B**, left element, mirrored) in cranial view, illustrating alternative states for character 138 (complete character description in text). Arrows indicate the dorsal epicondyle or approximate homologous site. Scale bars = 5 mm.

139. Humerus, distal end, cranial view, proximal margin of dorsal condyle (Fig. S86; Livezey and Zusi, 2006: 1451; Musser and Clarke, 2020: 476):

0 – rounded

1 – pointed

*140. Humerus, distal end, cranial view, proximal extent of dorsal condyle relative to distal margin of brachial fossa (Fig. S102; Livezey and Zusi, 2005: 1447; Musser and Cracraft, 2019: 244; Musser and Clarke, 2020: 475):

0 – condyle distal to distal terminus of fossa and typically separated from fossa by smooth area of bone

1 – condyle proximal margin typically extending at least proximal to distal margin of fossa

**Figure S102.** Humeri of *Phoenicopterus ruber* (**A**, right element) and *Aegotheles cristatus* (**B**, right element) in cranial view, illustrating alternative states for character 140 (complete character description in text). Dotted lines indicate the proximal extent of dorsal condyle relative to distal margin of brachial fossa. Scale bars = 5 mm.

141. Humerus, distal end, cranial view, ventral supracondylar tubercle (Fig. S103; Ksepka et al., 2019: 63):

0 – moderately developed

1 – expanded, forming large triangular platform approximately equal in width to ventral condyle and in dorsal extent to dorsal condyle

**Figure S103.** Humeri of *Dendrocygna bicolor* (**A**, right element) and *Trogon melanurus* (**B**, left element, mirrored) in cranial view, illustrating alternative states for character 141 (complete character description in text). Arrows indicate the ventral supracondylar tubercle. Scale bars = 5 mm.

142. Humerus, distal end, ventral view, origin of m. flexor carpi ulnaris on flexor process (Fig. S104; Ericson, 1997: 60; Worthy et al., 2017: 143):

0 – scar indistinct

1 – one scar

2 – two scars

**Figure S104.** Humeri of *Jynx torquilla* (**A**, right element), *Rollulus rouloul* (**B**, right element), and *Aegotheles cristatus* (**C**, right element) in cranioventral view, illustrating alternative states for character 142 (complete character description in text). Arrows indicate scar(s) for the origin of m. flexor carpi ulnaris on flexor process. Scale bars = 5 mm.

143. Humerus, distal end, ventral view, origin of m. flexor carpi ulnaris on flexor process (if present as two scars) (Fig. S105):

0 – both approximately equal depth

1 – caudal scar shallow, cranial scar deep

**Figure S105.** Humeri of *Phoenicopterus ruber* (**A**, right element) and *Podargus strigoides* (**B**, left element, mirrored) in cranioventral view, illustrating alternative states for character 143 (complete character description in text). Arrows indicate scars for the origin of m. flexor carpi ulnaris on flexor process. Scale bars = 5 mm.

144. Radius, shaft, craniocaudal curvature (Fig. S106):

0 – straight

1 – curved

**Figure S106.** Radii of *Rollulus rouloul* (**A**, right element) and *Aegotheles cristatus* (**B**, right element) in dorsal view, illustrating alternative states for character 144 (complete character description in text). Red lines indicate approximate craniocaudal curvature of the radial shaft. Scale bars = 5 mm.

145. Radius, osseous loop from shaft (Fig. S107; Ksepka et al., 2019: 67):

0 – absent

1 – present

**Figure S107.** Radii of *Aegotheles cristatus* (**A**, right element) and *Ninox novaeseelandiae* (**B**, right element) in caudal view, illustrating alternative states for characters 145 and 146 (complete character descriptions in text). Arrows for character 145 indicate the radial shaft and its osseous loop (when present), whereas arrows for character 146 indicate the radial bicipital tubercle. Scale bars = 5 mm.

146. Radius, proximal end, radial bicipital tubercle (Fig. S107; Ksepka et al., 2019: 66):

0 – small

1 – crest-like projection

147. Radius, distal end, dorsoventral curvature (Fig. S108):

0 – absent or slight

1 – pronounced

**Figure S108.** Radii of *Phoenicopterus ruber* (**A**, right element) and *Jynx torquilla* (**B**, right element) in cranial view, illustrating alternative states for character 147 (complete character description in text). Red lines indicate approximate dorsoventral curvature of the distal radius. Scale bars = 5 mm.

148. Radius, distal end, radial tendinal sulcus (Fig. S109):

0 – weak

1 – well defined

**Figure S109.** Radii of *Rollulus rouloul* (**A**, right element) and *Aegotheles cristatus* (**B**, right element) in dorsal view, illustrating alternative states for character 148 (complete character description in text). Arrows indicate the radial tendinal sulcus. Scale bars = 5 mm.

149. Radius, distal end, ventral aponeurosis tubercle shape (Fig. S110):

0 – round

1 – pointed

**Figure S110.** Radii of *Jynx torquilla* (**A**, right element) and *Aegotheles cristatus* (**B**, right element) in dorsal view, illustrating alternative states for character 149 (complete character description in text). Arrows indicate the ventral aponeurosis tubercle. Scale bars = 5 mm.

150. Ulna, proximal end, dorsal cotylar process (Fig. S111; Livezey and Zusi, 2006: 1492; Musser and Cracraft, 2019: 245; Musser and Clarke, 2020: 478):

0 – apex variably elevated dorsally compared to ulnar main body

1 – apex approximately coplanar with dorsal surface of ulnar main body

**Figure S111.** Ulnae of *Aegotheles cristatus* (**A**, right element) and *Dendrocygna bicolor* (**B**, right element) in cranial view, illustrating alternative states for character 150 (complete character description in text). Arrows indicate the dorsal cotylar process where it joins with the ulnar main body. Scale bars = 5 mm.

151. Ulna, proximal end, dorsal cotylar process, articular facet relative to ventral cotyle (Fig. S112; Livezey and Zusi, 2006: 1496; Musser and Cracraft, 2019: 246; Musser and Clarke, 2020: 479):

0 – facet of dorsal cotyle less expansive than that of ventral cotyle

1 – facets of both cotyles subequal

2 – facet of ventral cotyle less expansive than that of dorsal cotyle

**Figure S112.** Ulnae of *Aegotheles cristatus* (**A**, right element), *Rollulus rouloul* (**B**, right element), and *Podilymbus podiceps* (**C**, right element) in proximal view, illustrating alternative states for character 151 (complete character description in text). Red lines span the dorsoventral widths of the dorsal and ventral cotyles. Scale bars = 5 mm.

152. Ulna, proximal end, dorsal cotylar process, intercotylar crest (Fig. S113; Livezey and Zusi, 2006: 1497; Musser and Cracraft, 2019: 247; Musser and Clarke, 2020: 480):

0 – poorly developed

1 – prominent, dorsal and ventral cotyles clearly demarcated

**Figure S113.** Ulnae of *Phoenicopterus ruber* (**A**, right element) and *Aegotheles cristatus* (**B**, right element) in proximal view, illustrating alternative states for character 152 (complete character description in text). Arrows indicate the intercotylar crest. Scale bars = 5 mm.

153. Ulna, proximal end, deep pit distal to rim of ventral cotyle (Fig. S114; Ksepka et al., 2019: 71):

0 – absent

1 – present

**Figure S114.** Ulnae of *Aegotheles cristatus* (**A**, right element) and *Menura novaehollandiae* (**B**, right element) in ventral view, illustrating alternative states for character 153 (complete character description in text). Arrows indicate the deep pit distal to rim of ventral cotyle, or approximate homologous site to the pit. Scale bars = 5 mm.

154. Ulna, proximal end, humeroulnar trochlea (Fig. S115; Musser and Clarke, 2020: 483)

0 – absent or shallow

1 – deep, forming prominent depression or groove

**Figure S115.** Ulnae of *Aegotheles cristatus* (**A**, right element) and *Podilymbus podiceps* (**B**, right element) in ventral view, illustrating alternative states for characters 154, 155, 158, and 162 (complete character descriptions in text). Arrows for character 154 indicate the humeroulnar trochlea, arrows for character 155 indicate the impression of m. brachialis, arrows for character 158 indicate the ventral collateral ligamental tubercle, and arrows for character 162 indicate the radial depression. Scale bars = 5 mm.

155. Ulna, proximal end, cranial margin, impression of m. brachialis (Fig. S115; Livezey and Zusi, 2006: 1502; Musser and Cracraft, 2019: 248; Musser and Clarke, 2020: 481):

0 – flat, proximocaudal margin only slightly elevated

1 – modestly concave, deep with proximocaudal margin or brachial crest elevated

156. Ulna, proximal end, dorsal view, radial incisure (Fig. S116; Livezey and Zusi, 2006: 1505; Musser and Cracraft, 2019: 249; Musser and Clarke, 2020: 482):

0 – absent or indistinct

1 – prominent

**Figure S116.** Ulnae of *Aegotheles cristatus* (**A**, right element) and *Podilymbus podiceps* (**B**, right element) in cranial view, illustrating alternative states for character 156 (complete character description in text). Arrows indicate the radial incisure. Scale bars = 5 mm.

157. Ulna, proximal end, scapulotricipital impression (Fig. S117; Musser and Clarke, 2020: 484):

0 – shallow

1 – deep

**Figure S117.** Ulnae of *Aegotheles cristatus* (**A**, right element) and *Podilymbus podiceps* (**B**, right element) in dorsal view, illustrating alternative states for characters 157 and 161 (complete character descriptions in text). Arrows for character 157 indicate the scapulotricipital impression, whereas arrows for character 161 indicate the ulnar shaft and one of its quill knobs (when present). Scale bars = 5 mm.

158. Ulna, proximal end, ventral collateral ligamental tubercle (Fig. S115; Ksepka et al., 2019: 72):

0 – weakly developed

1 – well developed and strongly projecting

159. Ulna, proximal end, olecranon (Fig. S118; Ksepka et al., 2019: 68; Musser and Clarke, 2020: 485):

0 – blunt and rounded

1 – narrow and sharply projected

**Figure S118.** Ulnae of *Aegotheles cristatus* (**A**, right element) and *Jynx torquilla* (**B**, right element) in dorsal view, illustrating alternative states for character 159 (complete character description in text). Arrows indicate the olecranon process. Scale bars = 5 mm.

160. Ulna, shaft, craniocaudal curvature (Fig. S119):

0 – absent or slight

1 – prominently curved

**Figure S119.** Ulnae of *Aegotheles cristatus* (**A**, right element) and *Columba livia* (**B**, left element, mirrored) in ventral view, illustrating alternative states for characters 160 and 163 (complete character descriptions in text). Red lines for character 160 indicate the approximate craniocaudal curvature of the ulnar shaft, whereas red lines for character 163 span the craniocaudal width of the carpal tubercle. Scale bars = 5 mm.

161. Ulna, shaft, quill knobs (Fig. S117; Ksepka et al., 2019: 70):

0 – absent or weakly developed

1 – strongly developed, prominently elevated

162. Ulna, distal end, radial depression (Fig. S115; Mayr and Clarke, 2003: 84; Worthy et al., 2017: 153; Musser and Clarke, 2020: 488):

0 – weak

1 – prominent

163. Ulna, distal end, carpal tubercle (Fig. S119; Worthy et al., 2017: 154):

0 – projects less than half the craniocaudal width of the dorsal and ventral condyles

1 – projects more than half the craniocaudal width of the dorsal and ventral condyles

164. Scapholunare, sulcus for tendon of m. extensor longus alulae (Fig. S120; Ksepka et al., 2019: 74):

0 – absent or weakly developed

1 – well-defined

Scapholunare characters scored based on *Colius* for *Urocolius*.

**Figure S120.** Scapholunares of *Aegotheles cristatus* (**A**, right element) and *Jynx torquilla* (**B**, left element, mirrored) in cranial view, illustrating alternative states for character 164 (complete character description in text). Arrows indicate the sulcus for tendon of m. extensor longus alulae. Scale bars = 5 mm.

165. Scapholunare, pneumatic foramina on distal surface (Fig. S121; Livezey and Zusi, 2006: 1563; Smith, 2010: 268):

0 – absent

1 – present

**Figure S121.** Scapholunares of *Aegotheles cristatus* (**A**, right element) and *Coragyps atratus* (**B**, right element) in distal view, illustrating alternative states for character 165 (complete character description in text). Arrows indicate the distal surface and its pneumatic foramen (when present). Scale bars = 5 mm.

*166. Pisiform, cranial extent of dorsal and ventral rami (Fig. S122; Mayr and Clarke, 2003: 87; Livezey and Zusi, 2006: 1570; Mayr, 2010: 41; Smith, 2010: 266; Nesbitt et al., 2011: 70):

0 – ventral ramus (crus longum) extends cranial to dorsal ramus (crus breve)

1 – ventral ramus subequal in cranial extent to dorsal ramus

2 – ventral ramus extends cranial to dorsal ramus

Scored state 1 in *Tapera*, but is reported to be state 2 in other cuckoos (Mayr and Ericson, 2004). Pisiform characters scored based on *Colius* for *Urocolius*.

**Figure S122.** Pisiforms of *Aegotheles cristatus* (**A**, right element), *Phoenicopterus ruber* (**B**, left element, mirrored), and *Corythaeola cristata* (**C**, right element) in distal view, illustrating alternative states for character 166 (complete character description in text). Dotted lines indicate the cranial extent of the ventral ramus relative to that of the dorsal ramus. Scale bars = 5 mm.

167. Pisiform, tubercle at insertion point of humerocarpal ligament (Fig. S123; Mayr and Clarke, 2003: 88; Livezey and Zusi, 2006: 1573; Smith, 2010: 264):

0 – absent

1 – present

Scored 1 in *Sterna* but is reduced.

**Figure S123.** Pisiforms of *Scopus umbretta* (**A**, right element) and *Aegotheles cristatus* (**B**, right element) in proximal view, illustrating alternative states for characters 167 and 168 (complete character descriptions in text). Arrows for character 167 indicate the tubercle at insertion point of humerocarpal ligament, or approximate homologous site to the tubercle; whereas arrows for character 168 indicate the ligamental groove on ventral ramus. Scale bars = 5 mm.

168. Pisiform, ligamental groove on ventral ramus (Fig. S123):

0 – absent or slight

1 – prominent

169. Carpometacarpus, dentiform process (metacarpal protuberance) (Fig. S124; Ksepka et al., 2019: 78):

0 – absent

1 – present

**Figure S124.** Carpometacarpi of *Podilymbus podiceps* (**A**, right element) and *Bucco capensis* (**B**, left element, mirrored) in dorsal view, illustrating alternative states for characters 169 and 170 (complete character descriptions in text). Arrows for character 169 indicate the cranial margin of the carpometacarpus and the dentiform process (when present), whereas arrows for character 170 indicate the dorsal tendinal sulcus. Scale bars = 5 mm.

170. Carpometacarpus, dorsal view, tendinal sulcus (Fig. S124; Ksepka et al., 2019: 82):

0 – barely perceptible

1 – distinct

171. Carpometacarpus, dorsal view, tendinal sulcus length (if distinct) (Fig. S125):

0 – extends less than half of major metacarpal length

1 – extends half of major metacarpal length or more

**Figure S125.** Carpometacarpi of *Monias benschi* (**A**, left element, mirrored) and *Aegotheles cristatus* (**B**, right element) in dorsal view, illustrating alternative states for character 171 (complete character description in text). Red lines span the proximodistal extent of the dorsal tendinal sulcus. Scale bars = 5 mm.

172. Carpometacarpus, dorsal view, tendinal sulcus length (if extending half of major metacarpal length or more) (Fig. S126):

0 – continues along dorsal surface of major digit

1 – wraps around cranial surface of major digit

**Figure S126.** Carpometacarpi of *Aegotheles cristatus* (**A**, right element) and *Jynx torquilla* (**B**, right element) in cranial view, illustrating alternative states for character 172 (complete character description in text). Arrows indicate the dorsal tendinal sulcus. Scale bars = 5 mm.

173. Carpometacarpus, distinct ridge from the caudal end of minor metacarpal to pisiform process (Fig. S127; Ksepka et al., 2019: 84):

0 – absent

1 – present

**Figure S127.** Carpometacarpi of *Aegotheles cristatus* (**A**, right element) and *Psophia crepitans* (**B**, right element) in ventral view, illustrating alternative states for characters 173, 186, and 194 (complete character descriptions in text). Arrows for character 173 indicate the region between the caudal end of minor metacarpal and the pisiform process, and the ridge between the two (when present); whereas arrows for character 186 indicate the small ventral tubercle on the minor metacarpal. Dotted lines indicate the distal extent of the minor metacarpal relative to that of the major metacarpal. Scale bars = 5 mm.

174. Carpometacarpus, bowing of minor metacarpal (Fig. S128; Mayr and Clarke, 2003: 85; Musser et al., 2019: 73; Musser and Clarke, 2020: 492):

0 – weak, delimiting a narrow intermetacarpal space with nearly parallel cranial and caudal margins

1 – strong, delimiting a large, ovoid intermetacarpal space

**Figure S128.** Carpometacarpi of *Podilymbus podiceps* (**A**, right element) and *Aegotheles cristatus* (**B**, right element) in ventral view, illustrating alternative states for characters 174, 188, and 198 (complete character descriptions in text). Arrows for character 174 indicate the intermetacarpal space, whereas arrows for character 188 indicate the cranial carpal fossa or approximately homologous site. Red lines span the proximodistal length of synostosis between the major and minor metacarpals. Scale bars = 5 mm.

175. Carpometacarpus, proximal end, carpal trochlea, trochlear sulcus (Fig. S129; Livezey, 1998: 236; Musser and Cracraft, 2019: 251; Musser and Clarke, 2020: 490):

0 - shallow, rounded in cranial or caudal view, or is somewhat deep laterally but not cranially

1 – deep, subangular in cranial or caudal view

**Figure S129.** Carpometacarpi of *Chauna chavaria* (**A**, left element, mirrored) and *Gavia stellata* (**B**, left element, mirrored) in caudal view, illustrating alternative states for character 175 (complete character description in text). Red lines indicate the approximate shape of the proximal margin of the trochlear sulcus. Scale bars = 5 mm.

176. Carpometacarpus, proximal end, dorsal view, proximal termination of dorsal rim of carpal trochlea (Fig. S130; Livezey, 1998: 236; Musser and Cracraft, 2019: 252; Musser and Clarke, 2020: 493):

0 – weakly angular

1 – rounded

2 – strongly angular, almost pointed, elongated proximally

**Figure S130.** Carpometacarpi of *Phoenicopterus ruber* (**A**, left element, mirrored), *Podilymbus podiceps* (**B**, right element), and *Aegotheles cristatus* (**C**, right element) in dorsal view, illustrating alternative states for character 176 (complete character description in text). Red lines indicate the approximate shape of the proximal dorsal margin of the trochlear sulcus. Scale bars = 5 mm.

*177. Carpometacarpus, proximal end, caudal view, distal extent of dorsal rim of carpal trochlea (Fig. S131; Ericson, 1997: 64; Worthy et al., 2017: 156):

0 – ending considerably short of ventral rim

1 – falling only slightly short of ventral rim

2 – equals or exceeds ventral rim

**Figure S131.** Carpometacarpi of *Rollulus rouloul* (**A**, right element), *Aegotheles cristatus* (**B**, right element), and *Chauna chavaria* (**C**, left element, mirrored) in caudal view, illustrating alternative states for character 177 (complete character description in text). Dotted lines indicate the distal extent of the dorsal rim of the trochlear sulcus relative to that of the ventral rim. Scale bars = 5 mm.

178. Carpometacarpus, proximal end, caudal view, position of ventral rim of carpal trochlea relative to synostosed area of minor and major metacarpals (Fig. S132; Worthy et al., 2017: 159):

0 – minor metacarpal extends dorsad of ventral rim

1 – minor metacarpal entirely ventral to ventral rim

**Figure S132.** Carpometacarpi of *Balearica pavonina* (**A**, right element) and *Rollulus rouloul* (**B**, right element) in caudal view, illustrating alternative states for characters 178 and 180 (complete character descriptions in text). Dotted lines indicate the dorsal extent of the ventral margin of the trochlear sulcus, whereas solid lines indicate the dorsal and ventral margins of the minor metacarpal. Scale bars = 5 mm.

179. Carpometacarpus, proximal end, caudal view, minor metacarpal dorsoventral thickness at proximal third of length (Fig. S133):

0 – less than half as thick as that of major metacarpal

1 – more than half as thick as that of major metacarpal

**Figure S133.** Carpometacarpi of *Florisuga mellivora* (**A**, left element, mirrored) and *Aegotheles cristatus* (**B**, right element) in caudal view, illustrating alternative states for character 179 (complete character description in text). Red lines span the dorsoventral thickness of the minor metacarpal at proximal third of its length. Scale bars = 5 mm.

180. Carpometacarpus, proximal end, caudal view, minor metacarpal dorsoventral thickness (Fig. S132):

0 – maintains approximate thickness throughout

1 – prominently narrows distally

181. Carpometacarpus, proximal end, caudal view, minor metacarpal, caudal groove (Fig. S134; Livezey, 1986: 44; Livezey, 1996: 59; Livezey, 1998: 252; Worthy and Lee, 2008: 80; Worthy et al., 2017: 164):

0 – absent or weak

1 – prominent, extending to synostosis with major metacarpal

2 – prominent, distal to synostosis with major metacarpal

**Figure S134.** Carpometacarpi of *Aegotheles cristatus* (**A**, right element), *Balearica pavonina* (**B**, right element), and *Rollulus rouloul* (**C**, right element) in caudal view, illustrating alternative states for character 181 (complete character description in text). Arrows indicate the caudal groove or approximate homologous site. Scale bars = 5 mm.

*182. Carpometacarpus, proximal end, extensor process (Fig. S135; Clarke, 2004: 142; Ksepka et al., 2019: 85):

0 – absent

1 – present, just surpassing distal articular facet for phalanx 1 in cranial extent

2 – present, surpasses articular facet by approximately half the width of facet

3 – present, surpasses articular facet by approximately width of facet

4 – present, cranial extent beyond articular facet surpasses width of facet

**Figure S135.** Carpometacarpi of *Gavia stellata* (**A**, left element, mirrored), *Podilymbus podiceps* (**B**, right element), *Charadrius vociferus* (**C**, left element, mirrored), *Alca torda* (**D**, left element, mirrored), and *Bucco capensis* (**E**, left element, mirrored) in dorsal view, illustrating alternative states for character 182 (complete character description in text). Red lines span the craniocaudal heights of the extensor process and the articular facet for phalanx I-1. Arrow indicates approximate homologous site to the extensor process in (**A**). Scale bars = 5 mm.

183. Carpometacarpus, proximal end, intermetacarpal process (Fig. S136; Ksepka et al., 2019: 77):

0 – absent or inconspicuous

1 – well-developed, contacting minor metacarpal

**Figure S136.** Carpometacarpi of *Aegotheles cristatus* (**A**, right element) and *Rollulus rouloul* (**B**, right element) in dorsal view, illustrating alternative states for character 183 (complete character description in text). Arrows indicate the intermetacarpal process. Scale bars = 5 mm.

184. Carpometacarpus, proximal end, proximal extent of intermetacarpal space (Fig. S137; Clarke, 2004: 147):

0 – reaches distal end of alular metacarpal

1 – distal to distal end of alular metacarpal

**Figure S137.** Carpometacarpi of *Gavia stellata* (**A**, left element, mirrored) and *Aegotheles cristatus* (**B**, right element) in dorsal view, illustrating alternative states for character 184 (complete character description in text). Dotted lines indicate the distal extent of the alular metacarpal relative to the proximal extent of the intermetacarpal space. Scale bars = 5 mm.

185. Carpometacarpus, proximal end, dorsal view, caudal carpal fossa (cuniform fossa), pneumatic foramen (Fig. S138):

0 – absent

1 – present

**Figure S138.** Carpometacarpi of *Aegotheles cristatus* (**A**, right element) and *Chauna chavaria* (**B**, left element, mirrored) in caudal view, illustrating alternative states for character 185 (complete character description in text). Arrows indicate the caudal carpal fossa. Scale bars = 5 mm.

186. Carpometacarpus, proximal end, ventral view, small tubercle on minor metacarpal immediately distal to proximal synostosis of metacarpals (Fig. S127):

0 – absent or weak

1 – prominent

187. Carpometacarpus, proximal end, ventral view, small tubercle on minor metacarpal immediately distal to proximal synostosis of metacarpals (Fig. S139; Livezey, 1998: 244; Musser and Cracraft, 2019: 253; Musser and Clarke, 2020: 494):

0 – elongate

1 – distinct and rounded

**Figure S139.** Carpometacarpi of *Coragyps atratus* (**A**, right element) and *Psophia crepitans* (**B**, right element) in ventral view, illustrating alternative states for character 187 (complete character description in text). Arrows indicate the tubercle on the minor metacarpal. Scale bars = 5 mm.

188. Carpometacarpus, proximal end, ventral view, cranial carpal fossa (Fig. S128):

0 – absent

1 – present

189. Carpometacarpus, proximal end, ventral view, cranial carpal fossa, pneumatic foramen (Fig. S140; Worthy and Lee, 2008: 74; Worthy et al., 2017: 157):

0 – absent

1 – present

**Figure S140.** Carpometacarpi of *Podilymbus podiceps* (**A**, right element) and *Fregata aquila* (**B**, left element, mirrored) in ventral view, illustrating alternative states for characters 189 and 196 (complete character descriptions in text). Arrows for character 189 indicate the cranial carpal fossa, whereas arrows for character 196 indicate the ventral interosseus sulcus. Scale bars = 5 mm.

190. Carpometacarpus, proximal end, ventral view, infratrochlear fossa (Fig. S141):

0 – absent

1 – present

**Figure S141.** Carpometacarpi of *Rollulus rouloul* (**A**, right element) and *Aegotheles cristatus* (**B**, right element) in ventral view, illustrating alternative states for character 190 (complete character description in text). Arrows indicate the infratrochlear fossa or approximate homologous site. Scale bars = 5 mm.

191. Carpometacarpus, proximal end, ventral view, infratrochlear fossa, pneumatic foramen (Fig. S142):

0 – absent

1 – present

**Figure S142.** Carpometacarpi of *Aegotheles cristatus* (**A**, right element) and *Balearica pavonina* (**B**, right element) in ventral view, illustrating alternative states for character 191 (complete character description in text). Arrows indicate the infratrochlear fossa. Scale bars = 5 mm.

*192. Carpometacarpus, proximal end, ventral view, ridge linking ventral rim of carpal trochlea and pisiform process, relationship to ventral surface of extensor process (Fig. S143; Worthy and Lee, 2008: 77; Worthy et al., 2017: 161):

0 – rounded profile, little elevated

1 – sharp drop-off

2 – overhangs ventral surface of extensor process with resultant pit under ledge

**Figure S143.** Carpometacarpi of *Aegotheles cristatus* (**A**, right element), *Podilymbus podiceps* (**B**, right element), and *Rollulus rouloul* (**C**, right element) in proximal view, illustrating alternative states for character 192 (complete character description in text). Red lines indicate the cranioventral margin of the ridge linking the ventral rim of carpal trochlea and the pisiform process. Scale bars = 5 mm.

193. Carpometacarpus, proximal end, ventral view, pisiform process, cranial overhang (Fig. S144):

0 – absent or weak

1 – prominent, forming strongly concave cranial surface

**Figure S144.** Carpometacarpi of *Corythaeola cristata* (**A**, left element, mirrored) and *Caprimulgus macrurus* (**B**, right element) in proximal view, illustrating alternative states for character 193 (complete character description in text). Red lines indicate the cranial margin of the pisiform process. Scale bars = 5 mm.

194. Carpometacarpus, distal end, relative distal projection of minor and major metacarpals (Fig. S127; Ksepka et al., 2019: 79):

0 – subequal

1 – minor metacarpal projects substantially beyond major metacarpal

195. Carpometacarpus, distal end, dorsal view, distal synostosis of metacarpals, dorsal interosseal sulcus (Fig. S145; Livezey, 1998: 254):

0 – absent or shallow, nearly flat

1 – deep

**Figure S145.** Carpometacarpi of *Upupa epops* (**A**, left element, mirrored) and *Bucco capensis* (**B**, left element, mirrored) in dorsal view, illustrating alternative states for character 195 (complete character description in text). Arrows indicate the dorsal interosseal sulcus or homologous site. Scale bars = 5 mm.

196. Carpometacarpus, distal end, ventral view, distal synostosis of metacarpals, ventral interosseal sulcus (Fig. S140; Livezey, 1998: 253; Musser and Cracraft, 2019: 254):

0 – absent or inconspicuous

1 – prominent groove

197. Carpometacarpus, distal end, ventral view, cranial projection on major metacarpal (Fig. S146):

0 – weak
1 – prominent

**Figure S146.** Carpometacarpi of *Monias benschi* (**A**, left element, mirrored) and *Aegotheles cristatus* (**B**, right element) in ventral view, illustrating alternative states for character 197 (complete character description in text). Arrows indicate the cranial projection on major metacarpal, whereas dotted lines indicate the cranial extent of the rest of the major metacarpal. Scale bars = 5 mm.

198. Carpometacarpus, distal end, maximum length of synostosis between metacarpals, measured from distal end of intermetacarpal space to articular facet for minor digit (Fig. S128; Worthy and Lee, 2008: 84; Worthy et al., 2017: 171):

0 – short, length less than width measured just distad of intermetacarpal space

1 – long, length greater than width of synostosis

199. Major digit, phalanx 1, large proximally directed process on ventral side of proximal end (Fig. S147; Ksepka et al., 2019: 86):

0 – absent

1 – present

Phalanx characters scored based on *Colius* for *Urocolius*.

**Figure S147.** Manual phalanges II-1 of *Aegotheles cristatus* (**A**, right element) and *Jynx torquilla* (**B**, right element) in cranial view, illustrating alternative states for character 199 (complete character description in text). Arrows indicate the proximally directed process on ventral side of proximal end. Scale bars = 5 mm.

200. Major digit, phalanx 1, proximally hooked process projecting from caudal edge of distal end (Fig. S148; Ksepka et al., 2019: 87):

0 – absent

1 – present

**Figure S148.** Manual phalanges II-1 of *Aegotheles cristatus* (**A**, right element) and *Jynx torquilla* (**B**, right element) in dorsal view, illustrating alternative states for characters 200, 201, and 204 (complete character descriptions in text). Arrows for character 200 indicate the proximally hooked process on caudal edge of distal end or approximate homologous site to the process, arrows for character 201 indicate the internal index process, and arrows for character 204 indicate fenestration in the phalanx or approximate homologous site to the fenestra. Scale bars = 5 mm.

201. Major digit, phalanx 1, internal index process (Fig. S148; Mayr, 2010: 44; Ksepka et al., 2013: 77; Musser and Clarke, 2020: 496):

0 – poorly developed, does not project appreciably beyond rest of bone

1 – well developed, forming prominent lobe

202. Major digit, phalanx 1, proportions (Fig. S149; Mayr, 2004b: 46):

0 – short (ratio length to craniocaudal width less than 4.5)

1 – elongate (ratio length to craniocaudal width more than 4.5)

**Figure S149.** Manual phalanges II-1 of *Aegotheles cristatus* (**A**, right element) and *Podilymbus podiceps* (**B**, right element) in dorsal view, illustrating alternative states for character 202 (complete character description in text). Red lines the proximodistal length of the phalanx. Scale bars = 5 mm.

203. Major digit, phalanx 1, dorsal fossa (Fig. S150; Mayr, 2010: 43; Ksepka et al., 2013: 79):

0 – single depression

1 – divided into two depressions or fenestrae separated by a distinct oblique bulge

2 – depressions indistinct

**Figure S150.** Manual phalanges II-1 of *Corythaeola cristata* (**A**, right element), *Aegotheles cristatus* (**B**, right element), and *Jynx torquilla* (**C**, right element) in dorsal view, illustrating alternative states for character 203 (complete character description in text). Arrows indicate the depressions in the dorsal fossa, or approximate homologous site to the depressions. Scale bars = 5 mm.

204. Major digit, phalanx 1, fenestration (Fig. S148; Ksepka et al., 2013: 80; Musser and Clarke, 2020: 497):

0 – solid

1 – fenestrated
