## Appendix S3 for "Towards a comprehensive anatomical matrix for crown birds: phylogenetic insights from the pectoral girdle and forelimb skeleton"

**Appendix S3: List of Specimens Examined**

Institutional abbreviations: ALMNH, Alabama Museum of Natural History, University of Alabama, Tuscaloosa, AL, USA; FHSM, Sternberg Museum of Natural History, Hays, KS, USA; FMNH, Field Museum of Natural History, Chicago, IL, USA; KUVP, University of Kansas Biodiversity Institute and Natural History Museum, Lawrence, KS, USA; MSC and RMM, McWane Science Center (formerly Red Mountain Museum), Birmingham, AL, USA; NHMUK, Natural History Museum, Tring, UK; OUMNH, Oxford University Museum of Natural History, Oxford, UK; UMMZ, University of Michigan Museum of Zoology, Ann Arbor, MI, USA; UMZC, University Museum of Zoology, Cambridge, UK; YPM, Yale Peabody Museum, New Haven, CT, USA.

- *Ichthyornis dispar*: ALMNH PV 1988.20.427.1, ALMNH PV 1988.20.5, ALMNH PV 1993.2.133, FHSM VP 18702, KUVP 2281, KUVP 2300, KUVP 25469, KUVP 25471, KUVP 25472, KUVP 119673, KUVP 123459, MSC 34426, MSC 34427, NHMUK A 905, RMM 2841, RMM 3394, RMM 5794, RMM 5895, RMM 5916, RMM 5937, RMM 6200, RMM 6201, RMM 6202, RMM 7841, RMM 7842, RMM 7844, YPM 1450, YPM 1461, YPM 1724, YPM 1733, YPM 1740, YPM 1741.
- *Eudromia elegans*: UMMZ 156966.
- *Chauna chavaria*: OUMNH 23790.
- *Anseranas semipalmata*: NHMUK 1852.7.22.1.
- *Dendrocygna bicolor*: UMMZ 219885.
- *Alectura lathami*: NHMUK S2010.1.31.
- *Ortalis ruficauda*: UMMZ 155489.
- *Rollulus rouloul*: NHMUK 1871.7.20.87.
- *Phoenicopterus ruber*: UMZC 346.B.
- *Podilymbus podiceps*: UMMZ 205087.
- *Monias benschi*: NHMUK 1924.11.
- *Pterocles quadricinctus*: FMNH 319937.
- *Columba livia*: FMNH 347273.
- *Corythaeola cristata*: NHMUK 1923.11.12.363.
- *Ardeotis australis*: UMMZ 214165.
- *Tapera naevia*: UMMZ 222217.
- *Caprimulgus macrurus*: FMNH 392245.
- *Nyctibius griseus*: UMMZ 136371.
- *Podargus strigoides*: UMZC 493B.
- *Aegotheles cristatus*: YPM 124258.
- *Streptoprocne zonaris*: FMNH 85767.
- *Florisuga mellivora*: YPM 101545.
- *Opisthocomus hoazin*: NHMUK S1961.6.1.
- *Psophia crepitans*: FMNH 105783.
- *Aramus guarauna*: FMNH 376078.
- *Balearica pavonina*: NHMUK 1859.10.26.
- *Podica senegalensis*: UMZC 209A.
- *Sarothrura elegans*: NHMUK S1997.34.2.
- *Rallus limicola*: FMNH 501812.
- *Burhinus senegalensis*: FMNH 313704.
- *Charadrius vociferus*: FMNH 470173.
- *Rostratula benghalensis*: FMNH 319933.
- *Limosa lapponica*: NHMUK S1994.46.20.
- *Turnix varius*: NHMUK S1952.2.142.
- *Alca torda*: UMZC 187.AA.
- *Sterna hirundo*: NHMUK S1975.65.
- *Eurypyga helias*: FMNH 317341.
- *Phaethon lepturus*: NHMUK 1876.3.16.3.
- *Gavia stellata*: NHMUK 1891.7.20.132.
- *Spheniscus humboldti*: NHMUK S2000.7.1.
- *Phoebastria irrorata*: NHMUK S1963.28.4.
- *Oceanites oceanicus*: NHMUK S1957.11.2.
- *Pagodroma nivea*: NHMUK S1998.55.
- *Leptoptilos crumenifer*: NHMUK S1952.3.182.
- *Fregata aquila*: NHMUK 1890.11.3.3.
- *Sula dactylatra*: NHMUK 1890.11.3.10.
- *Leucocarbo atriceps*: NHMUK S2012.36.
- *Eudocimus ruber*: NHMUK S1999.8.1.
- *Scopus umbretta*: FMNH 313701.
- *Pelecanus occidentalis*: NHMUK S1973.66.16.
- *Tigrisoma lineatum*: FMNH 105528.
- *Coragyps atratus*: UMMZ 71891.
- *Pandion haliaetus*: FMNH 437336.
- *Elanus caeruleus*: NHMUK 1850.8.15.159.
- *Tyto alba*: NHMUK S1989.22.1.
- *Ninox novaeseelandiae*: UMZC 497.B.
- *Urocolius macrourus*: FMNH 368959.
- *Leptosomus discolor*: NHMUK A1968.30.38.
- *Trogon melanurus*: FMNH 290496.
- *Upupa epops*: FMNH 352821.
- *Bucorvus abyssinicus*: NHMUK S2006.31.20.
- *Coracias benghalensis*: NHMUK S1987.19.15.
- *Merops orientalis*: UMMZ 216577.
- *Alcedo atthis*: NHMUK S1994.7.1.
- *Bucco capensis*: FMNH 330305.
- *Psilopogon chrysopogon*: NHMUK 1850.8.15.
- *Jynx torquilla*: NHMUK S1986.36.10.
- *Cariama cristata*: FMNH 105653.
- *Micrastur ruficollis*: FMNH 330226.
- *Nestor notabilis*: FMNH 23530.
- *Psittacus erithacus*: NHMUK S1992.41.60.
- *Acanthisitta chloris*: NHMUK 1940.12.8.146.
- *Erythropitta erythrogaster*: FMNH 344970.
- *Neopelma chrysocephalum*: YPM 139662.
- *Menura novaehollandiae*: FMNH 336751.
- *Climacteris melanurus*: UMMZ 214303.
