## Appendix S5 for "Towards a comprehensive anatomical matrix for crown birds: phylogenetic insights from the pectoral girdle and forelimb skeleton": RF_calculations_code.R

```
library(ape)
library(geiger)
library(phangorn)

#Read in molecular trees

treeJarvis<-read.tree(file="Jarvisetal.tre")
treePrum<-read.tree(file="Prumetal.tre")
treeKuhl<-read.tree(file="Kuhletal.tre")
treeWu<-read.tree(file="Wuetal.tre")
treeStiller<-read.tree(file="Stilleretal.tre")

#Read in trees from analyses of pectoral dataset

tree1<-read.tree(file="Analysis1.tre")
tree2<-read.tree(file="Analysis2.tre")
tree3<-read.tree(file="Analysis3.tre")
tree4<-read.tree(file="Analysis4.tre")
tree5<-read.tree(file="Analysis5.tre")
tree6<-read.tree(file="Analysis6.tre")
tree7<-read.tree(file="Analysis7.tre")
tree8<-read.tree(file="Analysis8.tre")
tree9<-read.tree(file="Analysis9.tre")
tree10<-read.tree(file="Analysis10.tre")
tree11<-read.tree(file="Analysis11.tre")
tree12<-read.tree(file="Analysis12.tre")

#Calculate RF distances for trees from Analysis 1

RF.dist.Jarvis.tree1=RF.dist(tree1,treeJarvis)
RF.dist.Prum.tree1=RF.dist(tree1,treePrum)
RF.dist.Kuhl.tree1=RF.dist(tree1,treeKuhl)
RF.dist.Wu.tree1=RF.dist(tree1,treeWu)
RF.dist.Stiller.tree1=RF.dist(tree1,treeStiller)

#Calculate RF distances for tree from Analysis 2

RF.dist.Jarvis.tree2=RF.dist(tree2,treeJarvis)
RF.dist.Prum.tree2=RF.dist(tree2,treePrum)
RF.dist.Kuhl.tree2=RF.dist(tree2,treeKuhl)
RF.dist.Wu.tree2=RF.dist(tree2,treeWu)
RF.dist.Stiller.tree2=RF.dist(tree2,treeStiller)

#Calculate RF distances for tree from Analysis 3

RF.dist.Jarvis.tree3=RF.dist(tree3,treeJarvis)
RF.dist.Prum.tree3=RF.dist(tree3,treePrum)
RF.dist.Kuhl.tree3=RF.dist(tree3,treeKuhl)
RF.dist.Wu.tree3=RF.dist(tree3,treeWu)
RF.dist.Stiller.tree3=RF.dist(tree3,treeStiller)

#Calculate RF distances for trees from Analysis 4
#Summarize function used when a range of values is obtained from different MPTs

RF.dist.Jarvis.tree4=RF.dist(tree4,treeJarvis)
summary(RF.dist.Jarvis.tree4)
RF.dist.Prum.tree4=RF.dist(tree4,treePrum)
summary(RF.dist.Prum.tree4)
RF.dist.Kuhl.tree4=RF.dist(tree4,treeKuhl)
summary(RF.dist.Kuhl.tree4)
RF.dist.Wu.tree4=RF.dist(tree4,treeWu)
summary(RF.dist.Wu.tree4)
RF.dist.Stiller.tree4=RF.dist(tree4,treeStiller)
summary(RF.dist.Stiller.tree4)

#Calculate RF distances for tree from Analysis 5

RF.dist.Jarvis.tree5=RF.dist(tree5,treeJarvis)
RF.dist.Prum.tree5=RF.dist(tree5,treePrum)
RF.dist.Kuhl.tree5=RF.dist(tree5,treeKuhl)
RF.dist.Wu.tree5=RF.dist(tree5,treeWu)
RF.dist.Stiller.tree5=RF.dist(tree5,treeStiller)

#Calculate RF distances for tree from Analysis 6

RF.dist.Jarvis.tree6=RF.dist(tree6,treeJarvis)
RF.dist.Prum.tree6=RF.dist(tree6,treePrum)
RF.dist.Kuhl.tree6=RF.dist(tree6,treeKuhl)
RF.dist.Wu.tree6=RF.dist(tree6,treeWu)
RF.dist.Stiller.tree6=RF.dist(tree6,treeStiller)

#Calculate RF distances for trees from Analysis 7
#Summarize function used when a range of values is obtained from different MPTs

RF.dist.Jarvis.tree7=RF.dist(tree7,treeJarvis)
RF.dist.Prum.tree7=RF.dist(tree7,treePrum)
summary(RF.dist.Prum.tree7)
RF.dist.Kuhl.tree7=RF.dist(tree7,treeKuhl)
RF.dist.Wu.tree7=RF.dist(tree7,treeWu)
RF.dist.Stiller.tree7=RF.dist(tree7,treeStiller)

#Calculate RF distances for tree from Analysis 8

RF.dist.Jarvis.tree8=RF.dist(tree8,treeJarvis)
RF.dist.Prum.tree8=RF.dist(tree8,treePrum)
RF.dist.Kuhl.tree8=RF.dist(tree8,treeKuhl)
RF.dist.Wu.tree8=RF.dist(tree8,treeWu)
RF.dist.Stiller.tree8=RF.dist(tree8,treeStiller)

#Calculate RF distances for tree from Analysis 9

RF.dist.Jarvis.tree9=RF.dist(tree9,treeJarvis)
RF.dist.Prum.tree9=RF.dist(tree9,treePrum)
RF.dist.Kuhl.tree9=RF.dist(tree9,treeKuhl)
RF.dist.Wu.tree9=RF.dist(tree9,treeWu)
RF.dist.Stiller.tree9=RF.dist(tree9,treeStiller)

#Calculate RF distances for tree from Analysis 10

RF.dist.Jarvis.tree10=RF.dist(tree10,treeJarvis)
RF.dist.Prum.tree10=RF.dist(tree10,treePrum)
RF.dist.Kuhl.tree10=RF.dist(tree10,treeKuhl)
RF.dist.Wu.tree10=RF.dist(tree10,treeWu)
RF.dist.Stiller.tree10=RF.dist(tree10,treeStiller)

#Calculate RF distances for tree from Analysis 11

RF.dist.Jarvis.tree11=RF.dist(tree11,treeJarvis)
RF.dist.Prum.tree11=RF.dist(tree11,treePrum)
RF.dist.Kuhl.tree11=RF.dist(tree11,treeKuhl)
RF.dist.Wu.tree11=RF.dist(tree11,treeWu)
RF.dist.Stiller.tree11=RF.dist(tree11,treeStiller)

#Calculate RF distances for tree from Analysis 12

RF.dist.Jarvis.tree12=RF.dist(tree12,treeJarvis)
RF.dist.Prum.tree12=RF.dist(tree12,treePrum)
RF.dist.Kuhl.tree12=RF.dist(tree12,treeKuhl)
RF.dist.Wu.tree12=RF.dist(tree12,treeWu)
RF.dist.Stiller.tree12=RF.dist(tree12,treeStiller)
```
