## Appendix S6 for "Towards a comprehensive anatomical matrix for crown birds: phylogenetic insights from the pectoral girdle and forelimb skeleton": Similarity_calculations_code.R

```
#Procedures follow those of Evers and Benson (2019)
#Load character-taxon matrix 

data.matrix<-read.table("Similarity_matrix.txt", header=T, na.strings=c("-","?"))

#Transpose matrix

data.matrix<-t(data.matrix)

#Make a matrix of same dimensions is which the entries are numerical

data2<-matrix(nrow=dim(data.matrix)[1],ncol=dim(data.matrix)[2])
for(i in 1:ncol(data.matrix)) data2[,i]<-as.numeric(data.matrix[,i])
colnames(data2)<-colnames(data.matrix)

#Create a dissimilarity matrix based on 'data2' with taxa in numeric columns

sim.data<-matrix(nrow=ncol(data2),ncol=ncol(data2))
rownames(sim.data)<-colnames(data2)
colnames(sim.data)<-colnames(data2)

#Calculate dissmilarity values

for(i in 1:ncol(data2)) {for(j in 1:ncol(data2)) {sim.data[j,i]<-length(grep(TRUE,data2[,j]==data2[,i]))/length(grep(TRUE,complete.cases(data2[,j]==data2[,i])))}}

#Load required packages for coloring matrices

require(gclus)

#Load packages for additional colors

library(RColorBrewer)
display.brewer.all(type="seq")

#Assign colours to the similarity matrix

hm.cols<-dmat.color(1-sim.data, byrank=FALSE, colors=brewer.pal(9, "PuRd"))

#Plot and export matrix as PDF

dev.new()
pdf("Heatmap.Sim.Matrix.pdf", useDingbats=FALSE)
plotcolors(hm.cols,rlabels=rownames(sim.data),clabels=colnames(sim.data))
dev.off()

#Subsetting data to only sternal characters

sternum.data<-data.matrix[14:60,]

#Calculate percentages of missing data
#No taxon with 10% or more missing data present for sternal data, so nothing else done, but see furcular subset for a case in which a taxon was removed

perc.NA<-as.matrix(apply(sternum.data, 2, function(col)sum(is.na(col)/length(col))))

#Make a matrix of same dimensions in which the entries are numerical

data3<-matrix(nrow=dim(sternum.data)[1],ncol=dim(sternum.data)[2])
for(i in 1:ncol(sternum.data)) data3[,i]<-as.numeric(sternum.data[,i])
colnames(data3)<-colnames(sternum.data)

#Create a similarity matrix based on 'data3' with taxa in numeric columns

sim.data.sternum<-matrix(nrow=ncol(data3),ncol=ncol(data3))
rownames(sim.data.sternum)<-colnames(data3)
colnames(sim.data.sternum)<-colnames(data3)

#Calculate smilarity values

for(i in 1:ncol(data3)) {for(j in 1:ncol(data3)) {sim.data.sternum[j,i]<-length(grep(TRUE,data3[,j]==data3[,i]))/length(grep(TRUE,complete.cases(data3[,j]==data3[,i])))}}

#Plot sternal similarity data as heatmap

hm.cols2<-dmat.color(1-sim.data.sternum, byrank=FALSE, colors=brewer.pal(9, "PuRd"))
dev.new()
pdf("Heatmap.Sim.Matrix.sternum.pdf", useDingbats=FALSE)
plotcolors(hm.cols2,rlabels=rownames(sim.data.sternum),clabels=colnames(sim.data.sternum))
dev.off()

#Subsetting data to only coracoid characters

coracoid.data<-data.matrix[62:87,]

#Calculate percentages of missing data

perc.NA<-as.matrix(apply(coracoid.data, 2, function(col)sum(is.na(col)/length(col))))

#Make a matrix of same dimensions in which the entries are numerical

data4<-matrix(nrow=dim(coracoid.data)[1],ncol=dim(coracoid.data)[2])
for(i in 1:ncol(coracoid.data)) data4[,i]<-as.numeric(coracoid.data[,i])
colnames(data4)<-colnames(coracoid.data)

#Create a similarity matrix based on 'data4' with taxa in numeric columns

sim.data.coracoid<-matrix(nrow=ncol(data4),ncol=ncol(data4))
rownames(sim.data.coracoid)<-colnames(data4)
colnames(sim.data.coracoid)<-colnames(data4)

#Calculate smilarity values

for(i in 1:ncol(data4)) {for(j in 1:ncol(data4)) {sim.data.coracoid[j,i]<-length(grep(TRUE,data4[,j]==data4[,i]))/length(grep(TRUE,complete.cases(data4[,j]==data4[,i])))}}

#Plot coracoid similarity data as heatmap

hm.cols3<-dmat.color(1-sim.data.coracoid, byrank=FALSE, colors=brewer.pal(9, "PuRd"))
dev.new()
pdf("Heatmap.Sim.Matrix.coracoid.pdf", useDingbats=FALSE)
plotcolors(hm.cols3,rlabels=rownames(sim.data.coracoid),clabels=colnames(sim.data.coracoid))
dev.off()

#Subsetting data to only humeral characters

humerus.data<-data.matrix[c(97, 100:143),]

#Calculate percentages of missing data

perc.NA<-as.matrix(apply(humerus.data, 2, function(col)sum(is.na(col)/length(col))))

#Make a matrix of same dimensions in which the entries are numerical

data5<-matrix(nrow=dim(humerus.data)[1],ncol=dim(humerus.data)[2])
for(i in 1:ncol(humerus.data)) data5[,i]<-as.numeric(humerus.data[,i])
colnames(data5)<-colnames(humerus.data)

#Create a similarity matrix based on 'data5' with taxa in numeric columns

sim.data.humerus<-matrix(nrow=ncol(data5),ncol=ncol(data5))
rownames(sim.data.humerus)<-colnames(data5)
colnames(sim.data.humerus)<-colnames(data5)

#Calculate smilarity values

for(i in 1:ncol(data5)) {for(j in 1:ncol(data5)) {sim.data.humerus[j,i]<-length(grep(TRUE,data5[,j]==data5[,i]))/length(grep(TRUE,complete.cases(data5[,j]==data5[,i])))}}

#Plot humeral similarity data as heatmap

hm.cols4<-dmat.color(1-sim.data.humerus, byrank=FALSE, colors=brewer.pal(9, "PuRd"))
dev.new()
pdf("Heatmap.Sim.Matrix.humerus.pdf", useDingbats=FALSE)
plotcolors(hm.cols4,rlabels=rownames(sim.data.humerus),clabels=colnames(sim.data.humerus))
dev.off()

#Subsetting data to only ulnar characters

ulna.data<-data.matrix[c(98, 150:163),]

#Calculate percentages of missing data

perc.NA<-as.matrix(apply(ulna.data, 2, function(col)sum(is.na(col)/length(col))))

#Make a matrix of same dimensions in which the entries are numerical

data6<-matrix(nrow=dim(ulna.data)[1],ncol=dim(ulna.data)[2])
for(i in 1:ncol(ulna.data)) data6[,i]<-as.numeric(ulna.data[,i])
colnames(data6)<-colnames(ulna.data)

#Create a similarity matrix based on 'data6' with taxa in numeric columns

sim.data.ulna<-matrix(nrow=ncol(data6),ncol=ncol(data6))
rownames(sim.data.ulna)<-colnames(data6)
colnames(sim.data.ulna)<-colnames(data6)

#Calculate smilarity values

for(i in 1:ncol(data6)) {for(j in 1:ncol(data6)) {sim.data.ulna[j,i]<-length(grep(TRUE,data6[,j]==data6[,i]))/length(grep(TRUE,complete.cases(data6[,j]==data6[,i])))}}

#Plot ulnar similarity data as heatmap

hm.cols5<-dmat.color(1-sim.data.ulna, byrank=FALSE, colors=brewer.pal(9, "PuRd"))
dev.new()
pdf("Heatmap.Sim.Matrix.ulna.pdf", useDingbats=FALSE)
plotcolors(hm.cols5,rlabels=rownames(sim.data.ulna),clabels=colnames(sim.data.ulna))
dev.off()

#Subsetting data to only carpometacarpal characters

carpometacarpus.data<-data.matrix[c(99, 169:198),]

#Calculate percentages of missing data

perc.NA<-as.matrix(apply(carpometacarpus.data, 2, function(col)sum(is.na(col)/length(col))))

#Make a matrix of same dimensions in which the entries are numerical

data7<-matrix(nrow=dim(carpometacarpus.data)[1],ncol=dim(carpometacarpus.data)[2])
for(i in 1:ncol(carpometacarpus.data)) data7[,i]<-as.numeric(carpometacarpus.data[,i])
colnames(data7)<-colnames(carpometacarpus.data)

#Create a similarity matrix based on 'data7' with taxa in numeric columns

sim.data.carpometacarpus<-matrix(nrow=ncol(data7),ncol=ncol(data7))
rownames(sim.data.carpometacarpus)<-colnames(data7)
colnames(sim.data.carpometacarpus)<-colnames(data7)

#Calculate smilarity values

for(i in 1:ncol(data7)) {for(j in 1:ncol(data7)) {sim.data.carpometacarpus[j,i]<-length(grep(TRUE,data7[,j]==data7[,i]))/length(grep(TRUE,complete.cases(data7[,j]==data7[,i])))}}

#Plot carpometacarpal similarity data as heatmap

hm.cols6<-dmat.color(1-sim.data.carpometacarpus, byrank=FALSE, colors=brewer.pal(9, "PuRd"))
dev.new()
pdf("Heatmap.Sim.Matrix.carpometacarpus.pdf", useDingbats=FALSE)
plotcolors(hm.cols6,rlabels=rownames(sim.data.carpometacarpus),clabels=colnames(sim.data.carpometacarpus))
dev.off()

#Subsetting data to only furcular characters

furcula.data<-data.matrix[1:13,]

#Calculate percentages of missing data

perc.NA<-as.matrix(apply(furcula.data, 2, function(col)sum(is.na(col)/length(col))))
perc.NA>=0.9

#Create object with names of taxa with more than 90% missing data

drop.taxa<-which(perc.NA %% 2 >= 0.9, arr.ind=TRUE, useNames=TRUE)
drop.taxa<-attributes(drop.taxa)$dimnames[[1]]

#Create object with numbers of columns to be dropped from dataset

drop.cols<-which(match(colnames(furcula.data), drop.taxa) != "NA")

#Create reduced dataset that only contains taxa with more than 90% scored characters

furcula.data.red <- furcula.data[,-drop.cols]

#Make a matrix of same dimensions in which the entries are numerical

data8<-matrix(nrow=dim(furcula.data.red)[1],ncol=dim(furcula.data.red)[2])
for(i in 1:ncol(furcula.data.red)) data8[,i]<-as.numeric(furcula.data.red[,i])
colnames(data8)<-colnames(furcula.data.red)

#Create a similarity matrix based on 'data8' with taxa in numeric columns

sim.data.furcula.red<-matrix(nrow=ncol(data8),ncol=ncol(data8))
rownames(sim.data.furcula.red)<-colnames(data8)
colnames(sim.data.furcula.red)<-colnames(data8)

#Calculate smilarity values

for(i in 1:ncol(data8)) {for(j in 1:ncol(data8)) {sim.data.furcula.red[j,i]<-length(grep(TRUE,data8[,j]==data8[,i]))/length(grep(TRUE,complete.cases(data8[,j]==data8[,i])))}}

#Plot furcular similarity data as heatmap

hm.cols7<-dmat.color(1-sim.data.furcula.red, byrank=FALSE, colors=brewer.pal(9, "PuRd"))
dev.new()
pdf("Heatmap.Sim.Matrix.furcula.pdf", useDingbats=FALSE)
plotcolors(hm.cols7,rlabels=rownames(sim.data.furcula.red),clabels=colnames(sim.data.furcula.red))
dev.off()
```
