## Appendix S7 for "Towards a comprehensive anatomical matrix for crown birds: phylogenetic insights from the pectoral girdle and forelimb skeleton": Homoplasy_analyses_Jun24-edit.R

```
###### 2024-Jun-11 ######

# Pectoral girdle homoplasy analyses

packages <- c("ape", "phangorn", "phytools", "stringr", "TreeTools", "geiger")
lapply(packages, library, character.only=TRUE)

### Pectoral dataset ###

# Trees
prum <- read.tree("Prumetal.phy")
jarv <- read.tree("Jarvisetal.phy")
kuhl <- read.tree("Kuhletal.phy")
wu <- read.tree("Wuetal.phy")
still <- read.tree("Stilleretal.phy") 

# Matrix
pec <- ReadCharacters("pectoral_simple.nex")

get.states(pec)
contrast.matrix(pec)
statef(pec) # Invariant character

# Remove invariant characters
pec <- remove.invar(pec) #38

dim(pec)

pec_phydat <- morph.phydat(pec)

rhi_prum <- RHI(pec_phydat, prum, 1000)
rhi_jarv <- RHI(pec_phydat, jarv, 1000)
rhi_kuhl <- RHI(pec_phydat, kuhl, 1000)
rhi_wu <- RHI(pec_phydat, wu, 1000)
rhi_still <- RHI(pec_phydat, still, 1000)

ci_prum <- CI(prum, pec_phydat)
ri_prum <- RI(prum, pec_phydat)

ci_jarv <- CI(jarv, pec_phydat)
ri_jarv <- RI(jarv, pec_phydat)

ci_kuhl <- CI(kuhl, pec_phydat)
ri_kuhl <- RI(kuhl, pec_phydat)

ri_wu <- RI(wu, pec_phydat)
ci_wu <- CI(wu, pec_phydat)

ri_still <- RI(still, pec_phydat)
ci_still <- CI(still, pec_phydat)

### Calculate RHI for ordered characters

# Ordered characters

ord <- c(9, 10, 15, 17, 24, 35, 36, 50, 53, 54, 66, 68, 72, 78, 97, 104, 115, 140, 166, 177, 182, 191, 192)

# after character 38 is removed, adjust numbers

ord <- c(9, 10, 15, 17, 24, 35, 36, 50, 53, 54, 66, 68, 72, 78, 97, 104, 115, 140, 166, 177, 182, 191, 192)

ord_ <- c(50, 53, 54, 66, 68, 72, 78, 97, 104, 115, 140, 166, 177, 182, 191, 192)-1

ord_1 <- c(c(9, 10, 15, 17, 24, 35, 36), ord_)

# Make a cost matrix

cost <- cost.matrix(pec)

rhi_prum_ord <- RHI(pec_phydat, prum, 1000, ord=ord_1, cost=cost)
rhi_jarv_ord <- RHI(pec_phydat, jarv, 1000, ord=ord_1, cost=cost)
rhi_kuhl_ord <- RHI(pec_phydat, kuhl, 1000, ord=ord_1, cost=cost)
rhi_wu_ord <- RHI(pec_phydat, wu, 1000, ord=ord_1, cost=cost)
rhi_still_ord <- RHI(pec_phydat, still, 1000, ord=ord_1, cost=cost)

#####

### Strisores Chen et al 2019 ###

chen <- read.tree("strisoresconstraint-new.phy")

strisores <- ReadCharacters("strisores-simple.nex")

get.states(strisores)
contrast.matrix(strisores)
statef(strisores) # One char with one state
strisores <- remove.invar(strisores)
 # Remove 98, 104, 117
dim(strisores)

stris_phydat <- morph.phydat(strisores)

rhi_chen <- RHI(stris_phydat, chen, 1000)

ci_chen <- CI(chen, stris_phydat)
ri_chen <- RI(chen, stris_phydat)

### Passerines Steell et al 2023 ###

passerines <- ReadCharacters("Passeriformes_matrix_a.nex")

steell <- read.tree("Passeriformes_a_tree.phy")

# Remove fossil taxon Zygodactylus
steell <- drop.tip(steell, "Zygodactylus_luberonensis_SMFAv519")
name.check(steell, passerines)
passerines <- passerines[rownames(passerines) != "Zygodactylus_luberonensis_SMFAv519",]
name.check(steell, passerines)
statef(passerines)

dim(passerines)

passerine_phydat <- morph.phydat(passerines)

rhi_steell <- RHI(passerine_phydat, steell, 1000)
ri_steell <- RI(steell, passerine_phydat)
ci_steell <- CI(steell, passerine_phydat)

#####

### Musser and Clarke 2020 - Charadriformes/Rails/Neornithes

# Matrix
charad <- ReadCharacters("Musser_Clarke_2020_simple.nex")

# Tree with Prum constraint
musser_prum <- read.tree("Musser_Clarke_2020_simple-Prum.phy")

charad <- as.data.frame(charad)
name.check(musser_prum, charad)
charad <- charad[musser_prum$tip.label,]
charad <- as.matrix(charad)

statef(charad) # One invariant
charad <- remove.invar(charad)
# Remove 81
dim(charad)

charad_phydat <- morph.phydat(charad)

rhi_musser_prum <- RHI(charad_phydat, musser_prum, 1000)
ci_musser_prum <- CI(musser_prum, charad_phydat)
ri_musser_prum <- RI(musser_prum, charad_phydat)

# Tree with Jarvis constraint
musser_jarv <- read.tree("Musser_Clarke_2020_simple-Jarvis.phy")
name.check(musser_jarv, charad)

rhi_musser_jarv <- RHI(charad_phydat, musser_jarv, 1000)
ci_musser_jarv <- CI(musser_jarv, charad_phydat)
ri_musser_jarv <- RI(musser_jarv, charad_phydat)

# Stiller constraint

musser_still <- read.tree("Musser_Clarke_2020_simple-Stiller.phy")

rhi_musser_still <- RHI(charad_phydat, musser_still, 1000)
ci_musser_still <- CI(musser_still, charad_phydat)
ri_musser_still <- RI(musser_still, charad_phydat)

######

### Ksepka et al 2022 - Galliformes ###

galliformes <- ReadCharacters("Galliformes_Ksepka2022_simple.nex")
get.states(galliformes)
contrast.matrix(galliformes)

ksepka_2022 <- read.tree("Galliformes_Ksepka2022.phy")
name.check(ksepka_2022, galliformes)
galliformes <- as.data.frame(galliformes)
galliformes <- galliformes[ksepka_2022$tip.label,]
galliformes <- as.matrix(galliformes)
statef(galliformes) # invariants present
galliformes <- remove.invar(galliformes)
#32


dim(galliformes)

gall_phydat <- morph.phydat(galliformes)

rhi_ksepka_gall <- RHI(gall_phydat, ksepka_2022, 1000)
ci_ksepka_gall <- CI(ksepka_2022, gall_phydat)
ri_ksepka_gall <- RI(ksepka_2022, gall_phydat)

#####

### Field et al 2020 - Neornithes ###

neornithes <- ReadCharacters("Neornithes_Field_2020.nex")

### Prum tree

field2020_prum <- read.tree("Neornithes_Field_2020-Prum.phy")

name.check(field2020_prum, neornithes)

# Remove taxa from matrix
neornithes <- as.data.frame(neornithes)
neornithes <- neornithes[field2020_prum$tip.label,]
neornithes <- as.matrix(neornithes)

statef(neornithes) # invariants present
neornithes <- remove.invar(neornithes)
#194, 201, 282, 287

dim(neornithes)

neorn_phydat <- morph.phydat(neornithes)

rhi_field_neorn_prum <- RHI(neorn_phydat, field2020_prum, 1000)
ci_field_neorn_prum <- CI(field2020_prum, neorn_phydat)
ri_field_neorn_prum <- RI(field2020_prum, neorn_phydat)

### Jarvis tree

field2020_jarv <- read.tree("Neornithes_Field_2020-Jarvis.phy")
name.check(field2020_jarv, neornithes)

rhi_field_neorn_jarv <- RHI(neorn_phydat, field2020_jarv, 1000)
ci_field_neorn_jarv <- CI(field2020_jarv, neorn_phydat)
ri_field_neorn_jarv <- RI(field2020_jarv, neorn_phydat)

#####

# Livesy and Zusi 2006 - Neornithes

lz_neorn <- ReadCharacters("Livsey_Zusi_2007_simple.nex")
dim(lz_neorn)

### Prum

lz_neorn_prum <- read.tree("Livsey_Zusi_2007_Prum.phy")

name.check(lz_neorn_prum, lz_neorn)

# Remove taxa from matrix
lz_neorn <- as.data.frame(lz_neorn)
lz_neorn <- lz_neorn[lz_neorn_prum$tip.label,]
lz_neorn <- as.matrix(lz_neorn)

get.states(lz_neorn)
statef(lz_neorn) # 'N' comes up as a state, so replace with ?

lz_neorn[lz_neorn == "N"] <- "?"
get.states(lz_neorn)
statef(lz_neorn)

lz_neorn <- remove.invar(lz_neorn)
# The normal method didn't work because this matrix has NAs.

# Need to delete chars with sd = 0 and sd = NA

vals <- c()
for(i in 1:ncol(lz_neorn)){
	vals[[i]] <- sd(lz_neorn[,i], na.rm=TRUE)
	}
vals <- unlist(vals)

vals[is.na(vals)] <- 0
range(vals) # NAs removed

zeros <- c()
for(i in 1:length(vals)){
	if (vals[i] == 0){
		zeros[[i]] <- i
	}
}

zeros <- unlist(zeros)
length(zeros) #659

lz_neorn <- lz_neorn[,-zeros]
dim(lz_neorn)
statef(lz_neorn) # Problem solved
get.states(lz_neorn)
contrast.matrix(lz_neorn)

lz_phydat <- morph.phydat(lz_neorn)

rhi_lz_prum <- RHI(lz_phydat, lz_neorn_prum, 1000)
ci_lz_prum <- CI(lz_neorn_prum, lz_phydat)
ri_lz_prum <- RI(lz_neorn_prum, lz_phydat)

# Jarvis

lz_neorn_jarv <- read.tree("Livsey_Zusi_2007_Jarvis.phy")

name.check(lz_neorn_jarv, lz_neorn)

rhi_lz_jarv <- RHI(lz_phydat, lz_neorn_jarv, 1000)
ci_lz_jarv <- CI(lz_neorn_jarv, lz_phydat)
ri_lz_jarv <- RI(lz_neorn_jarv, lz_phydat)

lz_neorn_still <- read.tree("Livsey_Zusi_2007_simple-Stiller.phy")

rhi_lz_still <- RHI(lz_phydat, lz_neorn_still, 1000)
ci_lz_still <- CI(lz_neorn_still, lz_phydat)
ri_lz_still <- RI(lz_neorn_still, lz_phydat)


#####

### Ksepka et al 2019 Telluraves

# Prum constraint topology (from original paper) and Jarvis constraint tree (made by hand)

ksepka_tell_prum <- read.tree("Ksepka_2019_Telluraves_prum.phy")

ksepka_tell_jarv <- read.tree("Ksepka_2019_jarvis.phy")

ksepka_tell <- ReadCharacters("Ksepka_Telluraves_matrix.nex")

# Remove taxa from matrix
name.check(ksepka_tell_prum, ksepka_tell)

ksepka_tell <- as.data.frame(ksepka_tell)
ksepka_tell <- ksepka_tell[ksepka_tell_prum$tip.label,]
ksepka_tell <- as.matrix(ksepka_tell)

statef(ksepka_tell) # invariants present
ksepka_tell <- remove.invar(ksepka_tell)
# 28, 85

dim(ksepka_tell)

tell_phydat <- morph.phydat(ksepka_tell)

rhi_tell_jarv <- RHI(tell_phydat, ksepka_tell_jarv, 1000)
ci_tell_jarv <- CI(ksepka_tell_jarv, tell_phydat)
ri_tell_jarv <- RI(ksepka_tell_jarv, tell_phydat)

rhi_tell_prum <- RHI(tell_phydat, ksepka_tell_prum, 1000)
ci_tell_prum <- CI(ksepka_tell_prum, tell_phydat)
ri_tell_prum <- RI(ksepka_tell_prum, tell_phydat)

### Mayr & Clarke 2003

mayrclarke <- ReadCharacters("Mayr_Clarke_2003.nex")

mayrclarke_jarv <- read.tree("Mayr_Clarke_2003-Jarvis.phy")

mayrclarke_prum <- read.tree("Mayr_Clarke_2003-Prum.phy")

mayrclarke_stiller <- read.tree("Mayr_Clarke_2003-Stiller.phy")


name.check(mayrclarke_jarv, mayrclarke)

#remove taxa
mayrclarke <- as.data.frame(mayrclarke)
mayrclarke <- mayrclarke[mayrclarke_jarv$tip.label,]
mayrclarke <- as.matrix(mayrclarke)

statef(mayrclarke)
mayrclarke <- remove.invar(mayrclarke)
#1, 41, 42

dim(mayrclarke)

contrast.matrix(mayrclarke)
mayrclarke_phydat <- morph.phydat(mayrclarke)

rhi_mc_jarv <- RHI(mayrclarke_phydat, mayrclarke_jarv, 1000)
ci_mc_jarv <- CI(mayrclarke_jarv, mayrclarke_phydat)
ri_mc_jarv <- RI(mayrclarke_jarv, mayrclarke_phydat)

rhi_mc_prum <- RHI(mayrclarke_phydat, mayrclarke_prum, 1000)
ci_mc_prum <- CI(mayrclarke_prum, mayrclarke_phydat)
ri_mc_prum <- RI(mayrclarke_prum, mayrclarke_phydat)

rhi_mc_stiller <- RHI(mayrclarke_phydat, mayrclarke_stiller, 1000)
ci_mc_stiller <- CI(mayrclarke_stiller, mayrclarke_phydat)
ri_mc_stiller <- RI(mayrclarke_stiller, mayrclarke_phydat)


### Ksepka et al 2023 penguins

penguins <- ReadCharacters("PenguinMatrix.nex")

penguin_tree <- read.tree("PenguinMatrix.phy")

name.check(penguin_tree, penguins)
penguins <- as.data.frame(penguins)
penguins <- penguins[penguin_tree$tip.label,]
penguins <- as.matrix(penguins)

statef(penguins)
penguins <- remove.invar(penguins)
#[1] "Following invariant characters removed:"
 #[1]  19  79  89  96 101 115 118 126 130 144 151 157 164 173 176 179 181 183 184 192 199
#[22] 202 206 216 224 225 228 230 242 243 249 250 253 255 279

dim(penguins)

contrast.matrix(penguins)

peng_phydat <- morph.phydat(penguins)

peng_ci <- CI(penguin_tree, peng_phydat)
peng_ri <- RI(penguin_tree, peng_phydat)
peng_rhi <- RHI(peng_phydat, penguin_tree, 1000)

########

### Plot

# Pectoral matrix values
rhi_jarv
rhi_prum
rhi_kuhl
rhi_wu
rhi_still

# Mayr & Clarke 2003
rhi_mc_jarv
rhi_mc_prum
rhi_mc_stiller

# Livezy & Zusi 2007
rhi_lz_jarv
rhi_lz_prum
rhi_lz_still

# Field et al 2020
rhi_field_neorn_jarv
rhi_field_neorn_prum

# Musser & Clarke 2020
rhi_musser_jarv
rhi_musser_prum
rhi_musser_still

# Chen et al 2019
rhi_chen

# Ksepka et al 2019
rhi_tell_jarv
rhi_tell_prum

# Ksepka et al 2022
rhi_ksepka_gall

# Ksepka et al 2023
peng_rhi

# Steell et al 2023
rhi_steell

# RHI values (this is definitely the long-winded version...)

rhi_vals <- c(rhi_jarv[2], rhi_prum[2], rhi_kuhl[2], rhi_wu[2], rhi_still[2], rhi_mc_jarv[2], rhi_mc_prum[2], rhi_mc_stiller[2], rhi_lz_jarv[2], rhi_lz_prum[2], rhi_lz_still[2], rhi_field_neorn_jarv[2], rhi_field_neorn_prum[2], rhi_musser_jarv[2], rhi_musser_prum[2], rhi_musser_still[2], rhi_chen[2], rhi_tell_jarv[2], rhi_tell_prum[2], rhi_ksepka_gall[2], peng_rhi[2], rhi_steell[2])

vals5 <- c(rhi_jarv[1], rhi_prum[1], rhi_kuhl[1], rhi_wu[1], rhi_still[1], rhi_mc_jarv[1], rhi_mc_prum[1], rhi_mc_stiller[1], rhi_lz_jarv[1], rhi_lz_prum[1], rhi_lz_still[1], rhi_field_neorn_jarv[1], rhi_field_neorn_prum[1], rhi_musser_jarv[1], rhi_musser_prum[1], rhi_musser_still[1], rhi_chen[1], rhi_tell_jarv[1], rhi_tell_prum[1], rhi_ksepka_gall[1], peng_rhi[1], rhi_steell[1])

vals95 <- c(rhi_jarv[3], rhi_prum[3], rhi_kuhl[3], rhi_wu[3], rhi_still[3], rhi_mc_jarv[3], rhi_mc_prum[3], rhi_mc_stiller[3], rhi_lz_jarv[3], rhi_lz_prum[3], rhi_lz_still[3], rhi_field_neorn_jarv[3], rhi_field_neorn_prum[3], rhi_musser_jarv[3], rhi_musser_prum[3], rhi_musser_still[3], rhi_chen[3], rhi_tell_jarv[3], rhi_tell_prum[3], rhi_ksepka_gall[3], peng_rhi[3], rhi_steell[3])

names(rhi_vals) <- c("Pectoral", "Pectoral", "Pectoral", "Pectoral", "Pectoral", "Mayr & Clarke 2003", "Mayr & Clarke 2003", "Mayr & Clarke 2003", "Livezy & Zusi 2007", "Livezy & Zusi 2007", "Livezy & Zusi 2007", "Field et al 2020", "Field et al 2020", "Musser & Clarke 2020", "Musser & Clarke 2020", "Musser & Clarke 2020", "Chen et al 2019", "Ksepka et al 2019", "Ksepka et al 2019", "Ksepka et al 2022", "Ksepka et al 2023", "Steell et al 2023")

# COLOUR BY MATRIX
pec <- c("dark gray")
pec <- sample(pec_cols, 5, replace=TRUE)

mc <- c("dark green")
mc <- sample(mc, 3, replace=TRUE)

lz <- c("light blue")
lz <- sample(lz, 3, replace=TRUE)

field <- c("pink")
field <- sample(field, 2, replace=T)

musser <- c("light green")
musser <- sample(musser, 3, replace=T)

chen <- c("purple")

tell <- c("dark blue")
tell <- sample(tell, 2, replace=T)

gall <- c("red")

peng <- c("cyan")

steell <- c("orange")

cols <- c(pec, mc, lz, field, musser, chen, tell, gall, peng, steell)

rhis <- rbind(rhi_vals, vals5, vals95, cols)
class(rhis)
dim(rhis)

# COLOUR BY TOPOLOGY

jarv <- c("orange")
prum <- c("dark green")
kuhl <- ("purple")
wu <- c("light green")
still <- c("light blue")
other <- c("grey")

cols <- c(jarv, prum, kuhl, wu, still, jarv, prum, still, jarv, prum, still, jarv, prum, jarv, prum, still, other, jarv, prum, other, other, other)

legend <- c(jarv, prum, kuhl, wu, still, other)
names(legend) <- c("Jarvis et al 2014", "Prum et al 2015", "Kuhl et al 2021", "Wu et al 2024", "Stiller et al 2024", "Other")


bars <- barplot(rhi_vals, ylab="Relative Homoplasy", ylim=c(0,1), names.arg=names(rhi_vals), cex.names=0.6, las=2, border=NA, col=cols)
legend(x="topright", legend=names(legend), fill=legend, col=legend)

# Add error bars signifying the 5% and 95% quantiles
segments(bars, vals5, bars, vals95, lwd=1.5)
arrows(bars, vals5, bars, vals95, lwd=1.5, angle=90, code=3, length=0.05)

### Plot by topology colour but in decreasing homoplasy

# Order the matrix by the median RHI value so the 5 and 95% quantiles are ordered accordingly

cols <- c(jarv, prum, kuhl, wu, still, jarv, prum, still, jarv, prum, still, jarv, prum, jarv, prum, still, other, jarv, prum, other, other, other)
rhis <- rbind(rhi_vals, vals5, vals95, cols)
class(rhis)
dim(rhis)

rhis_df <- data.frame(t(rhis))
sorted_rhis <- rhis_df[order(rhis_df$rhi_vals, decreasing=TRUE),]

pdf("Pectoral_homoplasy_barplot.pdf")
bars <- barplot(as.numeric(sorted_rhis$rhi_vals), ylab="Relative Homoplasy", ylim=c(0,1), names.arg=rownames(sorted_rhis), cex.names=0.6, las=2, border=NA, col=sorted_rhis$cols)
legend(x="topright", legend=names(legend), fill=legend, col=legend)
segments(bars, as.numeric(sorted_rhis$vals5), bars, as.numeric(sorted_rhis$vals95), lwd=1.5)
arrows(bars, as.numeric(sorted_rhis$vals5), bars, as.numeric(sorted_rhis$vals95), lwd=1.5, angle=90, code=3, length=0.05)
dev.off()

###

# Test RHI score for randomised pectoral dataset

pec_r <- apply(pec, 2, sample, replace=FALSE)
rownames(pec_r) <- row.names(pec)

statef(pec_r)
pec_r <- remove.invar(pec_r)
pec_r_phydat <- morph.phydat(pec_r)

RHI(pec_r_phydat, prum, 1000)

#5% quantile          RHI 95% quantile  Tree Length         Lmin 
#   0.9865330    0.9974668    1.0086617 3395.0000000  245.0000000

RHI(pec_r_phydat, jarv, 1000) #0.9952487

RHI(pec_r_phydat, still, 1000) #0.9939835

RHI(pec_r_phydat, wu, 1000) #0.9961989

RHI(pec_r_phydat, kuhl, 1000) #0.9931843
```
